# Mouse Adaptation of Human Inflammatory Bowel Diseases Microbiota Enhances Colonization Efficiency and Alters Microbiome Aggressiveness Depending on Recipient Colonic Inflammatory Environment

**DOI:** 10.1101/2024.01.23.576862

**Authors:** Simon M. Gray, Anh D. Moss, Jeremy W. Herzog, Saori Kashiwagi, Bo Liu, Jacqueline B. Young, Shan Sun, Aadra Bhatt, Anthony A. Fodor, R. Balfour Sartor

## Abstract

Understanding the cause vs consequence relationship of gut inflammation and microbial dysbiosis in inflammatory bowel diseases (IBD) requires a reproducible mouse model of human-microbiota-driven experimental colitis. Our study demonstrated that human fecal microbiota transplant (FMT) transfer efficiency is an underappreciated source of experimental variability in human microbiota associated (HMA) mice. Pooled human IBD patient fecal microbiota engrafted germ-free (GF) mice with low amplicon sequence variant (ASV)-level transfer efficiency, resulting in high recipient-to-recipient variation of microbiota composition and colitis severity in HMA *Il-10^-/-^* mice. In contrast, mouse-to-mouse transfer of mouse-adapted human IBD patient microbiota transferred with high efficiency and low compositional variability resulting in highly consistent and reproducible colitis phenotypes in recipient *Il-10^-/-^* mice. Human-to-mouse FMT caused a population bottleneck with reassembly of microbiota composition that was host inflammatory environment specific. Mouse-adaptation in the inflamed *Il-10^-/-^*host reassembled a more aggressive microbiota that induced more severe colitis in serial transplant to *Il-10^-/-^* mice than the distinct microbiota reassembled in non-inflamed WT hosts. Our findings support a model of IBD pathogenesis in which host inflammation promotes aggressive resident bacteria, which further drives a feed-forward process of dysbiosis exacerbated gut inflammation. This model implies that effective management of IBD requires treating both the dysregulated host immune response and aggressive inflammation-driven microbiota. We propose that our mouse-adapted human microbiota model is an optimized, reproducible, and rigorous system to study human microbiome-driven disease phenotypes, which may be generalized to mouse models of other human microbiota-modulated diseases, including metabolic syndrome/obesity, diabetes, autoimmune diseases, and cancer.

## Introduction

Human inflammatory bowel diseases (IBD) are heterogeneous chronic inflammatory conditions driven by microbial activation of dysregulated intestinal immune responses in genetically susceptible hosts^1^. Host genetic susceptibility loci, such as polymorphisms in *Nod2*, *Il23r*, *Il-10r*, and *Il-10*, explain <20% of IBD variance^2-4^ and disease incidence is rising globally^5^, suggesting that environmental factors (diet, microbiome) are important drivers of IBD. IBD patients have altered intestinal microbiota composition (dysbiosis), functionally characterized by reduced diversity, unstable community structure over time and following perturbation, and expanded aggressive (*Gammaproteobacteria, Enterococcaceae*, sulfur-reducing bacteria) but reduced beneficial (short-chain fatty acid [SCFA]- producing *Clostridiales*, *Blautia*) resident bacteria^6-10^. Viable microbes are required to develop chronic T-cell mediated intestinal inflammation in most experimental colitis models (i.e. *Il-10^-/-^, Il2^-/-^, Tcrab ^-/-^,* Naïve CD4^+^ T cell transfer to *Rag1/2^-/-^*, *Tlr5^-/-^, Tnf^τ.ARE^* mice) in which GF mice have no inflammation but develop progressive intestinal inflammation after colonization with complex microbiota^11-16^. Aggressive resident bacteria (pathobionts) within the complex gut microbiota are the key drivers of intestinal inflammation^17-22^; however, whether dysbiotic expansion of pathobionts is a cause or consequence of intestinal inflammation and how the host environment shapes microbial ecology in IBD remain poorly understood.

Colonization of GF animals with defined human bacterial consortia or human fecal microbiota transplant (FMT) are the gold-standard methods to demonstrate causality and investigate mechanisms of human microbiome-driven disease phenotypes^23-29^. Defined consortia enable strict control of microbiota composition, which facilitates mechanistic studies using genetically modified consortium members but requires selection of bacterial strains by variable criteria^28, 30-32^. Strain-level genetic and functional variation are human disease-state specific, strongly impact host-microbe interaction, and alter disease severity in experimental colitis models^22, 33-38^. Because defined consortia may omit strain-specific genetic and functional attributes responsible for human disease phenotypes, direct transplant of human disease-associated feces to GF rodents is an appealing method to study human microbiome-driven diseases.

Human IBD patient FMT to colitis-prone, GF mice (*Il-10^-/-^* and *Rag1^-/-^* T-cell transfer models) transfers enhanced colitis severity compared to healthy patient FMT and induces a T_H_17- and T_H_2- dominant immune phenotype that is characteristic of human IBD^26, 39-42^. These fecal transplant studies clearly transfer disease phenotype to susceptible mice by human IBD-associated microbes. Importantly, human-to-mouse fecal transplant causes a microbial population bottleneck that engrafts a compositionally distinct microbiome in recipient mice compared to human donor stool, likely due to low human-to-mouse strain-level transfer efficiency (∼40%) and host-specific microbe preferences^43-46^. We took advantage of the microbiota reassembly associated with human-to-mouse FMT to ask if 1) the host environment controls microbiota assembly and inflammatory potential, and 2) mouse-adaptation of human fecal microbiota forms a microbial community that is stable in serial transplant to GF mice and leads to more reproducible experimental phenotypes.

To evaluate the impact of the host inflammatory environment on gut microbiota assembly we transferred pooled feces from human IBD patients with active disease to wild-type (WT) or *Il-10^-/-^* mice. Human microbiota-associated (HMA) *Il-10^-/-^* mice had lower microbial alpha diversity, higher compositional variability, and expansion of pathobionts compared to HMA WT mice, illustrating the influence of an inflammatory colonic environment on dysbiosis. Serial transfer of non-inflamed (WT) mouse-adapted human microbiota to GF *Il-10^-/-^* mice induced less severe colitis than inflamed (*Il-10^-/-^*) mouse-adapted human microbiota. Transplant of human fecal microbiota to GF mice resulted in low human-to-mouse transfer efficiency at the strain level, while mouse-adapted human microbiota yielded high strain level transfer efficiency. High microbiota compositional variability in HMA *Il-10^-/-^* mice was associated with variable colitis severity, but recipient mice colonized with mouse-adapted human microbiota exhibited low compositional variability and more consistent colitis phenotypes. Our findings suggest that the reproducibility and rigor of HMA animal studies are impacted by the variability of human-to-mouse FMT; however, experimental design can be improved by first adapting the human microbiota to the mouse host followed by transfer of mouse-adapted human microbiota for subsequent highly reproducible mechanistic studies.

## Methods

### Mouse Lines

GF 129S6/SvEv background wildtype (WT) and *Il-10*-deficient (*Il-10^-/-^*) mice^13^ were obtained from the National Gnotobiotic Rodent Resource Center (NGRRC) at the University of North Carolina at Chapel Hill. All animal experiments were conducted under approved Institutional Animal Care and Use Committee protocols.

### Human fecal samples

Human fecal samples from 5 adult patients with active Crohn’s disease (CD) (4 donors) or ulcerative colitis (UC) (1 donor) without prior intestinal surgery or antibiotic exposure within 3 months were collected under an Institutional Review Board approved protocol (Fig S1A). De-identified stool samples were aliquoted immediately after collection in an anaerobic chamber and stored without preservatives at -80°C until use.

### Human fecal microbiota and mouse-adapted fecal microbiota colonization of GF mice

Human fecal material from two sets of 3 human donors with active IBD (HM1: Donors 1, 2, 3; HM2: Donors 3, 4, 5) was thawed and pooled in equal proportions by weight under anaerobic conditions (N_2_:H_2_:CO_2_ = 80:10:10), diluted with anaerobically reduced phosphate-buffered saline (PBS) to generate a fecal slurry, and administered by 150μl oral gavage to recipient GF 129 WT or 129 *Il-10^-/-^* mice at 2mg pooled human donor stool per mouse. Mouse-adapted fecal pellets from HMA 129 WT or 129 *Il-10^-/-^* mice were freshly collected and pooled daily between 14- and 21-days post-colonization and frozen at -80°C without preservatives. To generate a standardized slurry of mouse-adapted microbiota, mass collected fecal pellets from all mice in a group were pooled, homogenized, and diluted to 100mg/ml under anaerobic conditions in sterile anaerobically reduced lysogeny broth (LB) with 20% glycerol. Solid particulate matter was pelleted by brief slow centrifugation and slurry supernatant was aliquoted to cryovials for storage at -80°C. Slurry supernatant contains the same microbial community composition as whole fecal material^43, 47^. Mouse-adapted microbiota slurry generated as above from fecal pellets of HMA 129 WT mice is called non-inflamed mouse adapted microbiota (NIMM), while slurry generated from fecal pellets of HMA colitis-prone 129 *Il-10^-/-^*mice is called inflamed mouse adapted microbiota (IMM) (Fig 1A). To colonize GF mice with mouse-adapted microbiota, standardized aliquots of 100mg/ml fecal slurry were thawed under anaerobic conditions, diluted with anaerobically reduced PBS, and administered by oral gavage to recipient GF 129 WT or 129 *Il-10^-/-^*mice at 2mg per mouse in 150μl. Fecal pellets from IMM or NIMM associated 129 WT or 129 *Il-10^-/-^* mice were collected daily between 14- and 21-days post-colonization when the *Il-10^-/-^* recipient microbiota has stabilized and before cage effects are reported to develop^48, 49^, processed and frozen in aliquots as above to generate standardized slurries of serial passages (-g1, -g2, and -g3) of mouse-adapted microbiota (Fig 1A). All experiments were performed using aliquots from a single production batch of mouse-adapted microbiota. All mouse fecal transplant experiments were performed in BSL-2 isolation cubicles with HEPA-filtered air on a 12-hour dark/light cycle with ad libitum access to autoclaved water and mouse chow (Purina Advanced Protocol Select Rodent 50 IF/6F Auto Diet) using the sterile out-of-isolator gnotobiotic cage technique (Complete cage GM500, Green Line, Tecniplast)^50^. Cage changes and all animal handling were performed in a laminar flow biosafety cabinet under sterile technique following ultraviolet light treatment and 10-minute Peroxigard sterilization of all equipment and surfaces. We maintained strict GF conditions with the out-of-isolator gnotobiotic technique for at least 2 weeks. We consider the complex microbiota fecal transplant experiments reported here to be ‘near-gnotobiotic’ with low risk of environmental contamination, but not strictly gnotobiotic since they are performed with out-of-isolator gnotobiotic cage technique for durations >2 weeks and sterility could not be monitored due to complex microbiota transplants.

**Figure 1.**
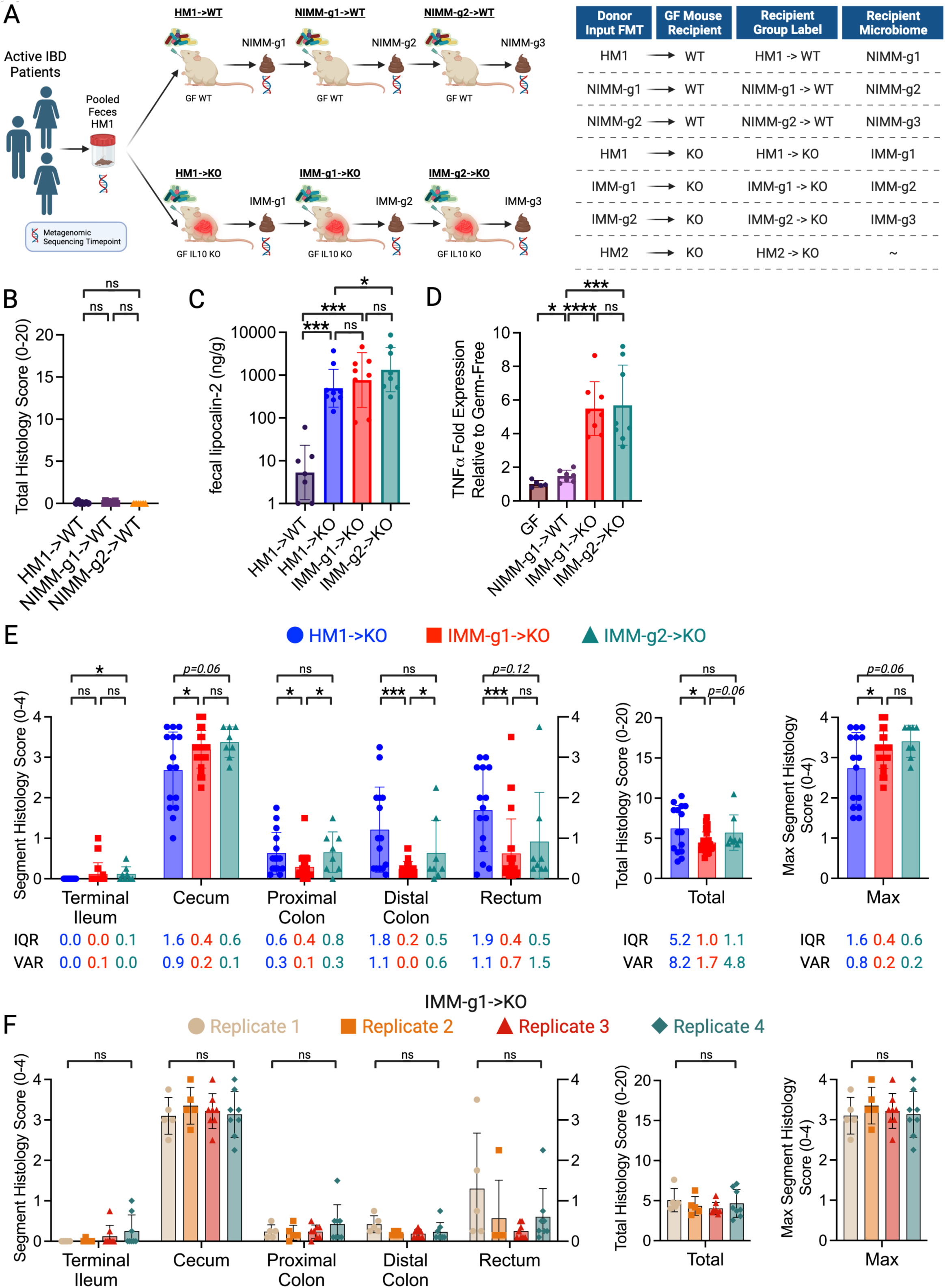
Mouse-adapted human microbiota induces more consistent and reproducible colitis than directly transplanted human microbiota. A) Experimental design. Pooled feces from 3 humans with active IBD (2 CD, 1UC) were transplanted to non-inflamed WT or colitis-susceptible *Il-10*^-/-^ (IL-10KO, KO) GF recipient mice. Mouse-adapted microbiotas were serial transplanted to non-inflamed WT or colitis-susceptible *Il-10*^-/-^ GF recipient mice. B) Total colon and ileum histology score for WT mice at day 28 post-colonization. C) f-LCN2 level at day 28 post-colonization. D) TNFα mRNA levels in cecal tissue at day 28 post-colonization. E) Segment, total colon and ileum, and max segment histology score for *Il-10^-/-^* mice at day 28 post-colonization. F) Segment, total colon and ileum, and max segment histology score for IMM-g1 colonized *Il-10^-/-^* mice at day 28 post-colonization from 4 independent experiments. Data shown are representative of (C-D) or cumulative (B, E-F) from 2-4 independent experiments. n=7-9 (B-D), n=15-26 (E), n=5-8 (F) mice per group. Data are expressed as mean±SD or geometric mean ± geometric SD (C). Statistical significance calculated by unpaired t-test or Mann-Whitney test (C) with *p<0.05, **p<0.01, ***p<0.001.

### Gene expression by qRT-PCR, Intestine histopathology score, and Fecal lipocalin-2 quantification

Standard molecular assays and histopathology scoring were performed as previously described^51-53^. Details of these procedures and a list of qPCR primers are found in the Supplemental Experimental Procedures.

### Statistical Analyses

Non-sequencing based statistical analyses were performed with Prism 10 (GraphPad) with statistical tests and significance thresholds indicated in figure legends.

### 16S rRNA Amplicon Metagenomic Sequencing and Analysis

16S rRNA amplicon (variable regions 3-4) sequencing was performed on the Illumina NextSeq 2000 platform, processed, and taxonomically classified through QIIME2 by the UNC Microbiome Core^54^. Additional details of these procedures are found in the Supplemental Experimental Procedures. Sequence counts data at both the genus and phylum level were extracted from the respective QIIME2 artifact files. The amplicon sequence variant (ASV)-level counts table was generated with forward reads using the following parameters with single-end DADA2 on the QIIME2 (version 2021.2) platform: the first 10 base pairs of each sequence were trimmed, and the sequences were truncated to 180 base pairs as determined by sequence quality using FastQC (version 0.11.9)^54, 55^. Statistical analysis was conducted with the *vegan* package (ver.2.6-2) in R (ver. 4.2.2) and visualized with the Shiny application *Plotmicrobiome* and custom R code (Sun et al. GitHub https://github.com/ssun6/plotmicrobiome, Supplemental File 1). To ensure reproducibility and rigor, the results of our analyses were independently reproduced with custom Python code by a second bioinformatician (JBY) with replicated key figures and reproducible tested code available in a Jupyter Notebook file (Supplemental File 1). R and Python code used in our analyses are available at https://github.com/anhmoss/Mouse-Adaptation-of-Human-Inflammatory-Bowel-Disease-Microbiota-Enhances-Colonization-Efficiency and in Supplemental File 1. 16S rRNA amplicon sequencing data are available at https://github.com/anhmoss/Mouse-Adaptation-of-Human-Inflammatory-Bowel-Disease-Microbiota-Enhances-Colonization-Efficiency.

To account for varying sequencing depth, all counts data were normalized according to the following formula prior to downstream statistical analyses:

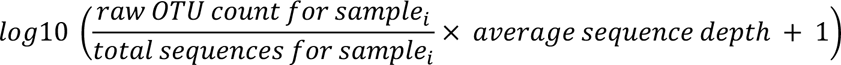

This formula adjusts the pseudo-count to have a similar effect across samples by scaling all samples to the average sequencing depth. ASV transfer efficiency was measured as Pearson correlation coefficient (r) for pairs of samples within a given group or between two groups.

## Results

### Mouse-adapted human microbiota induces more consistent and reproducible colitis than directly transplanted human microbiota

Human fecal microbiota transplantation into GF mice can transfer microbe-dependent pathological phenotypes to recipient animals, allowing investigation of microbial mechanisms of human diseases such as IBD^23, 26, 56^. The large interpersonal variation of human gut microbiota, host-specificity of gut microbial ecology, and variable engraftment of human gut microbes into GF mice pose challenges to transplanted phenotype reproducibility and interpretation^43, 45, 46, 57^. To understand the impact of recipient host environment on human fecal microbiota engraftment and phenotype transfer in a mouse model of experimental colitis, we transplanted pooled feces from 3 humans with active IBD (2 CD, 1UC) to non-inflamed WT or colitis-susceptible *Il-10*^-/-^ GF mice (Fig 1A, S1A). We then transplanted these mouse-adapted microbiota to sequential cohorts of non-inflamed WT or colitis-susceptible *Il-10*^-/-^ GF recipient mice, generating serial transfers of mouse-adapted human microbiota identified as -g1, -g2, and -g3 (Fig 1A). In our nomenclature, different human IBD patient fecal pools are called Human Microbiota (HM1 or HM2) (shown in S1A), feces from HMA WT mice are called Non-Inflamed Mouse-adapted Microbiota (NIMM), and feces from HMA *Il-10*^-/-^ mice are called Inflamed Mouse-adapted Microbiota (IMM) (Fig 1A). Serial mouse-adapted fecal transplant experiments were only conducted with HM1-dervied HMA mouse stool due to resource constraints; HM1 was selected because the cohort contained both UC and CD donors (Fig 1A; S1A). Because colonic immune stimulation of GF mice is equivalent following transplant of human or mouse microbiota, HMA mice are a clinically relevant model of experimental colitis^44, 45^.

GF 129 WT mice receiving HM1, NIMM-g1, or NIMM-g2 fecal transplant did not develop colitis as assessed by colon histology, non-invasive fecal lipocalin-2 (f-LCN2), and tissue inflammatory cytokine levels (Fig 1B-D; S1B-C). Transplantation of both human microbiota HM1 and mouse-adapted microbiota IMM-g1 or IMM-g2 to GF 129 *Il-10^-/-^* mice induced severe colitis as assessed by colon histology, non-invasive f-LCN2, and inflammatory cytokine levels (Fig 1C-F; S1B,D). IMM-g1 and IMM-g2 induced cecal predominant colitis that was equivalent in severity and kinetics to colitis induced by HM1 (Fig 1E; S1B,D). However, HM1 induced colitis was more variable than IMM-g1 or IMM-g2 induced colitis as quantified by segment and total histology score variance and interquartile range (Fig 1E; Fig S1G). The high phenotypic variance of human microbiome-induced colitis was replicated by a separate cohort (HM2) of pooled feces from 3 humans with active CD transplanted to *Il-10*^-/-^ GF mice (Fig S1E,F). In contrast to the highly variable phenotype of human microbiome-induced colitis, mouse-adapted microbiome IMM-g1 induced colitis had little variation in severity or distribution within or across independent experiments (Fig 1E,F; S1G,H). To evaluate whether variability in colitis phenotype was related to microbiome composition, we performed 16S amplicon sequencing of input donor microbiota and fecal samples collected from ex-GF 129 WT and 129 *Il-10^-/-^* mice colonized for 28 days with human microbiota or mouse-adapted microbiotas (Fig 1A). As we show later in the results (Fig 3), human microbiota transplant to GF mice was associated with significantly lower microbiota engraftment consistency than mouse-adapted microbiota transplant, suggesting that variability in engrafted human microbiota composition may cause variability in colitis phenotypes.

### Human microbiome restructures with transplant to GF mice

To investigate how the recipient host intestinal environment shapes human microbiota engraftment in GF mice, we assessed microbiome compositional variation by calculating the average relative abundance of genera across all samples for each fecal transplant condition. Fig 2 shows taxonomic barplots of the 8 most abundant genera across groups with the remaining lower abundance taxa grouped as “Other” (Fig 2, S2A). The 30 most abundant genera across groups and the relative abundance of genera for individual mice are visualized in barplots in Fig S2. We performed pairwise t-tests to assess differential abundance between groups, excluding genera present in less than 10% of the samples (Table S1).

**Figure 2.**
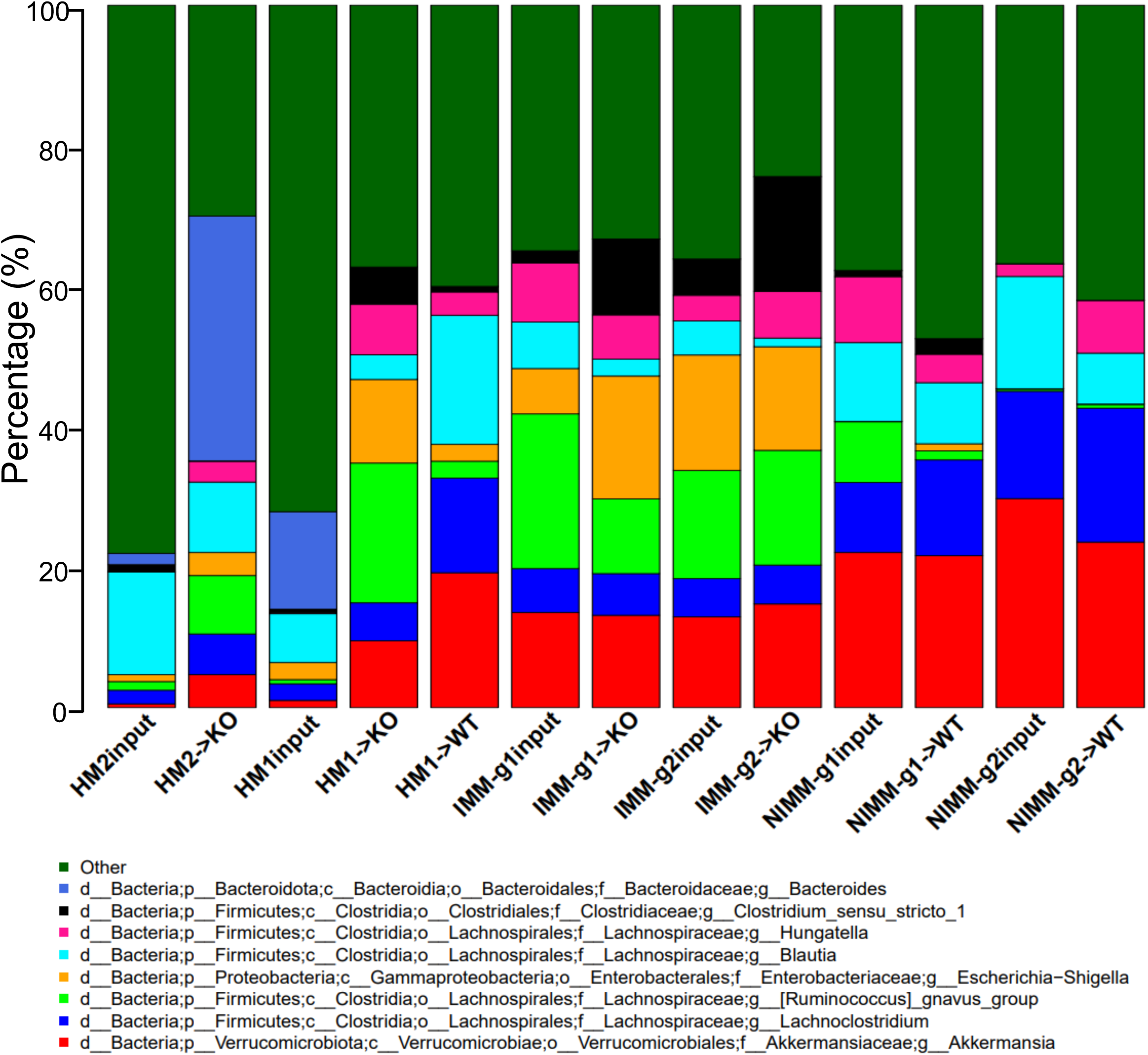
Recipient host environment influences engraftment composition of human-microbiome associated mice. A) 16S Seq taxonomic bar plots show top 8 most abundant genera in FMT inputs and recipient mouse feces at day 28 post-colonization. For mouse recipient groups, bar plots are average of 16S seq data from n=7-18 mice/group.

Pooled human microbiome composition (HM1 input and HM2 input) was compositionally distinct from all colonized mouse groups as visualized by taxonomic barplots and principal coordinates analysis (PCoA) clustering (aka multidimensional scaling), with the strongest separation existing between human and mouse-adapted microbiotas along the first MDS axis (Fig 2, 3A). Compared to human microbiomes, HMA mouse and MA-FMT mouse microbiomes had increased relative abundance of *Akkermansia*, *Lachnoclostridium*, *Ruminococcus gnavus* group, and *Hungatella*, a low-abundance member of the human gut that was not detectable by 16S in HM1 or HM2 inputs (Fig 2). *Bacteroides*, a major constituent of the human gut microbiome, was present in HM1 input and HM2 input, and expanded in HM2-associated *Il-10^-/-^* mice but reduced in HM1-associated WT and *Il-10^-/-^* mice (Fig 2, Table S1). The expansion of *Bacteroides* in HM2-but reduction in HM1-associated *Il-10^-/-^* mice was surprising because *Bacteroides* was more abundant in HM1 input compared to HM2 input, suggesting stochastic factors influence engraftment of human microbiota in GF mice (Fig 2). These data demonstrate that human microbiota association of GF mice results in major compositional restructuring of the engrafted microbiome that may be partially stochastic.

**Figure 3.**
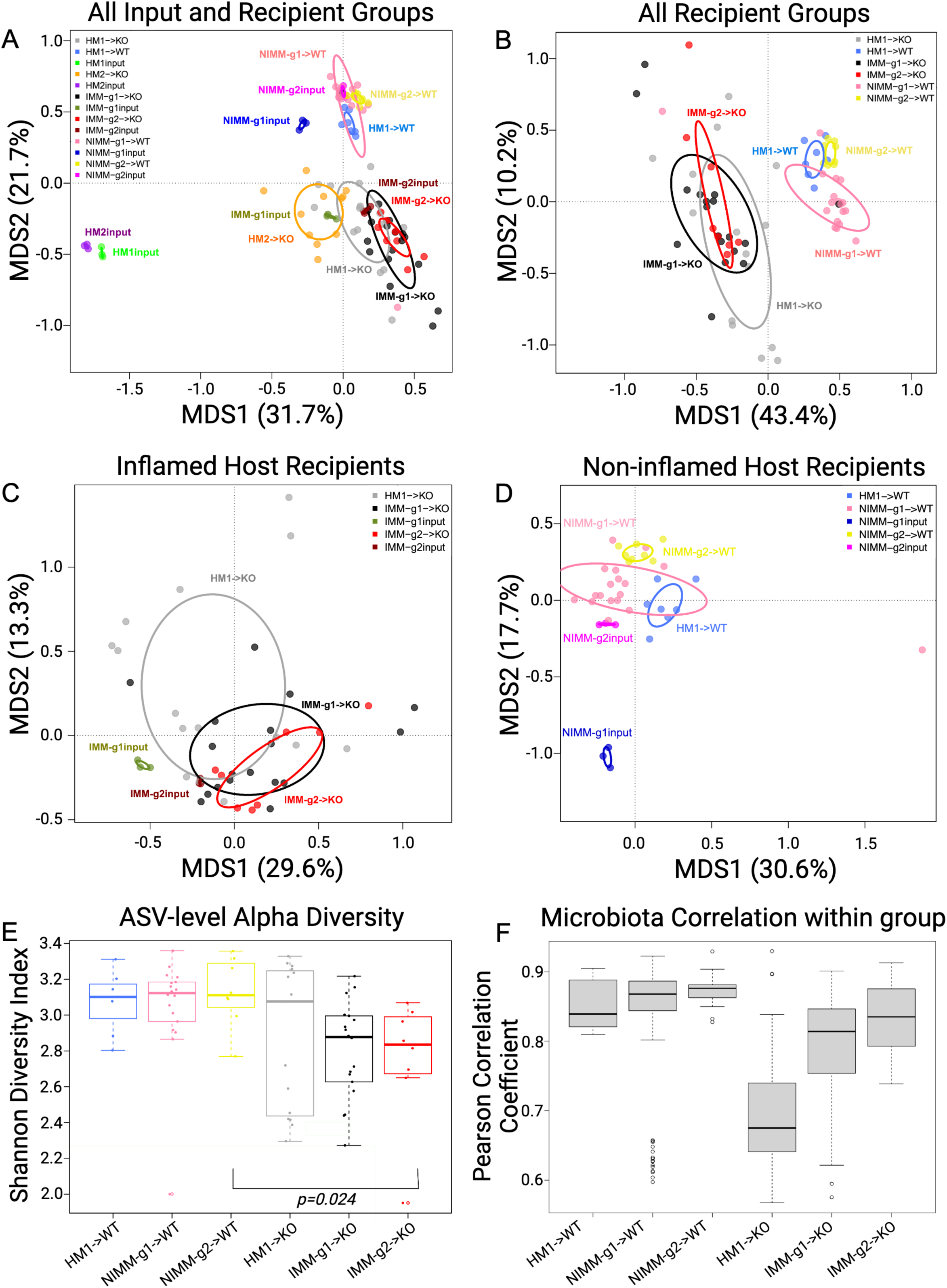
Human microbiome restructuring with transplant to GF mice is host inflammatory environment specific. A) Principal coordinates analysis, PCoA, of 16S Seq data for human and mouse-adapted FMT inputs and FMT recipient WT and KO mouse groups. B) PCoA of FMT recipient WT and KO mouse groups. C) PCoA of FMT recipient KO mouse groups. D) PCoA of FMT recipient WT mouse groups. E) Shannon index at ASV level for FMT recipient WT and KO mouse groups. F) Pearson correlation coefficients (r) within group for FMT recipient WT and KO mouse groups quantify variability of microbiota composition between mice in the same group (microbiota engraftment consistency). Dots in PCoA plots represent individual mice for FMT recipient WT and *Il-10^-/-^* (KO) mouse groups. For FMT inputs, a single input slurry was used in each experiment and input dots represent sequencing data from three 16S amplicon PCR technical replicates. Analysis conclusions did not change when using average input vs individual technical replicates, so technical replicates are displayed to demonstrate the high consistency of 16S amplicon PCR in our dataset.

### Recipient host environment influences engraftment composition of human-microbiota associated mice

The recipient host environment shapes the engrafted microbiome composition of HMA mice (Fig 2, S2, 3A, B). After removing HM1 and HM2 inputs, PCoA showed separation of inflamed mouse-adapted microbiota (IMM) and non-inflamed mouse-adapted microbiota (NIMM) along the first MDS axis (Fig 3B). PERMANOVA test with all mouse recipient groups as the model term demonstrated that approximately 43% of the variation in the data is explained by the recipient host environment (coefficient of determination, R^2^ = 0.43, p=0.001). Mouse adaptation in the inflamed *Il-10^-/^*^-^ host (IMM) enriched for significantly higher relative abundance of *Escherichia-Shigella, Parasutterella*, *Enterococcus, Clostridium_sensu_stricto_1*, *Ruminococcus gnavus group,* and *Bifidobacterium* but significantly lower relative abundance of *Clostridium innocuum, Blautia*, *Lachnoclostridium,* and multiple other genera within *Lachnospiraceae* and *Ruminococaceae* when compared to mouse adaptation in the non-inflamed WT host (NIMM) (Fig 2, S2, Table S1). PCoA of serial microbiota passage within the non-inflamed WT host environment (NIMM-g1, -g2) showed that the global microbiome structure remained stable with no distinct clustering of groups (Fig 3D) and only 7 operational taxonomic units (OTUs) demonstrated significantly differential abundance between NIMM-g1 and -g2 using a cutoff of FDR<0.1 (Fig 2, S2, Table S1). PCoA of serial microbiota passage within the inflamed *Il-10^-/-^* host environment (IMM-g1, -g2) similarly showed globally stable microbiome structure with no distinct clustering of groups (Fig 3C, S3A), while no OTUs were differentially abundant between IMM-g1 and -g2 (Fig 2, S2, Table S1). Mouse adaptation in the inflamed *Il-10^-/-^* host (IMM) was associated with significantly lower alpha diversity at the amplicon sequence variant (ASV) level compared to the non-inflamed WT host (NIMM), consistent with observations that human IBD patients have lower alpha diversity than healthy humans (Fig 3E, S3B,C)^8^. Together, these data demonstrate that the composition of the human microbiome is fundamentally restructured with transplant to GF mice and that the recipient host environment strongly shapes the relative abundance of engrafted strains with the inflamed *Il-10^-/-^*host (IMM) driving a dysbiotic microbiome defined by lower alpha-diversity, enrichment of pathobionts, and reduction of protective SCFA-producing bacteria relative to the non-inflamed WT host (NIMM).

### Human microbiota engrafts with variable composition compared to more consistent engraftment by mouse-adapted microbiota

Since the human microbiome restructures with transplant to GF mice, we speculated that variability in engrafted microbiota composition may explain the colitis phenotype variability of HMA *Il-10^-/-^*mice (Fig 1E, S1E-F). Variability of microbiota composition was quantified by pairwise calculation of Pearson correlation coefficient for all samples within the same group (i.e., all mice within HM1->KO). A high Pearson correlation coefficient indicates compositional similarity between samples in a group, while a low coefficient indicates compositional variability between samples in a group. Human microbiota transplant to 129 *Il-10^-/-^* mice (HM1->KO) was associated with significantly lower Pearson correlation coefficients than mouse-adapted microbiota transplants to 129 *Il-10^-/-^*mice (IMM-g1->KO, IMM-g2->KO) (Fig 3F, S3D). A similar trend was seen with human microbiota or mouse-adapted microbiota transplant to 129 WT mice (Fig 3F). Pearson correlation coefficients for 129 *Il-10^-/-^*recipient mice were consistently lower than 129 WT recipients at each stage of serial passage (i.e., HM1->WT vs HM1->KO or NIMM-g1->WT vs IMM-g1->KO), demonstrating that inflammation promotes variability of microbiome composition while health is associated with microbiome stability (Fig 3F). These results are consistent with observations in humans that the composition of IBD patient microbiomes fluctuate more than healthy controls over time^28, 58^. Together, these data suggest that 1) inflammation promotes microbiome variability and 2) variability in colitis phenotype with human microbiota transplant may be due to variability in engrafted human microbiota composition, while the more consistent colitis induced by mouse-adapted microbiota may be due to homogeneity of engraftment of mouse-adapted microbiota.

### Mouse-adapted human IBD microbiota transfers with higher efficiency than human fecal transplant

Since HMA mice had significantly different microbiome composition than human donor stool but mouse-adapted FMT mice had highly consistent microbiomes between serial transfer, we evaluated whether mouse-adapted microbiota transfers to GF mice more efficiently than human fecal transplant (Fig 4A-D). To quantify transfer efficiency, we detected all ASVs across all samples and compared ASV abundance between human stool, HMA mice, and mouse-adapted FMT mice across serial transfers (Fig 4A-D). We visualized these data using scatter plots where each dot represents a unique ASV plotted by log_10_ relative abundance in the input microbiome (x-axis) vs recipient mouse microbiome (y-axis). We quantified transfer efficiency using Pearson correlation coefficient (r), where high Pearson r indicates consistent ASV abundances between samples and high transfer efficiency. We used deep 16S amplicon sequencing rather than whole genomic shot gun sequencing (WGS) because repeat sequencing of the same region allows for exact identification of ASVs in a database-independent manner without reliance on classification algorithms.

**Figure 4.**
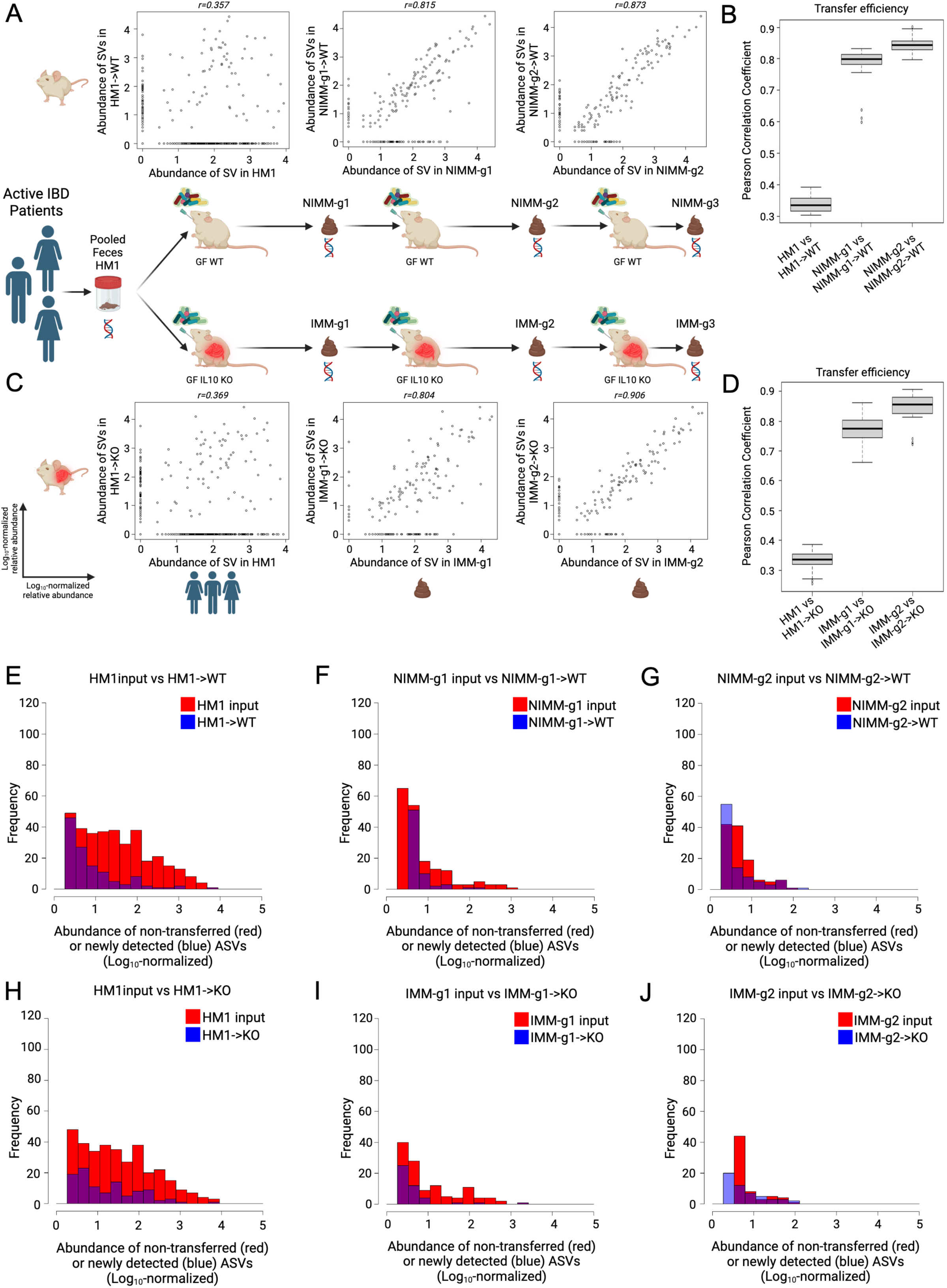
Mouse-adapted human IBD microbiota transfers with higher efficiency than human fecal transplant. A) ASV level log_10_-normalized relative abundance correlations for FMT input and WT recipient mice where each dot represents a unique ASV plotted in the input microbiome (x-axis) vs recipient mouse microbiome (y-axis). B) Transfer efficiency quantified by Pearson correlation coefficient (r) between FMT input and WT recipient mouse groups at the ASV level. C) ASV level log_10_-normalized relative abundance correlations for FMT input and KO recipient mice. D) Transfer efficiency quantified by Pearson correlation coefficient (r) between FMT input and KO recipient mouse groups at the ASV level. E-J) Representative histograms of non-transferring ASVs (red, representing y=0 ASVs in above dot plots) and newly detected *in vivo* ASVs (blue, representing x=0 ASVs in above dot plots) binned by log_10_-normalized relative abundance for (E) HM1->WT, (F) NIMM-g1->WT, (G) NIMM-g2->WT, (H) HM1->KO, (I) IMM-g1->KO, and (J) IMM-g2->KO FMT recipient mouse groups.

Human fecal transplant to WT or *Il-10^-/-^* mice was associated with low transfer efficiency and poor transfer of relative composition to recipient mice (Fig 4A-D, S5D-E). Very similar results were seen in WT and *Il-10^-/-^*mice. A large proportion of ASVs present in human stool did not transfer to recipient mice, which is illustrated by ASVs falling on the x-axis (Fig 4A,C). The relative abundance (log_10_ normalized) of non-transferring ASVs demonstrated that even moderately to highly abundant ASVs in human stool did not transfer efficiently to GF mice (Fig 4E,H). ASV relative abundance in human stool had little correlation with relative abundance in recipient mice (Fig 4A,C), leading to a very low ASV level transfer efficiency for human fecal transplant to WT (r=0.34±0.03) or *Il-10^-/-^* (r=0.33±0.03) mice (Fig 4B,D). For mouse-adapted FMT, however, ASV relative abundance in MA-FMT input (IMM or NIMM) was highly correlated with relative abundance in recipient mice (Fig 4A,C). Only a small proportion of ASVs present in mouse-adapted microbiota did not transfer to recipient mice, and those non-transferring ASVs were primarily low-abundance strains (Fig 4A,C,F-G,I-J). Serial transfer of mouse-adapted microbiota further improved the correlation between input and recipient microbiomes and reduced non-transferring ASV numbers, leading to very high ASV level transfer efficiency for mouse-adapted FMT to WT (r=0.84±0.02) or *Il-10^-/-^* mice (r=0.85±0.05) (Fig 4 A-D). Similar results were found when comparing human microbiome input to mouse-adapted microbiome inputs (Fig S4A-F). Analysis of transfer efficiency at the genus level also demonstrated low transfer efficiency for human fecal transplant but high transfer efficiency for mouse-adapted FMT; however, phylum level analysis showed high transfer efficiency for all conditions, giving a misleading perception of transfer efficiency (Fig 5A-H, S5F).

**Figure 5.**
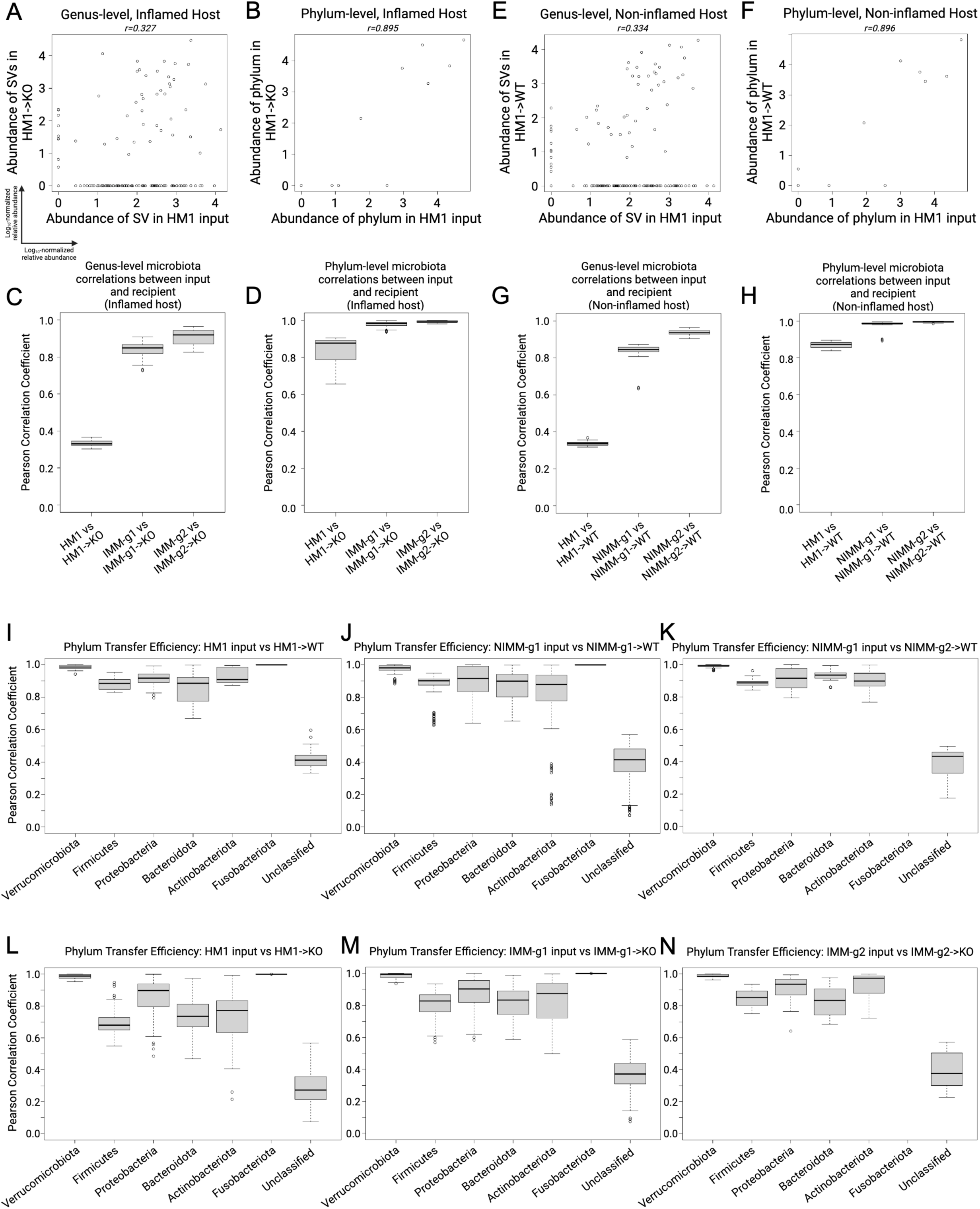
Transfer efficiency varies between taxa. A) Genus-level and B) phylum-level log_10_- normalized relative abundance correlations comparing HM1 input to HM1->KO, C-D) Pearson correlation coefficient (r) between input and inflamed (KO) recipient at the (C) genus- and (D) phylum-level. E) Genus-level and F) phylum-level log_10_-normalized relative abundance correlations comparing HM1 input to HM1->WT, G-H) Pearson correlation coefficient (r) between input and non-inflamed (WT) recipient at the (G) genus- and (H) phylum-level. I-K) Pearson correlation coefficient (r) between input and non-inflamed (WT) recipients by phylum. L-N) Pearson correlation coefficient (r) between input and inflamed (KO) recipients by phylum.

Some ASVs (falling on the y-axis) in HMA mice were not detected in human stool, representing either mutation of the V3-V4 sequence, *in vivo* expansion of very low abundance strains undetected at the depth of 16S sequencing utilized, or environmental contamination (Fig 4A,C). To rule out environmental contamination, we performed human FMT to GF mice in strictly gnotobiotic isolators and still detected many ASVs in HMA mice that were not detected in human input stool by 16S Seq (Fig S5D-E). We analyzed an independently published 16S Seq dataset of HMA WT mice colonized and then bred in a gnotobiotic isolator and found similar results of low human-to-mouse but high mouse-adapted-to-offspring mouse transfer efficiency at the ASV level (Fig S5A-C)^44^. These data demonstrate that a large fraction of the human microbiome does not efficiently engraft GF mice; however, once engrafting strains adapt to the mouse gut they transfer with very high efficiency in serial fecal transplant.

### Transfer efficiency varies between taxa

To assess the transfer efficiency of different taxa from transplant of human microbiota or mouse-adapted microbiota to GF mice, we compared Pearson correlation coefficients (r) between phyla (Fig 5I-N). Unclassified bacteria had the lowest transfer efficiency in all groups, consistent with prior reports (Fig 5I-N)^43^. *Verrucomicrobiota* and *Fusobacteriota* consistently had very high transfer efficiency, which likely reflected that a single species from each phylum was present in donor stool (Fig 5I-N). *Akkermansia muciniphila*, a known keystone species, is the only human gut member of *Verrucomicrobiota* and transferred highly efficiently across all transplant conditions and recipients. Transfer efficiencies trended lower for *Firmicutes, Bacteroidota,* and *Actinobacteriota* and trended somewhat lower in *Proteobacteria* in all *Il-10^-/-^* mice compared to WT mice (Fig 5I-N).

### Inflamed mouse-adapted microbiome induces faster onset colitis than non-inflamed mouse adapted microbiome

Since mouse-adaptation of human microbiota in the non-inflamed (WT) host reduced the frequency of pathobionts while expanding putatively protective bacteria, we investigated whether NIMM-g1 induces less severe colitis than IMM-g1 when transplanted to *Il-10*^-/-^ GF mice (Fig 6A). At 14 days post-colonization, NIMM-g1 colonized *Il-10*^-/-^ mice had significantly lower f-LCN2 levels, cecum- and total colon histologic inflammation than IMM-g1 colonized *Il-10*^-/-^ mice (Fig 6B, S6A-B). At 28 days post-colonization, NIMM-g1 colonized *Il-10*^-/-^ mice continued to have significantly reduced cecal inflammation scores and trend toward lower cecal inflammatory cytokine levels but had developed increased rectal inflammation compared to IMM-g1 colonized *Il-10*^-/-^ mice (Fig 6C). NIMM-g1 colonized *Il-10*^-/-^ mice had significantly lower maximum segment inflammation on a per-mouse basis compared to IMM-g1 colonized *Il-10*^-/-^ mice (Fig 6C). However, the increase in rectal inflammation resulted in a non-significant trend toward lower f-LCN2 levels and no difference in total colon histology scores between NIMM-g1 and IMM-g1 colonized *Il-10*^-/-^ mice at 28 days post-colonization (Fig 6C; S6C). PCoA demonstrated that the microbiome of NIMM-g1 colonized *Il-10*^-/-^ mice (NIMM-g1->KO) clustered with WT HMA and MA-FMT mice, rather than *Il-10*^-/-^ HMA or MA-FMT mice (Fig 6E). Alpha diversity of NIMM-g1 colonized *Il-10*^-/-^ mice was equal to NIMM-g1 colonized WT mice and non-significantly higher than IMM-g1 colonized *Il-10*^-/-^ mice (Fig 6G). These data suggest that major changes in community restructuring occur during initial adaptation of human microbiota to the non-inflamed mouse host, but that once a stable mouse-adapted community forms it transfers with stable global structure in serial transplant to subsequently inflamed GF host mice. Although PCoA demonstrated that global microbiome structure of NIMM was stable between the inflamed and non-inflamed environments, taxonomic bar plots and differential abundance analysis demonstrate that several taxa undergo changes in frequency (Fig 6F; S6D; Table S1). Putatively protective *Blautia* and *Lachnospiraceae NK4A136* group were significantly reduced while the pathobiont containing genera *Ruminococcus gnavus* group and *Hungatella* were significantly expanded in *Il-10*^-/-^ compared to WT mice colonized with NIMM-g1, suggesting that these genera may be particularly responsive to the inflammatory environment – consistent with observations in human IBD microbiome profiling studies (Fig 6F, S6E-H)^7, 9, 59, 60^. Together, these data demonstrate that human microbiome adaptation is dependent on the host environment, but once a stable mouse-adapted microbiome has been established it remains remarkably stable in composition despite an altered host environment.

**Figure 6.**
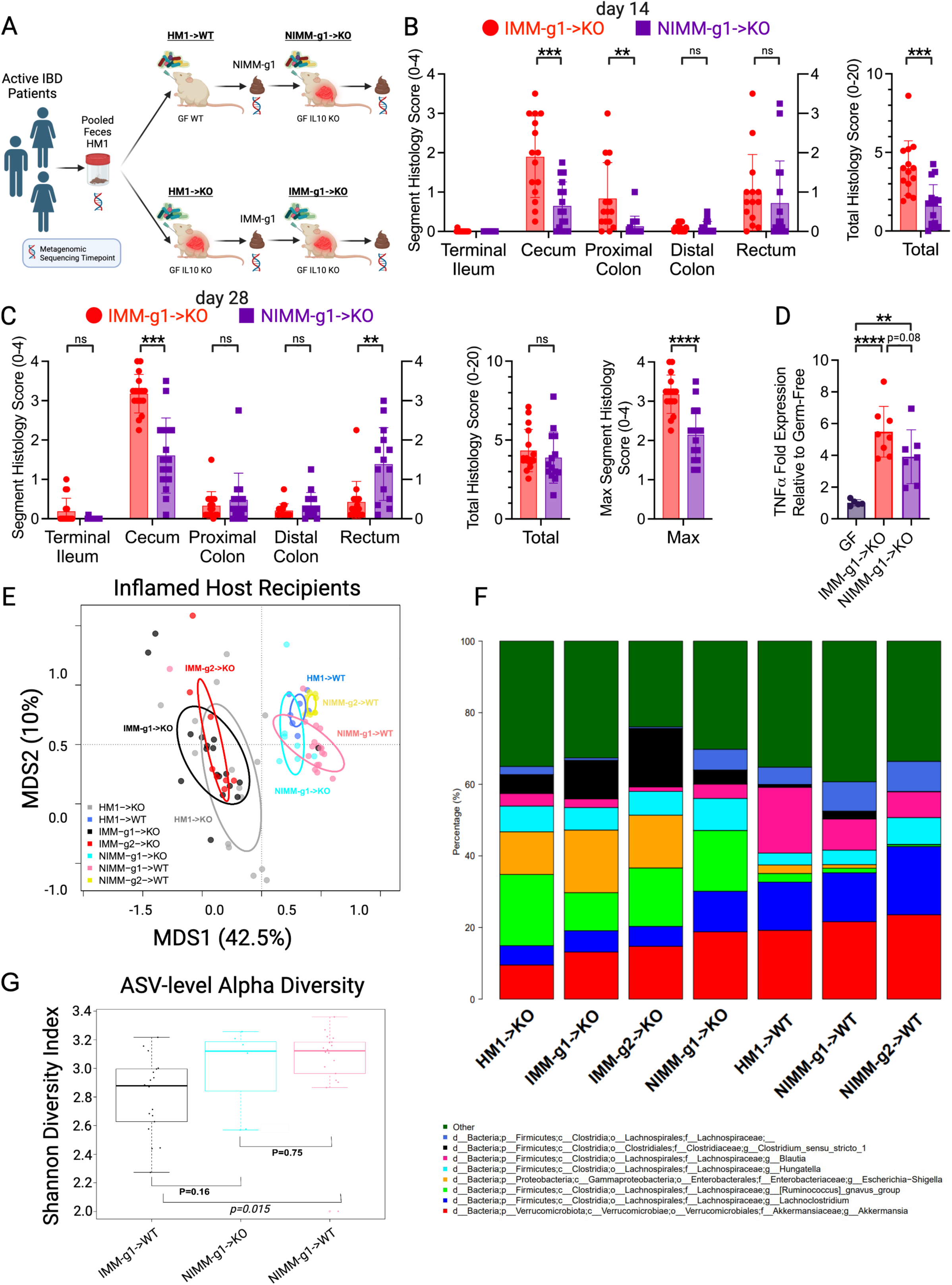
Inflamed mouse-adapted microbiome more rapidly induces severe colitis than non-inflamed mouse adapted microbiome. A). Experimental design. Human IBD patient microbiota (HM1) was adapted in the inflamed (IMM-g1) or non-inflamed (NIMM-g1) host, then transplanted to *Il-10^-/-^* (KO) GF recipient mice. B) Segment and total colon + ileum histology score for KO mice at day 14 post-colonization. C) Segment, total colon + ileum, and max segment histology score for KO mice at day 28 post-colonization. D) TNFα mRNA levels in cecal tissue at day 28 post-colonization. E) PCoA of FMT recipient WT and KO mouse groups, including NIMM-g1->KO group. F) 16S Seq taxonomic barplots show top 8 most abundant genera in FMT inputs and recipient mouse feces at day 28 post-colonization. For mouse recipient groups, barplots are average of 16S seq data from n=7-18 mice/group. G) Shannon diversity index at ASV level for IMM-g1->WT, NIMM-g1->KO and NIMM-g1->WT groups. Data shown are representative of (D) or cumulative (B-C, E-F) from 2-4 independent experiments. n=15-16 (B-C), n=5-8 (D), n=7-16 (E-G) mice per group. Data are expressed as mean±SD. Statistical significance calculated by unpaired t-test (B-D, G) with *p<0.05, **p<0.01, ***p<0.001, ****p<0.0001.

## Discussion

The role of gut microbiota dysbiosis as cause or consequence of intestinal inflammation is an area of active investigation and debate with clinical importance for the management of IBD^1^. Transplant of human disease-associated feces to GF rodents is an approach that captures strain-specific functional and genetic variation responsible for human-microbiome driven disease phenotypes without biased selection of defined input strains. Although widely accepted, this approach is complicated by phenotypic and experimental variability of unclear etiology. Our study identified that FMT transfer efficiency is an underappreciated source of experimental variability. Using high depth, low-error rate Illumina 16S amplicon sequencing (16S Seq), we showed that pooled human IBD patient fecal microbiota engrafts GF mice with low ASV-level transfer efficiency, resulting in high recipient-to-recipient variation of microbiota composition and colitis severity in HMA *Il-10^-/-^* mice. Human-to-mouse FMT caused a population bottleneck with reassembly of microbiota composition that was host inflammatory environment specific. In the inflamed environment of HMA *Il-10^-/-^* mice, the microbiota reassembled with lower microbial alpha diversity, higher recipient-to-recipient microbiota compositional variability, and expansion of pathobionts compared to the distinct microbiota reassembled in the non-inflamed environment of HMA WT mice. Following the initial human-to-mouse population bottleneck and microbiota reassembly, the mouse-adapted human IBD patient microbiota transferred with high efficiency and low compositional variability to GF recipients, which correlated with highly consistent and reproducible colitis phenotypes in *Il-10^-/-^* recipient mice. The mouse-adapted microbiota composition was remarkably stable in serial transplant to both inflamed and non-inflamed host environments. We replicated the key finding of low human-to-mouse but high mouse-adapted-to-mouse transfer efficiency at the ASV level by analysis of an independently published 16S Seq dataset of HMA WT mice bred in a gnotobiotic isolator^44^. Microbiota adaptation in the inflamed environment assembled a more aggressive microbiota than adaptation in the non-inflamed environment, demonstrating that the genetically determined host inflammatory environment shapes dysbiosis that subsequently drives more severe inflammation. Our data demonstrate that host gut inflammation is both a cause and consequence of microbial dysbiosis.

Our data support recent criticism that stochastic ecological processes and donor heterogeneity influence phenotypes in HMA murine models^61^. We found that OTU based metrics, especially at higher taxonomic levels, over-estimated transfer efficiency compared to ASV analysis^23, 44, 61^. The low transfer efficiency and population bottleneck of human-to-mouse FMT led to high variability in engrafted microbiota composition between individual recipients of the same human input stool, which correlated with significant variability in colitis severity in recipient *Il-10^-/-^* mice. We speculate that stochastic differences in engraftment were accentuated by the bottleneck of human-to-mouse FMT and drove phenotypic variability^61^. Large interindividual variability of human donor microbiota likely exacerbates this phenomenon in HMA murine studies^8, 57, 61^. We used pooled human IBD donor stool to mitigate the impact of individual human donor microbiota heterogeneity and replicated our results with 2 pooled human donor pools. Our pooling approach is suitable for experimental designs that require a representative human disease associated microbiome to interrogate mechanistic questions (i.e., the impact of diet or host genetic background) or test therapeutics (i.e., live biotherapeutics or novel biologics); however, studies evaluating microbiome-driven phenotype transfer require an appropriately powered number of individual human donors to establish causality and avoid bias from pseudo-replication^61^. In our study, both pooled human fecal cohorts HM1 and HM2 contained *Bacteroides* genus at high abundance. However, following human-to-mouse transplant, *Bacteroides* abundance dramatically decreased in HM1 recipients but expanded in HM2 recipients. Although our study was not powered to distinguish whether this divergent engraftment arose from microbial ecology of the donor microbiota or stochastic processes, our data suggest that low human-to-mouse transfer efficiency in the setting of donor heterogeneity and stochastic ecological processes is an underappreciated source of variability in HMA animal models. In contrast, transplant of mouse-adapted human microbiota yielded highly reproducible and consistent microbiota composition and colitis phenotypes – an improved model for studying human microbiota driven diseases.

Our data demonstrated that mouse-adaptation of human fecal microbiota was shaped by the host inflammatory environment to form stable microbial communities that reproducibly engrafted GF mice with high efficiency to drive distinct colitis phenotypes. Mouse-adaptation in the inflamed genetically susceptible host assembled an aggressive microbiota with low alpha-diversity and high pathobiont abundance (*Enterobacteriaceae, R. gnavus*) that drove more severe colitis in serial transplant to *Il-10^-/-^*mice than microbiota adapted in the non-inflamed host. Gut inflammation induces host-derived metabolites, such as nitrate, lactate, and ethanolamine, that enhance fitness, abundance, and virulence of aggressive *E. coli* and promote ectopic gut colonization of inflammation-associated *Veillonella* species^62-66^. Adherent and invasive *E. coli* and other inflammation-associated aggressive resident bacteria drive intestinal inflammation in murine colitis models^1, 18, 19, 21^. Together with the literature, our data support a model of IBD pathogenesis in which host inflammation in genetically susceptible hosts promotes the expansion, fitness, and virulence of aggressive resident bacteria, which further drive a feed-forward process of dysbiosis exacerbated gut inflammation. This model implies that effective management of IBD requires treating both the dysregulated host immune response and aggressive inflammation-associated microbiota.

Our study benefitted from several strengths including an experimental design that incorporated multiple serial FMT, high recipient mouse numbers, and application of high-depth low error rate sequencing for accurate ASV tracking; however, there were some limitations. First, most experiments were conducted in out-of-isolator gnotobiotic cages, where contamination risk is extremely low but could not be monitored due to the complex FMT inputs. To address this, we replicated key experiments in strict gnotobiotic isolators, confirming our findings of low human-to-mouse ASV-level transfer efficiency and the emergence of ASVs in HMA mice not detected in human input stool. Second, we did not analyze WGS data to compare transfer efficiency of microbial functions vs taxonomic composition. We used 16S Seq rather than WGS because repeat sequencing of the same region allows for exact identification of ASVs in a database-independent manner without reliance on classification algorithms. Future WGS studies are needed to evaluate the impact of taxonomic transfer efficiency on transfer of microbial functions. Third, we did not evaluate the impact of mouse diet on the initial human-to-mouse engraftment bottleneck – an important topic for follow up studies.

Our mouse-adapted human microbiota model is an optimized, reproducible, and rigorous system to study human microbiome-driven disease phenotypes. Multiple approaches (human microbiome profiling, defined consortia animal studies, HMA animal models) can investigate causality and identify mechanisms of microbiota-driven diseases^1, 29, 67^. Mono-association and defined consortium studies are reductionist approaches where a single variable (i.e., single-gene mutations) can interrogate bacterial mechanisms^1, 29^. Representative synthetic microbiota, such as hCOM2, PedsCom, and SIHUMI, provide a more ecologically complex system with known input strain identity and the ability to easily track relative abundance by simplified metagenomic sequencing approaches^28, 31, 32, 67, 68^. However, even large complex defined consortia do not capture the understudied strain level variation that exists in heterogeneous human resident microbiota and contributes to important differences in strain dependent microbiota aggressiveness^22, 36, 37, 66^. The high transfer efficiency of mouse-adapted human microbiota transplant to GF mice improves phenotype consistency, experiment reproducibility and rigor of mouse models of human microbiota-driven disease. Homogenous repositories of mouse-adapted human microbiota provide an identical microbial starting point for every experiment that can be replicated over time and between institutions/collaborators without transfer of human host genetic material present in human feces to collaborators^69, 70^. Because of high transfer efficiency and reproducible engraftment, mouse-adapted human microbiota repositories can be expanded *in vivo* when stocks run low, mitigating limitations of finite human fecal samples. While this study focused on colitis, our mouse-adapted human microbiota approach is a framework that may be generalized to mouse FMT models of other human microbiota-modulated diseases, such as metabolic syndrome/obesity, diabetes, autoimmune diseases, and cancer.

## Acknowledgements

We thank the members of the Sartor and Fodor laboratories for helpful comments, suggestions, and support. This study was supported in part by NIH/NIDDK T32DK007737 (SMG), P01DK094779 (RBS), P40OD010995 (RBS), CGIBD P30DK034987 (RBS), Crohn’s & Colitis Foundation Gnotobiotic Facility (RBS), UNC Physician Scientist Training Program Fellowship Award (SMG), Crohn’s & Colitis Foundation Career Development Award (SK). We thank the staff and manager, Josh Frost, of the National Gnotobiotic Rodent Resource Center for gnotobiotic husbandry and mice. The UNC Microbiome Core is funded in part by the UNC Nutrition Obesity Research Center (NORC P30 DK056350).

## Data Availability Statement

The datasets, R code, and Python code are publicly available at https://github.com/anhmoss/Mouse-Adaptation-of-Human-Inflammatory-Bowel-Disease-Microbiota-Enhances-Colonization-Efficiency and from the corresponding authors upon reasonable request.

## Tables

**Supplementary Table S1:** Differential abundance analysis between groups excluding genera present in less than 10% of the samples.

## Figure Legends

**Supplemental Figure S1. (related to Figure 1).**
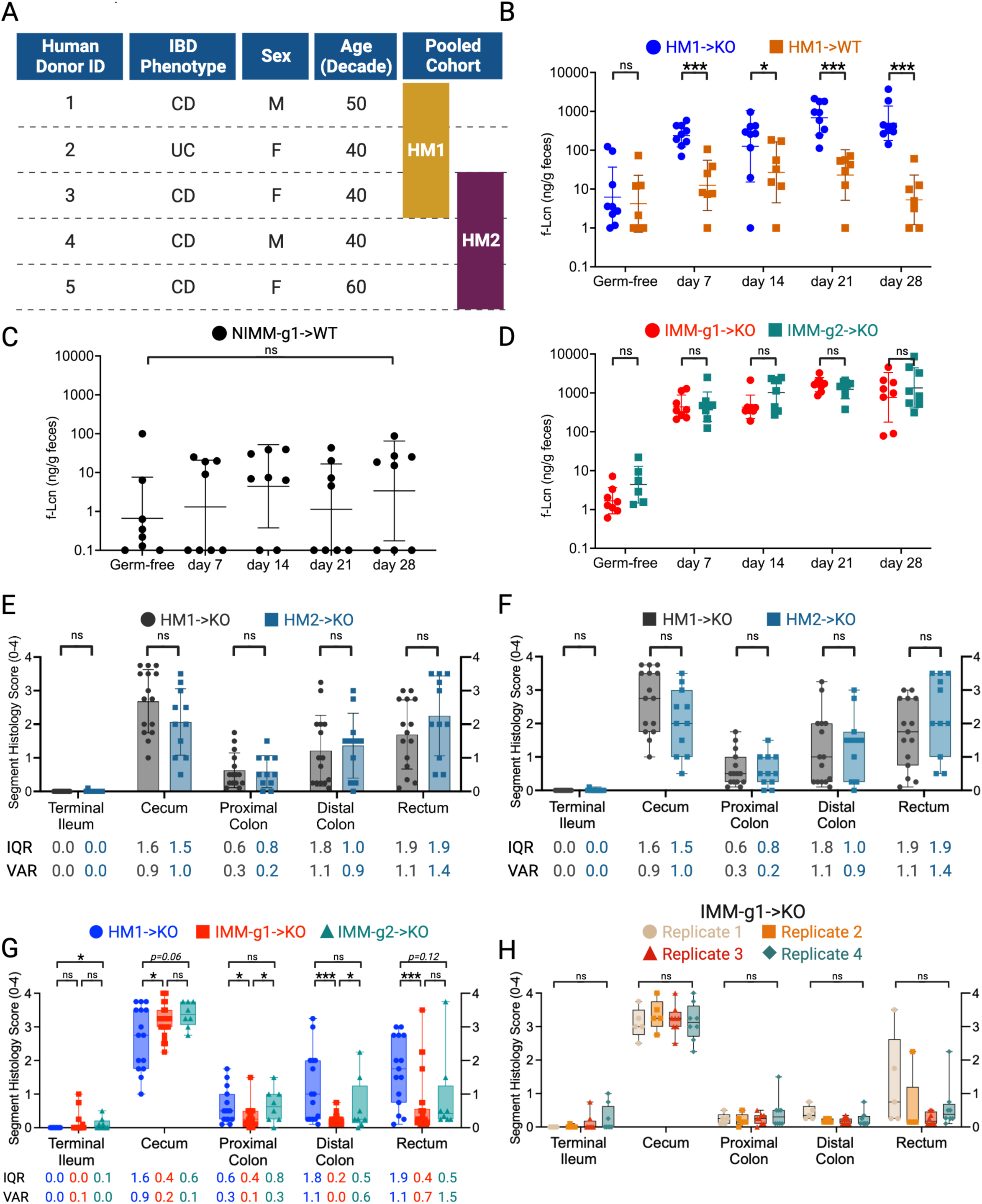
Mouse-adapted human microbiota induces more consistent and reproducible colitis than directly transplanted human microbiota. A) Demographics and cohort membership of human IBD donors. B-D) f-LCN2 time-course of (B) HM1 colonized WT and KO mice, (C) NIMM-g1 colonized WT mice, (D) IMM-g1 or IMM-g2 colonized KO mice. E-F) Bar plot (E) and Box-and-whisker (F) plot of segment histology score for HM1 and HM2 colonized KO mice at day 28 post-colonization. G) Box-and-whisker plot of segment histology score for KO mice at day 28 post-colonization. H) Box-and-whisker plot of segment histology score for IMM-g1 colonized KO mice at day 28 post-colonization from 4 independent experiments. Data shown are representative of (B-D, H) or cumulative (E-G) from 2-4 independent experiments. n=7-9 (B-D), n=11-15 (E-F), n=15-26 (G), n=5-8 (H) mice per group. Data are expressed as mean±SD (E) or geometric mean ± geometric SD (B-D). In box-and-whisker plots, box represents lower, median, and upper quartiles; whiskers are min to max. Statistical significance calculated by Mann-Whitney test (B-D) or unpaired t-test (E-H) with *p<0.05, **p<0.01, ***p<0.001.

**Supplemental Figure S2 (related to Figure 2).**
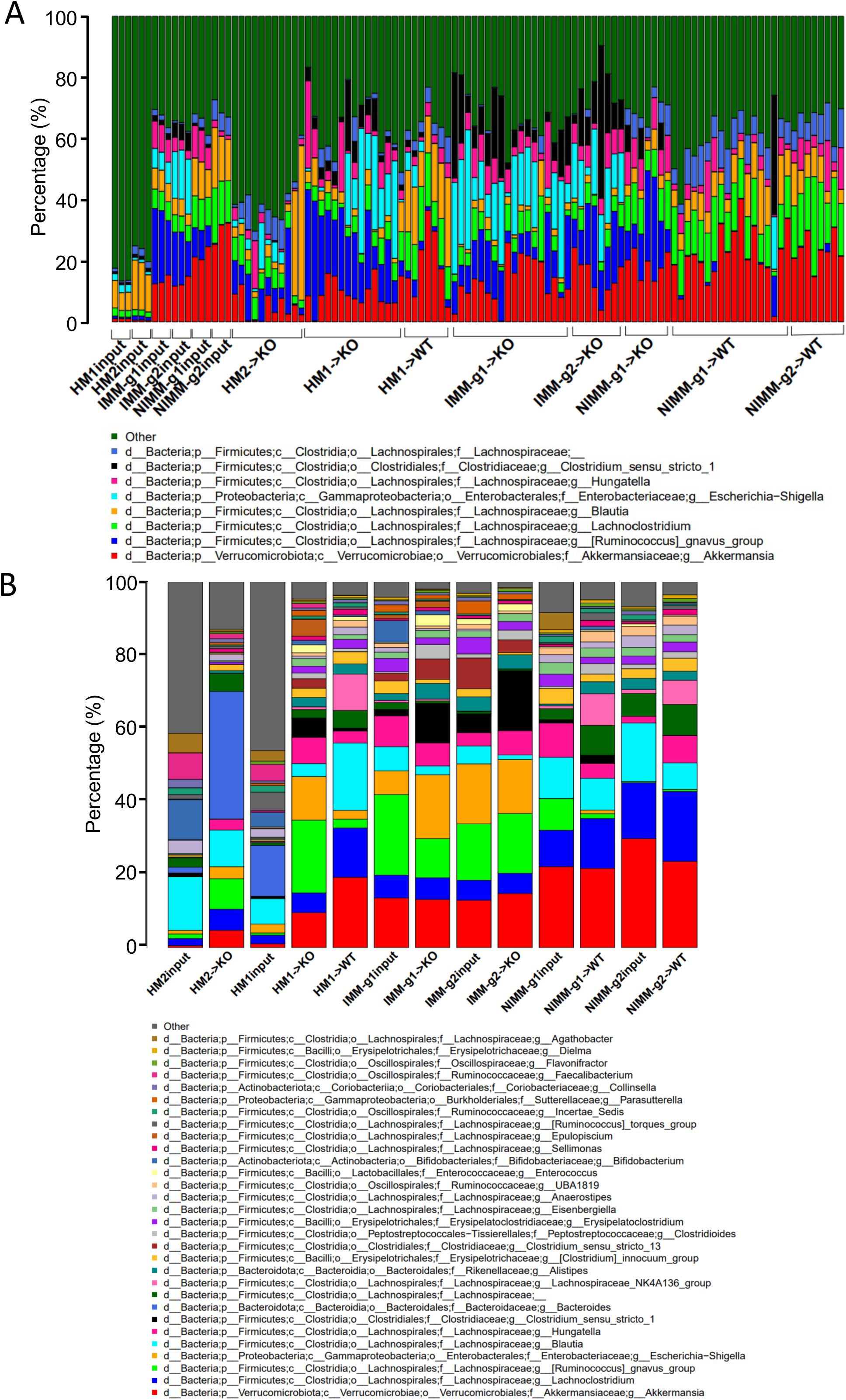
Recipient host environment influences engraftment composition of human-microbiome associated mice. A) 16S Seq taxonomic bar plots show top 8 most abundant genera in human and mouse-adapted FMT inputs and individual recipient mouse feces at day 28 post-colonization. B) 16S Seq taxonomic bar plots show top 30 most abundant genera in human and mouse-adapted FMT inputs and recipient mouse feces at day 28 post-colonization. For mouse recipient groups, bar plots are average of 16S Seq data from n=7-18 mice/group.

**Supplemental Figure S3 (related to Figure 3).**
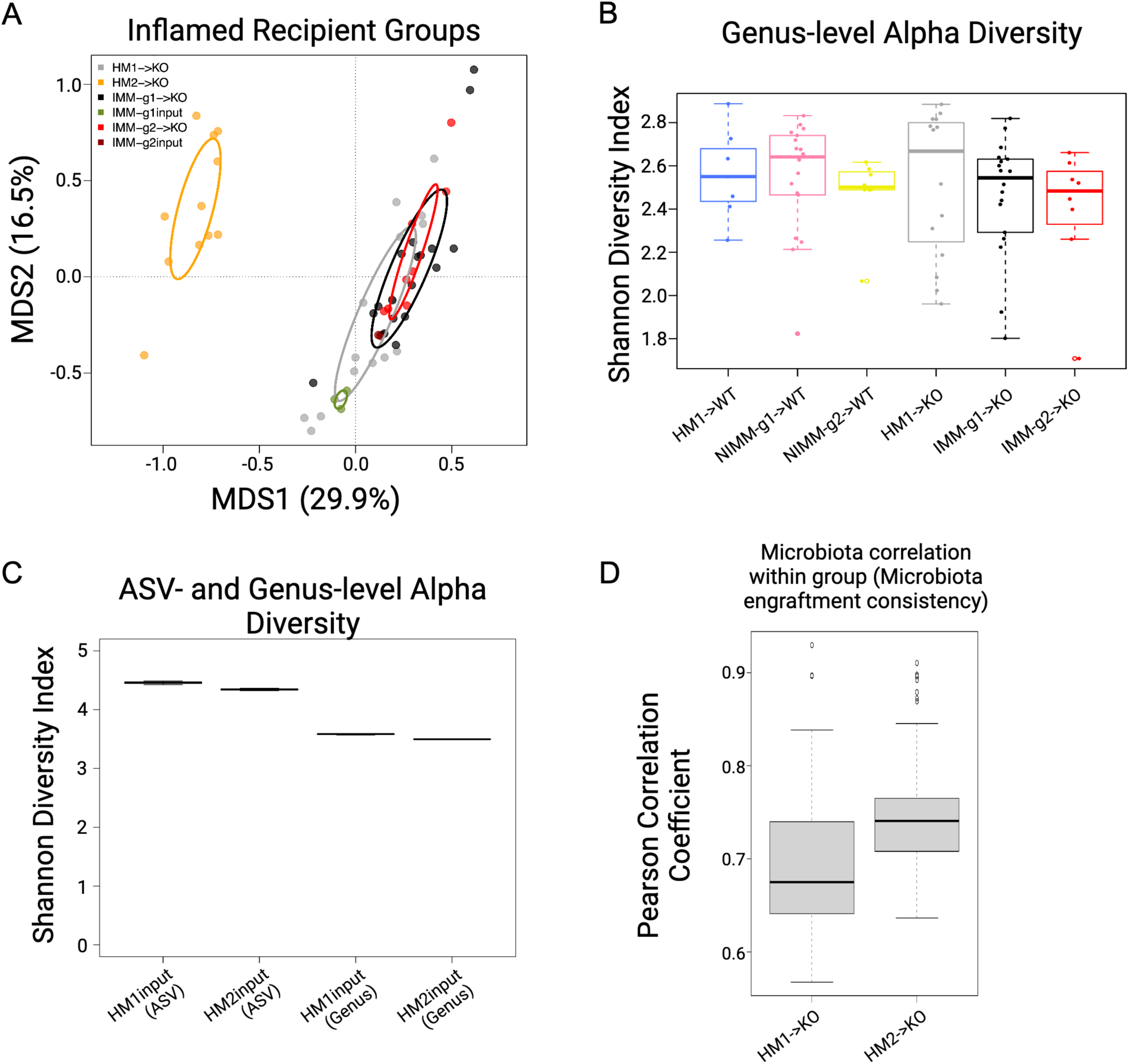
Human microbiome restructuring with transplant to GF mice is host inflammatory environment specific. A) PCoA of 16S Seq data for HM1, HM2, and mouse-adapted FMT inputs and FMT recipient KO mouse groups. B) Shannon index at genus level for FMT recipient WT and KO mouse groups. C) Shannon index at ASV and genus level for HM1 and HM2 FMT inputs. D) Pearson correlation coefficient (r) within group for HM1->KO and HM2->KO recipient mouse groups quantifies variability of microbiota composition between mice in the same group (microbiota engraftment consistency).

**Supplemental Figure S4 (related to Figure 4).**
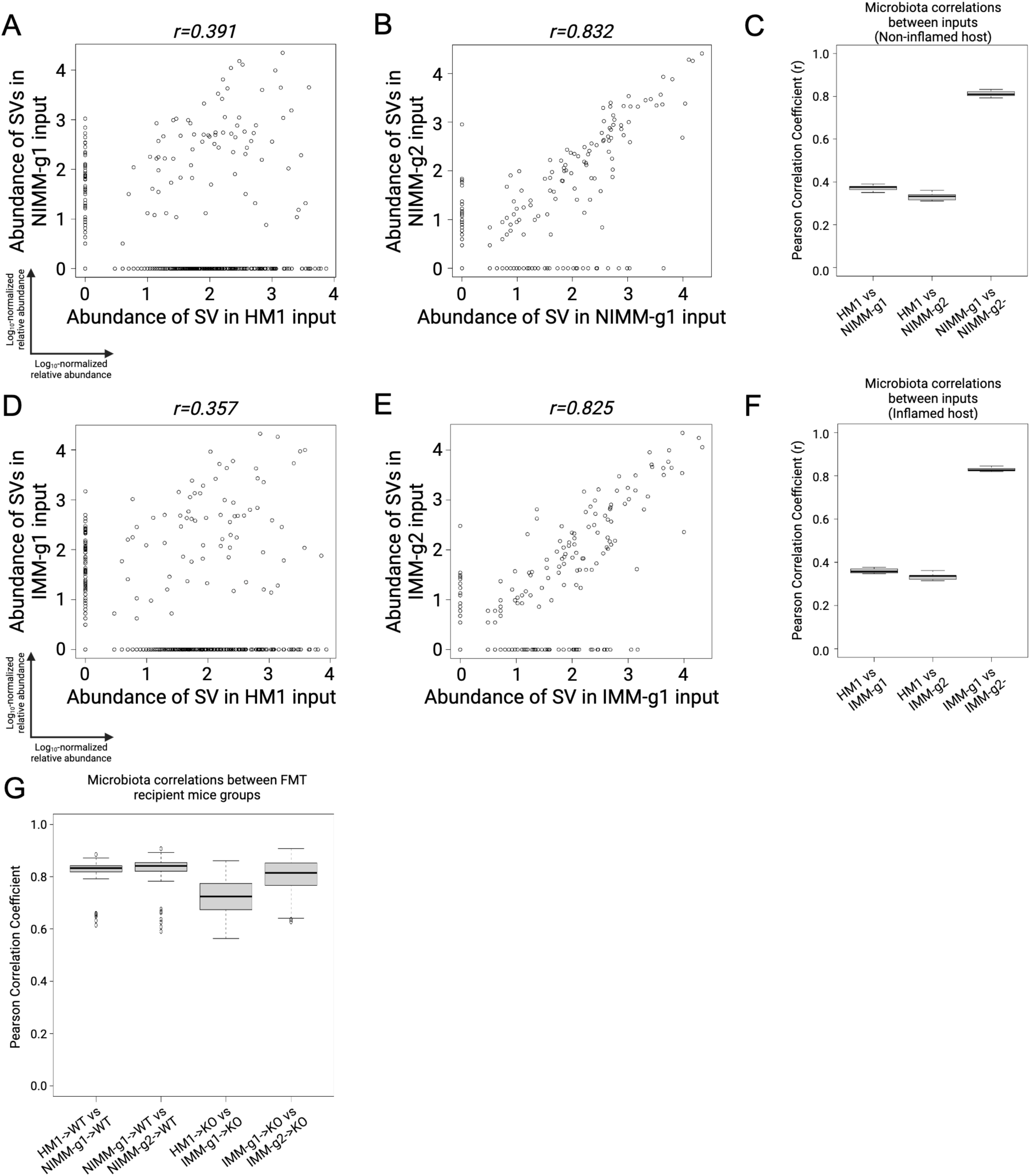
Mouse-adapted human IBD microbiota transfers with higher efficiency than human fecal transplant. A-B) ASV level log_10_-normalized relative abundance correlations comparing (A) HM1 input to NIMM-g1 input and (B) NIMM-g1 input to NIMM-g2 input. C) Pearson correlation coefficient (r) between HM1 input, NIMM-g1 input, and NIMM-g2 inputs at the ASV level. D-E) ASV level log_10_-normalized relative abundance correlations comparing (D) HM1 input to IMM-g1 input and (E) IMM-g1 input to IMM-g2 input. F) Pearson correlation coefficient (r) between HM1 input, IMM-g1 input, and IMM-g2 inputs at the ASV level. G) Pearson correlation coefficient (r) between WT recipient and KO recipient groups at the ASV level.

**Supplemental Figure S5 (related to Figure 5).**
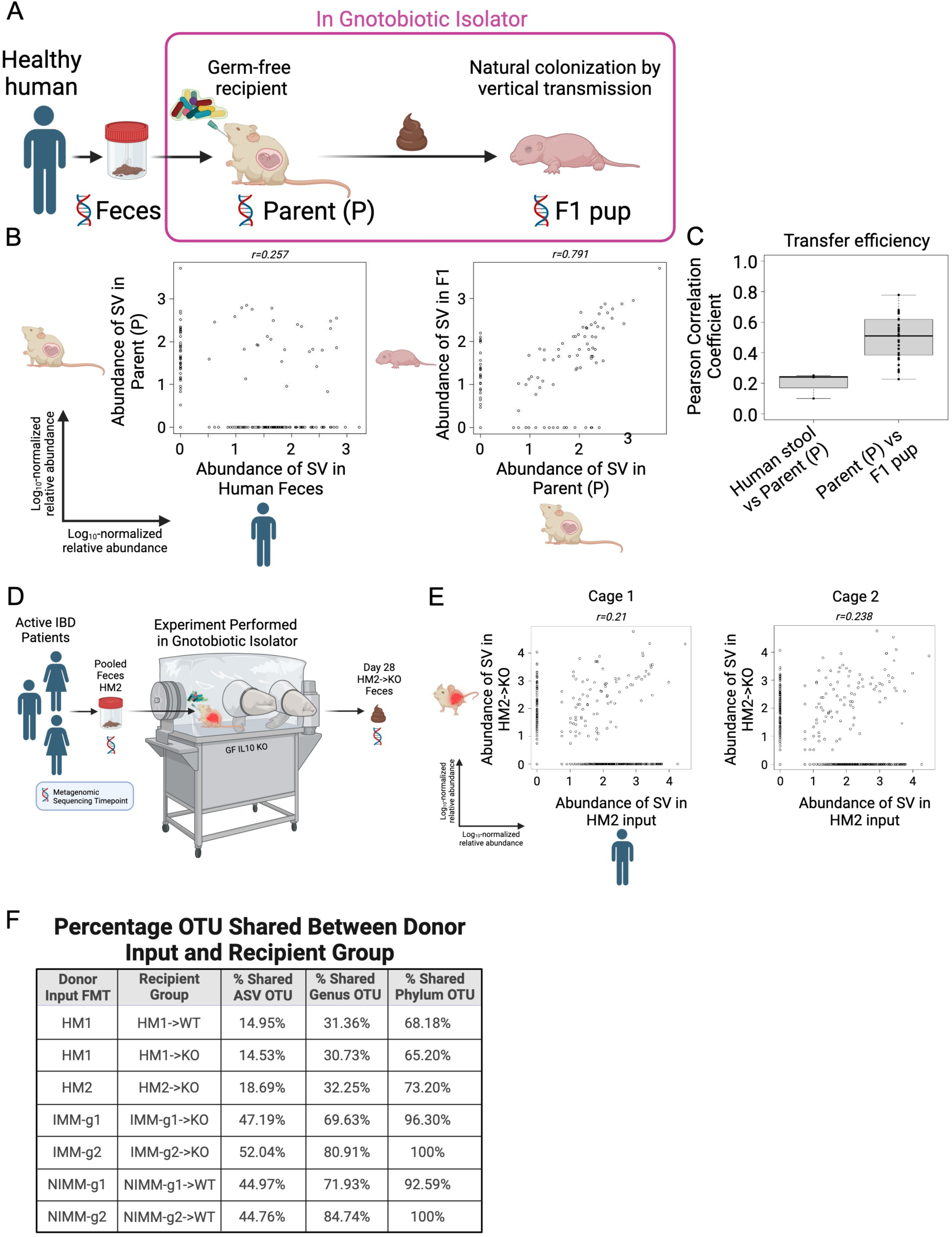
Transfer efficiency varies between taxa. A-C) Analysis of published 16S Seq data from Lundberg *et al*.^44^ (A) Experimental design. Feces from a single healthy human donor were transplanted to GF adult WT mice (Parent, P) in a gnotobiotic isolator. HMA WT mice were bred in-isolator to generate F1 pups. B) ASV level log_10_-normalized relative abundance correlations for FMT input and recipient mice. For F1 pups, the input was natural colonization by vertical transmission in-isolator from Parent (P). C) Transfer efficiency quantified by Pearson correlation coefficient (r) between FMT input and recipient mouse groups at the ASV level. D) Experimental design. Pooled feces from 3 humans with active IBD (3 CD, HM2) were transplanted to colitis-susceptible *Il-10*^-/-^ (KO) GF recipient mice in a gnotobiotic isolator. E) ASV level log_10_-normalized relative abundance correlations for HM2 input and HM2->KO recipient mice. F) Table comparing percentage shared OTU between input and recipient group at the ASV, Genus, and Phylum level.

**Supplemental Figure S6 (related to Figure 6).**
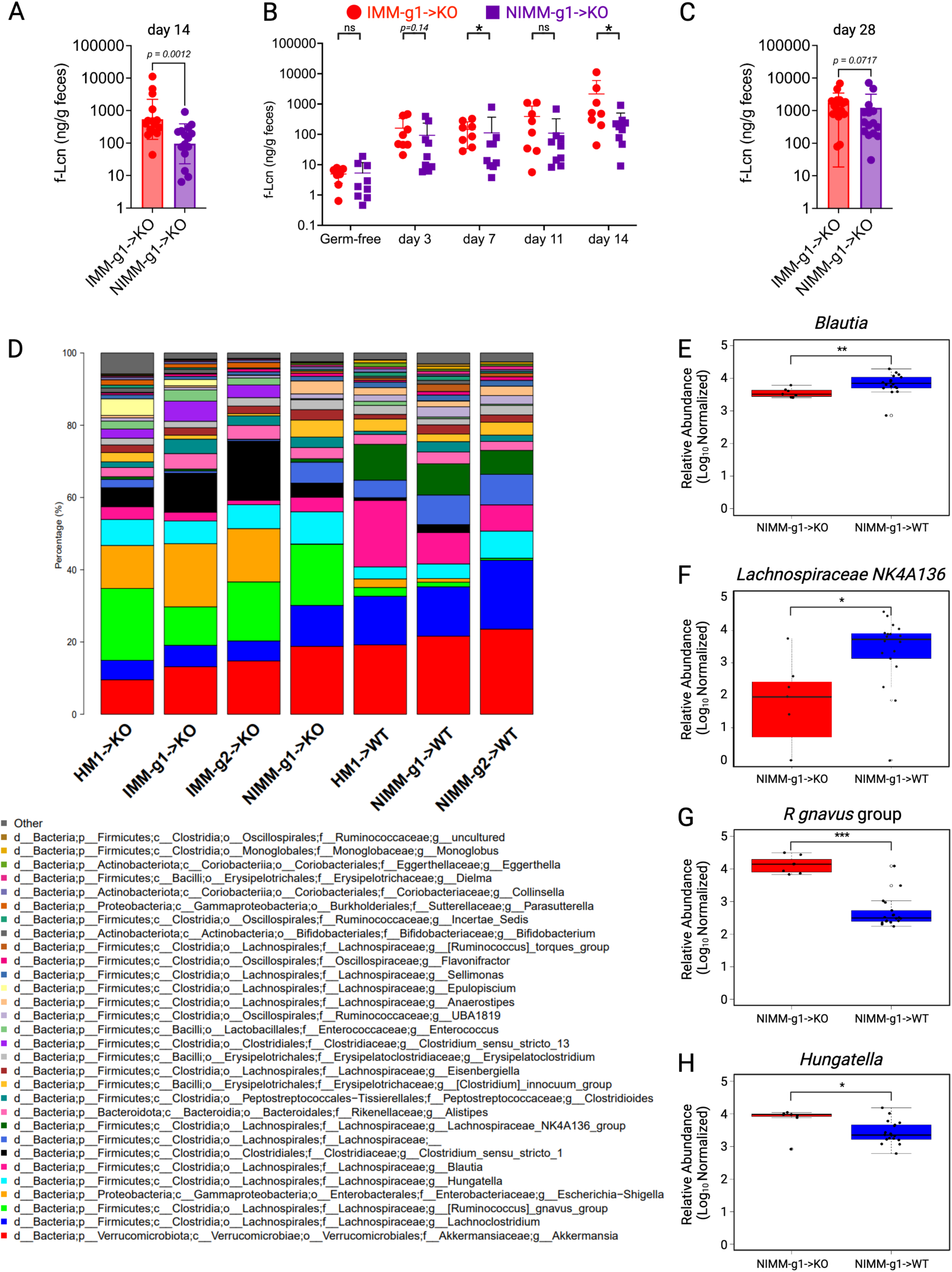
Inflamed mouse-adapted microbiome more rapidly induces severe colitis than non-inflamed mouse adapted microbiome. A-C) f-LCN2 level of IMM-g1->KO and NIMM-g1->KO mice at (A) day 14, (B) time-course from day 0 to day 14, and (C) day 28 post-colonization. D) 16S Seq taxonomic bar plots show top 30 most abundant genera in human and mouse-adapted FMT inputs and recipient mouse feces at day 28 post-colonization. E-G) log_10_- normalized relative abundance of (E) *Blautia,* (F) *Lachnospiraceae NK4A136* group, (G) *R gnavus* group, (H) *Hungatella* in NIMM-g1->KO vs NIMM-g1->WT mice. For mouse recipient groups, barplots are average of 16S Seq data from n=7-18 mice/group. Data shown are cumulative (A) or representative (B-C) of 2 independent experiments. n=15-16 (A), n=8-9 (B), n=7-8 (C) mice per group. Data are expressed as geometric mean ± geometric SD (A-C). Statistical significance calculated by Mann-Whitney test (A-C) or unpaired t-test (E-H) with *p<0.05, **p<0.01, ***p<0.001.

## Supplemental Information

**Supplemental File 1**: RMarkdown notebook of R Code Analysis and Jupyter Notebook of Python Code Analysis. Also publicly available at https://github.com/anhmoss/Mouse-Adaptation-of-Human-Inflammatory-Bowel-Disease-Microbiota-Enhances-Colonization-Efficiency.

## Supplemental Experimental Procedures

### Gene expression by qRT-PCR

Tissues were immediately placed in RNAprotect cell reagent to stabilize RNA. Total RNA extraction from tissues (AllPrep PowerViral DNA/RNA Kit, Qiagen) and cDNA generation (iScript cDNA Synthesis Kit, Bio-Rad) were performed according to the manufacturer’s protocols. Quantitative RT-PCR was performed on cDNA in duplicate or triplicate with a QuantStudio3 machine (ThermoFisher) using iTaq™ Universal SYBR Green Supermix (Bio-Rad). Target gene expression was quantified relative to internal control b-actin and expressed using the comparative Ct method (2^-ΔΔCt^). qPCR primer sequences are: Tnfa-F 5’-ACCCTCACACTCAGATCATCTTCTC-3’, Tnfa-R 5’-TGAGATCCATGCCGTTGG-3’. Actb-F 5’-AGCCATGTACGTAGCCATCCAG-3’; Actb-R 5’-TGGCGTGAGGGAGAGCATAG-3’

### Intestine histopathology scoring

Small bowel and colon tissue sections were fixed in 10% phosphate-buffered formalin. Fixed tissue was paraffin-embedded, sectioned at 5μm thickness, and stained with hematoxylin and eosin by the UNC Center for Gastrointestinal Biology and Disease Histology Core. Histologic tissue inflammation was quantified by blinded scoring as previously described on a scale of 0-4 for 5 tissue segments (terminal ileum, cecum, proximal colon, distal colon, rectum)^1^. Total inflammatory score was the summation of the 5 segments. Max segment score was the single highest score from the 5 segments.

### Fecal lipocalin-2 quantification

Fecal samples (10-30mg) were homogenized in PBS with 0.1% Tween 20 and incubated at 4°C overnight, followed by centrifugation to pellet solid debris. Lipocalin-2 ELISA was performed on clear fecal supernatant according to manufacturer’s instructions (DY1857, R&D Systems)^2^.

### Microbial and Statistical Analyses

16S rRNA amplicon (variable regions 3-4) PCR and sequencing were performed by the UNC Microbiome Core. Sequencing was performed on an Illumina MiSeq platform. Sequencing outputs were converted to fastq format and demultiplexed using Illumina Bcl2Fastq 2.20.0. The resulting paired-end reads were processed with the QIIME2 2022-2 wrapper for DADA2 including merging paired ends, quality filtering, error correction, and chimera detection^3, 4^. Amplicon sequencing variants from DADA2 were assigned taxonomy with respect to the Silva databases, their sequences were aligned using maFFT in QIIME2, and a phylogenetic tree was built with FastTree in QIIME2^5-7^.

