## Supplemental Table S1 for "Mouse Adaptation of Human Inflammatory Bowel Diseases Microbiota Enhances Colonization Efficiency and Alters Microbiome Aggressiveness Depending on Recipient Colonic Inflammatory Environment"

**Supplemental Table 1**

**Explanation of analysis**: We performed pairwise t-tests to assess differential abundance between groups, excluding genera present in less than 10% of the samples

**Explanation of tables in tabs**: Tabs represent pairwise comparisons of mouse FMT recipient groups. See Fig 1A for detailed experimental design and explanation of nomenclature. Statistics are calculated for differential abundance comparisons between mouse FMT recipient groups.

**Nomenclature Explanation**

| Donor Input FMT | GF Mouse Recipient | Recipient Group Label | Recipient Microbiome Name |
|---|---|---|---|
| HM1 | WT | HM1->WT | NIMM-g1 |
| NIMM-g1 | WT | NIMM-g1->WT | NIMM-g2 |
| NIMM-g2 | WT | NIMM-g2->WT | NIMM-g3 |
| HM1 | KO | HM1->KO | IMM-g1 |
| IMM-g1 | KO | IMM-g1->KO | IMM-g2 |
| IMM-g2 | KO | IMM-g2->KO | IMM-g3 |
| HM2 | KO | HM2->KO | ~ |

**Tabs in Workbook**

HM1->KO vs HM1->WT
HM1->KO vs HM2->KO
HM1->KO vs IMM-g1->KO
IMM-g1->KO vs IMM-g2->KO
IMM-g1->KO vs NIMM-g1->KO
IMM-g1->KO vs NIMM-g1->WT
HM1->WT vs NIMM-g1->WT
NIMM-g1->WT vs NIMM-g2->WT
NIMM-g1->KO vs NIMM-g1->WT

| Operational Taxonomic Unit (Genus) for HM1->KO vs HM1->WT Comparison | T-value | P-value | FDR | Enriched |
|---|---|---|---|---|
| d__Bacteria;p__Firmicutes;c__Clostridia;o__Lachnospirales;f__Lachnospiraceae;g__Lachnospiraceae | -8.488550283 | 6.33E-08 | 3.12E-06 | HM1->WT |
| d__Bacteria;p__Firmicutes;c__Clostridia;o__Lachnospirales;f__Lachnospiraceae;g__[Eubacterium]_fissicatena_group | -9.134589714 | 1.24E-07 | 3.12E-06 | HM1->WT |
| d__Bacteria;p__Firmicutes;c__Clostridia;o__Lachnospirales;f__Lachnospiraceae;g__Lachnospiraceae_NK4A136_group | -9.135735196 | 1.25E-07 | 3.12E-06 | HM1->WT |
| d__Bacteria;p__Firmicutes;c__Clostridia;o__Lachnospirales;f__Lachnospiraceae;g__Epulopiscium | 7.138571583 | 1.07E-06 | 1.69E-05 | HM1->KO |
| d__Bacteria;p__Actinobacteriota;c__Coriobacteriia;o__Coriobacteriales;f__Eggerthellaceae;g__Adlercreutzia | -7.254382295 | 1.12E-06 | 1.69E-05 | HM1->WT |
| d__Bacteria;p__Firmicutes;c__Clostridia;o__Lachnospirales;f__Lachnospiraceae;g__Blautia | -6.773520585 | 2.10E-06 | 2.62E-05 | HM1->WT |
| d__Bacteria;p__Firmicutes;c__Clostridia;o__Lachnospirales;f__Lachnospiraceae;g__[Ruminococcus]_gnavus_group | 7.951232103 | 3.16E-06 | 3.39E-05 | HM1->KO |
| d__Bacteria;p__Actinobacteriota;c__Coriobacteriia;o__Coriobacteriales;f__Eggerthellaceae;g__Gordonibacter | -6.145960666 | 1.24E-05 | 0.000112071 | HM1->WT |
| d__Bacteria;p__Firmicutes;c__Clostridia;o__Lachnospirales;f__Lachnospiraceae;g__Lachnoclostridium | -5.788720245 | 1.34E-05 | 0.000112071 | HM1->WT |
| d__Bacteria;p__Actinobacteriota;c__Coriobacteriia;o__Coriobacteriales;f__Eggerthellaceae;g__Eggerthella | -5.661915153 | 3.94E-05 | 0.000295402 | HM1->WT |
| d__Bacteria;p__Firmicutes;c__Clostridia;o__Oscillospirales;f__Oscillospiraceae;g__UCG-005 | -4.598911261 | 0.000179748 | 0.001225555 | HM1->WT |
| d__Bacteria;p__Firmicutes;c__Clostridia;o__Clostridiales;f__Clostridiaceae;g__Clostridium_sensu_stricto_13 | 4.524472145 | 0.000236423 | 0.001477646 | HM1->KO |
| d__Bacteria;p__Firmicutes;c__Clostridia;o__Oscillospirales;f__Ruminococcaceae;g__Paludicola | -4.332486538 | 0.000432429 | 0.002494784 | HM1->WT |
| d__Bacteria;p__Firmicutes;c__Clostridia;o__Oscillospirales;f__Ruminococcaceae;g__Negativibacillus | 4.278339537 | 0.000764954 | 0.004097967 | HM1->KO |
| d__Bacteria;p__Firmicutes;c__Clostridia;o__Lachnospirales;f__Lachnospiraceae;g__Anaerostipes | -4.139976324 | 0.000900938 | 0.004504692 | HM1->WT |
| d__Bacteria;p__Proteobacteria;c__Gammaproteobacteria;o__Vibrionales;f__Vibrionaceae;g__Vibrio | -3.745828966 | 0.001400114 | 0.006563034 | HM1->WT |
| d__Bacteria;p__Firmicutes;c__Clostridia;o__Oscillospirales;f__Ruminococcaceae;g__Phocea | 3.449202222 | 0.003910844 | 0.017253722 | HM1->KO |
| d__Bacteria;p__Firmicutes;c__Clostridia;o__Lachnospirales;f__Lachnospiraceae;__ | -3.208690271 | 0.005289446 | 0.020941013 | HM1->WT |
| d__Bacteria;p__Firmicutes;c__Clostridia;o__Peptococcales;f__Peptococcaceae;g__uncultured | 3.295931565 | 0.005305057 | 0.020941013 | HM1->KO |
| d__Bacteria;p__Firmicutes;c__Clostridia;o__Monoglobales;f__Monoglobaceae;g__Monoglobus | -2.922851884 | 0.008434306 | 0.031628648 | HM1->WT |
| d__Bacteria;p__Firmicutes;c__Clostridia;o__Peptostreptococcales-Tissierellales;f__Peptostreptococcaceae;g__Terrisporobacter | 2.950212748 | 0.010541452 | 0.034754231 | HM1->KO |
| d__Bacteria;p__Firmicutes;c__Clostridia;o__Lachnospirales;f__Lachnospiraceae;g__[Ruminococcus]_torques_group | -3.01158123 | 0.010549957 | 0.034754231 | HM1->WT |
| d__Bacteria;p__Firmicutes;c__Clostridia;o__Peptostreptococcales-Tissierellales;f__Anaerovoracaceae;g__Family_XIII_AD3011_group | -2.816774219 | 0.010657964 | 0.034754231 | HM1->WT |
| d__Bacteria;p__Firmicutes;c__Clostridia;o__Lachnospirales;f__Lachnospiraceae;g__Lactonifactor | 2.81786847 | 0.013690093 | 0.04278154 | HM1->KO |
| d__Bacteria;p__Firmicutes;c__Clostridia;o__Clostridiales;f__Clostridiaceae;g__Clostridium_sensu_stricto_1 | 3.069360849 | 0.020296601 | 0.060889804 | HM1->KO |
| d__Bacteria;p__Firmicutes;c__Bacilli;o__Lactobacillales;f__Streptococcaceae;g__Streptococcus | 2.596328893 | 0.021127732 | 0.06094538 | HM1->KO |
| d__Bacteria;p__Bacteroidota;c__Bacteroidia;o__Bacteroidales;f__Rikenellaceae;g__Alistipes | -2.517879932 | 0.024064763 | 0.066846565 | HM1->WT |
| d__Bacteria;p__Firmicutes;c__Clostridia;o__Oscillospirales;f__Oscillospiraceae;g__Oscillibacter | 2.419595914 | 0.032886627 | 0.08808918 | HM1->KO |
| d__Bacteria;p__Firmicutes;c__Bacilli;o__Erysipelotrichales;f__Erysipelotrichaceae;g__Turicibacter | 2.22759033 | 0.042823474 | 0.110750364 | HM1->KO |
| d__Bacteria;p__Actinobacteriota;c__Coriobacteriia;o__Coriobacteriales;f__Coriobacteriaceae;g__Collinsella | -2.124507252 | 0.047745904 | 0.119364759 | HM1->WT |
| d__Bacteria;p__Verrucomicrobiota;c__Verrucomicrobiae;o__Verrucomicrobiales;f__Akkermansiaceae;g__Akkermansia | -2.001235886 | 0.062939453 | 0.152272869 | HM1->WT |
| d__Bacteria;p__Firmicutes;c__Clostridia;o__Lachnospirales;f__Lachnospiraceae;g__GCA-900066755 | -1.917141007 | 0.069698258 | 0.157373258 | HM1->WT |
| d__Bacteria;p__Firmicutes;c__Clostridia;o__Oscillospirales;f__Ruminococcaceae;g__Ruminococcus | 1.921115965 | 0.070260433 | 0.157373258 | HM1->KO |
| d__Bacteria;p__Firmicutes;c__Clostridia;o__Oscillospirales;f__[Eubacterium]_coprostanoligenes_group;g__[Eubacterium]_coprostanoligenes_group | 1.904714488 | 0.071342544 | 0.157373258 | HM1->KO |
| d__Bacteria;p__Firmicutes;c__Clostridia;o__Lachnospirales;f__Lachnospiraceae;g__uncultured | 1.891683194 | 0.077301629 | 0.164910925 | HM1->KO |
| d__Bacteria;p__Firmicutes;c__Clostridia;o__Lachnospirales;f__Lachnospiraceae;g__Lachnospiraceae_UCG-008 | -1.991940783 | 0.080261553 | 0.164910925 | HM1->WT |
| d__Bacteria;p__Firmicutes;c__Clostridia;o__Oscillospirales;f__Ruminococcaceae;g__Incertae_Sedis | -1.846780654 | 0.083487288 | 0.164910925 | HM1->WT |
| d__Bacteria;p__Firmicutes;c__Clostridia;o__Oscillospirales;f__Ruminococcaceae;g__Subdoligranulum | 1.863142456 | 0.083554869 | 0.164910925 | HM1->KO |
| d__Bacteria;p__Firmicutes;c__Clostridia;o__Oscillospirales;f__Ruminococcaceae;g__Faecalibacterium | 1.841035519 | 0.086903937 | 0.167122956 | HM1->KO |
| d__Bacteria;p__Proteobacteria;c__Gammaproteobacteria;o__Burkholderiales;f__Sutterellaceae;g__Parasutterella | 1.777696261 | 0.093397828 | 0.175120928 | HM1->KO |
| d__Bacteria;p__Bacteroidota;c__Bacteroidia;o__Bacteroidales;f__Tannerellaceae;g__Parabacteroides | -1.959500036 | 0.097762888 | 0.178834551 | HM1->WT |
| d__Bacteria;p__Firmicutes;c__Bacilli;o__Erysipelotrichales;f__Erysipelatoclostridiaceae;g__Erysipelatoclostridium | -1.723035528 | 0.105053803 | 0.187596077 | HM1->WT |
| d__Bacteria;p__Firmicutes;c__Bacilli;o__Erysipelotrichales;f__Erysipelotrichaceae;g__[Clostridium]_innocuum_group | -1.689270557 | 0.108200975 | 0.18872263 | HM1->WT |
| d__Bacteria;p__Firmicutes;c__Clostridia;o__Peptostreptococcales-Tissierellales;f__Peptostreptococcaceae;g__Clostridioides | 1.693141168 | 0.124264566 | 0.211814601 | HM1->KO |
| d__Bacteria;p__Firmicutes;c__Clostridia;o__Lachnospirales;f__Lachnospiraceae;g__Eisenbergiella | 1.617884358 | 0.134850552 | 0.22475092 | HM1->KO |
| d__Bacteria;p__Firmicutes;c__Clostridia;o__Lachnospirales;f__Lachnospiraceae;g__Sellimonas | -1.516020725 | 0.145196507 | 0.234861695 | HM1->WT |
| d__Bacteria;p__Firmicutes;c__Clostridia;o__Oscillospirales;f__Ruminococcaceae;g__Anaerotruncus | -1.506298725 | 0.147695236 | 0.234861695 | HM1->WT |
| d__Bacteria;p__Firmicutes;c__Clostridia;o__Oscillospirales;f__Ruminococcaceae;g__UBA1819 | -1.546946505 | 0.150311485 | 0.234861695 | HM1->WT |
| d__Bacteria;p__Firmicutes;c__uncultured;o__uncultured;f__uncultured;g__uncultured | 1.467462799 | 0.164354338 | 0.249863757 | HM1->KO |
| d__Bacteria;p__Firmicutes;c__Clostridia;o__Lachnospirales;f__Lachnospiraceae;g__Marvinbryantia | 1.459220545 | 0.166575838 | 0.249863757 | HM1->KO |
| d__Bacteria;p__Proteobacteria;c__Gammaproteobacteria;o__Enterobacterales;f__Enterobacteriaceae;g__Escherichia-Shigella | 1.354936281 | 0.197656354 | 0.290671108 | HM1->KO |
| d__Bacteria;p__Firmicutes;c__Bacilli;o__Erysipelotrichales;f__Erysipelotrichaceae;g__Dielma | -1.250680558 | 0.231613284 | 0.334057621 | HM1->WT |
| d__Bacteria;p__Firmicutes;c__Clostridia;o__Lachnospirales;f__Lachnospiraceae;g__GCA-900066575 | -1.236556149 | 0.241293998 | 0.341453771 | HM1->WT |
| d__Bacteria;__;__;__;__;__ | -1.217579016 | 0.262918564 | 0.365164672 | HM1->WT |
| d__Bacteria;p__Firmicutes;c__Clostridia;o__Oscillospirales;f__Oscillospiraceae;g__uncultured | -1.12878595 | 0.274527839 | 0.374356144 | HM1->WT |
| d__Bacteria;p__Firmicutes;c__Clostridia;o__Oscillospirales;f__Butyricicoccaceae;g__Butyricicoccus | 1.020982739 | 0.323503001 | 0.426149105 | HM1->KO |
| d__Bacteria;p__Firmicutes;c__Clostridia;o__Lachnospirales;f__Lachnospiraceae;g__Roseburia | 1 | 0.334281943 | 0.426149105 | HM1->KO |
| d__Bacteria;p__Actinobacteriota;c__Coriobacteriia;o__Coriobacteriales;f__Eggerthellaceae;__ | 1 | 0.334281943 | 0.426149105 | HM1->KO |
| d__Bacteria;p__Firmicutes;c__Bacilli;o__Staphylococcales;f__Staphylococcaceae;g__Staphylococcus | 0.98906855 | 0.335237296 | 0.426149105 | HM1->KO |
| d__Bacteria;p__Firmicutes;c__Clostridia;o__Oscillospirales;f__Ruminococcaceae;g__DTU089 | -1 | 0.355917684 | 0.444897105 | HM1->WT |
| d__Bacteria;p__Bacteroidota;c__Bacteroidia;o__Bacteroidales;f__Bacteroidaceae;g__Bacteroides | 0.922140617 | 0.373105981 | 0.458736862 | HM1->KO |
| d__Bacteria;p__Firmicutes;c__Clostridia;o__Oscillospirales;f__Ruminococcaceae;g__uncultured | -0.845682315 | 0.410382339 | 0.496430248 | HM1->WT |
| d__Bacteria;p__Firmicutes;c__Clostridia;o__Lachnospirales;f__Lachnospiraceae;g__Hungatella | 0.81083642 | 0.427050135 | 0.508393018 | HM1->KO |
| d__Bacteria;p__Firmicutes;c__Clostridia;o__Lachnospirales;f__Lachnospiraceae;g__Agathobacter | -0.801601825 | 0.440987563 | 0.512652274 | HM1->WT |
| d__Bacteria;p__Firmicutes;c__Clostridia;o__Oscillospirales;f__Oscillospiraceae;g__Colidextribacter | -0.782595773 | 0.444298637 | 0.512652274 | HM1->WT |
| Unassigned;__;__;__;__;__ | -0.751948976 | 0.466089744 | 0.529647436 | HM1->WT |
| d__Bacteria;p__Actinobacteriota;c__Actinobacteria;o__Bifidobacteriales;f__Bifidobacteriaceae;g__Bifidobacterium | -0.718439805 | 0.487012031 | 0.545162722 | HM1->WT |
| d__Bacteria;p__Firmicutes;c__Bacilli;o__Lactobacillales;f__Enterococcaceae;g__Enterococcus | -0.558138604 | 0.583017737 | 0.643034269 | HM1->WT |
| d__Bacteria;p__Firmicutes;c__Clostridia;o__Oscillospirales;f__[Clostridium]_methylpentosum_group;g__[Clostridium]_methylpentosum_group | 0.534227501 | 0.60256167 | 0.654958337 | HM1->KO |
| d__Bacteria;p__Firmicutes;c__Clostridia;o__Oscillospirales;f__Ruminococcaceae;g__Candidatus_Soleaferrea | 0.38705943 | 0.706997445 | 0.757497262 | HM1->KO |
| d__Bacteria;p__Firmicutes;c__Clostridia;o__Clostridia_vadinBB60_group;f__Clostridia_vadinBB60_group;g__Clostridia_vadinBB60_group | -0.240589576 | 0.813879015 | 0.859731354 | HM1->WT |
| d__Bacteria;p__Firmicutes;c__Clostridia;o__Peptostreptococcales-Tissierellales;f__Anaerovoracaceae;g__[Eubacterium]_nodatum_group | 0.188868185 | 0.853371495 | 0.888928641 | HM1->KO |
| d__Bacteria;p__Firmicutes;c__Clostridia;o__Oscillospirales;f__Oscillospiraceae;g__Flavonifractor | 0.167986622 | 0.869748875 | 0.893577612 | HM1->KO |
| d__Bacteria;p__Firmicutes;c__Bacilli;o__Erysipelotrichales;f__Erysipelotrichaceae;g__Holdemania | -0.045494883 | 0.964414277 | 0.977446902 | HM1->WT |
| d__Bacteria;p__Firmicutes;c__Clostridia;o__Oscillospirales;f__Ruminococcaceae;__ | -0.023778803 | 0.981520909 | 0.981520909 | HM1->WT |

| Operational Taxonomic Unit (Genus) for HM1->KO vs HM2->KO Comparison | T-value | P-value | FDR | Enriched |
|---|---|---|---|---|
| d__Bacteria;p__Firmicutes;c__Clostridia;o__Peptostreptococcales-Tissierellales;f__Peptostreptococcaceae;g__Clostridioides | 14.26313282 | 2.12E-11 | 1.22E-09 | HM1->KO |
| d__Bacteria;p__Bacteroidota;c__Bacteroidia;o__Bacteroidales;f__Bacteroidaceae;g__Bacteroides | -17.87030689 | 3.16E-11 | 1.22E-09 | HM2->KO |
| d__Bacteria;p__Bacteroidota;c__Bacteroidia;o__Bacteroidales;f__Marinifilaceae;g__Odoribacter | -24.80141475 | 2.60E-10 | 6.66E-09 | HM2->KO |
| d__Bacteria;p__Fusobacteriota;c__Fusobacteriia;o__Fusobacteriales;f__Fusobacteriaceae;g__Fusobacterium | -19.0423166 | 3.47E-09 | 6.67E-08 | HM2->KO |
| d__Bacteria;p__Firmicutes;c__Clostridia;o__Clostridiales;f__Clostridiaceae;g__Clostridium_sensu_stricto_1 | 9.98485579 | 7.17E-08 | 1.10E-06 | HM1->KO |
| d__Bacteria;p__Firmicutes;c__Clostridia;o__Oscillospirales;f__Ruminococcaceae;g__Candidatus_Soleaferrea | 6.972448656 | 5.35E-07 | 6.86E-06 | HM1->KO |
| d__Bacteria;p__Firmicutes;c__Clostridia;o__Lachnospirales;f__Lachnospiraceae;g__Epulopiscium | 7.741674625 | 9.37E-07 | 1.03E-05 | HM1->KO |
| d__Bacteria;p__Bacteroidota;c__Bacteroidia;o__Bacteroidales;f__Tannerellaceae;__ | -9.818312334 | 1.88E-06 | 1.81E-05 | HM2->KO |
| d__Bacteria;p__Firmicutes;c__Clostridia;o__Lachnospirales;f__Lachnospiraceae;g__Eisenbergiella | 5.781160253 | 1.32E-05 | 0.000112899 | HM1->KO |
| Unassigned;__;__;__;__;__ | 4.912539491 | 0.000141111 | 0.001086554 | HM1->KO |
| d__Bacteria;p__Firmicutes;c__Bacilli;o__Erysipelotrichales;f__Erysipelotrichaceae;g__Holdemania | -4.206661286 | 0.000652422 | 0.004566953 | HM2->KO |
| d__Bacteria;p__Firmicutes;c__Clostridia;o__Oscillospirales;f__Ruminococcaceae;__ | -4.07876278 | 0.00095628 | 0.005808614 | HM2->KO |
| d__Bacteria;p__Firmicutes;c__Clostridia;o__Peptostreptococcales-Tissierellales;f__Anaerovoracaceae;g__[Eubacterium]_nodatum_group | 3.753911257 | 0.000980675 | 0.005808614 | HM1->KO |
| d__Bacteria;p__Firmicutes;c__Clostridia;o__Clostridiales;f__Clostridiaceae;g__Clostridium_sensu_stricto_13 | 3.523369649 | 0.001745827 | 0.009602049 | HM1->KO |
| d__Bacteria;p__Actinobacteriota;c__Coriobacteriia;o__Coriobacteriales;f__Eggerthellaceae;g__Eggerthella | 3.580994299 | 0.002021722 | 0.010378173 | HM1->KO |
| d__Bacteria;p__Bacteroidota;c__Bacteroidia;o__Bacteroidales;f__Tannerellaceae;g__Parabacteroides | -3.993899197 | 0.002543243 | 0.012239356 | HM2->KO |
| d__Bacteria;p__Firmicutes;c__Clostridia;o__Oscillospirales;f__Ruminococcaceae;g__UBA1819 | 3.401552663 | 0.002999929 | 0.013587914 | HM1->KO |
| d__Bacteria;p__Firmicutes;c__Clostridia;o__Oscillospirales;f__Oscillospiraceae;g__Oscillibacter | -3.18731025 | 0.004157136 | 0.017783306 | HM2->KO |
| d__Bacteria;p__Firmicutes;c__Bacilli;o__Erysipelotrichales;f__Erysipelatoclostridiaceae;g__Erysipelatoclostridium | 3.611739416 | 0.004507559 | 0.018267476 | HM1->KO |
| d__Bacteria;p__Firmicutes;c__Clostridia;o__Lachnospirales;f__Lachnospiraceae;g__[Ruminococcus]_gnavus_group | 2.944066808 | 0.011738644 | 0.045193781 | HM1->KO |
| d__Bacteria;p__Firmicutes;c__Clostridia;o__Lachnospirales;f__Lachnospiraceae;g__Lactonifactor | 2.81786847 | 0.013690093 | 0.050197007 | HM1->KO |
| d__Bacteria;p__Firmicutes;c__Clostridia;o__Lachnospirales;f__Lachnospiraceae;__ | -2.506777523 | 0.020257547 | 0.070731972 | HM2->KO |
| d__Bacteria;p__Firmicutes;c__Bacilli;o__Lactobacillales;f__Streptococcaceae;g__Streptococcus | 2.596328893 | 0.021127732 | 0.070731972 | HM1->KO |
| d__Bacteria;p__Firmicutes;c__Clostridia;o__Lachnospirales;f__Lachnospiraceae;g__[Ruminococcus]_torques_group | -2.211960625 | 0.039284883 | 0.126039001 | HM2->KO |
| d__Bacteria;p__Firmicutes;c__Clostridia;o__Lachnospirales;f__Lachnospiraceae;g__uncultured | 2.18733172 | 0.046183188 | 0.139338884 | HM1->KO |
| d__Bacteria;p__Verrucomicrobiota;c__Verrucomicrobiae;o__Verrucomicrobiales;f__Akkermansiaceae;g__Akkermansia | 2.162106428 | 0.047049493 | 0.139338884 | HM1->KO |
| d__Bacteria;p__Actinobacteriota;c__Coriobacteriia;o__Coriobacteriales;f__Coriobacteriaceae;g__Collinsella | -2.053617019 | 0.051198604 | 0.146010833 | HM2->KO |
| d__Bacteria;p__Firmicutes;c__Clostridia;o__Oscillospirales;f__[Eubacterium]_coprostanoligenes_group;g__[Eubacterium]_coprostanoligenes_group | 2.024729773 | 0.055034767 | 0.147884295 | HM1->KO |
| d__Bacteria;p__Firmicutes;c__Clostridia;o__Lachnospirales;f__Lachnospiraceae;g__Hungatella | 2.043281969 | 0.055696682 | 0.147884295 | HM1->KO |
| d__Bacteria;p__Actinobacteriota;c__Coriobacteriia;o__Coriobacteriales;f__Eggerthellaceae;g__Gordonibacter | -1.839002148 | 0.08015382 | 0.205728139 | HM2->KO |
| d__Bacteria;p__Firmicutes;c__Clostridia;o__Lachnospirales;f__Lachnospiraceae;g__Agathobacter | -1.819699174 | 0.087098347 | 0.216341054 | HM2->KO |
| d__Bacteria;p__Firmicutes;c__Clostridia;o__Oscillospirales;f__Oscillospiraceae;g__Colidextribacter | -1.729349121 | 0.098488302 | 0.236987477 | HM2->KO |
| d__Bacteria;p__Firmicutes;c__Clostridia;o__Oscillospirales;f__Ruminococcaceae;g__Paludicola | -1.659269314 | 0.110080716 | 0.256855005 | HM2->KO |
| d__Bacteria;p__Firmicutes;c__Clostridia;o__Oscillospirales;f__Ruminococcaceae;g__uncultured | 1.574261347 | 0.128746442 | 0.291572824 | HM1->KO |
| d__Bacteria;p__Firmicutes;c__Clostridia;o__Lachnospirales;f__Lachnospiraceae;g__Lachnospiraceae | 1.507614561 | 0.1473003 | 0.32406066 | HM1->KO |
| d__Bacteria;p__Firmicutes;c__Clostridia;o__Lachnospirales;f__Lachnospiraceae;g__Sellimonas | -1.477179344 | 0.159100682 | 0.340298681 | HM2->KO |
| d__Bacteria;p__Firmicutes;c__Clostridia;o__Lachnospirales;f__Lachnospiraceae;g__[Eubacterium]_fissicatena_group | 1.466955106 | 0.164490455 | 0.342317975 | HM1->KO |
| d__Bacteria;p__Firmicutes;c__Clostridia;o__Oscillospirales;f__Oscillospiraceae;g__UCG-005 | 1.396045647 | 0.17548393 | 0.355585857 | HM1->KO |
| d__Bacteria;p__Firmicutes;c__Clostridia;o__Lachnospirales;f__Lachnospiraceae;g__Blautia | -1.369527934 | 0.185884527 | 0.367002783 | HM2->KO |
| d__Bacteria;p__Firmicutes;c__Clostridia;o__Monoglobales;f__Monoglobaceae;g__Monoglobus | -1.350744801 | 0.193417568 | 0.372328818 | HM2->KO |
| d__Bacteria;p__Firmicutes;c__Clostridia;o__Lachnospirales;f__Lachnospiraceae;g__Lachnospiraceae_UCG-008 | -1.325382297 | 0.20384263 | 0.382826402 | HM2->KO |
| d__Bacteria;p__Firmicutes;c__Clostridia;o__Oscillospirales;f__[Clostridium]_methylpentosum_group;g__[Clostridium]_methylpentosum_group | 1.219282986 | 0.234662247 | 0.430214119 | HM1->KO |
| d__Bacteria;p__Firmicutes;c__Clostridia;o__Oscillospirales;f__Ruminococcaceae;g__Negativibacillus | 1.069382956 | 0.296420825 | 0.522611967 | HM1->KO |
| d__Bacteria;p__Proteobacteria;c__Gammaproteobacteria;o__Vibrionales;f__Vibrionaceae;g__Vibrio | 1.064675611 | 0.29863541 | 0.522611967 | HM1->KO |
| d__Bacteria;p__Firmicutes;c__Clostridia;o__Lachnospirales;f__Lachnospiraceae;g__Roseburia | 1 | 0.334281943 | 0.551863394 | HM1->KO |
| d__Bacteria;p__Firmicutes;c__Clostridia;o__Clostridia_vadinBB60_group;f__Clostridia_vadinBB60_group;g__Clostridia_vadinBB60_group | 1 | 0.334281943 | 0.551863394 | HM1->KO |
| d__Bacteria;p__Firmicutes;c__Clostridia;o__Lachnospirales;f__Lachnospiraceae;g__Marvinbryantia | -0.984753599 | 0.339523669 | 0.551863394 | HM2->KO |
| d__Bacteria;p__Firmicutes;c__Clostridia;o__Lachnospirales;f__Lachnospiraceae;g__Lachnospiraceae_NK4A136_group | 0.963508718 | 0.34541735 | 0.551863394 | HM1->KO |
| d__Bacteria;__;__;__;__;__ | -0.966990889 | 0.351185796 | 0.551863394 | HM2->KO |
| d__Bacteria;p__Firmicutes;c__Clostridia;o__Oscillospirales;f__Ruminococcaceae;g__Anaerotruncus | 0.906254736 | 0.374275676 | 0.576384541 | HM1->KO |
| d__Bacteria;p__Bacteroidota;c__Bacteroidia;o__Bacteroidales;f__Rikenellaceae;g__Alistipes | -0.864746183 | 0.400222641 | 0.604257713 | HM2->KO |
| d__Bacteria;p__Firmicutes;c__Clostridia;o__Peptococcales;f__Peptococcaceae;g__uncultured | -0.841442192 | 0.409222706 | 0.605964392 | HM2->KO |
| d__Bacteria;p__Firmicutes;c__Clostridia;o__Oscillospirales;f__Ruminococcaceae;g__Incertae_Sedis | 0.804593342 | 0.431754047 | 0.627265313 | HM1->KO |
| d__Bacteria;p__Firmicutes;c__uncultured;o__uncultured;f__uncultured;g__uncultured | -0.780443252 | 0.449866516 | 0.641476328 | HM2->KO |
| d__Bacteria;p__Firmicutes;c__Bacilli;o__Erysipelotrichales;f__Erysipelotrichaceae;g__[Clostridium]_innocuum_group | -0.742246601 | 0.46854053 | 0.644829222 | HM2->KO |
| d__Bacteria;p__Actinobacteriota;c__Coriobacteriia;o__Coriobacteriales;f__Eggerthellaceae;__ | -0.740719761 | 0.468966707 | 0.644829222 | HM2->KO |
| d__Bacteria;p__Firmicutes;c__Clostridia;o__Oscillospirales;f__Ruminococcaceae;g__Subdoligranulum | 0.700012291 | 0.490907692 | 0.663156005 | HM1->KO |
| d__Bacteria;p__Firmicutes;c__Clostridia;o__Lachnospirales;f__Lachnospiraceae;g__GCA-900066755 | 0.666059371 | 0.511993147 | 0.67971504 | HM1->KO |
| d__Bacteria;p__Firmicutes;c__Bacilli;o__Erysipelotrichales;f__Erysipelotrichaceae;g__Turicibacter | -0.633106475 | 0.535119186 | 0.698375887 | HM2->KO |
| d__Bacteria;p__Firmicutes;c__Clostridia;o__Lachnospirales;f__Lachnospiraceae;g__Anaerostipes | 0.608866911 | 0.552139132 | 0.703269348 | HM1->KO |
| d__Bacteria;p__Firmicutes;c__Bacilli;o__Erysipelotrichales;f__Erysipelotrichaceae;g__Dielma | 0.59034896 | 0.562664997 | 0.703269348 | HM1->KO |
| d__Bacteria;p__Firmicutes;c__Clostridia;o__Lachnospirales;f__Lachnospiraceae;g__GCA-900066575 | 0.582673928 | 0.566268826 | 0.703269348 | HM1->KO |
| d__Bacteria;p__Firmicutes;c__Clostridia;o__Lachnospirales;f__Lachnospiraceae;g__Lachnoclostridium | -0.567306575 | 0.57583686 | 0.703800606 | HM2->KO |
| d__Bacteria;p__Firmicutes;c__Clostridia;o__Peptostreptococcales-Tissierellales;f__Anaerovoracaceae;g__Family_XIII_AD3011_group | 0.553722574 | 0.586721754 | 0.705899611 | HM1->KO |
| d__Bacteria;p__Firmicutes;c__Bacilli;o__Staphylococcales;f__Staphylococcaceae;g__Staphylococcus | -0.489944864 | 0.630196466 | 0.746540429 | HM2->KO |
| d__Bacteria;p__Firmicutes;c__Clostridia;o__Oscillospirales;f__Ruminococcaceae;g__Faecalibacterium | -0.46916379 | 0.644137535 | 0.74889516 | HM2->KO |
| d__Bacteria;p__Proteobacteria;c__Gammaproteobacteria;o__Enterobacterales;f__Enterobacteriaceae;g__Escherichia-Shigella | 0.457353246 | 0.651636049 | 0.74889516 | HM1->KO |
| d__Bacteria;p__Actinobacteriota;c__Coriobacteriia;o__Coriobacteriales;f__Eggerthellaceae;g__Adlercreutzia | -0.401153298 | 0.692283737 | 0.783909525 | HM2->KO |
| d__Bacteria;p__Firmicutes;c__Clostridia;o__Oscillospirales;f__Butyricicoccaceae;g__Butyricicoccus | 0.367034373 | 0.717066592 | 0.79399903 | HM1->KO |
| d__Bacteria;p__Actinobacteriota;c__Actinobacteria;o__Bifidobacteriales;f__Bifidobacteriaceae;g__Bifidobacterium | -0.34049406 | 0.736486169 | 0.79399903 | HM2->KO |
| d__Bacteria;p__Firmicutes;c__Clostridia;o__Oscillospirales;f__Ruminococcaceae;g__Ruminococcus | -0.338612796 | 0.73786616 | 0.79399903 | HM2->KO |
| d__Bacteria;p__Firmicutes;c__Clostridia;o__Oscillospirales;f__Oscillospiraceae;g__Flavonifractor | 0.332612266 | 0.742440651 | 0.79399903 | HM1->KO |
| d__Bacteria;p__Firmicutes;c__Bacilli;o__Lactobacillales;f__Enterococcaceae;g__Enterococcus | 0.282293763 | 0.781292642 | 0.824103198 | HM1->KO |
| d__Bacteria;p__Firmicutes;c__Clostridia;o__Peptostreptococcales-Tissierellales;f__Peptostreptococcaceae;g__Terrisporobacter | -0.245466845 | 0.80845691 | 0.832217349 | HM2->KO |
| d__Bacteria;p__Firmicutes;c__Clostridia;o__Oscillospirales;f__Oscillospiraceae;g__uncultured | 0.242519985 | 0.810601314 | 0.832217349 | HM1->KO |
| d__Bacteria;p__Firmicutes;c__Clostridia;o__Oscillospirales;f__Ruminococcaceae;g__Phocea | -0.205150521 | 0.839295404 | 0.850338765 | HM2->KO |
| d__Bacteria;p__Proteobacteria;c__Gammaproteobacteria;o__Burkholderiales;f__Sutterellaceae;g__Parasutterella | -0.140368696 | 0.889590594 | 0.889590594 | HM2->KO |

| Operational Taxonomic Unit (Genus) for HM1->KO vs IMM-g1->KO Comparison | T-value | P-value | FDR | Enriched |
|---|---|---|---|---|
| d__Bacteria;p__Firmicutes;c__Clostridia;o__Peptostreptococcales-Tissierellales;f__Peptostreptococcaceae;g__Clostridioides | -3.856040953 | 0.00089122 | 0.03917372 | IMM-g1->KO |
| d__Bacteria;p__Firmicutes;c__Clostridia;o__Lachnospirales;f__Lachnospiraceae;g__Lactonifactor | -3.686744698 | 0.001044633 | 0.03917372 | IMM-g1->KO |
| d__Bacteria;p__Firmicutes;c__Clostridia;o__Lachnospirales;f__Lachnospiraceae;g__Epulopiscium | 3.349822035 | 0.00213719 | 0.053429747 | HM1->KO |
| d__Bacteria;p__Firmicutes;c__Clostridia;o__Lachnospirales;f__Lachnospiraceae;g__[Ruminococcus]_gnavus_group | 2.8058424 | 0.009057392 | 0.169826099 | HM1->KO |
| d__Bacteria;p__Bacteroidota;c__Bacteroidia;o__Bacteroidales;f__Rikenellaceae;g__Alistipes | -2.71838275 | 0.015717848 | 0.235767713 | IMM-g1->KO |
| d__Bacteria;p__Firmicutes;c__Clostridia;o__Oscillospirales;f__Oscillospiraceae;g__UCG-005 | 2.41279103 | 0.024888774 | 0.247332915 | HM1->KO |
| d__Bacteria;p__Firmicutes;c__Bacilli;o__Lactobacillales;f__Streptococcaceae;g__Streptococcus | 2.495832846 | 0.02543905 | 0.247332915 | HM1->KO |
| d__Bacteria;p__Firmicutes;c__Clostridia;o__Oscillospirales;f__Ruminococcaceae;g__Ruminococcus | 2.415175635 | 0.029506351 | 0.247332915 | HM1->KO |
| d__Bacteria;p__Firmicutes;c__Clostridia;o__Peptococcales;f__Peptococcaceae;g__uncultured | 2.295103433 | 0.032188589 | 0.247332915 | HM1->KO |
| d__Bacteria;p__Firmicutes;c__Clostridia;o__Oscillospirales;f__[Clostridium]_methylpentosum_group;g__[Clostridium]_methylpentosum_group | -2.234990296 | 0.032977722 | 0.247332915 | IMM-g1->KO |
| d__Bacteria;p__Proteobacteria;c__Gammaproteobacteria;o__Vibrionales;f__Vibrionaceae;g__Vibrio | 2.12133112 | 0.043517013 | 0.257871953 | HM1->KO |
| d__Bacteria;p__Firmicutes;c__Clostridia;o__Lachnospirales;f__Lachnospiraceae;g__uncultured | 2.18733172 | 0.046183188 | 0.257871953 | HM1->KO |
| d__Bacteria;p__Firmicutes;c__Clostridia;o__Lachnospirales;f__Lachnospiraceae;__ | 2.096382452 | 0.047768137 | 0.257871953 | HM1->KO |
| d__Bacteria;p__Firmicutes;c__Clostridia;o__Clostridiales;f__Clostridiaceae;g__Clostridium_sensu_stricto_13 | -2.078827973 | 0.048136098 | 0.257871953 | IMM-g1->KO |
| d__Bacteria;p__Firmicutes;c__Clostridia;o__Oscillospirales;f__Ruminococcaceae;g__uncultured | 2.0392844 | 0.051884758 | 0.259423792 | HM1->KO |
| d__Bacteria;p__Proteobacteria;c__Gammaproteobacteria;o__Enterobacterales;f__Enterobacteriaceae;g__Escherichia-Shigella | -2.004925076 | 0.057984473 | 0.271802217 | IMM-g1->KO |
| d__Bacteria;p__Firmicutes;c__Bacilli;o__Erysipelotrichales;f__Erysipelotrichaceae;g__Turicibacter | 1.912877331 | 0.074052439 | 0.326701937 | HM1->KO |
| d__Bacteria;p__Firmicutes;c__Clostridia;o__Oscillospirales;f__Ruminococcaceae;g__Subdoligranulum | 1.863142456 | 0.083554869 | 0.343067686 | HM1->KO |
| d__Bacteria;p__Firmicutes;c__Clostridia;o__Oscillospirales;f__Ruminococcaceae;__ | 1.840993078 | 0.086910481 | 0.343067686 | HM1->KO |
| d__Bacteria;p__Firmicutes;c__Clostridia;o__Oscillospirales;f__Ruminococcaceae;g__Faecalibacterium | 1.668124266 | 0.116502524 | 0.436884464 | HM1->KO |
| d__Bacteria;p__Firmicutes;c__Clostridia;o__Peptostreptococcales-Tissierellales;f__Peptostreptococcaceae;g__Romboutsia | -1.565815034 | 0.135815799 | 0.485056424 | IMM-g1->KO |
| d__Bacteria;p__Firmicutes;c__Bacilli;o__Staphylococcales;f__Staphylococcaceae;g__Staphylococcus | 1.459459783 | 0.15919922 | 0.499727514 | HM1->KO |
| d__Bacteria;__;__;__;__;__ | -1.433571826 | 0.163828866 | 0.499727514 | IMM-g1->KO |
| d__Bacteria;p__Firmicutes;c__uncultured;o__uncultured;f__uncultured;g__uncultured | 1.467462799 | 0.164354338 | 0.499727514 | HM1->KO |
| d__Bacteria;p__Firmicutes;c__Clostridia;o__Lachnospirales;f__Lachnospiraceae;g__Marvinbryantia | 1.459220545 | 0.166575838 | 0.499727514 | HM1->KO |
| d__Bacteria;p__Firmicutes;c__Bacilli;o__Erysipelotrichales;f__Erysipelotrichaceae;g__Holdemania | -1.346325213 | 0.189712286 | 0.530431814 | IMM-g1->KO |
| d__Bacteria;p__Firmicutes;c__Clostridia;o__Oscillospirales;f__Ruminococcaceae;g__Paludicola | 1.3398498 | 0.190955453 | 0.530431814 | HM1->KO |
| d__Bacteria;p__Firmicutes;c__Clostridia;o__Oscillospirales;f__Oscillospiraceae;g__Colidextribacter | 1.297912214 | 0.20802158 | 0.55720066 | HM1->KO |
| d__Bacteria;p__Firmicutes;c__Clostridia;o__Clostridiales;f__Clostridiaceae;g__Clostridium_sensu_stricto_1 | -1.238615385 | 0.225586803 | 0.561116334 | IMM-g1->KO |
| d__Bacteria;p__Firmicutes;c__Clostridia;o__Oscillospirales;f__Oscillospiraceae;g__Oscillibacter | 1.23432051 | 0.226359055 | 0.561116334 | HM1->KO |
| d__Bacteria;p__Firmicutes;c__Bacilli;o__Lactobacillales;f__Enterococcaceae;g__Enterococcus | -1.204916202 | 0.239896047 | 0.561116334 | IMM-g1->KO |
| Unassigned;__;__;__;__;__ | 1.197899689 | 0.240301084 | 0.561116334 | HM1->KO |
| d__Bacteria;p__Actinobacteriota;c__Coriobacteriia;o__Coriobacteriales;f__Coriobacteriaceae;g__Collinsella | 1.170917151 | 0.252524226 | 0.561116334 | HM1->KO |
| d__Bacteria;p__Firmicutes;c__Clostridia;o__Lachnospirales;f__Lachnospiraceae;g__Blautia | 1.132584264 | 0.266269756 | 0.561116334 | HM1->KO |
| d__Bacteria;p__Actinobacteriota;c__Coriobacteriia;o__Coriobacteriales;f__Eggerthellaceae;g__Adlercreutzia | 1.121524959 | 0.272226554 | 0.561116334 | HM1->KO |
| d__Bacteria;p__Firmicutes;c__Clostridia;o__Oscillospirales;f__Ruminococcaceae;g__UBA1819 | 1.107937942 | 0.276457817 | 0.561116334 | HM1->KO |
| d__Bacteria;p__Firmicutes;c__Clostridia;o__Monoglobales;f__Monoglobaceae;g__Monoglobus | 1.10972619 | 0.276817392 | 0.561116334 | HM1->KO |
| d__Bacteria;p__Bacteroidota;c__Bacteroidia;o__Bacteroidales;f__Tannerellaceae;g__Parabacteroides | -1 | 0.331332762 | 0.583081486 | IMM-g1->KO |
| d__Bacteria;p__Firmicutes;c__Clostridia;o__Lachnospirales;f__Lachnospiraceae;g__Agathobacter | 1 | 0.334281943 | 0.583081486 | HM1->KO |
| d__Bacteria;p__Firmicutes;c__Clostridia;o__Lachnospirales;f__Lachnospiraceae;g__Roseburia | 1 | 0.334281943 | 0.583081486 | HM1->KO |
| d__Bacteria;p__Actinobacteriota;c__Coriobacteriia;o__Coriobacteriales;f__Eggerthellaceae;__ | 1 | 0.334281943 | 0.583081486 | HM1->KO |
| d__Bacteria;p__Firmicutes;c__Clostridia;o__Clostridia_vadinBB60_group;f__Clostridia_vadinBB60_group;g__Clostridia_vadinBB60_group | 1 | 0.334281943 | 0.583081486 | HM1->KO |
| d__Bacteria;p__Firmicutes;c__Clostridia;o__Peptostreptococcales-Tissierellales;f__Anaerovoracaceae;g__[Eubacterium]_nodatum_group | -0.981186609 | 0.334300052 | 0.583081486 | IMM-g1->KO |
| d__Bacteria;p__Actinobacteriota;c__Coriobacteriia;o__Coriobacteriales;f__Eggerthellaceae;g__Gordonibacter | 0.960157989 | 0.344414235 | 0.587069719 | HM1->KO |
| d__Bacteria;p__Firmicutes;c__Clostridia;o__Oscillospirales;f__Ruminococcaceae;g__Incertae_Sedis | 0.89039153 | 0.384120226 | 0.6389558 | HM1->KO |
| d__Bacteria;p__Firmicutes;c__Clostridia;o__Lachnospirales;f__Lachnospiraceae;g__GCA-900066575 | 0.868640425 | 0.391892891 | 0.6389558 | HM1->KO |
| d__Bacteria;p__Firmicutes;c__Bacilli;o__Erysipelotrichales;f__Erysipelotrichaceae;g__[Clostridium]_innocuum_group | 0.824635642 | 0.417910413 | 0.666878318 | HM1->KO |
| d__Bacteria;p__Firmicutes;c__Clostridia;o__Oscillospirales;f__Oscillospiraceae;g__Flavonifractor | -0.801893401 | 0.428813778 | 0.670021528 | IMM-g1->KO |
| d__Bacteria;p__Firmicutes;c__Clostridia;o__Lachnospirales;f__Lachnospiraceae;g__Anaerostipes | 0.772055967 | 0.446438745 | 0.678917518 | HM1->KO |
| d__Bacteria;p__Firmicutes;c__Clostridia;o__Lachnospirales;f__Lachnospiraceae;g__Lachnospiraceae | -0.756452971 | 0.455091636 | 0.678917518 | IMM-g1->KO |
| d__Bacteria;p__Firmicutes;c__Clostridia;o__Lachnospirales;f__Lachnospiraceae;g__GCA-900066755 | -0.715234254 | 0.48028339 | 0.678917518 | IMM-g1->KO |
| d__Bacteria;p__Firmicutes;c__Clostridia;o__Lachnospirales;f__Lachnospiraceae;g__Sellimonas | -0.710072501 | 0.486101793 | 0.678917518 | IMM-g1->KO |
| d__Bacteria;p__Firmicutes;c__Clostridia;o__Oscillospirales;f__Ruminococcaceae;g__Anaerotruncus | -0.696298354 | 0.491705112 | 0.678917518 | IMM-g1->KO |
| d__Bacteria;p__Firmicutes;c__Clostridia;o__Lachnospirales;f__Lachnospiraceae;g__[Eubacterium]_fissicatena_group | 0.691887651 | 0.495501591 | 0.678917518 | HM1->KO |
| d__Bacteria;p__Verrucomicrobiota;c__Verrucomicrobiae;o__Verrucomicrobiales;f__Akkermansiaceae;g__Akkermansia | -0.685908423 | 0.497872847 | 0.678917518 | IMM-g1->KO |
| d__Bacteria;p__Firmicutes;c__Clostridia;o__Oscillospirales;f__Oscillospiraceae;g__uncultured | 0.656303268 | 0.516658962 | 0.691953968 | HM1->KO |
| d__Bacteria;p__Firmicutes;c__Clostridia;o__Peptostreptococcales-Tissierellales;f__Peptostreptococcaceae;g__Terrisporobacter | -0.581534204 | 0.565091832 | 0.743541884 | IMM-g1->KO |
| d__Bacteria;p__Firmicutes;c__Clostridia;o__Oscillospirales;f__[Eubacterium]_coprostanoligenes_group;g__[Eubacterium]_coprostanoligenes_group | -0.551821834 | 0.585311965 | 0.750118692 | IMM-g1->KO |
| d__Bacteria;p__Firmicutes;c__Clostridia;o__Lachnospirales;f__Lachnospiraceae;g__[Ruminococcus]_torques_group | 0.531959872 | 0.598667293 | 0.750118692 | HM1->KO |
| d__Bacteria;p__Firmicutes;c__Clostridia;o__Lachnospirales;f__Lachnospiraceae;g__Eisenbergiella | 0.512590536 | 0.611999985 | 0.750118692 | HM1->KO |
| d__Bacteria;p__Firmicutes;c__Clostridia;o__Lachnospirales;f__Lachnospiraceae;g__Hungatella | -0.51164699 | 0.613615223 | 0.750118692 | IMM-g1->KO |
| d__Bacteria;p__Proteobacteria;c__Gammaproteobacteria;o__Burkholderiales;f__Sutterellaceae;g__Parasutterella | -0.497956441 | 0.622113434 | 0.750118692 | IMM-g1->KO |
| d__Bacteria;p__Firmicutes;c__Clostridia;o__Lachnospirales;f__Lachnospiraceae;g__Lachnospiraceae_UCG-008 | 0.486821294 | 0.630099701 | 0.750118692 | HM1->KO |
| d__Bacteria;p__Firmicutes;c__Clostridia;o__Oscillospirales;f__Ruminococcaceae;g__Negativibacillus | 0.471068794 | 0.641004997 | 0.751177731 | HM1->KO |
| d__Bacteria;p__Actinobacteriota;c__Coriobacteriia;o__Coriobacteriales;f__Eggerthellaceae;g__Eggerthella | 0.410292429 | 0.684708294 | 0.790048031 | HM1->KO |
| d__Bacteria;p__Firmicutes;c__Clostridia;o__Oscillospirales;f__Ruminococcaceae;g__Candidatus_Soleaferrea | -0.379858862 | 0.706709436 | 0.803078905 | IMM-g1->KO |
| d__Bacteria;p__Actinobacteriota;c__Actinobacteria;o__Bifidobacteriales;f__Bifidobacteriaceae;g__Bifidobacterium | -0.324069667 | 0.748403481 | 0.83776509 | IMM-g1->KO |
| d__Bacteria;p__Firmicutes;c__Clostridia;o__Lachnospirales;f__Lachnospiraceae;g__Lachnospiraceae_NK4A136_group | 0.275979662 | 0.784557567 | 0.865320846 | HM1->KO |
| d__Bacteria;p__Bacteroidota;c__Bacteroidia;o__Bacteroidales;f__Bacteroidaceae;g__Bacteroides | -0.183268014 | 0.855826342 | 0.930246024 | IMM-g1->KO |
| d__Bacteria;p__Firmicutes;c__Bacilli;o__Erysipelotrichales;f__Erysipelatoclostridiaceae;g__Erysipelatoclostridium | 0.104038163 | 0.917857235 | 0.969016661 | HM1->KO |
| d__Bacteria;p__Firmicutes;c__Clostridia;o__Oscillospirales;f__Ruminococcaceae;g__Phocea | -0.102947512 | 0.918775047 | 0.969016661 | IMM-g1->KO |
| d__Bacteria;p__Firmicutes;c__Bacilli;o__Erysipelotrichales;f__Erysipelotrichaceae;g__Dielma | 0.085214168 | 0.932742771 | 0.969016661 | HM1->KO |
| d__Bacteria;p__Firmicutes;c__Clostridia;o__Oscillospirales;f__Butyricicoccaceae;g__Butyricicoccus | -0.071948258 | 0.943176217 | 0.969016661 | IMM-g1->KO |
| d__Bacteria;p__Firmicutes;c__Clostridia;o__Peptostreptococcales-Tissierellales;f__Anaerovoracaceae;g__Family_XIII_AD3011_group | -0.032997316 | 0.973906877 | 0.984360086 | IMM-g1->KO |
| d__Bacteria;p__Firmicutes;c__Clostridia;o__Lachnospirales;f__Lachnospiraceae;g__Lachnoclostridium | 0.019761961 | 0.984360086 | 0.984360086 | HM1->KO |

| Operational Taxonomic Unit (Genus) for IMM-g1->KO vs IMM-g2->KO Comparison | T-value | P-value | FDR | Enriched |
|---|---|---|---|---|
| d__Bacteria;p__Firmicutes;c__Clostridia;o__Oscillospirales;f__Oscillospiraceae;g__Oscillibacter | 2.891090917 | 0.012901144 | 0.602381974 | IMM-g1->KO |
| d__Bacteria;p__Firmicutes;c__Clostridia;o__Lachnospirales;f__Lachnospiraceae;g__[Ruminococcus]_gnavus_group | -2.517319021 | 0.019093938 | 0.602381974 | IMM-g2->KO |
| d__Bacteria;p__Actinobacteriota;c__Actinobacteria;o__Bifidobacteriales;f__Bifidobacteriaceae;g__Bifidobacterium | 2.269032686 | 0.03254679 | 0.602381974 | IMM-g1->KO |
| d__Bacteria;p__Firmicutes;c__Clostridia;o__Lachnospirales;f__Lachnospiraceae;g__Epulopiscium | 2.194877388 | 0.039244245 | 0.602381974 | IMM-g1->KO |
| d__Bacteria;p__Firmicutes;c__Clostridia;o__Peptostreptococcales-Tissierellales;f__Peptostreptococcaceae;g__Terrisporobacter | 2.014395302 | 0.05655272 | 0.602381974 | IMM-g1->KO |
| d__Bacteria;p__Firmicutes;c__Clostridia;o__Oscillospirales;f__Butyricicoccaceae;g__Butyricicoccus | -1.998559223 | 0.058189089 | 0.602381974 | IMM-g2->KO |
| d__Bacteria;p__Firmicutes;c__Clostridia;o__Oscillospirales;f__Oscillospiraceae;g__Colidextribacter | -1.943802056 | 0.06587104 | 0.602381974 | IMM-g2->KO |
| d__Bacteria;p__Firmicutes;c__Clostridia;o__Oscillospirales;f__Ruminococcaceae;g__Negativibacillus | -1.917348644 | 0.070868467 | 0.602381974 | IMM-g2->KO |
| d__Bacteria;p__Firmicutes;c__Clostridia;o__Lachnospirales;f__Lachnospiraceae;g__Lachnospiraceae | 1.771821476 | 0.089668067 | 0.637251673 | IMM-g1->KO |
| d__Bacteria;p__Actinobacteriota;c__Coriobacteriia;o__Coriobacteriales;f__Eggerthellaceae;g__Adlercreutzia | 1.738821657 | 0.100141377 | 0.637251673 | IMM-g1->KO |
| d__Bacteria;p__Firmicutes;c__Clostridia;o__Peptostreptococcales-Tissierellales;f__Peptostreptococcaceae;g__Clostridioides | 1.710674669 | 0.112830371 | 0.637251673 | IMM-g1->KO |
| d__Bacteria;p__Firmicutes;c__Clostridia;o__Peptostreptococcales-Tissierellales;f__Peptostreptococcaceae;g__Romboutsia | 1.565815034 | 0.135815799 | 0.637251673 | IMM-g1->KO |
| d__Bacteria;p__Firmicutes;c__Clostridia;o__Clostridiales;f__Clostridiaceae;g__Clostridium_sensu_stricto_1 | -1.529158148 | 0.14066884 | 0.637251673 | IMM-g2->KO |
| d__Bacteria;p__Firmicutes;c__Clostridia;o__Lachnospirales;f__Lachnospiraceae;g__Anaerostipes | 1.455248308 | 0.159986006 | 0.637251673 | IMM-g1->KO |
| d__Bacteria;p__Firmicutes;c__Clostridia;o__Peptococcales;f__Peptococcaceae;g__uncultured | 1.443422083 | 0.167076745 | 0.637251673 | IMM-g1->KO |
| d__Bacteria;p__Firmicutes;c__Clostridia;o__Lachnospirales;f__Lachnospiraceae;g__Blautia | 1.442714984 | 0.174014077 | 0.637251673 | IMM-g1->KO |
| d__Bacteria;p__Firmicutes;c__Clostridia;o__Lachnospirales;f__Lachnospiraceae;g__Roseburia | -1.47921969 | 0.182617258 | 0.637251673 | IMM-g2->KO |
| d__Bacteria;p__Firmicutes;c__Clostridia;o__Lachnospirales;f__Lachnospiraceae;__ | 1.430061495 | 0.184327749 | 0.637251673 | IMM-g1->KO |
| d__Bacteria;p__Firmicutes;c__Bacilli;o__Erysipelotrichales;f__Erysipelotrichaceae;g__[Clostridium]_innocuum_group | 1.365357588 | 0.197712806 | 0.637251673 | IMM-g1->KO |
| d__Bacteria;p__Firmicutes;c__Clostridia;o__Oscillospirales;f__[Clostridium]_methylpentosum_group;g__[Clostridium]_methylpentosum_group | 1.28716507 | 0.224020387 | 0.637251673 | IMM-g1->KO |
| d__Bacteria;p__Firmicutes;c__Clostridia;o__Oscillospirales;f__Ruminococcaceae;g__uncultured | 1.244477421 | 0.225459763 | 0.637251673 | IMM-g1->KO |
| d__Bacteria;p__Firmicutes;c__Clostridia;o__Monoglobales;f__Monoglobaceae;g__Monoglobus | 1.090234327 | 0.294686066 | 0.637251673 | IMM-g1->KO |
| d__Bacteria;p__Firmicutes;c__Clostridia;o__Oscillospirales;f__Ruminococcaceae;g__UBA1819 | -1.05523528 | 0.3027141 | 0.637251673 | IMM-g2->KO |
| d__Bacteria;p__Firmicutes;c__Clostridia;o__Lachnospirales;f__Lachnospiraceae;g__[Ruminococcus]_torques_group | -1.029167277 | 0.315713803 | 0.637251673 | IMM-g2->KO |
| d__Bacteria;p__Firmicutes;c__Clostridia;o__Clostridiales;f__Clostridiaceae;g__Clostridium_sensu_stricto_13 | 1.024826689 | 0.329858118 | 0.637251673 | IMM-g1->KO |
| d__Bacteria;p__Firmicutes;c__Bacilli;o__Erysipelotrichales;f__Erysipelotrichaceae;g__Turicibacter | 1 | 0.331332762 | 0.637251673 | IMM-g1->KO |
| d__Bacteria;p__Firmicutes;c__Clostridia;o__Oscillospirales;f__Ruminococcaceae;g__Ruminococcus | 1 | 0.331332762 | 0.637251673 | IMM-g1->KO |
| d__Bacteria;p__Firmicutes;c__Bacilli;o__Lactobacillales;f__Streptococcaceae;g__Streptococcus | 1 | 0.331332762 | 0.637251673 | IMM-g1->KO |
| d__Bacteria;p__Bacteroidota;c__Bacteroidia;o__Bacteroidales;f__Tannerellaceae;g__Parabacteroides | 1 | 0.331332762 | 0.637251673 | IMM-g1->KO |
| d__Bacteria;p__Firmicutes;c__Clostridia;o__Oscillospirales;f__Oscillospiraceae;g__UCG-005 | 1 | 0.331332762 | 0.637251673 | IMM-g1->KO |
| d__Bacteria;p__Firmicutes;c__Bacilli;o__Staphylococcales;f__Staphylococcaceae;g__Staphylococcus | 1 | 0.331332762 | 0.637251673 | IMM-g1->KO |
| d__Bacteria;p__Firmicutes;c__Clostridia;o__Lachnospirales;f__Lachnospiraceae;g__[Eubacterium]_fissicatena_group | 1 | 0.331332762 | 0.637251673 | IMM-g1->KO |
| d__Bacteria;p__Firmicutes;c__Clostridia;o__Lachnospirales;f__Lachnospiraceae;g__Lachnospiraceae_UCG-008 | 1 | 0.331332762 | 0.637251673 | IMM-g1->KO |
| d__Bacteria;p__Firmicutes;c__Clostridia;o__Lachnospirales;f__Lachnospiraceae;g__Lachnospiraceae_NK4A136_group | -1.00322518 | 0.332159702 | 0.637251673 | IMM-g2->KO |
| d__Bacteria;p__Firmicutes;c__Clostridia;o__Lachnospirales;f__Lachnospiraceae;g__GCA-900066755 | -0.987297316 | 0.338209233 | 0.637251673 | IMM-g2->KO |
| d__Bacteria;p__Actinobacteriota;c__Coriobacteriia;o__Coriobacteriales;f__Eggerthellaceae;g__Eggerthella | -0.96136368 | 0.347080484 | 0.637251673 | IMM-g2->KO |
| d__Bacteria;p__Bacteroidota;c__Bacteroidia;o__Bacteroidales;f__Tannerellaceae;__ | -1 | 0.350616663 | 0.637251673 | IMM-g2->KO |
| d__Bacteria;p__Firmicutes;c__Clostridia;o__Oscillospirales;f__Oscillospiraceae;g__Flavonifractor | -0.917882723 | 0.372517322 | 0.637251673 | IMM-g2->KO |
| d__Bacteria;p__Firmicutes;c__Clostridia;o__Oscillospirales;f__Ruminococcaceae;g__Incertae_Sedis | 0.910553243 | 0.37454912 | 0.637251673 | IMM-g1->KO |
| d__Bacteria;p__Firmicutes;c__Clostridia;o__Peptostreptococcales-Tissierellales;f__Anaerovoracaceae;g__[Eubacterium]_nodatum_group | 0.921228249 | 0.374853925 | 0.637251673 | IMM-g1->KO |
| d__Bacteria;p__Verrucomicrobiota;c__Verrucomicrobiae;o__Verrucomicrobiales;f__Akkermansiaceae;g__Akkermansia | -0.883961369 | 0.385718221 | 0.639727781 | IMM-g2->KO |
| d__Bacteria;p__Firmicutes;c__Bacilli;o__Lactobacillales;f__Enterococcaceae;g__Enterococcus | -0.851968971 | 0.403008872 | 0.652490555 | IMM-g2->KO |
| d__Bacteria;p__Firmicutes;c__Bacilli;o__Erysipelotrichales;f__Erysipelatoclostridiaceae;g__Erysipelatoclostridium | -0.821580616 | 0.428335615 | 0.677367949 | IMM-g2->KO |
| d__Bacteria;p__Firmicutes;c__Clostridia;o__Lachnospirales;f__Lachnospiraceae;g__Hungatella | -0.785244203 | 0.441923486 | 0.68297266 | IMM-g2->KO |
| d__Bacteria;p__Firmicutes;c__Clostridia;o__Oscillospirales;f__Ruminococcaceae;g__Anaerotruncus | 0.749570506 | 0.460820206 | 0.696350533 | IMM-g1->KO |
| d__Bacteria;p__Proteobacteria;c__Gammaproteobacteria;o__Burkholderiales;f__Sutterellaceae;g__Parasutterella | -0.689642997 | 0.502162076 | 0.741763364 | IMM-g2->KO |
| d__Bacteria;p__Firmicutes;c__Clostridia;o__Lachnospirales;f__Lachnospiraceae;g__Lactonifactor | 0.677643701 | 0.512689384 | 0.741763364 | IMM-g1->KO |
| d__Bacteria;p__Firmicutes;c__Clostridia;o__Oscillospirales;f__Ruminococcaceae;g__Faecalibacterium | -0.598638656 | 0.564330581 | 0.786128896 | IMM-g2->KO |
| d__Bacteria;p__Firmicutes;c__Clostridia;o__Oscillospirales;f__[Eubacterium]_coprostanoligenes_group;g__[Eubacterium]_coprostanoligenes_group | 0.576649923 | 0.573364314 | 0.786128896 | IMM-g1->KO |
| d__Bacteria;p__Firmicutes;c__Clostridia;o__Lachnospirales;f__Lachnospiraceae;g__Eisenbergiella | -0.535838649 | 0.59798976 | 0.786128896 | IMM-g2->KO |
| d__Bacteria;p__Proteobacteria;c__Gammaproteobacteria;o__Enterobacterales;f__Enterobacteriaceae;g__Escherichia-Shigella | -0.530965414 | 0.600325986 | 0.786128896 | IMM-g2->KO |
| d__Bacteria;p__Firmicutes;c__Clostridia;o__Oscillospirales;f__Ruminococcaceae;g__Paludicola | -0.508862295 | 0.619123601 | 0.786128896 | IMM-g2->KO |
| d__Bacteria;p__Firmicutes;c__Clostridia;o__Peptostreptococcales-Tissierellales;f__Anaerovoracaceae;g__Family_XIII_AD3011_group | -0.505118577 | 0.620467035 | 0.786128896 | IMM-g2->KO |
| d__Bacteria;p__Firmicutes;c__Clostridia;o__Lachnospirales;f__Lachnospiraceae;g__Lachnoclostridium | -0.496368254 | 0.624278829 | 0.786128896 | IMM-g2->KO |
| d__Bacteria;__;__;__;__;__ | -0.459210883 | 0.655926913 | 0.810964183 | IMM-g2->KO |
| d__Bacteria;p__Firmicutes;c__Clostridia;o__Lachnospirales;f__Lachnospiraceae;g__Sellimonas | -0.420755898 | 0.678665264 | 0.812239258 | IMM-g2->KO |
| Unassigned;__;__;__;__;__ | -0.422510114 | 0.680847613 | 0.812239258 | IMM-g2->KO |
| d__Bacteria;p__Firmicutes;c__Bacilli;o__Erysipelotrichales;f__Erysipelotrichaceae;g__Dielma | -0.399973451 | 0.693937679 | 0.813582107 | IMM-g2->KO |
| d__Bacteria;p__Actinobacteriota;c__Coriobacteriia;o__Coriobacteriales;f__Eggerthellaceae;g__Gordonibacter | 0.33975278 | 0.737124739 | 0.84245897 | IMM-g1->KO |
| d__Bacteria;p__Bacteroidota;c__Bacteroidia;o__Bacteroidales;f__Rikenellaceae;g__Alistipes | 0.334615787 | 0.74334615 | 0.84245897 | IMM-g1->KO |
| d__Bacteria;p__Actinobacteriota;c__Coriobacteriia;o__Coriobacteriales;f__Coriobacteriaceae;g__Collinsella | 0.30377837 | 0.767013077 | 0.85014437 | IMM-g1->KO |
| d__Bacteria;p__Firmicutes;c__Clostridia;o__Oscillospirales;f__Ruminococcaceae;g__Phocea | -0.292222728 | 0.775131632 | 0.85014437 | IMM-g2->KO |
| d__Bacteria;p__Firmicutes;c__Clostridia;o__Oscillospirales;f__Oscillospiraceae;g__uncultured | -0.246610801 | 0.808589807 | 0.872763601 | IMM-g2->KO |
| d__Bacteria;p__Proteobacteria;c__Gammaproteobacteria;o__Vibrionales;f__Vibrionaceae;g__Vibrio | 0.161833766 | 0.873783511 | 0.906851302 | IMM-g1->KO |
| d__Bacteria;p__Firmicutes;c__Clostridia;o__Lachnospirales;f__Lachnospiraceae;g__GCA-900066575 | -0.159018072 | 0.876375611 | 0.906851302 | IMM-g2->KO |
| d__Bacteria;p__Bacteroidota;c__Bacteroidia;o__Bacteroidales;f__Bacteroidaceae;g__Bacteroides | -0.153958675 | 0.880179204 | 0.906851302 | IMM-g2->KO |
| d__Bacteria;p__Firmicutes;c__Bacilli;o__Erysipelotrichales;f__Erysipelotrichaceae;g__Holdemania | -0.065367679 | 0.948903843 | 0.963066587 | IMM-g2->KO |
| d__Bacteria;p__Firmicutes;c__Clostridia;o__Oscillospirales;f__Ruminococcaceae;g__Candidatus_Soleaferrea | -0.036532421 | 0.971406292 | 0.971406292 | IMM-g2->KO |

| **Operational Taxonomic Unit (Genus) for IMM-g1->KO vs NIMM-g1->KO Comparison** | **T-value** | **P-value** | **FDR** | **Enriched** |
|---|---|---|---|---|
| d__Bacteria;p__Firmicutes;c__Clostridia;o__Lachnospirales;f__Lachnospiraceae;__ | -9.259623812 | 3.64E-09 | 2.51E-07 | NIMM-g1->KO |
| d__Bacteria;p__Firmicutes;c__Clostridia;o__Lachnospirales;f__Lachnospiraceae;g__Lactonifactor | 10.34935696 | 9.34E-09 | 3.22E-07 | IMM-g1->KO |
| d__Bacteria;p__Proteobacteria;c__Gammaproteobacteria;o__Enterobacterales;f__Enterobacteriaceae;g__Escherichia-Shigella | 8.962358466 | 1.09E-06 | 2.52E-05 | IMM-g1->KO |
| d__Bacteria;p__Firmicutes;c__Clostridia;o__Clostridiales;f__Clostridiaceae;g__Clostridium_sensu_stricto_13 | 7.95976078 | 1.50E-06 | 2.59E-05 | IMM-g1->KO |
| d__Bacteria;p__Firmicutes;c__Bacilli;o__Erysipelotrichales;f__Erysipelotrichaceae;g__[Clostridium]_innocuum_group | -5.584566259 | 1.56E-05 | 0.00021484 | NIMM-g1->KO |
| d__Bacteria;p__Firmicutes;c__Clostridia;o__Lachnospirales;f__Lachnospiraceae;g__GCA-900066575 | -5.480629775 | 2.82E-05 | 0.000324568 | NIMM-g1->KO |
| d__Bacteria;p__Actinobacteriota;c__Coriobacteriia;o__Coriobacteriales;f__Eggerthellaceae;g__Gordonibacter | -4.886326205 | 6.97E-05 | 0.000686563 | NIMM-g1->KO |
| d__Bacteria;p__Firmicutes;c__Clostridia;o__Lachnospirales;f__Lachnospiraceae;g__Lachnoclostridium | -4.419148454 | 0.000207034 | 0.001785671 | NIMM-g1->KO |
| d__Bacteria;p__Firmicutes;c__Clostridia;o__Lachnospirales;f__Lachnospiraceae;g__[Ruminococcus]_torques_group | -4.798140769 | 0.000266507 | 0.002043224 | NIMM-g1->KO |
| d__Bacteria;p__Firmicutes;c__Clostridia;o__Oscillospirales;f__[Eubacterium]_coprostanoligenes_group;g__[Eubacterium]_coprostanoligenes_group | 4.436407564 | 0.000361803 | 0.002496442 | IMM-g1->KO |
| d__Bacteria;p__Bacteroidota;c__Bacteroidia;o__Bacteroidales;f__Tannerellaceae;g__Parabacteroides | -5.769982356 | 0.00056108 | 0.003251283 | NIMM-g1->KO |
| d__Bacteria;p__Firmicutes;c__Clostridia;o__Monoglobales;f__Monoglobaceae;g__Monoglobus | -4.105826006 | 0.000588822 | 0.003251283 | NIMM-g1->KO |
| d__Bacteria;p__Firmicutes;c__Clostridia;o__Oscillospirales;f__Ruminococcaceae;g__UBA1819 | -4.007035018 | 0.000612561 | 0.003251283 | NIMM-g1->KO |
| d__Bacteria;p__Firmicutes;c__Clostridia;o__Lachnospirales;f__Lachnospiraceae;g__Anaerostipes | -4.154617217 | 0.000714635 | 0.003522128 | NIMM-g1->KO |
| d__Bacteria;p__Firmicutes;c__Clostridia;o__Oscillospirales;f__Ruminococcaceae;g__Negativibacillus | 3.982815193 | 0.000962385 | 0.004426971 | IMM-g1->KO |
| d__Bacteria;p__Firmicutes;c__Clostridia;o__Lachnospirales;f__Lachnospiraceae;g__[Eubacterium]_fissicatena_group | -4.853582686 | 0.00113223 | 0.004882742 | NIMM-g1->KO |
| d__Bacteria;p__Firmicutes;c__Clostridia;o__Oscillospirales;f__Oscillospiraceae;g__UCG-005 | -4.549822677 | 0.002019342 | 0.008196153 | NIMM-g1->KO |
| d__Bacteria;p__Firmicutes;c__Clostridia;o__Peptostreptococcales-Tissierellales;f__Peptostreptococcaceae;g__Terrisporobacter | 3.583072925 | 0.002291561 | 0.008784315 | IMM-g1->KO |
| d__Bacteria;p__Firmicutes;c__Clostridia;o__Oscillospirales;f__Oscillospiraceae;g__Colidextribacter | -3.302578334 | 0.003286828 | 0.011471775 | NIMM-g1->KO |
| d__Bacteria;p__Firmicutes;c__Clostridia;o__Oscillospirales;f__Ruminococcaceae;g__Paludicola | -3.668368015 | 0.003325152 | 0.011471775 | NIMM-g1->KO |
| d__Bacteria;p__Proteobacteria;c__Gammaproteobacteria;o__Burkholderiales;f__Sutterellaceae;g__Parasutterella | 3.279859474 | 0.004418018 | 0.01413377 | IMM-g1->KO |
| d__Bacteria;p__Firmicutes;c__Clostridia;o__Lachnospirales;f__Lachnospiraceae;g__Blautia | -3.143208659 | 0.004664629 | 0.01413377 | NIMM-g1->KO |
| d__Bacteria;p__Firmicutes;c__Clostridia;o__Oscillospirales;f__[Clostridium]_methylpentosum_group;g__[Clostridium]_methylpentosum_group | 3.340531384 | 0.004711257 | 0.01413377 | IMM-g1->KO |
| d__Bacteria;p__Actinobacteriota;c__Actinobacteria;o__Bifidobacteriales;f__Bifidobacteriaceae;g__Bifidobacterium | 3.009219717 | 0.006295862 | 0.018100604 | IMM-g1->KO |
| d__Bacteria;p__Firmicutes;c__Clostridia;o__Lachnospirales;f__Lachnospiraceae;g__Epulopiscium | 2.973947901 | 0.008514394 | 0.023499727 | IMM-g1->KO |
| d__Bacteria;p__Firmicutes;c__Clostridia;o__Lachnospirales;f__Lachnospiraceae;g__Lachnospiraceae | -2.875306401 | 0.013403759 | 0.03376681 | NIMM-g1->KO |
| d__Bacteria;p__Firmicutes;c__Bacilli;o__Erysipelotrichales;f__Erysipelotrichaceae;g__Dielma | -2.712784197 | 0.013487139 | 0.03376681 | NIMM-g1->KO |
| d__Bacteria;p__Actinobacteriota;c__Coriobacteriia;o__Coriobacteriales;f__Eggerthellaceae;g__Eggerthella | -2.67305957 | 0.013702474 | 0.03376681 | NIMM-g1->KO |
| d__Bacteria;p__Actinobacteriota;c__Coriobacteriia;o__Coriobacteriales;f__Eggerthellaceae;g__Adlercreutzia | -2.678754809 | 0.027129443 | 0.064549365 | NIMM-g1->KO |
| d__Bacteria;p__Firmicutes;c__Clostridia;o__Oscillospirales;f__Oscillospiraceae;g__uncultured | -2.279698609 | 0.033237763 | 0.075452355 | NIMM-g1->KO |
| d__Bacteria;p__Firmicutes;c__Clostridia;o__Oscillospirales;f__Oscillospiraceae;g__Flavonifractor | -2.256124314 | 0.033898884 | 0.075452355 | NIMM-g1->KO |
| d__Bacteria;p__Firmicutes;c__Clostridia;o__Oscillospirales;f__Ruminococcaceae;g__Anaerotruncus | -2.28866896 | 0.035517805 | 0.076444737 | NIMM-g1->KO |
| d__Bacteria;p__Verrucomicrobiota;c__Verrucomicrobiae;o__Verrucomicrobiales;f__Akkermansiaceae;g__Akkermansia | -2.23860747 | 0.036560526 | 0.076444737 | NIMM-g1->KO |
| d__Bacteria;p__Firmicutes;c__Bacilli;o__Lactobacillales;f__Enterococcaceae;g__Enterococcus | 2.405076659 | 0.03768028 | 0.076468803 | IMM-g1->KO |
| d__Bacteria;p__Firmicutes;c__Clostridia;o__Oscillospirales;f__Ruminococcaceae;g__uncultured | -2.435762134 | 0.039254703 | 0.077387844 | NIMM-g1->KO |
| d__Bacteria;__;__;__;__;__ | 2.157570254 | 0.045561675 | 0.087326544 | IMM-g1->KO |
| d__Bacteria;p__Firmicutes;c__Clostridia;o__Clostridiales;f__Clostridiaceae;g__Clostridium_sensu_stricto_1 | 2.258441239 | 0.059863022 | 0.111636446 | IMM-g1->KO |
| d__Bacteria;p__Firmicutes;c__Clostridia;o__Lachnospirales;f__Lachnospiraceae;g__Eisenbergiella | -1.924129325 | 0.067126149 | 0.121886955 | NIMM-g1->KO |
| d__Bacteria;p__Firmicutes;c__Clostridia;o__Lachnospirales;f__Lachnospiraceae;g__[Ruminococcus]_gnavus_group | -1.882062701 | 0.075443177 | 0.13347639 | NIMM-g1->KO |
| d__Bacteria;p__Actinobacteriota;c__Coriobacteriia;o__Coriobacteriales;f__Coriobacteriaceae;g__Collinsella | -1.738527978 | 0.095835182 | 0.165315689 | NIMM-g1->KO |
| d__Bacteria;p__Firmicutes;c__Clostridia;o__Peptostreptococcales-Tissierellales;f__Peptostreptococcaceae;g__Romboutsia | 1.565815034 | 0.135815799 | 0.221848209 | IMM-g1->KO |
| d__Bacteria;p__Firmicutes;c__Clostridia;o__Lachnospirales;f__Lachnospiraceae;g__Lachnospiraceae_NK4A136_group | -1.625334703 | 0.137285691 | 0.221848209 | NIMM-g1->KO |
| d__Bacteria;p__Proteobacteria;c__Gammaproteobacteria;o__Vibrionales;f__Vibrionaceae;g__Vibrio | 1.552994099 | 0.138253232 | 0.221848209 | IMM-g1->KO |
| d__Bacteria;p__Firmicutes;c__Bacilli;o__Erysipelotrichales;f__Erysipelotrichaceae;g__Holdemania | -1.561865468 | 0.148269884 | 0.232514136 | NIMM-g1->KO |
| d__Bacteria;p__Firmicutes;c__Clostridia;o__Lachnospirales;f__Lachnospiraceae;g__Lachnospiraceae_UCG-008 | -1.526158281 | 0.163969609 | 0.24929489 | NIMM-g1->KO |
| d__Bacteria;p__Firmicutes;c__Clostridia;o__Peptococcales;f__Peptococcaceae;g__uncultured | 1.443422083 | 0.167076745 | 0.24929489 | IMM-g1->KO |
| Unassigned;__;__;__;__;__ | -1.415163697 | 0.170418489 | 0.24929489 | NIMM-g1->KO |
| d__Bacteria;p__Firmicutes;c__Bacilli;o__Erysipelotrichales;f__Erysipelatoclostridiaceae;g__Erysipelatoclostridium | -1.474632177 | 0.174640448 | 0.24929489 | NIMM-g1->KO |
| d__Bacteria;p__Bacteroidota;c__Bacteroidia;o__Bacteroidales;f__Bacteroidaceae;g__Bacteroides | 1.418952189 | 0.177035502 | 0.24929489 | IMM-g1->KO |
| d__Bacteria;p__Firmicutes;c__Clostridia;o__Peptostreptococcales-Tissierellales;f__Peptostreptococcaceae;g__Clostridioides | 1.277417102 | 0.243537011 | 0.336081075 | IMM-g1->KO |
| d__Bacteria;p__Firmicutes;c__Clostridia;o__Oscillospirales;f__Ruminococcaceae;g__Faecalibacterium | 1 | 0.331332762 | 0.416242715 | IMM-g1->KO |
| d__Bacteria;p__Firmicutes;c__Bacilli;o__Erysipelotrichales;f__Erysipelotrichaceae;g__Turicibacter | 1 | 0.331332762 | 0.416242715 | IMM-g1->KO |
| d__Bacteria;p__Firmicutes;c__Clostridia;o__Oscillospirales;f__Ruminococcaceae;g__Ruminococcus | 1 | 0.331332762 | 0.416242715 | IMM-g1->KO |
| d__Bacteria;p__Firmicutes;c__Bacilli;o__Lactobacillales;f__Streptococcaceae;g__Streptococcus | 1 | 0.331332762 | 0.416242715 | IMM-g1->KO |
| d__Bacteria;p__Firmicutes;c__Clostridia;o__Lachnospirales;f__Lachnospiraceae;g__Hungatella | -1.010987234 | 0.336855455 | 0.416242715 | NIMM-g1->KO |
| d__Bacteria;p__Firmicutes;c__Clostridia;o__Lachnospirales;f__Lachnospiraceae;g__GCA-900066755 | -0.962798357 | 0.35385772 | 0.416242715 | NIMM-g1->KO |
| d__Bacteria;p__Firmicutes;c__Clostridia;o__Lachnospirales;f__Lachnospiraceae;g__Agathobacter | -1 | 0.355917684 | 0.416242715 | NIMM-g1->KO |
| d__Bacteria;p__Firmicutes;c__Clostridia;o__Oscillospirales;f__Ruminococcaceae;g__DTU089 | -1 | 0.355917684 | 0.416242715 | NIMM-g1->KO |
| d__Bacteria;p__Actinobacteriota;c__Coriobacteriia;o__Coriobacteriales;f__Eggerthellaceae;__ | -1 | 0.355917684 | 0.416242715 | NIMM-g1->KO |
| d__Bacteria;p__Firmicutes;c__Clostridia;o__Oscillospirales;f__Oscillospiraceae;g__Oscillibacter | -0.916456157 | 0.377129376 | 0.433698782 | NIMM-g1->KO |
| d__Bacteria;p__Firmicutes;c__Clostridia;o__Oscillospirales;f__Butyricicoccaceae;g__Butyricicoccus | -0.773590318 | 0.448291594 | 0.507083934 | NIMM-g1->KO |
| d__Bacteria;p__Firmicutes;c__Clostridia;o__Peptostreptococcales-Tissierellales;f__Anaerovoracaceae;g__[Eubacterium]_nodatum_group | -0.776837693 | 0.456165858 | 0.507668455 | NIMM-g1->KO |
| d__Bacteria;p__Bacteroidota;c__Bacteroidia;o__Bacteroidales;f__Rikenellaceae;g__Alistipes | 0.566996645 | 0.577298706 | 0.632279535 | IMM-g1->KO |
| d__Bacteria;p__Firmicutes;c__Clostridia;o__Oscillospirales;f__Ruminococcaceae;g__Incertae_Sedis | 0.522997364 | 0.6178284 | 0.666096244 | IMM-g1->KO |
| d__Bacteria;p__Firmicutes;c__Clostridia;o__Peptostreptococcales-Tissierellales;f__Anaerovoracaceae;g__Family_XIII_AD3011_group | -0.489729471 | 0.631971287 | 0.670861828 | NIMM-g1->KO |
| d__Bacteria;p__Firmicutes;c__Bacilli;o__Staphylococcales;f__Staphylococcaceae;g__Staphylococcus | -0.415177608 | 0.687157852 | 0.7183923 | NIMM-g1->KO |
| d__Bacteria;p__Firmicutes;c__Clostridia;o__Oscillospirales;f__Ruminococcaceae;g__Phocea | 0.346109708 | 0.736972828 | 0.758972017 | IMM-g1->KO |
| d__Bacteria;p__Firmicutes;c__Clostridia;o__Lachnospirales;f__Lachnospiraceae;g__Sellimonas | -0.331236364 | 0.748401633 | 0.759407539 | NIMM-g1->KO |
| d__Bacteria;p__Firmicutes;c__Clostridia;o__Oscillospirales;f__Ruminococcaceae;g__Candidatus_Soleaferrea | 0.019708413 | 0.984649104 | 0.984649104 | IMM-g1->KO |

| **Operational Taxonomic Unit (Genus) for IMM-g1->KO vs NIMM-g1->WT Comparison** | **T-value** | **P-value** | **FDR** | **Enriched** |
|---|---|---|---|---|
| d__Bacteria;p__Firmicutes;c__Clostridia;o__Lachnospirales;f__Lachnospiraceae;g__Lachnospiraceae_UCG-008 | -10.622979 | 3.30E-12 | 1.20E-10 | NIMM-g1->WT |
| d__Bacteria;p__Firmicutes;c__Clostridia;o__Lachnospirales;f__Lachnospiraceae;g__[Eubacterium]_fissicatena_group | -10.496807 | 3.43E-12 | 1.20E-10 | NIMM-g1->WT |
| d__Bacteria;p__Firmicutes;c__Clostridia;o__Clostridiales;f__Clostridiaceae;g__Clostridium_sensu_stricto_13 | 9.80382396 | 3.05E-11 | 7.11E-10 | IMM-g1->KO |
| d__Bacteria;p__Proteobacteria;c__Gammaproteobacteria;o__Enterobacterales;f__Enterobacteriaceae;g__Escherichia-Shigella | 8.84514842 | 3.52E-10 | 6.16E-09 | IMM-g1->KO |
| d__Bacteria;p__Firmicutes;c__Clostridia;o__Clostridiales;f__Clostridiaceae;g__Clostridium_sensu_stricto_1 | 9.16653359 | 1.39E-09 | 1.95E-08 | IMM-g1->KO |
| d__Bacteria;p__Bacteroidota;c__Bacteroidia;o__Bacteroidales;f__Tannerellaceae;g__Parabacteroides | -8.8426351 | 3.28E-09 | 3.82E-08 | NIMM-g1->WT |
| d__Bacteria;p__Firmicutes;c__Clostridia;o__Lachnospirales;f__Lachnospiraceae;g__[Ruminococcus]_gnavus_group | 7.68212631 | 6.29E-09 | 6.29E-08 | IMM-g1->KO |
| d__Bacteria;p__Firmicutes;c__Clostridia;o__Lachnospirales;f__Lachnospiraceae;g__Lactonifactor | 10.349357 | 9.34E-09 | 8.17E-08 | IMM-g1->KO |
| d__Bacteria;p__Firmicutes;c__Clostridia;o__Lachnospirales;f__Lachnospiraceae;__ | -7.2656175 | 2.74E-08 | 2.13E-07 | NIMM-g1->WT |
| d__Bacteria;p__Firmicutes;c__Clostridia;o__Lachnospirales;f__Lachnospiraceae;g__[Ruminococcus]_torques_group | -7.1152672 | 3.21E-08 | 2.24E-07 | NIMM-g1->WT |
| d__Bacteria;p__Firmicutes;c__Clostridia;o__Lachnospirales;f__Lachnospiraceae;g__Lachnospiraceae_NK4A136_group | -7.0472329 | 3.91E-08 | 2.49E-07 | NIMM-g1->WT |
| d__Bacteria;p__Firmicutes;c__Clostridia;o__Oscillospirales;f__Oscillospiraceae;g__UCG-005 | -7.4736582 | 6.33E-08 | 3.69E-07 | NIMM-g1->WT |
| d__Bacteria;p__Firmicutes;c__Clostridia;o__Lachnospirales;f__Lachnospiraceae;g__Lachnospiraceae | -5.7018746 | 2.11E-06 | 1.14E-05 | NIMM-g1->WT |
| d__Bacteria;p__Firmicutes;c__Clostridia;o__Oscillospirales;f__[Clostridium]_methylpentosum_group;g__[Clostridium]_methylpentosum_group | 6.52211668 | 5.22E-06 | 2.61E-05 | IMM-g1->KO |
| d__Bacteria;p__Firmicutes;c__Clostridia;o__Monoglobales;f__Monoglobaceae;g__Monoglobus | -5.6144412 | 7.09E-06 | 3.31E-05 | NIMM-g1->WT |
| d__Bacteria;p__Actinobacteriota;c__Coriobacteriia;o__Coriobacteriales;f__Eggerthellaceae;g__Gordonibacter | -5.2161289 | 1.19E-05 | 5.19E-05 | NIMM-g1->WT |
| d__Bacteria;p__Firmicutes;c__Clostridia;o__Lachnospirales;f__Lachnospiraceae;g__Blautia | -4.970322 | 2.61E-05 | 0.0001076 | NIMM-g1->WT |
| d__Bacteria;p__Firmicutes;c__Clostridia;o__Lachnospirales;f__Lachnospiraceae;g__Anaerostipes | -5.2687516 | 3.20E-05 | 0.00012436 | NIMM-g1->WT |
| d__Bacteria;p__Firmicutes;c__Clostridia;o__Lachnospirales;f__Lachnospiraceae;g__Lachnoclostridium | -4.6894869 | 4.49E-05 | 0.00016532 | NIMM-g1->WT |
| d__Bacteria;p__Firmicutes;c__Clostridia;o__Oscillospirales;f__Ruminococcaceae;g__Incertae_Sedis | -4.4849006 | 8.34E-05 | 0.00029203 | NIMM-g1->WT |
| d__Bacteria;p__Firmicutes;c__Clostridia;o__Oscillospirales;f__Ruminococcaceae;g__UBA1819 | -4.3767712 | 0.00011279 | 0.00037065 | NIMM-g1->WT |
| d__Bacteria;p__Firmicutes;c__Bacilli;o__Erysipelotrichales;f__Erysipelotrichaceae;g__Dielma | -4.3782033 | 0.00011649 | 0.00037065 | NIMM-g1->WT |
| d__Bacteria;p__Firmicutes;c__Clostridia;o__Lachnospirales;f__Lachnospiraceae;g__GCA-900066575 | -4.2099113 | 0.00017773 | 0.00054093 | NIMM-g1->WT |
| d__Bacteria;p__Firmicutes;c__Clostridia;o__Oscillospirales;f__Ruminococcaceae;g__uncultured | -4.1846877 | 0.0002568 | 0.00074901 | NIMM-g1->WT |
| d__Bacteria;p__Firmicutes;c__Clostridia;o__Oscillospirales;f__Ruminococcaceae;g__Paludicola | -3.7970765 | 0.00057855 | 0.00161993 | NIMM-g1->WT |
| d__Bacteria;p__Firmicutes;c__Clostridia;o__Oscillospirales;f__Oscillospiraceae;g__Colidextribacter | -3.7150317 | 0.00076959 | 0.00207197 | NIMM-g1->WT |
| d__Bacteria;p__Firmicutes;c__Clostridia;o__Oscillospirales;f__Ruminococcaceae;g__Negativibacillus | 3.98281519 | 0.00096239 | 0.00249507 | IMM-g1->KO |
| d__Bacteria;p__Firmicutes;c__Clostridia;o__Oscillospirales;f__Oscillospiraceae;g__uncultured | -3.6131424 | 0.00134621 | 0.00336553 | NIMM-g1->WT |
| d__Bacteria;p__Actinobacteriota;c__Coriobacteriia;o__Coriobacteriales;f__Eggerthellaceae;g__Adlercreutzia | -3.5159124 | 0.00154775 | 0.00373596 | NIMM-g1->WT |
| d__Bacteria;p__Firmicutes;c__Bacilli;o__Lactobacillales;f__Enterococcaceae;g__Enterococcus | 3.36074582 | 0.00200746 | 0.00468408 | IMM-g1->KO |
| d__Bacteria;p__Firmicutes;c__Clostridia;o__Peptostreptococcales-Tissierellales;f__Peptostreptococcaceae;g__Terrisporobacter | 3.58307293 | 0.00229156 | 0.00506356 | IMM-g1->KO |
| d__Bacteria;p__Firmicutes;c__Clostridia;o__Oscillospirales;f__Ruminococcaceae;__ | -3.5784293 | 0.00231477 | 0.00506356 | NIMM-g1->WT |
| d__Bacteria;p__Firmicutes;c__Clostridia;o__Lachnospirales;f__Lachnospiraceae;g__GCA-900066755 | -2.9600009 | 0.00563095 | 0.01194445 | NIMM-g1->WT |
| d__Bacteria;p__Firmicutes;c__Bacilli;o__Erysipelotrichales;f__Erysipelotrichaceae;g__[Clostridium]_innocuum_group | -2.9372395 | 0.00720693 | 0.01443456 | NIMM-g1->WT |
| Unassigned;__;__;__;__;__ | -2.8984143 | 0.00721728 | 0.01443456 | NIMM-g1->WT |
| d__Bacteria;p__Firmicutes;c__Bacilli;o__Staphylococcales;f__Staphylococcaceae;g__Staphylococcus | -2.8349148 | 0.00972336 | 0.01890654 | NIMM-g1->WT |
| d__Bacteria;p__Firmicutes;c__Clostridia;o__Oscillospirales;f__Oscillospiraceae;g__Flavonifractor | -2.7233817 | 0.01104222 | 0.02089068 | NIMM-g1->WT |
| d__Bacteria;p__Firmicutes;c__Clostridia;o__Lachnospirales;f__Lachnospiraceae;g__Sellimonas | -2.6506861 | 0.01240324 | 0.02284807 | NIMM-g1->WT |
| d__Bacteria;p__Actinobacteriota;c__Actinobacteria;o__Bifidobacteriales;f__Bifidobacteriaceae;g__Bifidobacterium | 2.560816 | 0.01515593 | 0.02720296 | IMM-g1->KO |
| d__Bacteria;p__Firmicutes;c__Clostridia;o__Clostridia_vadinBB60_group;f__Clostridia_vadinBB60_group;g__Clostridia_vadinBB60_group | -2.5134119 | 0.02232601 | 0.03907051 | NIMM-g1->WT |
| d__Bacteria;p__Proteobacteria;c__Gammaproteobacteria;o__Burkholderiales;f__Sutterellaceae;g__Parasutterella | 2.33179872 | 0.0291878 | 0.04983282 | IMM-g1->KO |
| d__Bacteria;p__Firmicutes;c__Clostridia;o__Lachnospirales;f__Lachnospiraceae;g__Epulopiscium | 2.16163136 | 0.04184777 | 0.06974629 | IMM-g1->KO |
| d__Bacteria;p__Verrucomicrobiota;c__Verrucomicrobiae;o__Verrucomicrobiales;f__Akkermansiaceae;g__Akkermansia | -2.0211687 | 0.05276367 | 0.08589434 | NIMM-g1->WT |
| d__Bacteria;p__Firmicutes;c__Clostridia;o__Peptostreptococcales-Tissierellales;f__Anaerovoracaceae;g__[Eubacterium]_nodatum_group | -1.9807615 | 0.05931652 | 0.09325662 | NIMM-g1->WT |
| d__Bacteria;p__Firmicutes;c__Clostridia;o__Peptostreptococcales-Tissierellales;f__Peptostreptococcaceae;g__Clostridioides | 1.94641402 | 0.05995068 | 0.09325662 | IMM-g1->KO |
| d__Bacteria;p__Firmicutes;c__Clostridia;o__Lachnospirales;f__Lachnospiraceae;g__Hungatella | 1.93168726 | 0.06177156 | 0.0940002 | IMM-g1->KO |
| d__Bacteria;p__Actinobacteriota;c__Coriobacteriia;o__Coriobacteriales;f__Eggerthellaceae;g__Eggerthella | -1.8933582 | 0.06687929 | 0.09960745 | NIMM-g1->WT |
| d__Bacteria;p__Firmicutes;c__Clostridia;o__Oscillospirales;f__Ruminococcaceae;g__Phocea | 1.76456316 | 0.08663238 | 0.12633889 | IMM-g1->KO |
| d__Bacteria;p__Bacteroidota;c__Bacteroidia;o__Bacteroidales;f__Muribaculaceae;g__Muribaculaceae | -1.7949105 | 0.09046804 | 0.12924006 | NIMM-g1->WT |
| d__Bacteria;p__Firmicutes;c__Clostridia;o__Oscillospirales;f__Ruminococcaceae;g__Candidatus_Soleaferrea | -1.6888999 | 0.10091471 | 0.14128059 | NIMM-g1->WT |
| d__Bacteria;p__Firmicutes;c__Clostridia;o__Oscillospirales;f__Ruminococcaceae;g__Faecalibacterium | -1.6356599 | 0.1155753 | 0.15863276 | NIMM-g1->WT |
| d__Bacteria;p__Firmicutes;c__Clostridia;o__Peptostreptococcales-Tissierellales;f__Peptostreptococcaceae;g__Romboutsia | 1.56581503 | 0.1358158 | 0.18282896 | IMM-g1->KO |
| d__Bacteria;p__Firmicutes;c__Clostridia;o__Peptococcales;f__Peptococcaceae;g__uncultured | 1.44342208 | 0.16707675 | 0.2206674 | IMM-g1->KO |
| d__Bacteria;p__Firmicutes;c__Clostridia;o__Peptostreptococcales-Tissierellales;f__Anaerovoracaceae;g__Family_XIII_AD3011_group | -1.2835328 | 0.20881018 | 0.27067987 | NIMM-g1->WT |
| d__Bacteria;p__Firmicutes;c__Clostridia;o__Lachnospirales;f__Lachnospiraceae;g__Eisenbergiella | -1.0839114 | 0.28662371 | 0.36479382 | NIMM-g1->WT |
| d__Bacteria;p__Firmicutes;c__Bacilli;o__Erysipelotrichales;f__Erysipelotrichaceae;g__Holdemania | -1.042551 | 0.30451493 | 0.37592945 | NIMM-g1->WT |
| d__Bacteria;p__Firmicutes;c__Clostridia;o__Oscillospirales;f__Ruminococcaceae;g__Anaerotruncus | -1.0421147 | 0.30611398 | 0.37592945 | NIMM-g1->WT |
| d__Bacteria;p__Firmicutes;c__Bacilli;o__Lactobacillales;f__Streptococcaceae;g__Streptococcus | 1 | 0.33133276 | 0.39310667 | IMM-g1->KO |
| d__Bacteria;p__Firmicutes;c__Clostridia;o__Oscillospirales;f__Ruminococcaceae;g__DTU089 | -1 | 0.33133276 | 0.39310667 | NIMM-g1->WT |
| d__Bacteria;p__Proteobacteria;c__Gammaproteobacteria;o__Vibrionales;f__Vibrionaceae;g__Vibrio | -0.968239 | 0.33995625 | 0.39661562 | NIMM-g1->WT |
| d__Bacteria;p__Actinobacteriota;c__Coriobacteriia;o__Coriobacteriales;f__Coriobacteriaceae;g__Collinsella | -0.750356 | 0.46168351 | 0.52980075 | NIMM-g1->WT |
| d__Bacteria;p__Firmicutes;c__Clostridia;o__Oscillospirales;f__Butyricicoccaceae;g__Butyricicoccus | -0.6452283 | 0.52478864 | 0.59250331 | NIMM-g1->WT |
| d__Bacteria;p__Firmicutes;c__Clostridia;o__Oscillospirales;f__Oscillospiraceae;g__Oscillibacter | -0.4558583 | 0.65171703 | 0.72413003 | NIMM-g1->WT |
| d__Bacteria;p__Bacteroidota;c__Bacteroidia;o__Bacteroidales;f__Rikenellaceae;g__Alistipes | 0.43916399 | 0.66359204 | 0.72580379 | IMM-g1->KO |
| d__Bacteria;p__Firmicutes;c__Clostridia;o__Oscillospirales;f__[Eubacterium]_coprostanoligenes_group;g__[Eubacterium]_coprostanoligenes_group | 0.37428924 | 0.71052413 | 0.76517983 | IMM-g1->KO |
| d__Bacteria;__;__;__;__;__ | -0.3094416 | 0.75894443 | 0.80494106 | NIMM-g1->WT |
| d__Bacteria;p__Firmicutes;c__Clostridia;o__Oscillospirales;f__Ruminococcaceae;g__Ruminococcus | 0.2628287 | 0.79447934 | 0.83005304 | IMM-g1->KO |
| d__Bacteria;p__Firmicutes;c__Bacilli;o__Erysipelotrichales;f__Erysipelotrichaceae;g__Turicibacter | -0.2430675 | 0.80942435 | 0.83323095 | NIMM-g1->WT |
| d__Bacteria;p__Bacteroidota;c__Bacteroidia;o__Bacteroidales;f__Bacteroidaceae;g__Bacteroides | -0.2064573 | 0.83767204 | 0.84981221 | NIMM-g1->WT |
| d__Bacteria;p__Firmicutes;c__Bacilli;o__Erysipelotrichales;f__Erysipelatoclostridiaceae;g__Erysipelatoclostridium | 0.03618591 | 0.9713523 | 0.9713523 | IMM-g1->KO |

| Operational Taxonomic Unit (Genus) for HM1->WT vs NIMM-g1->WT Comparison | T-value | P-value | FDR | Enriched |
|---|---|---|---|---|
| d__Bacteria;p__Proteobacteria;c__Gammaproteobacteria;o__Vibrionales;f__Vibrionaceae;g__Vibrio | 5.39001366 | 4.40E-05 | 0.002903623 | HM1->WT |
| d__Bacteria;p__Actinobacteriota;c__Coriobacteriia;o__Coriobacteriales;f__Coriobacteriaceae;g__Collinsella | 4.61690007 | 0.000128257 | 0.004232496 | HM1->WT |
| d__Bacteria;p__Firmicutes;c__Clostridia;o__Lachnospirales;f__Lachnospiraceae;g__Blautia | 4.286841959 | 0.00029081 | 0.005215912 | HM1->WT |
| d__Bacteria;p__Firmicutes;c__Clostridia;o__Lachnospirales;f__Lachnospiraceae;g__[Ruminococcus]_gnavus_group | 4.32005584 | 0.000316116 | 0.005215912 | HM1->WT |
| d__Bacteria;p__Actinobacteriota;c__Coriobacteriia;o__Coriobacteriales;f__Eggerthellaceae;g__Adlercreutzia | 3.262255261 | 0.003907314 | 0.051576548 | HM1->WT |
| d__Bacteria;p__Firmicutes;c__Clostridia;o__Oscillospirales;f__Oscillospiraceae;g__Flavonifractor | -3.08615967 | 0.013720324 | 0.126423555 | NIMM-g1->WT |
| d__Bacteria;p__Firmicutes;c__Clostridia;o__Lachnospirales;f__Lachnospiraceae;g__Lachnospiraceae_UCG-008 | -3.089147584 | 0.014565678 | 0.126423555 | NIMM-g1->WT |
| d__Bacteria;p__Firmicutes;c__Clostridia;o__Peptostreptococcales-Tissierellales;f__Peptostreptococcaceae;g__Clostridioides | -3.19237101 | 0.015324067 | 0.126423555 | NIMM-g1->WT |
| d__Bacteria;p__Firmicutes;c__Clostridia;o__Oscillospirales;f__Ruminococcaceae;g__Phocea | -2.540800575 | 0.021105423 | 0.14119397 | NIMM-g1->WT |
| d__Bacteria;p__Firmicutes;c__Bacilli;o__Lactobacillales;f__Enterococcaceae;g__Enterococcus | 2.472945475 | 0.022995662 | 0.14119397 | HM1->WT |
| d__Bacteria;p__Actinobacteriota;c__Coriobacteriia;o__Coriobacteriales;f__Eggerthellaceae;g__Eggerthella | 2.427214297 | 0.023532328 | 0.14119397 | HM1->WT |
| d__Bacteria;p__Firmicutes;c__Bacilli;o__Staphylococcales;f__Staphylococcaceae;g__Staphylococcus | -2.355208076 | 0.02743539 | 0.150894646 | NIMM-g1->WT |
| d__Bacteria;p__Bacteroidota;c__Bacteroidia;o__Bacteroidales;f__Tannerellaceae;g__Parabacteroides | -2.55581732 | 0.031892879 | 0.158836814 | NIMM-g1->WT |
| d__Bacteria;p__Firmicutes;c__Clostridia;o__Oscillospirales;f__[Eubacterium]_coprostanoligenes_group;g__[Eubacterium]_coprostanoligenes_group | -2.267871021 | 0.033692658 | 0.158836814 | NIMM-g1->WT |
| d__Bacteria;p__Firmicutes;c__Clostridia;o__Oscillospirales;f__Ruminococcaceae;g__Faecalibacterium | -2.200106398 | 0.041915882 | 0.178200937 | NIMM-g1->WT |
| d__Bacteria;p__Firmicutes;c__Clostridia;o__Peptostreptococcales-Tissierellales;f__Anaerovoracaceae;g__[Eubacterium]_nodatum_group | -2.459525947 | 0.043200227 | 0.178200937 | NIMM-g1->WT |
| d__Bacteria;p__Firmicutes;c__Clostridia;o__Lachnospirales;f__Lachnospiraceae;g__Lachnospiraceae_NK4A136_group | 2.071102057 | 0.051716146 | 0.190008287 | HM1->WT |
| d__Bacteria;p__Firmicutes;c__Clostridia;o__Oscillospirales;f__Ruminococcaceae;g__Paludicola | 2.043850934 | 0.053198161 | 0.190008287 | HM1->WT |
| d__Bacteria;p__Firmicutes;c__Clostridia;o__Lachnospirales;f__Lachnospiraceae;g__Eisenbergiella | -2.114279629 | 0.058545294 | 0.190008287 | NIMM-g1->WT |
| d__Bacteria;p__Firmicutes;c__Clostridia;o__Oscillospirales;f__Butyricicoccaceae;g__Butyricicoccus | -2.071718584 | 0.059472499 | 0.190008287 | NIMM-g1->WT |
| d__Bacteria;p__Firmicutes;c__Bacilli;o__Erysipelotrichales;f__Erysipelatoclostridiaceae;g__Erysipelatoclostridium | 2.089330198 | 0.060457182 | 0.190008287 | HM1->WT |
| d__Bacteria;p__Firmicutes;c__Clostridia;o__Lachnospirales;f__Lachnospiraceae;g__Anaerostipes | 1.950706497 | 0.063380005 | 0.190140016 | HM1->WT |
| d__Bacteria;p__Actinobacteriota;c__Actinobacteria;o__Bifidobacteriales;f__Bifidobacteriaceae;g__Bifidobacterium | 2.053364079 | 0.073394543 | 0.207469585 | HM1->WT |
| d__Bacteria;p__Firmicutes;c__Clostridia;o__Lachnospirales;f__Lachnospiraceae;g__[Ruminococcus]_torques_group | -1.957613775 | 0.075443485 | 0.207469585 | NIMM-g1->WT |
| d__Bacteria;p__Proteobacteria;c__Gammaproteobacteria;o__Enterobacterales;f__Enterobacteriaceae;g__Escherichia-Shigella | 1.921094562 | 0.086524281 | 0.222844518 | HM1->WT |
| d__Bacteria;p__Bacteroidota;c__Bacteroidia;o__Bacteroidales;f__Muribaculaceae;g__Muribaculaceae | -1.794910476 | 0.090468043 | 0.222844518 | NIMM-g1->WT |
| d__Bacteria;p__Firmicutes;c__Clostridia;o__Lachnospirales;f__Lachnospiraceae;g__GCA-900066755 | -1.773878376 | 0.093269946 | 0.222844518 | NIMM-g1->WT |
| d__Bacteria;p__Firmicutes;c__Clostridia;o__Oscillospirales;f__Oscillospiraceae;g__Oscillibacter | -1.857945506 | 0.094540099 | 0.222844518 | NIMM-g1->WT |
| d__Bacteria;p__Firmicutes;c__Bacilli;o__Erysipelotrichales;f__Erysipelotrichaceae;g__Holdemania | -1.728775775 | 0.1162651 | 0.26460333 | NIMM-g1->WT |
| d__Bacteria;p__Firmicutes;c__Bacilli;o__Erysipelotrichales;f__Erysipelotrichaceae;g__[Clostridium]_innocuum_group | 1.677405449 | 0.120867192 | 0.265907822 | HM1->WT |
| d__Bacteria;p__Firmicutes;c__Clostridia;o__Oscillospirales;f__Ruminococcaceae;g__Candidatus_Soleaferrea | -1.697506755 | 0.127904314 | 0.27231241 | NIMM-g1->WT |
| d__Bacteria;p__Firmicutes;c__Clostridia;o__Clostridiales;f__Clostridiaceae;g__Clostridium_sensu_stricto_1 | 1.589127531 | 0.145287224 | 0.2996549 | HM1->WT |
| d__Bacteria;p__Firmicutes;c__Bacilli;o__Erysipelotrichales;f__Erysipelotrichaceae;g__Turicibacter | -1.449402349 | 0.165422371 | 0.330844743 | NIMM-g1->WT |
| d__Bacteria;p__Firmicutes;c__Clostridia;o__Oscillospirales;f__[Clostridium]_methylpentosum_group;g__[Clostridium]_methylpentosum_group | 1.530185112 | 0.176845092 | 0.343287532 | HM1->WT |
| d__Bacteria;p__Firmicutes;c__Clostridia;o__Lachnospirales;f__Lachnospiraceae;g__Agathobacter | 1.508617827 | 0.182128537 | 0.343442385 | HM1->WT |
| d__Bacteria;p__Firmicutes;c__Bacilli;o__Erysipelotrichales;f__Erysipelotrichaceae;g__Dielma | -1.383357557 | 0.201061003 | 0.368611839 | NIMM-g1->WT |
| d__Bacteria;p__Firmicutes;c__Clostridia;o__Oscillospirales;f__Ruminococcaceae;__ | -1.323175747 | 0.208608909 | 0.372113189 | NIMM-g1->WT |
| d__Bacteria;p__Firmicutes;c__Clostridia;o__Lachnospirales;f__Lachnospiraceae;g__GCA-900066575 | -1.264310475 | 0.2312595 | 0.394940103 | NIMM-g1->WT |
| d__Bacteria;p__Bacteroidota;c__Bacteroidia;o__Bacteroidales;f__Bacteroidaceae;g__Bacteroides | -1.249381891 | 0.233373697 | 0.394940103 | NIMM-g1->WT |
| d__Bacteria;p__Firmicutes;c__Clostridia;o__Oscillospirales;f__Ruminococcaceae;g__UBA1819 | -1.196364252 | 0.252690272 | 0.413570479 | NIMM-g1->WT |
| d__Bacteria;p__Firmicutes;c__Clostridia;o__Oscillospirales;f__Ruminococcaceae;g__uncultured | -1.16963908 | 0.256914994 | 0.413570479 | NIMM-g1->WT |
| d__Bacteria;p__Firmicutes;c__Clostridia;o__Oscillospirales;f__Oscillospiraceae;g__uncultured | -1.102286837 | 0.300198735 | 0.471740869 | NIMM-g1->WT |
| d__Bacteria;p__Firmicutes;c__Clostridia;o__Clostridia_vadinBB60_group;f__Clostridia_vadinBB60_group;g__Clostridia_vadinBB60_group | -0.96328753 | 0.349482906 | 0.533876526 | NIMM-g1->WT |
| d__Bacteria;p__Firmicutes;c__Clostridia;o__Lachnospirales;f__Lachnospiraceae;g__uncultured | 1 | 0.355917684 | 0.533876526 | HM1->WT |
| d__Bacteria;p__Firmicutes;c__Clostridia;o__Lachnospirales;f__Lachnospiraceae;__ | -0.812290313 | 0.425262743 | 0.623718689 | NIMM-g1->WT |
| d__Bacteria;p__Firmicutes;c__Clostridia;o__Oscillospirales;f__Ruminococcaceae;g__Ruminococcus | 0.795716089 | 0.454527612 | 0.652148313 | HM1->WT |
| Unassigned;__;__;__;__;__ | -0.745262697 | 0.472925771 | 0.655657466 | NIMM-g1->WT |
| d__Bacteria;p__Firmicutes;c__Clostridia;o__Peptostreptococcales-Tissierellales;f__Anaerovoracaceae;g__Family_XIII_AD3011_group | 0.723217783 | 0.476841793 | 0.655657466 | HM1->WT |
| d__Bacteria;p__Actinobacteriota;c__Coriobacteriia;o__Coriobacteriales;f__Eggerthellaceae;g__Gordonibacter | 0.671621236 | 0.509128444 | 0.685764843 | HM1->WT |
| d__Bacteria;p__Firmicutes;c__Clostridia;o__Oscillospirales;f__Ruminococcaceae;g__Incertae_Sedis | 0.605389713 | 0.551035729 | 0.727367162 | HM1->WT |
| d__Bacteria;p__Bacteroidota;c__Bacteroidia;o__Bacteroidales;f__Rikenellaceae;g__Alistipes | -0.588393214 | 0.563288887 | 0.728962089 | NIMM-g1->WT |
| d__Bacteria;p__Firmicutes;c__Clostridia;o__Lachnospirales;f__Lachnospiraceae;g__[Eubacterium]_fissicatena_group | 0.549987283 | 0.588260654 | 0.746638522 | HM1->WT |
| d__Bacteria;p__Firmicutes;c__Clostridia;o__Lachnospirales;f__Lachnospiraceae;g__Sellimonas | -0.540232664 | 0.602026882 | 0.749693854 | NIMM-g1->WT |
| d__Bacteria;p__Verrucomicrobiota;c__Verrucomicrobiae;o__Verrucomicrobiales;f__Akkermansiaceae;g__Akkermansia | -0.479388331 | 0.641158869 | 0.776990326 | NIMM-g1->WT |
| d__Bacteria;p__Firmicutes;c__Clostridia;o__Lachnospirales;f__Lachnospiraceae;g__Epulopiscium | -0.465478698 | 0.647491938 | 0.776990326 | NIMM-g1->WT |
| d__Bacteria;p__Proteobacteria;c__Gammaproteobacteria;o__Burkholderiales;f__Sutterellaceae;g__Parasutterella | -0.39162871 | 0.699670189 | 0.824611294 | NIMM-g1->WT |
| d__Bacteria;p__Firmicutes;c__Clostridia;o__Oscillospirales;f__Oscillospiraceae;g__Colidextribacter | -0.347661696 | 0.731901721 | 0.847465151 | NIMM-g1->WT |
| d__Bacteria;p__Firmicutes;c__Clostridia;o__Lachnospirales;f__Lachnospiraceae;g__Lachnoclostridium | 0.261478293 | 0.796259814 | 0.891299957 | HM1->WT |
| d__Bacteria;p__Firmicutes;c__Clostridia;o__Oscillospirales;f__Oscillospiraceae;g__UCG-005 | -0.260538174 | 0.796768144 | 0.891299957 | NIMM-g1->WT |
| d__Bacteria;p__Firmicutes;c__Clostridia;o__Oscillospirales;f__Ruminococcaceae;g__Anaerotruncus | -0.176877776 | 0.862200909 | 0.935929554 | NIMM-g1->WT |
| d__Bacteria;p__Firmicutes;c__Clostridia;o__Lachnospirales;f__Lachnospiraceae;g__Hungatella | 0.172374826 | 0.865025799 | 0.935929554 | HM1->WT |
| d__Bacteria;p__Firmicutes;c__Clostridia;o__Clostridiales;f__Clostridiaceae;g__Clostridium_sensu_stricto_13 | -0.151918333 | 0.881982787 | 0.938884903 | NIMM-g1->WT |
| d__Bacteria;__;__;__;__;__ | 0.119605242 | 0.906883262 | 0.95006818 | HM1->WT |
| d__Bacteria;p__Firmicutes;c__Clostridia;o__Lachnospirales;f__Lachnospiraceae;g__Lachnospiraceae | -0.064077578 | 0.94947373 | 0.968182957 | NIMM-g1->WT |
| d__Bacteria;p__Firmicutes;c__Clostridia;o__Monoglobales;f__Monoglobaceae;g__Monoglobus | 0.048715273 | 0.961997552 | 0.968182957 | HM1->WT |
| d__Bacteria;p__Firmicutes;c__Clostridia;o__Oscillospirales;f__Ruminococcaceae;g__DTU089 | 0.040487162 | 0.968182957 | 0.968182957 | HM1->WT |

| Operational Taxonomic Unit (Genus) for NIMM-g1->WT vs NIMM-g2->WT Comparison | T-value | P-value | FDR | Enriched |
|---|---|---|---|---|
| d__Bacteria;p__Actinobacteriota;c__Coriobacteriia;o__Coriobacteriales;f__Eggerthellaceae;g__Adlercreutzia | -4.6585333 | 0.00018745 | 0.01237152 | NIMM-g2->WT |
| d__Bacteria;p__Firmicutes;c__Clostridia;o__Oscillospirales;f__Ruminococcaceae;g__Paludicola | -3.895171 | 0.00075895 | 0.02471974 | NIMM-g2->WT |
| d__Bacteria;p__Firmicutes;c__Clostridia;o__Lachnospirales;f__Lachnospiraceae;g__Anaerostipes | -3.7763775 | 0.00112362 | 0.02471974 | NIMM-g2->WT |
| d__Bacteria;p__Bacteroidota;c__Bacteroidia;o__Bacteroidales;f__Muribaculaceae;g__Muribaculaceae | -3.8896756 | 0.00165171 | 0.0272532 | NIMM-g2->WT |
| d__Bacteria;p__Firmicutes;c__Clostridia;o__Oscillospirales;f__Ruminococcaceae;g__Incertae_Sedis | 3.45447145 | 0.00218211 | 0.02880383 | NIMM-g1->WT |
| d__Bacteria;p__Firmicutes;c__Bacilli;o__Staphylococcales;f__Staphylococcaceae;g__Staphylococcus | 3.40265658 | 0.00338824 | 0.03727066 | NIMM-g1->WT |
| d__Bacteria;p__Firmicutes;c__Bacilli;o__Erysipelotrichales;f__Erysipelotrichaceae;g__Holdemania | 3.4466557 | 0.00495422 | 0.04671125 | NIMM-g1->WT |
| d__Bacteria;p__Firmicutes;c__Bacilli;o__Erysipelotrichales;f__Erysipelotrichaceae;g__[Clostridium]_innocuum_group | -2.7147612 | 0.01367792 | 0.11284281 | NIMM-g2->WT |
| d__Bacteria;p__Firmicutes;c__Clostridia;o__Oscillospirales;f__Oscillospiraceae;g__Oscillibacter | 2.65870711 | 0.01971856 | 0.11811064 | NIMM-g1->WT |
| d__Bacteria;p__Firmicutes;c__Clostridia;o__Oscillospirales;f__Ruminococcaceae;g__uncultured | 2.49974674 | 0.02084999 | 0.11811064 | NIMM-g1->WT |
| d__Bacteria;p__Firmicutes;c__Clostridia;o__Oscillospirales;f__Ruminococcaceae;g__Phocea | 2.54080057 | 0.02110542 | 0.11811064 | NIMM-g1->WT |
| d__Bacteria;p__Firmicutes;c__Clostridia;o__Lachnospirales;f__Lachnospiraceae;g__Lachnoclostridium | -2.3964117 | 0.02478916 | 0.11811064 | NIMM-g2->WT |
| d__Bacteria;p__Firmicutes;c__Clostridia;o__Lachnospirales;f__Lachnospiraceae;g__GCA-900066755 | 2.6636851 | 0.02494031 | 0.11811064 | NIMM-g1->WT |
| d__Bacteria;p__Firmicutes;c__Clostridia;o__Lachnospirales;f__Lachnospiraceae;g__[Eubacterium]_fissicatena_group | 2.70750503 | 0.02551587 | 0.11811064 | NIMM-g1->WT |
| d__Bacteria;p__Firmicutes;c__Clostridia;o__Lachnospirales;f__Lachnospiraceae;g__Hungatella | -2.436846 | 0.02684333 | 0.11811064 | NIMM-g2->WT |
| d__Bacteria;p__Firmicutes;c__Clostridia;o__Oscillospirales;f__Oscillospiraceae;g__uncultured | 2.48799161 | 0.02901996 | 0.11970735 | NIMM-g1->WT |
| d__Bacteria;p__Firmicutes;c__Clostridia;o__Lachnospirales;f__Lachnospiraceae;g__Lachnospiraceae_UCG-008 | 2.58053435 | 0.03110141 | 0.12074665 | NIMM-g1->WT |
| d__Bacteria;p__Firmicutes;c__Clostridia;o__Oscillospirales;f__[Eubacterium]_coprostanoligenes_group;g__[Eubacterium]_coprostanoligenes_group | 2.2432036 | 0.03475528 | 0.12743603 | NIMM-g1->WT |
| d__Bacteria;p__Firmicutes;c__Clostridia;o__Oscillospirales;f__Ruminococcaceae;__ | 2.13536693 | 0.04376223 | 0.14761653 | NIMM-g1->WT |
| d__Bacteria;p__Actinobacteriota;c__Coriobacteriia;o__Coriobacteriales;f__Eggerthellaceae;g__Gordonibacter | -2.1235483 | 0.04473228 | 0.14761653 | NIMM-g2->WT |
| d__Bacteria;p__Firmicutes;c__Clostridia;o__Peptostreptococcales-Tissierellales;f__Peptostreptococcaceae;g__Clostridioides | 2.02444618 | 0.06090613 | 0.19141927 | NIMM-g1->WT |
| d__Bacteria;p__Firmicutes;c__Bacilli;o__Erysipelotrichales;f__Erysipelatoclostridiaceae;g__Erysipelatoclostridium | -1.9910908 | 0.07036294 | 0.21108881 | NIMM-g2->WT |
| d__Bacteria;p__Actinobacteriota;c__Coriobacteriia;o__Coriobacteriales;f__Eggerthellaceae;__ | -2.0438036 | 0.08026106 | 0.23031435 | NIMM-g2->WT |
| Unassigned;__;__;__;__;__ | 1.51190276 | 0.15020896 | 0.41307464 | NIMM-g1->WT |
| d__Bacteria;p__Firmicutes;c__Bacilli;o__Erysipelotrichales;f__Erysipelotrichaceae;g__Turicibacter | 1.44940235 | 0.16542237 | 0.43519711 | NIMM-g1->WT |
| d__Bacteria;p__Actinobacteriota;c__Actinobacteria;o__Bifidobacteriales;f__Bifidobacteriaceae;g__Bifidobacterium | 1.42787442 | 0.17144128 | 0.43519711 | NIMM-g1->WT |
| d__Bacteria;p__Firmicutes;c__Clostridia;o__Oscillospirales;f__Butyricicoccaceae;g__Butyricicoccus | 1.39531962 | 0.19909096 | 0.48416507 | NIMM-g1->WT |
| d__Bacteria;p__Firmicutes;c__Clostridia;o__Lachnospirales;f__Lachnospiraceae;g__[Ruminococcus]_torques_group | 1.34841788 | 0.20540336 | 0.48416507 | NIMM-g1->WT |
| d__Bacteria;p__Firmicutes;c__Clostridia;o__Lachnospirales;f__Lachnospiraceae;g__[Ruminococcus]_gnavus_group | 1.24611075 | 0.23202613 | 0.52805947 | NIMM-g1->WT |
| d__Bacteria;p__Verrucomicrobiota;c__Verrucomicrobiae;o__Verrucomicrobiales;f__Akkermansiaceae;g__Akkermansia | -1.1853832 | 0.24804173 | 0.53369154 | NIMM-g2->WT |
| d__Bacteria;p__Bacteroidota;c__Bacteroidia;o__Bacteroidales;f__Rikenellaceae;g__Alistipes | 1.16090678 | 0.25717063 | 0.53369154 | NIMM-g1->WT |
| d__Bacteria;p__Bacteroidota;c__Bacteroidia;o__Bacteroidales;f__Bacteroidaceae;g__Bacteroides | 1.17459067 | 0.25875954 | 0.53369154 | NIMM-g1->WT |
| d__Bacteria;p__Firmicutes;c__Clostridia;o__Clostridia_vadinBB60_group;f__Clostridia_vadinBB60_group;g__Clostridia_vadinBB60_group | 1.12689733 | 0.27275627 | 0.54023714 | NIMM-g1->WT |
| d__Bacteria;p__Firmicutes;c__Clostridia;o__Lachnospirales;f__Lachnospiraceae;g__GCA-900066575 | 1.12953518 | 0.27830398 | 0.54023714 | NIMM-g1->WT |
| d__Bacteria;p__Proteobacteria;c__Gammaproteobacteria;o__Enterobacterales;f__Enterobacteriaceae;g__Escherichia-Shigella | 1.04670003 | 0.30804695 | 0.58088853 | NIMM-g1->WT |
| d__Bacteria;p__Firmicutes;c__Clostridia;o__Oscillospirales;f__Ruminococcaceae;g__Ruminococcus | 1 | 0.33133276 | 0.59102601 | NIMM-g1->WT |
| d__Bacteria;p__Firmicutes;c__Clostridia;o__Oscillospirales;f__Ruminococcaceae;g__DTU089 | 1 | 0.33133276 | 0.59102601 | NIMM-g1->WT |
| d__Bacteria;p__Firmicutes;c__Clostridia;o__Oscillospirales;f__Ruminococcaceae;g__Subdoligranulum | -1 | 0.35061666 | 0.59335128 | NIMM-g2->WT |
| d__Bacteria;p__Firmicutes;c__Clostridia;o__Oscillospirales;f__[Clostridium]_methylpentosum_group;g__[Clostridium]_methylpentosum_group | -1 | 0.35061666 | 0.59335128 | NIMM-g2->WT |
| d__Bacteria;p__Firmicutes;c__Clostridia;o__Lachnospirales;f__Lachnospiraceae;__ | -0.8620836 | 0.39827894 | 0.64772488 | NIMM-g2->WT |
| d__Bacteria;p__Actinobacteriota;c__Coriobacteriia;o__Coriobacteriales;f__Eggerthellaceae;g__Eggerthella | -0.8524972 | 0.40237455 | 0.64772488 | NIMM-g2->WT |
| d__Bacteria;p__Firmicutes;c__Clostridia;o__Peptostreptococcales-Tissierellales;f__Anaerovoracaceae;g__[Eubacterium]_nodatum_group | 0.76647126 | 0.45133413 | 0.68706027 | NIMM-g1->WT |
| d__Bacteria;p__Firmicutes;c__Clostridia;o__Oscillospirales;f__Ruminococcaceae;g__Anaerotruncus | -0.7657525 | 0.45592057 | 0.68706027 | NIMM-g2->WT |
| d__Bacteria;p__Proteobacteria;c__Gammaproteobacteria;o__Vibrionales;f__Vibrionaceae;g__Vibrio | 0.76210553 | 0.45804018 | 0.68706027 | NIMM-g1->WT |
| d__Bacteria;p__Firmicutes;c__Clostridia;o__Oscillospirales;f__Oscillospiraceae;g__Colidextribacter | -0.7220582 | 0.47881533 | 0.70226249 | NIMM-g2->WT |
| d__Bacteria;p__Firmicutes;c__Clostridia;o__Oscillospirales;f__Ruminococcaceae;g__Candidatus_Soleaferrea | 0.65882565 | 0.52504515 | 0.74150447 | NIMM-g1->WT |
| d__Bacteria;p__Firmicutes;c__Bacilli;o__Erysipelotrichales;f__Erysipelotrichaceae;g__Dielma | -0.6287396 | 0.53870953 | 0.74150447 | NIMM-g2->WT |
| d__Bacteria;p__Firmicutes;c__Clostridia;o__Lachnospirales;f__Lachnospiraceae;g__Epulopiscium | 0.62376493 | 0.53927598 | 0.74150447 | NIMM-g1->WT |
| d__Bacteria;p__Firmicutes;c__Clostridia;o__Lachnospirales;f__Lachnospiraceae;g__Lachnospiraceae | 0.5990192 | 0.55522251 | 0.74785073 | NIMM-g1->WT |
| d__Bacteria;p__Firmicutes;c__Clostridia;o__Oscillospirales;f__Ruminococcaceae;g__Faecalibacterium | 0.57913639 | 0.57060582 | 0.75088343 | NIMM-g1->WT |
| d__Bacteria;p__Firmicutes;c__Bacilli;o__Lactobacillales;f__Enterococcaceae;g__Enterococcus | -0.562543 | 0.5802281 | 0.75088343 | NIMM-g2->WT |
| d__Bacteria;p__Actinobacteriota;c__Coriobacteriia;o__Coriobacteriales;f__Coriobacteriaceae;g__Collinsella | -0.4953289 | 0.62555144 | 0.78057489 | NIMM-g2->WT |
| d__Bacteria;p__Firmicutes;c__Clostridia;o__Lachnospirales;f__Lachnospiraceae;g__Blautia | 0.49558995 | 0.62682529 | 0.78057489 | NIMM-g1->WT |
| d__Bacteria;p__Firmicutes;c__Clostridia;o__Clostridiales;f__Clostridiaceae;g__Clostridium_sensu_stricto_13 | 0.43491233 | 0.67007359 | 0.81897883 | NIMM-g1->WT |
| d__Bacteria;p__Firmicutes;c__Clostridia;o__Lachnospirales;f__Lachnospiraceae;g__Eisenbergiella | 0.4092113 | 0.68659084 | 0.82390901 | NIMM-g1->WT |
| d__Bacteria;p__Firmicutes;c__Clostridia;o__Lachnospirales;f__Lachnospiraceae;g__Sellimonas | -0.3649062 | 0.71955064 | 0.84804183 | NIMM-g2->WT |
| d__Bacteria;p__Firmicutes;c__Clostridia;o__Clostridiales;f__Clostridiaceae;g__Clostridium_sensu_stricto_1 | -0.2874422 | 0.77679788 | 0.89945018 | NIMM-g2->WT |
| d__Bacteria;p__Firmicutes;c__Clostridia;o__Monoglobales;f__Monoglobaceae;g__Monoglobus | 0.2681524 | 0.79087335 | 0.89995933 | NIMM-g1->WT |
| d__Bacteria;p__Proteobacteria;c__Gammaproteobacteria;o__Burkholderiales;f__Sutterellaceae;g__Parasutterella | 0.23732631 | 0.81495445 | 0.9097345 | NIMM-g1->WT |
| d__Bacteria;p__Firmicutes;c__Clostridia;o__Oscillospirales;f__Oscillospiraceae;g__UCG-005 | 0.22076507 | 0.8271447 | 0.9097345 | NIMM-g1->WT |
| d__Bacteria;p__Firmicutes;c__Clostridia;o__Oscillospirales;f__Ruminococcaceae;g__UBA1819 | -0.2032767 | 0.84081522 | 0.9097345 | NIMM-g2->WT |
| d__Bacteria;p__Firmicutes;c__Clostridia;o__Oscillospirales;f__Oscillospiraceae;g__Flavonifractor | -0.1764315 | 0.86263069 | 0.91828428 | NIMM-g2->WT |
| d__Bacteria;p__Bacteroidota;c__Bacteroidia;o__Bacteroidales;f__Tannerellaceae;g__Parabacteroides | 0.10755545 | 0.91525449 | 0.95835708 | NIMM-g1->WT |
| d__Bacteria;p__Firmicutes;c__Clostridia;o__Lachnospirales;f__Lachnospiraceae;g__Lachnospiraceae_NK4A136_group | 0.08699432 | 0.93166198 | 0.95835708 | NIMM-g1->WT |
| d__Bacteria;__;__;__;__;__ | -0.0717879 | 0.94383652 | 0.95835708 | NIMM-g2->WT |
| d__Bacteria;p__Firmicutes;c__Clostridia;o__Peptostreptococcales-Tissierellales;f__Anaerovoracaceae;g__Family_XIII_AD3011_group | -0.0301803 | 0.97638519 | 0.97638519 | NIMM-g2->WT |

| Operational Taxonomic Unit (Genus) for NIMM-g1->KO vs NIMM-g1->WT Comparison | T-value | P-value | FDR | Enriched |
|---|---|---|---|---|
| d__Bacteria;p__Firmicutes;c__Clostridia;o__Lachnospirales;f__Lachnospiraceae;g__[Ruminococcus]_gnavus_group | 9.747517304 | 6.73E-09 | 4.44E-07 | NIMM-g1->KO |
| d__Bacteria;p__Firmicutes;c__Bacilli;o__Erysipelotrichales;f__Erysipelotrichaceae;g__[Clostridium]_innocuum_group | 4.583874401 | 0.000159032 | 0.005248049 | NIMM-g1->KO |
| d__Bacteria;p__Firmicutes;c__Clostridia;o__Lachnospirales;f__Lachnospiraceae;g__Lachnospiraceae_UCG-008 | -4.823476523 | 0.000868157 | 0.019099452 | NIMM-g1->WT |
| d__Bacteria;p__Firmicutes;c__Clostridia;o__Monoglobales;f__Monoglobaceae;g__Monoglobus | -3.556325945 | 0.001707663 | 0.025462462 | NIMM-g1->WT |
| d__Bacteria;p__Firmicutes;c__Clostridia;o__Oscillospirales;f__[Eubacterium]_coprostanoligenes_group;g__[Eubacterium]_coprostanoligenes_group | -3.631690877 | 0.002062062 | 0.025462462 | NIMM-g1->WT |
| d__Bacteria;p__Firmicutes;c__Clostridia;o__Oscillospirales;f__Ruminococcaceae;__ | -3.578429343 | 0.002314769 | 0.025462462 | NIMM-g1->WT |
| d__Bacteria;p__Firmicutes;c__Clostridia;o__Lachnospirales;f__Lachnospiraceae;g__Blautia | -3.040965083 | 0.005933843 | 0.055947659 | NIMM-g1->WT |
| d__Bacteria;p__Firmicutes;c__Clostridia;o__Lachnospirales;f__Lachnospiraceae;g__Lachnospiraceae_NK4A136_group | -2.879285755 | 0.017875566 | 0.147473423 | NIMM-g1->WT |
| d__Bacteria;p__Firmicutes;c__Clostridia;o__Clostridia_vadinBB60_group;f__Clostridia_vadinBB60_group;g__Clostridia_vadinBB60_group | -2.513411883 | 0.022326006 | 0.163724046 | NIMM-g1->WT |
| d__Bacteria;p__Proteobacteria;c__Gammaproteobacteria;o__Vibrionales;f__Vibrionaceae;g__Vibrio | -2.381112305 | 0.027144826 | 0.179155854 | NIMM-g1->WT |
| d__Bacteria;p__Firmicutes;c__Bacilli;o__Staphylococcales;f__Staphylococcaceae;g__Staphylococcus | -2.200388204 | 0.038382149 | 0.20808993 | NIMM-g1->WT |
| d__Bacteria;p__Firmicutes;c__Clostridia;o__Oscillospirales;f__Ruminococcaceae;g__Faecalibacterium | -2.200106398 | 0.041915882 | 0.20808993 | NIMM-g1->WT |
| d__Bacteria;__;__;__;__;__ | -2.174487418 | 0.044078131 | 0.20808993 | NIMM-g1->WT |
| d__Bacteria;p__Firmicutes;c__Clostridia;o__Lachnospirales;f__Lachnospiraceae;g__Hungatella | 2.294676006 | 0.045278876 | 0.20808993 | NIMM-g1->KO |
| d__Bacteria;p__Firmicutes;c__Clostridia;o__Clostridiales;f__Clostridiaceae;g__Clostridium_sensu_stricto_1 | 2.304306082 | 0.047293166 | 0.20808993 | NIMM-g1->KO |
| Unassigned;__;__;__;__;__ | -2.058412178 | 0.054260084 | 0.223822846 | NIMM-g1->WT |
| d__Bacteria;p__Proteobacteria;c__Gammaproteobacteria;o__Burkholderiales;f__Sutterellaceae;g__Parasutterella | -1.957228038 | 0.066947315 | 0.246400112 | NIMM-g1->WT |
| d__Bacteria;p__Firmicutes;c__Clostridia;o__Oscillospirales;f__Ruminococcaceae;g__UBA1819 | -1.924446848 | 0.068185995 | 0.246400112 | NIMM-g1->WT |
| d__Bacteria;p__Actinobacteriota;c__Coriobacteriia;o__Coriobacteriales;f__Coriobacteriaceae;g__Collinsella | 1.994736632 | 0.070933365 | 0.246400112 | NIMM-g1->KO |
| d__Bacteria;p__Bacteroidota;c__Bacteroidia;o__Bacteroidales;f__Muribaculaceae;g__Muribaculaceae | -1.794910476 | 0.090468043 | 0.298544541 | NIMM-g1->WT |
| d__Bacteria;p__Firmicutes;c__Clostridia;o__Oscillospirales;f__Ruminococcaceae;g__Incertae_Sedis | -1.897592221 | 0.10286771 | 0.305909421 | NIMM-g1->WT |
| d__Bacteria;p__Firmicutes;c__Clostridia;o__Lachnospirales;f__Lachnospiraceae;g__Epulopiscium | -1.71650829 | 0.104232808 | 0.305909421 | NIMM-g1->WT |
| d__Bacteria;p__Firmicutes;c__Clostridia;o__Lachnospirales;f__Lachnospiraceae;g__Lachnospiraceae | -1.744208467 | 0.106604798 | 0.305909421 | NIMM-g1->WT |
| d__Bacteria;p__Firmicutes;c__Bacilli;o__Erysipelotrichales;f__Erysipelotrichaceae;g__Dielma | -1.674504721 | 0.113254076 | 0.307069328 | NIMM-g1->WT |
| d__Bacteria;p__Firmicutes;c__Clostridia;o__Oscillospirales;f__Ruminococcaceae;g__Anaerotruncus | 1.713621156 | 0.116314139 | 0.307069328 | NIMM-g1->KO |
| d__Bacteria;p__Bacteroidota;c__Bacteroidia;o__Bacteroidales;f__Bacteroidaceae;g__Bacteroides | -1.561384996 | 0.138351126 | 0.351199013 | NIMM-g1->WT |
| d__Bacteria;p__Firmicutes;c__Clostridia;o__Lachnospirales;f__Lachnospiraceae;g__[Ruminococcus]_torques_group | -1.48006807 | 0.161093403 | 0.3764785 | NIMM-g1->WT |
| d__Bacteria;p__Firmicutes;c__Bacilli;o__Erysipelotrichales;f__Erysipelatoclostridiaceae;g__Erysipelatoclostridium | 1.539198792 | 0.16216034 | 0.3764785 | NIMM-g1->KO |
| d__Bacteria;p__Firmicutes;c__Bacilli;o__Erysipelotrichales;f__Erysipelotrichaceae;g__Turicibacter | -1.449402349 | 0.165422371 | 0.3764785 | NIMM-g1->WT |
| d__Bacteria;p__Firmicutes;c__Clostridia;o__Lachnospirales;f__Lachnospiraceae;g__GCA-900066755 | -1.371678652 | 0.197749065 | 0.435047943 | NIMM-g1->WT |
| d__Bacteria;p__Actinobacteriota;c__Coriobacteriia;o__Coriobacteriales;f__Eggerthellaceae;g__Gordonibacter | -1.253586731 | 0.222613633 | 0.473951606 | NIMM-g1->WT |
| d__Bacteria;p__Firmicutes;c__Clostridia;o__Oscillospirales;f__Ruminococcaceae;g__Candidatus_Soleaferrea | -1.163419826 | 0.275713582 | 0.548036428 | NIMM-g1->WT |
| d__Bacteria;p__Firmicutes;c__Clostridia;o__Lachnospirales;f__Lachnospiraceae;g__[Eubacterium]_fissicatena_group | -1.152886227 | 0.28006691 | 0.548036428 | NIMM-g1->WT |
| d__Bacteria;p__Firmicutes;c__Clostridia;o__Oscillospirales;f__Oscillospiraceae;g__uncultured | -1.132695171 | 0.282321796 | 0.548036428 | NIMM-g1->WT |
| d__Bacteria;p__Firmicutes;c__Clostridia;o__Lachnospirales;f__Lachnospiraceae;g__Sellimonas | -1.087198688 | 0.310640267 | 0.582743739 | NIMM-g1->WT |
| d__Bacteria;p__Firmicutes;c__Clostridia;o__Oscillospirales;f__Ruminococcaceae;g__Ruminococcus | -1 | 0.331332762 | 0.582743739 | NIMM-g1->WT |
| d__Bacteria;p__Firmicutes;c__Clostridia;o__Lachnospirales;f__Lachnospiraceae;g__Eisenbergiella | 0.970670853 | 0.344043706 | 0.582743739 | NIMM-g1->KO |
| d__Bacteria;p__Firmicutes;c__Clostridia;o__Lachnospirales;f__Lachnospiraceae;g__Agathobacter | 1 | 0.355917684 | 0.582743739 | NIMM-g1->KO |
| d__Bacteria;p__Firmicutes;c__Clostridia;o__Oscillospirales;f__[Clostridium]_methylpentosum_group;g__[Clostridium]_methylpentosum_group | 1 | 0.355917684 | 0.582743739 | NIMM-g1->KO |
| d__Bacteria;p__Actinobacteriota;c__Coriobacteriia;o__Coriobacteriales;f__Eggerthellaceae;__ | 1 | 0.355917684 | 0.582743739 | NIMM-g1->KO |
| d__Bacteria;p__Proteobacteria;c__Gammaproteobacteria;o__Enterobacterales;f__Enterobacteriaceae;g__Escherichia-Shigella | -0.928490344 | 0.367854278 | 0.582743739 | NIMM-g1->WT |
| d__Bacteria;p__Firmicutes;c__Clostridia;o__Oscillospirales;f__Oscillospiraceae;g__UCG-005 | -0.928463849 | 0.370836925 | 0.582743739 | NIMM-g1->WT |
| d__Bacteria;p__Firmicutes;c__Bacilli;o__Erysipelotrichales;f__Erysipelotrichaceae;g__Holdemania | 0.826996194 | 0.427102126 | 0.6555521 | NIMM-g1->KO |
| d__Bacteria;p__Firmicutes;c__Clostridia;o__Oscillospirales;f__Ruminococcaceae;g__Phocea | 0.770545325 | 0.460627699 | 0.676345039 | NIMM-g1->KO |
| d__Bacteria;p__Firmicutes;c__Clostridia;o__Lachnospirales;f__Lachnospiraceae;g__Lachnoclostridium | -0.750606434 | 0.461144345 | 0.676345039 | NIMM-g1->WT |
| d__Bacteria;p__Firmicutes;c__Clostridia;o__Oscillospirales;f__Ruminococcaceae;g__uncultured | -0.741531932 | 0.471685184 | 0.676765698 | NIMM-g1->WT |
| d__Bacteria;p__Firmicutes;c__Clostridia;o__Oscillospirales;f__Oscillospiraceae;g__Oscillibacter | 0.653705889 | 0.529535731 | 0.743603366 | NIMM-g1->KO |
| d__Bacteria;p__Firmicutes;c__Clostridia;o__Oscillospirales;f__Ruminococcaceae;g__DTU089 | 0.628660617 | 0.548469123 | 0.745117315 | NIMM-g1->KO |
| d__Bacteria;p__Firmicutes;c__Clostridia;o__Oscillospirales;f__Ruminococcaceae;g__Paludicola | 0.592159563 | 0.56442058 | 0.745117315 | NIMM-g1->KO |
| d__Bacteria;p__Firmicutes;c__Clostridia;o__Oscillospirales;f__Oscillospiraceae;g__Flavonifractor | -0.587725192 | 0.564482814 | 0.745117315 | NIMM-g1->WT |
| d__Bacteria;p__Firmicutes;c__Clostridia;o__Peptostreptococcales-Tissierellales;f__Peptostreptococcaceae;g__Clostridioides | -0.580659446 | 0.579961533 | 0.750538454 | NIMM-g1->WT |
| d__Bacteria;p__Firmicutes;c__Clostridia;o__Peptostreptococcales-Tissierellales;f__Anaerovoracaceae;g__Family_XIII_AD3011_group | -0.544174918 | 0.598025363 | 0.759032192 | NIMM-g1->WT |
| d__Bacteria;p__Firmicutes;c__Clostridia;o__Oscillospirales;f__Oscillospiraceae;g__Colidextribacter | -0.51542006 | 0.612471653 | 0.762700549 | NIMM-g1->WT |
| d__Bacteria;p__Firmicutes;c__Clostridia;o__Lachnospirales;f__Lachnospiraceae;g__Anaerostipes | 0.489935148 | 0.638669325 | 0.774242231 | NIMM-g1->KO |
| d__Bacteria;p__Firmicutes;c__Clostridia;o__Lachnospirales;f__Lachnospiraceae;g__GCA-900066575 | -0.468006719 | 0.645201859 | 0.774242231 | NIMM-g1->WT |
| d__Bacteria;p__Actinobacteriota;c__Coriobacteriia;o__Coriobacteriales;f__Eggerthellaceae;g__Eggerthella | 0.375326449 | 0.710874832 | 0.832007868 | NIMM-g1->KO |
| d__Bacteria;p__Bacteroidota;c__Bacteroidia;o__Bacteroidales;f__Tannerellaceae;g__Parabacteroides | -0.369345811 | 0.718552249 | 0.832007868 | NIMM-g1->WT |
| d__Bacteria;p__Firmicutes;c__Clostridia;o__Lachnospirales;f__Lachnospiraceae;__ | -0.33246333 | 0.742570256 | 0.84499374 | NIMM-g1->WT |
| d__Bacteria;p__Firmicutes;c__Clostridia;o__Peptostreptococcales-Tissierellales;f__Anaerovoracaceae;g__[Eubacterium]_nodatum_group | -0.25640813 | 0.805336608 | 0.900885019 | NIMM-g1->WT |
| d__Bacteria;p__Actinobacteriota;c__Actinobacteria;o__Bifidobacteriales;f__Bifidobacteriaceae;g__Bifidobacterium | -0.203624349 | 0.840545253 | 0.910427701 | NIMM-g1->WT |
| d__Bacteria;p__Bacteroidota;c__Bacteroidia;o__Bacteroidales;f__Rikenellaceae;g__Alistipes | -0.194816141 | 0.848360855 | 0.910427701 | NIMM-g1->WT |
| d__Bacteria;p__Firmicutes;c__Clostridia;o__Oscillospirales;f__Butyricicoccaceae;g__Butyricicoccus | 0.163676679 | 0.871421269 | 0.910427701 | NIMM-g1->KO |
| d__Bacteria;p__Firmicutes;c__Clostridia;o__Clostridiales;f__Clostridiaceae;g__Clostridium_sensu_stricto_13 | -0.151100628 | 0.882573638 | 0.910427701 | NIMM-g1->WT |
| d__Bacteria;p__Firmicutes;c__Bacilli;o__Lactobacillales;f__Enterococcaceae;g__Enterococcus | 0.150563299 | 0.882838983 | 0.910427701 | NIMM-g1->KO |
| d__Bacteria;p__Actinobacteriota;c__Coriobacteriia;o__Coriobacteriales;f__Eggerthellaceae;g__Adlercreutzia | -0.053127443 | 0.958447614 | 0.973192962 | NIMM-g1->WT |
| d__Bacteria;p__Verrucomicrobiota;c__Verrucomicrobiae;o__Verrucomicrobiales;f__Akkermansiaceae;g__Akkermansia | -0.030585223 | 0.975866273 | 0.975866273 | NIMM-g1->WT |
