## Supplemental File 1 for "Mouse Adaptation of Human Inflammatory Bowel Diseases Microbiota Enhances Colonization Efficiency and Alters Microbiome Aggressiveness Depending on Recipient Colonic Inflammatory Environment"

### Analysis of FMT Model for Humanized Mice

#### Introduction

In our examination of a more stable and reproducible fecal microbiome transfer (FMT) model between human and mice, we used 16S rRNA amplicon sequence data of stool samples from both humans and mice throughout various iterations of fecal transfers. The different iterations of fecal transfers include generational, serial transfers (human to mouse, human-adapted mouse to mouse) and in two different host environments (inflamed and non-inflamed).

This notebook includes R scripts that generated PCoA plots, boxplots, and scatterplots presented in the published manuscript.

Taxonomic barplots and alpha diversity plots were generated through the Plotmicrobiome application.

A core set of figures from the manuscript were independently replicated by Jack Young using Python; both the Python scripts and replicated figures are provided in a Jupyter Notebook file (PDF format).

The Github link to the Plotmicrobiome application and the Python validation files are provided at the end of our manuscript.

Please refer to the ReadMe file on our Github respository for additional notes on running the scripts in this R notebook.

#### Analysis Overview

Each code chunk follows a step of the analysis pipeline:

- Installing all relevant libraries
- Data import and parsing
- PCoA Plots
- Transfer efficiency calculations of samples within a given FMT group
- Transfer efficiency calculations of samples across a pair of FMT groups
- Visualizing the non-transferred ASV abundance in samples across a pair of FMT groups
- Testing homogeneity of variances among histology tissue scores
- Visualizing histology tissue scores as boxplots (could probably omit this chunk)
- Assessing cage effect on transfer efficiency of samples within groups
- Assessing protein assay data (fLCN2)

##### Installing libraries

The code below insalls and loads all dependencies used throughout this analysis.

```
packagesToInstall = c("ggplot2", "vegan", "dplyr", "testthat", "stats","car","here")
install.packages(packagesToInstall,repos = "http://cran.us.r-project.org")
```

```
## Warning: package 'stats' is in use and will not be installed
```

```
## Warning: cannot remove prior installation of package 'vegan'
```

```
## Warning in file.copy(savedcopy, lib, recursive = TRUE): problem copying
## C:\Users\aduong\AppData\Local\R\win-library\4.3\00LOCK\vegan\libs\x64\vegan.dll
## to C:\Users\aduong\AppData\Local\R\win-library\4.3\vegan\libs\x64\vegan.dll:
## Permission denied
```

```
## Warning: restored 'vegan'
```

```
## Warning: cannot remove prior installation of package 'dplyr'
```

```
## Warning in file.copy(savedcopy, lib, recursive = TRUE): problem copying
## C:\Users\aduong\AppData\Local\R\win-library\4.3\00LOCK\dplyr\libs\x64\dplyr.dll
## to C:\Users\aduong\AppData\Local\R\win-library\4.3\dplyr\libs\x64\dplyr.dll:
## Permission denied
```

```
## Warning: restored 'dplyr'
```

```
## Warning: cannot remove prior installation of package 'testthat'
```

```
## Warning in file.copy(savedcopy, lib, recursive = TRUE): problem copying
## C:\Users\aduong\AppData\Local\R\win-library\4.3\00LOCK\testthat\libs\x64\testthat.dll
## to
## C:\Users\aduong\AppData\Local\R\win-library\4.3\testthat\libs\x64\testthat.dll:
## Permission denied
```

```
## Warning: restored 'testthat'
```

```
lapply(packagesToInstall, library, character.only=TRUE)
```

```
## Warning: package 'ggplot2' was built under R version 4.3.1
```

```
## Warning: package 'vegan' was built under R version 4.3.1
```

```
## Warning: package 'dplyr' was built under R version 4.3.1
```

```
## Warning: package 'testthat' was built under R version 4.3.1
```

```
## Warning: package 'car' was built under R version 4.3.1
```

```
## Warning: package 'here' was built under R version 4.3.1
```

##### Data import and parsing

Next, the script below reads in all the functions that are used throughout this analysis.

We recommend organizing these functions, which exist as external, independent R scripts, in the same directory as the R Notebook file.

```
allScripts = c("lognorm_function.R","generatePCOAPlots.R",
               "getCorrelationCoefficient_acrossGroup.R",
               "getCorrelationCoefficient_withinGroup.R",
               "getCorrelationCoefficient_withinGroup_phylumLevel_byGroup.R",
               "getCorrelationCoefficient_withinGroup_perCage.R",
               "generateTableWithTaxonomy_NontransferringASVS_samplePairsAcrossGroup.R",
               "generateScatterPlots_samplePairsAcrossGroup_histogramOfNontransferringASVS.R",
               "calculateBartlettPvalue_histologyScoreData.R","calculateLevenesTestPvalue_histologyScoreData.R",
               "generateScatterplots_acrossGroup.R"
               )

for(i in 1:length(allScripts)){
  source(allScripts[i], local = knitr::knit_global())
}
```

The code below reads in the counts data and metadata files, starting with the counts data at the ASV level, with a total sample size 110 sequenced samples. This includes three technical replicates of pooled human donor slurries and for each input mice slurry group.

QIIME2 generates counts table in a feature x sample format, in which features (taxa) are in rows and samples are columns. In our first step of reading in these counts data files into our R scripts, we maintained the format integrity (features as rows and samples as columns). Later, we transpose the counts table to match the format of the metadata file, which has samples as rows and features as columns.

Note: Early in our analysis, we updated the nomenclature system of our FMT groups to improve readability. Hence, we provide the metadata with the original and the updated nomenclature, the latter which we use for our published results and manuscript. In addition to updating nomenclature, which we updated in Excel, we formatted the sample names to make it R-friendly (i.e. change "-" into "." in sample names because R interprets "-" between integers as a mathematics operation). If original counts tables are read into Python, these changes are not needed. The Python validation scripts uses the metadata and counts files with the original nomenclature.

```
file_asv_updatedName = read.table(here("files","countsTable_asv_importToR_updatedNomenclature.txt"), header = T,
sep = "\t")
metamapping_updatedName = read.table(here("files","mappingMetadata_updatedNomenclature.txt"), header = T, sep = "
\t")
asv_taxonomy_map= read.table(here("files","asv_taxonomy_map.txt"), sep = "\t", header = TRUE)

## group labels
labels_inputRecipientGroups = unique(metamapping_updatedName$FMTGroupFMTsource..Recipientbackground)

file_asv_updatedName_t = t(file_asv_updatedName)
file_asv_updatedName_t_samplesOnly = file_asv_updatedName_t[-(c(1,nrow(file_asv_updatedName_t))),]
colnames(file_asv_updatedName_t_samplesOnly) = file_asv_updatedName_t[1,]


file_asv_updatedName_t_samplesOnly_toMerge = as.data.frame(file_asv_updatedName_t_samplesOnly)
file_asv_updatedName_t_samplesOnly_toMerge$SampleID = rownames(file_asv_updatedName_t_samplesOnly)

#merge counts and metadata files
myASVandMetadata_newNames = full_join(file_asv_updatedName_t_samplesOnly_toMerge, metamapping_updatedName, by="Sa
mpleID")

names(myASVandMetadata_newNames)[names(myASVandMetadata_newNames)=="FMTGroupFMTsource..Recipientbackground"] = "F
MTGroupFMTsourcegtRecipientbackground"

##lognorm
file_asv_updatedName_t_samplesOnly_df = as.data.frame(file_asv_updatedName_t_samplesOnly)
file_asv_updatedName_t_samplesOnly_df[] <- lapply(file_asv_updatedName_t_samplesOnly_df, function(x) as.numeric(a
s.character(x)))

file_asv_updatedName_t_samplesOnly_lognorm = lognorm_function(file_asv_updatedName_t_samplesOnly_df)

# serial passage groups
labels_NIMM_lineage = c("HM1input", "HM1->WT", "NIMM-g1input", "NIMM-g1->WT", "NIMM-g2input", "NIMM-g2->WT")
labels_IMM_lineage = c("HM1input", "HM1->KO", "IMM-g1input", "IMM-g1->KO","IMM-g2input", "IMM-g2->KO")
```

The code below parses and log normalizes the genus counts table:

```
file_genusCountsTable_updatedName = read.table(here("files","countsTable_genus_importToR_updatedNomenclature.tx
t"), header = T, sep = "\t")
metamapping_updatedName = read.table(here("files","mappingMetadata_updatedNomenclature.txt"), header = T, sep = "
\t")

file_genusCountsTable_updatedName_t = t(file_genusCountsTable_updatedName)

genusCountsTable_samplebyfeatureFormat = file_genusCountsTable_updatedName_t[-1,]
colnames(genusCountsTable_samplebyfeatureFormat) = file_genusCountsTable_updatedName_t[1,]

genusCounts_samplesOnly_toMerge = as.data.frame(genusCountsTable_samplebyfeatureFormat)
genusCounts_samplesOnly_toMerge$SampleID = rownames(genusCountsTable_samplebyfeatureFormat)

#merge counts and metadata files
genusCountsandMetadata = full_join(genusCounts_samplesOnly_toMerge, metamapping_updatedName, by="SampleID")

names(genusCountsandMetadata)[names(genusCountsandMetadata)=="FMTGroupFMTsource..Recipientbackground"] = "FMTGrou
pFMTsourcegtRecipientbackground"

##lognorm
genusCounts_samplesOnly_df = as.data.frame(genusCountsTable_samplebyfeatureFormat)
genusCounts_samplesOnly_df[] <- lapply(genusCounts_samplesOnly_df, function(x) as.numeric(as.character(x)))

genusCounts_samplesOnly_df_lognorm = lognorm_function(genusCounts_samplesOnly_df)
```

Below, the code parses and log normalizes the phylum counts table:

```
file_phylumCountsTable_updatedName = read.table(here("files","countsTable_phylum_importToR_updatedNomenclature.tx
t"), header = T, sep = "\t")
metamapping_updatedName = read.table(here("files","mappingMetadata_updatedNomenclature.txt"), header = T, sep = "
\t")

file_phylumCountsTable_updatedName_t = t(file_phylumCountsTable_updatedName)

phylumCountsTable_samplebyfeatureFormat = file_phylumCountsTable_updatedName_t[-1,]
colnames(phylumCountsTable_samplebyfeatureFormat) = file_phylumCountsTable_updatedName_t[1,]

phylumCounts_samplesOnly_toMerge = as.data.frame(phylumCountsTable_samplebyfeatureFormat)
phylumCounts_samplesOnly_toMerge$SampleID = rownames(phylumCountsTable_samplebyfeatureFormat)

#merge counts and metadata files
phylumCountsandMetadata = full_join(phylumCounts_samplesOnly_toMerge, metamapping_updatedName, by="SampleID")

names(phylumCountsandMetadata)[names(phylumCountsandMetadata)=="FMTGroupFMTsource..Recipientbackground"] = "FMTGr
oupFMTsourcegtRecipientbackground"

##lognorm
phylumCounts_samplesOnly_df = as.data.frame(phylumCountsTable_samplebyfeatureFormat)
phylumCounts_samplesOnly_df[] <- lapply(phylumCounts_samplesOnly_df, function(x) as.numeric(as.character(x)))

phylumCounts_samplesOnly_df_lognorm = lognorm_function(phylumCounts_samplesOnly_df)
```

##### PCoA Plots

PCoA plots were generated with the capscale function from the vegan package using Bray-Curtis dissimilarity matrix.

```
dir.create(here("PcoaPlots"))
```

```
## Warning in dir.create(here("PcoaPlots")):
## 'C:\Users\aduong\Desktop\Balfour2022\PcoaPlots' already exists
```

```
plotColors_ver2 = c("gray65", "royalblue1", "green", "orange", "purple",
                    "black", "olivedrab", "red", "red4",
                      "cyan","palevioletred1", "blue3", "yellow2", "magenta")


metagroup = factor(myASVandMetadata_newNames$FMTGroupFMTsourcegtRecipientbackground)

micegroups_all = c("HM1->KO", "HM1->WT", "IMM-g1->KO", "IMM-g2->KO", "NIMM-g1->WT","NIMM-g2->WT")
micegroups_inflamed = c("HM1->KO", "IMM-g1->KO","IMM-g2->KO", "IMM-g1input", "IMM-g2input")
micegroups_noninflamed = c("HM1->WT", "NIMM-g1->WT","NIMM-g2->WT", "NIMM-g1input", "NIMM-g2input")
excludecrossoverGroup = unique(genusCountsandMetadata$FMTGroupFMTsourcegtRecipientbackground)
excludecrossoverGroup = excludecrossoverGroup[-11]
micegroups_inflamed_withbothHMCohorts = c("HM1->KO", "HM2->KO","IMM-g1->KO","IMM-g2->KO", "IMM-g1input", "IMM-g2i
nput")
micegroups_serialTransfersPlusCrossover = c("HM1->KO", "HM1->WT", "IMM-g1->KO", "IMM-g2->KO", "NIMM-g1->WT","NIMM
-g2->WT", "NIMM-g1->KO")

micegroups_all.samplesindex = genusCountsandMetadata$SampleID[genusCountsandMetadata$FMTGroupFMTsourcegtRecipient
background %in% micegroups_all]
micegroups_inflamed.samplesindex = genusCountsandMetadata$SampleID[genusCountsandMetadata$FMTGroupFMTsourcegtReci
pientbackground %in% micegroups_inflamed]
micegroups_noninflamed.samplesindex = genusCountsandMetadata$SampleID[genusCountsandMetadata$FMTGroupFMTsourcegtR
ecipientbackground %in% micegroups_noninflamed]
excludcrossoverGroup.samplesindex = genusCountsandMetadata$SampleID[genusCountsandMetadata$FMTGroupFMTsourcegtRec
ipientbackground %in% excludecrossoverGroup]
micegroups_inflamed_withbothHMCohorts.samplesindex = genusCountsandMetadata$SampleID[genusCountsandMetadata$FMTGr
oupFMTsourcegtRecipientbackground %in% micegroups_inflamed_withbothHMCohorts]
micegroups_serialTransfersPlusCrossover.samplesindex = genusCountsandMetadata$SampleID[genusCountsandMetadata$FMT
GroupFMTsourcegtRecipientbackground %in% micegroups_serialTransfersPlusCrossover]

micegroups_all.metagroup = factor(metagroup[metagroup %in% micegroups_all])
micegroups_inflamed.metagroup = factor(metagroup[metagroup %in% micegroups_inflamed])
micegroups_noninflamed.metagroup = factor(metagroup[metagroup %in% micegroups_noninflamed])
excludecrossoverGroup.metagroup = factor(metagroup[metagroup %in% excludecrossoverGroup])
micegroups_inflamed_withbothHMCohorts.metagroup = factor(metagroup[metagroup %in% micegroups_inflamed_withbothHMC
ohorts])
micegroups_serialTransfersPlusCrossover.metagroup = factor(metagroup[metagroup %in% micegroups_serialTransfersPlu
sCrossover])

  # all mice groups
micegroups_all_genusCounts_samplesOnly_lognorm_df = genusCounts_samplesOnly_df_lognorm[rownames(genusCounts_sampl
esOnly_df_lognorm) %in% micegroups_all.samplesindex,]
micegroups_all.metagroup
micegroups_all_colors = plotColors_ver2[levels(metagroup) %in% micegroups_all]
  # KO mice groups
micegroups_inflamed_genusCounts_samplesOnly_lognorm_df = genusCounts_samplesOnly_df_lognorm[rownames(genusCounts_
samplesOnly_df_lognorm) %in% micegroups_inflamed.samplesindex,]
micegroups_inflamed.metagroup
micegroups_inflamed_colors = plotColors_ver2[levels(metagroup) %in% micegroups_inflamed]
  # WT mice groups
micegroups_noninflamed_genusCounts_samplesOnly_lognorm_df = genusCounts_samplesOnly_df_lognorm[rownames(genusCoun
ts_samplesOnly_df_lognorm) %in% micegroups_noninflamed.samplesindex,]
micegroups_noninflamed.metagroup
micegroups_noninflamed_colors = plotColors_ver2[levels(metagroup) %in% micegroups_noninflamed]
  ## excluding crossover group
excludecrossoverGroup_genusCounts_samplesOnly_lognorm_df = genusCounts_samplesOnly_df_lognorm[rownames(genusCount
s_samplesOnly_df_lognorm) %in% excludcrossoverGroup.samplesindex,]
excludecrossoverGroup.metagroup
excludecrossoverGroup_colors = plotColors_ver2[levels(metagroup) %in% excludecrossoverGroup]
  # KO mice groups with both HM cohorts
micegroups_inflamed_withbothHMCohorts_genusCounts_samplesOnly_lognorm_df = genusCounts_samplesOnly_df_lognorm[row
names(genusCounts_samplesOnly_df_lognorm) %in% micegroups_inflamed_withbothHMCohorts.samplesindex,]
micegroups_inflamed_withbothHMCohorts.metagroup
micegroups_inflamed_withbothHMCohorts_colors = plotColors_ver2[levels(metagroup) %in% micegroups_inflamed_withbot
hHMCohorts]

## serial transfer groups plus crossover group
micegroups_serialTransfersPlusCrossover_genusCounts_samplesOnly_lognorm_df = genusCounts_samplesOnly_df_lognorm[r
ownames(genusCounts_samplesOnly_df_lognorm) %in% micegroups_serialTransfersPlusCrossover.samplesindex,]
micegroups_serialTransfersPlusCrossover.metagroup
micegroups_serialTransfersPlusCrossover_colors = plotColors_ver2[levels(metagroup) %in% micegroups_serialTransfer
sPlusCrossover]


pdf(here("PcoaPlots","PCoAPlots.pdf"))
generatePCOAPlots(micegroups_serialTransfersPlusCrossover_genusCounts_samplesOnly_lognorm_df,
                  micegroups_serialTransfersPlusCrossover.metagroup,
                  micegroups_serialTransfersPlusCrossover_colors, legendlocation = "topright",
                  plotTitle="Mice Groups, Including Crossover Group")

generatePCOAPlots(micegroups_inflamed_withbothHMCohorts_genusCounts_samplesOnly_lognorm_df,
                  micegroups_inflamed_withbothHMCohorts.metagroup,
                  micegroups_inflamed_withbothHMCohorts_colors, legendlocation = "topleft", plotTitle="Inflamed M
ice Groups, Both HM Cohorts")

generatePCOAPlots(genusCounts_samplesOnly_df_lognorm, metagroup,
                  plotColors_ver2, legendlocation = "bottomleft", plotTitle="All Groups")

generatePCOAPlots(micegroups_all_genusCounts_samplesOnly_lognorm_df, micegroups_all.metagroup,
                  micegroups_all_colors, legendlocation = "topright", plotTitle="Mice Groups Only")

generatePCOAPlots(excludecrossoverGroup_genusCounts_samplesOnly_lognorm_df, excludecrossoverGroup.metagroup,
                  excludecrossoverGroup_colors, legendlocation = "topleft", plotTitle="Excluding Cross-Over Grou
p")

generatePCOAPlots(micegroups_noninflamed_genusCounts_samplesOnly_lognorm_df, micegroups_noninflamed.metagroup,
                  micegroups_noninflamed_colors, legendlocation = "topright", plotTitle="Non-inflamed Mice Group
s")

generatePCOAPlots(micegroups_inflamed_genusCounts_samplesOnly_lognorm_df, micegroups_inflamed.metagroup,
                  micegroups_inflamed_colors, legendlocation = "topright", plotTitle="Inflamed Mice Groups")

dev.off()
```

#### Correlations and Scatterplots

The next few segments of code generates the following:

- Correlation coefficients for sample-pairs 1) within group and
  2. across groups
- Boxplots that summarize the correlation coefficients
- Scatterplots for each sample-pair comparison, which reflects relative abundance for each ASV between a given pair of samples

##### ASV Transfer Efficiencies Within FMT Groups

This first code chunk calculates the correlation of samples within the same group.

```
file_asv_updatedName_t_samplesOnly_lognorm
labels_inputRecipientGroups

# serial passage groups
labels_NIMM_lineage = c("HM1input", "HM1->WT", "NIMM-g1input", "NIMM-g1->WT", "NIMM-g2input", "NIMM-g2->WT")
labels_IMM_lineage = c("HM1input", "HM1->KO", "IMM-g1input", "IMM-g1->KO","IMM-g2input", "IMM-g2->KO")
labels_serialpassage_newNames = c("HM1input", "HM1->WT", "NIMM-g1input", "NIMM-g1->WT", "NIMM-g2input", "NIMM-g2-
>WT","HM1->KO", "IMM-g1input", "IMM-g1->KO","IMM-g2input", "IMM-g2->KO")
myASVandMetadata_newNames
names(myASVandMetadata_newNames)[names(myASVandMetadata_newNames)=="FMTGroupFMTsource..Recipientbackground"] = "F
MTGroupFMTsourcegtRecipientbackground"

## scatterplots
dir.create(here("boxplots","acrossGroups_pearson"),recursive = T)
```

```
## Warning in dir.create(here("boxplots", "acrossGroups_pearson"), recursive = T):
## 'C:\Users\aduong\Desktop\Balfour2022\boxplots\acrossGroups_pearson' already
## exists
```

```
dir.create(here("boxplots","withinGroups_pearson"),recursive = T)
```

```
## Warning in dir.create(here("boxplots", "withinGroups_pearson"), recursive = T):
## 'C:\Users\aduong\Desktop\Balfour2022\boxplots\withinGroups_pearson' already
## exists
```

```
dir.create(here("asvAbundance_scatterplots","acrossGroups_pearson"), recursive = T)
```

```
## Warning in dir.create(here("asvAbundance_scatterplots",
## "acrossGroups_pearson"), :
## 'C:\Users\aduong\Desktop\Balfour2022\asvAbundance_scatterplots\acrossGroups_pearson'
## already exists
```

```
  #within groups
withinGroup_pearson_newNames_serialpassage=getCorrelationCoefficient_withinGroup(groupList=labels_serialpassage_n
ewNames, tableWithMeta=myASVandMetadata_newNames, tableCountsOnly=file_asv_updatedName_t_samplesOnly_lognorm, cor
rTestMethod="pearson",sampleColumnName = "SampleID", variableName = "FMTGroupFMTsourcegtRecipientbackground")

withinGroup_pearson_newNames_NIMMlineage=getCorrelationCoefficient_withinGroup(groupList=labels_NIMM_lineage, tab
leWithMeta=myASVandMetadata_newNames, tableCountsOnly=file_asv_updatedName_t_samplesOnly_lognorm, corrTestMethod
="pearson",sampleColumnName = "SampleID", variableName = "FMTGroupFMTsourcegtRecipientbackground")

withinGroup_pearson_newNames_IMMlineage=getCorrelationCoefficient_withinGroup(groupList=labels_IMM_lineage, table
WithMeta=myASVandMetadata_newNames, tableCountsOnly=file_asv_updatedName_t_samplesOnly_lognorm, corrTestMethod="p
earson",sampleColumnName = "SampleID", variableName = "FMTGroupFMTsourcegtRecipientbackground")


## figure 4s
withinGroup_pearson_IMMlineage_genusCounts =getCorrelationCoefficient_withinGroup(groupList=labels_IMM_lineage, t
ableWithMeta=myASVandMetadata_newNames, tableCountsOnly=file_asv_updatedName_t_samplesOnly_lognorm, corrTestMetho
d="pearson",sampleColumnName = "SampleID", variableName = "FMTGroupFMTsourcegtRecipientbackground")


## create pdf of boxplots
pdf(here("boxplots","withinGroups_pearson","asv_boxplots_withinGroup_allSerialPassage.pdf"), height = 8, width =
10)
boxplot(withinGroup_pearson_newNames_serialpassage,names = labels_serialpassage_newNames, las=2, cex.axis=0.70,
        main="Correlation Coefficients Within Groups, All Serial Passage Groups", cex.main=0.8)
dev.off()

pdf(here("boxplots","withinGroups_pearson","asv_boxplots_withinGroup_SerialPassage_NIMM.pdf"), height = 8, width
= 10)
boxplot(withinGroup_pearson_newNames_NIMMlineage,names = labels_NIMM_lineage, las=2, cex.axis=0.70,
        main="Correlation Coefficients Within Groups, Serial Passage for NIMM Lineage", cex.main=0.8)
dev.off()

pdf(here("boxplots","withinGroups_pearson","asv_boxplots_withinGroup_SerialPassage_IMM.pdf"), height = 8, width =
10)
boxplot(withinGroup_pearson_newNames_IMMlineage,names = labels_IMM_lineage, las=2, cex.axis=0.70,
        main="Correlation Coefficients Within Groups, Serial Passage for IMM Lineage", cex.main=0.8)
dev.off()


#scatterplots, asv correlation for samples across groups
generateScatterplots_acrossGroup(groupList=labels_serialpassage_newNames, tableWithMeta=myASVandMetadata_newName
s,corrTestMethod = "pearson"  ,
                                  variableName = "FMTGroupFMTsourcegtRecipientbackground", sampleColumnName ="Samp
leID" ,tableCountsOnly=file_asv_updatedName_t_samplesOnly_lognorm, resultsDirectory = here("asvAbundance_scatterp
lots","acrossGroups_pearson","scatterplot_"))
```

##### ASV Transfer Efficiencies Within FMT Groups: Human Cohorts

Next, we generate correlations (within groups) for samples within only human cohorts (HM1, HM2), which are then visually summarized as boxplots.

```
labels_bothHumanCohorts = c("HM1->KO", "HM2->KO")

withinGroup_pearson_humanCohortsOnly = getCorrelationCoefficient_withinGroup(groupList=labels_bothHumanCohorts, t
ableWithMeta=myASVandMetadata_newNames, tableCountsOnly=file_asv_updatedName_t_samplesOnly_lognorm, corrTestMetho
d="pearson", variableName = "FMTGroupFMTsourcegtRecipientbackground", sampleColumnName = "SampleID")

pdf(here("boxplots","withinGroups_pearson", "ASV_boxplot_withinGroup_bothHumanCohortsOnly.pdf"))
boxplot(withinGroup_pearson_humanCohortsOnly,names = labels_bothHumanCohorts, las=2, cex.axis=0.70,
        main="Pearson Correlation Coefficients Within Groups, Human Cohorts 1 and 2", cex.main=0.8)
stripchart(withinGroup_pearson_humanCohortsOnly, add=TRUE, pch=16, col=c("green", "purple"), method="jitter", ver
tical = T)

dev.off()

myList =c("HM1->WT", "NIMM-g1->WT", "NIMM-g2->WT","HM1->KO", "IMM-g1->KO", "IMM-g2->KO")

withinGroup_pearson_theSixGroups = getCorrelationCoefficient_withinGroup(groupList=myList, tableWithMeta=myASVand
Metadata_newNames, tableCountsOnly=file_asv_updatedName_t_samplesOnly_lognorm, corrTestMethod="pearson", variable
Name = "FMTGroupFMTsourcegtRecipientbackground", sampleColumnName = "SampleID")

pdf(here("boxplots","withinGroups_pearson", "ASV_boxplot_withinGroup_WTandKOgroups.pdf"))

boxplot(withinGroup_pearson_theSixGroups,names = myList, las=2, cex.axis=0.70,
        main="Pearson Correlation Coefficients Within Groups, WT vs KO", cex.main=0.8)
stripchart(withinGroup_pearson_theSixGroups, add=TRUE, pch=16, col="blue", method="jitter", vertical = T)

dev.off()
```

##### ASV Transfer Efficiencies Across FMT Groups

The code below generates ASV scatterplots and boxplots for samples across groups. This first code chunk sets the FMT groups of interest for the following analysis.

```
file_asv_updatedName_t_samplesOnly_lognorm
labels_inputRecipientGroups

labels_NIMM_lineage = c("HM1input", "HM1->WT", "NIMM-g1input", "NIMM-g1->WT", "NIMM-g2input", "NIMM-g2->WT")
labels_IMM_lineage = c("HM1input", "HM1->KO", "IMM-g1input", "IMM-g1->KO","IMM-g2input", "IMM-g2->KO")
labels_serialpassage_newNames = c("HM1input", "HM1->WT", "NIMM-g1input", "NIMM-g1->WT", "NIMM-g2input", "NIMM-g2-
>WT","HM1->KO", "IMM-g1input", "IMM-g1->KO","IMM-g2input", "IMM-g2->KO")
myASVandMetadata_newNames

#indexing groups of interest for asv correlation boxplots
fig4a.wtlineage.index = c(1,20,35)
fig4a.wtlineage.supp.index = c(2,4,21)

fig4b.kolineage.index = c(6,50,55)
fig4b.kolineage.supp.index = c(7,9,51)

fig4c.index = c(12,29,47,54)
fig4c.supp.index = c(32,43,34,45)
```

Next, we generate the correlations between samples from across groups at three taxonomic levels: strain level (ASV), genus, and phylum.

```
#across groups
  ## asv level counts
corr_acrossGroups_pearson_asvCounts_withHuman = getCorrelationCoefficient_acrossGroup(groupList=labels_serialpass
age_newNames, tableWithMeta=myASVandMetadata_newNames, tableCountsOnly=file_asv_updatedName_t_samplesOnly_lognor
m, corrTestMethod="pearson",variableName = "FMTGroupFMTsourcegtRecipientbackground", sampleColumnName = "SampleI
D")

  ## genus level counts
corr_acrossGroups_pearson_GenusCounts_withHuman = getCorrelationCoefficient_acrossGroup(groupList=labels_serialpa
ssage_newNames, tableWithMeta=genusCountsandMetadata, tableCountsOnly=genusCounts_samplesOnly_df_lognorm,
corrTestMethod="pearson", variableName = "FMTGroupFMTsourcegtRecipientbackground", sampleColumnName = "SampleID")

    ## phylum level counts
corr_acrossGroups_pearson_PhylumCounts_withHuman = getCorrelationCoefficient_acrossGroup(groupList=labels_serialp
assage_newNames, tableWithMeta=phylumCountsandMetadata, tableCountsOnly=phylumCounts_samplesOnly_df_lognorm,
corrTestMethod="pearson", variableName = "FMTGroupFMTsourcegtRecipientbackground", sampleColumnName = "SampleID")
```

These boxplots are for inflamed (KO) groups in our SV transfer comparison across groups.

```
  #asv level counts
pdf(here("boxplots","acrossGroups_pearson", "fig4_inflamed_asv.pdf"))
boxplot(corr_acrossGroups_pearson_asvCounts_withHuman[[2]][fig4b.kolineage.index],names = corr_acrossGroups_pears
on_asvCounts_withHuman[[1]][fig4b.kolineage.index],  cex.axis=0.6,
        main="Correlation Coefficients Across Groups, Inflamed Groups", cex.main=0.8,
        ylim=c(0,1))
dev.off()

pdf(here("boxplots","acrossGroups_pearson","fig4_inflamed_asv_supp.pdf"))
boxplot(corr_acrossGroups_pearson_asvCounts_withHuman[[2]][fig4b.kolineage.supp.index],names = corr_acrossGroups_
pearson_asvCounts_withHuman[[1]][fig4b.kolineage.supp.index],  cex.axis=0.6,
        main="Correlation Coefficients Across Groups, Inflamed Groups (SUPPFIG)", cex.main=0.8,
        ylim=c(0,1))
dev.off()

  # genus level counts

pdf(here("boxplots","acrossGroups_pearson","fig4_inflamed_genus.pdf"))
boxplot(corr_acrossGroups_pearson_GenusCounts_withHuman[[2]][fig4b.kolineage.index],names = corr_acrossGroups_pea
rson_GenusCounts_withHuman[[1]][fig4b.kolineage.index],  cex.axis=0.6,
        main="Correlation Coefficients Across Groups, Inflamed Groups (Genus)", cex.main=0.8,
        ylim=c(0,1))
dev.off()

pdf(here("boxplots","acrossGroups_pearson","fig4_inflamed_genus_supp.pdf"))
boxplot(corr_acrossGroups_pearson_GenusCounts_withHuman[[2]][fig4b.kolineage.supp.index],names = corr_acrossGroup
s_pearson_GenusCounts_withHuman[[1]][fig4b.kolineage.supp.index],  cex.axis=0.6,
        main="Correlation Coefficients Across Groups, Inflamed Groups (Genus) (SUPPFIG)", cex.main=0.8,
        ylim=c(0,1))
dev.off()

    #phylum counts

pdf(here("boxplots","acrossGroups_pearson", "fig4_inflamed_phylum.pdf"))
boxplot(corr_acrossGroups_pearson_PhylumCounts_withHuman[[2]][fig4b.kolineage.index],names = corr_acrossGroups_pe
arson_PhylumCounts_withHuman[[1]][fig4b.kolineage.index],  cex.axis=0.6,
        main="Correlation Coefficients Across Groups, Inflamed Groups (Phylum)", cex.main=0.8,
        ylim=c(0,1))
dev.off()

pdf(here("boxplots","acrossGroups_pearson","fig4_inflamed_phylum_supp.pdf"))
boxplot(corr_acrossGroups_pearson_PhylumCounts_withHuman[[2]][fig4b.kolineage.supp.index],names = corr_acrossGrou
ps_pearson_PhylumCounts_withHuman[[1]][fig4b.kolineage.supp.index],  cex.axis=0.6,
        main="Correlation Coefficients Across Groups, Inflamed Groups (Phylum, SUPPFIG)", cex.main=0.8,
        ylim=c(0,1))
dev.off()
```

These boxplots are for non-inflamed (WT) groups in our SV transfer comparison across groups.

```
#asv counts

pdf(here("boxplots","acrossGroups_pearson","fig4_NONinflamed_asv.pdf"))
boxplot(corr_acrossGroups_pearson_asvCounts_withHuman[[2]][fig4a.wtlineage.index],names = corr_acrossGroups_pears
on_asvCounts_withHuman[[1]][fig4a.wtlineage.index],  cex.axis=0.6,
        main="Correlation Coefficients Across Groups, Non-inflamed Groups", cex.main=0.8,
        ylim=c(0,1))
dev.off()

pdf(here("boxplots","acrossGroups_pearson","fig4_NONinflamed_asv_supp.pdf"))
boxplot(corr_acrossGroups_pearson_asvCounts_withHuman[[2]][fig4a.wtlineage.supp.index],names = corr_acrossGroups_
pearson_asvCounts_withHuman[[1]][fig4a.wtlineage.supp.index],  cex.axis=0.6,
        main="Correlation Coefficients Across Groups, Non-inflamed Groups (SUPPFIG)", cex.main=0.8,
        ylim=c(0,1))
dev.off()

    #genus counts

pdf(here("boxplots","acrossGroups_pearson", "fig4_NONinflamed_genus.pdf"))
boxplot(corr_acrossGroups_pearson_GenusCounts_withHuman[[2]][fig4a.wtlineage.index],names = corr_acrossGroups_pea
rson_GenusCounts_withHuman[[1]][fig4a.wtlineage.index],  cex.axis=0.6,
        main="Correlation Coefficients Across Groups, Non-inflamed Groups (Genus)", cex.main=0.8,
        ylim=c(0,1))
dev.off()

pdf(here("boxplots","acrossGroups_pearson","fig4_NONinflamed_genus_supp.pdf"))
boxplot(corr_acrossGroups_pearson_GenusCounts_withHuman[[2]][fig4a.wtlineage.supp.index],names = corr_acrossGroup
s_pearson_GenusCounts_withHuman[[1]][fig4a.wtlineage.supp.index],  cex.axis=0.6,
        main="Correlation Coefficients Across Groups, Non-inflamed Groups (Genus,SUPPFIG)", cex.main=0.8,
        ylim=c(0,1))
dev.off()

  # phylum counts

pdf(here("boxplots","acrossGroups_pearson","fig4_NONinflamed_phylum.pdf"))
boxplot(corr_acrossGroups_pearson_PhylumCounts_withHuman[[2]][fig4a.wtlineage.index],names = corr_acrossGroups_pe
arson_PhylumCounts_withHuman[[1]][fig4a.wtlineage.index],  cex.axis=0.6,
        main="Correlation Coefficients Across Groups, Non-inflamed Groups (Phylum)", cex.main=0.8,
        ylim=c(0,1))
dev.off()

pdf(here("boxplots","acrossGroups_pearson","fig4_NONinflamed_phylum_supp.pdf"))
boxplot(corr_acrossGroups_pearson_PhylumCounts_withHuman[[2]][fig4a.wtlineage.supp.index],names = corr_acrossGrou
ps_pearson_PhylumCounts_withHuman[[1]][fig4a.wtlineage.supp.index],  cex.axis=0.6,
        main="Correlation Coefficients Across Groups, Non-inflamed Groups (Phylum,SUPPFIG)", cex.main=0.8,
        ylim=c(0,1))
dev.off()
```

These boxplots include the cross-over groups in our SV transfer comparison across groups.

```
#asv counts
pdf(here("boxplots","acrossGroups_pearson","fig4_crossover_asv.pdf"), width = 10)
boxplot(corr_acrossGroups_pearson_asvCounts_withHuman[[2]][fig4c.index],names = corr_acrossGroups_pearson_asvCoun
ts_withHuman[[1]][fig4c.index],  cex.axis=0.6,
        main="Correlation Coefficients Across Groups, Cross-over Group", cex.main=0.8,
        ylim=c(0,1))
dev.off()

pdf(here("boxplots","acrossGroups_pearson","fig4_crossover_supp_asv.pdf"))
boxplot(corr_acrossGroups_pearson_asvCounts_withHuman[[2]][fig4c.supp.index],names = corr_acrossGroups_pearson_as
vCounts_withHuman[[1]][fig4c.supp.index],  cex.axis=0.6,
        main="Correlation Coefficients Across Groups, Cross-over Group (SUPPFIG)", cex.main=0.8,
        ylim=c(0,1))
dev.off()

    #genus counts
pdf(here("boxplots","acrossGroups_pearson","fig4_crossover_genus.pdf"))
boxplot(corr_acrossGroups_pearson_GenusCounts_withHuman[[2]][fig4c.index],
        names = corr_acrossGroups_pearson_GenusCounts_withHuman[[1]][fig4c.index],  cex.axis=0.6,
        main="Correlation Coefficients Across Groups, Cross-over Group (Genus)", cex.main=0.8,
        ylim=c(0,1))
dev.off()

pdf(here("boxplots","acrossGroups_pearson","fig4_crossover_supp_genus.pdf"))
boxplot(corr_acrossGroups_pearson_GenusCounts_withHuman[[2]][fig4c.supp.index],names = corr_acrossGroups_pearson_
GenusCounts_withHuman[[1]][fig4c.supp.index],  cex.axis=0.6,
        main="Correlation Coefficients Across Groups, Cross-over Group (Genus, SUPPFIG)", cex.main=0.8,
        ylim=c(0,1))
dev.off()

    #phylum counts
pdf(here("boxplots","acrossGroups_pearson","fig4_crossover_phylum.pdf"))
boxplot(corr_acrossGroups_pearson_PhylumCounts_withHuman[[2]][fig4c.index],
        names = corr_acrossGroups_pearson_PhylumCounts_withHuman[[1]][fig4c.index],  cex.axis=0.6,
        main="Correlation Coefficients Across Groups, Cross-over Group (Phylum)", cex.main=0.8,
        ylim=c(0,1))
dev.off()

pdf(here("boxplots","acrossGroups_pearson","fig4_crossover_supp_phylum.pdf"))
boxplot(corr_acrossGroups_pearson_PhylumCounts_withHuman[[2]][fig4c.supp.index],names = corr_acrossGroups_pearson
_PhylumCounts_withHuman[[1]][fig4c.supp.index],  cex.axis=0.6,
        main="Correlation Coefficients Across Groups, Cross-over Group (Phylum, SUPPFIG)", cex.main=0.8,
        ylim=c(0,1))
dev.off()
```

##### ASV Transfer Efficiencies Across FMT Groups: Non-transferred ASVs

Here, we plot the abundance of the non-transferred ASVs as histograms.

```
dir.create(here("histograms_nontransferredASV","tablesWithTaxonomy"),recursive = T)
```

```
## Warning in dir.create(here("histograms_nontransferredASV",
## "tablesWithTaxonomy"), :
## 'C:\Users\aduong\Desktop\Balfour2022\histograms_nontransferredASV\tablesWithTaxonomy'
## already exists
```

```
labels_NIMM_lineage = c("HM1input", "HM1->WT", "NIMM-g1input", "NIMM-g1->WT", "NIMM-g2input", "NIMM-g2->WT")
labels_IMM_lineage = c("HM1input", "HM1->KO", "IMM-g1input", "IMM-g1->KO","IMM-g2input", "IMM-g2->KO")


generateScatterPlots_samplePairsAcrossGroup_histogramOfNontransferringASVS(groupList=labels_serialpassage_newName
s,
                                                                            tableWithMeta=myASVandMetadata_newNames,
                                                                               tableCountsOnly=file_asv_updatedName_t_s
amplesOnly_lognorm, resultsDirectory = here("histograms_nontransferredASV", "histogram_"))
```

```
#tables of non-transferred asvs with taxa name in each

generateTableWithTaxonomy_NontransferringASVS_samplePairsAcrossGroup(groupList=labels_serialpassage_newNames,
                                                                      taxonomyMap=asv_taxonomy_map,
                                                                      taxaColName="taxonomy",
                                                                      asvColName="OTUID",
                                                                      tableWithMeta=myASVandMetadata_newNames,
                                                                      tableCountsOnly=file_asv_updatedName_t_sampl
esOnly_lognorm,
                                                                      resultsDirectory=here("histograms_nontransfe
rredASV","tablesWithTaxonomy", "table_"))
```

##### ASV Transfer Efficiencies Within FMT Groups at the Phylum level

The code below parses the taxonomy column to count the number of different phylum present in this counts table. The phylum "NA" means unclassified.

There are a total of 13 phyla in the ASV counts table. Of these 13 phyla, only four had one ASV present. For statistical testing purposes, these phyla were excluded from the Pearson Correlation Coefficient test: Arthropoda, Cyanobacteria, Desulfobacterota, and Vertebrata. Campilobacterota and Patescibacteria were very sparse (present in 5 or less samples each), and since this led to empty results in the boxplot, we excluded these two phyla as well.

```
taxanames_all = file_asv_updatedName$taxonomy

taxanames_all_split = strsplit(taxanames_all, ";")

phylum_label = NULL

for(i in 1:length(taxanames_all_split)){
  phylum_label[i] = taxanames_all_split[[i]][2]
}

phylum_label_short = gsub("p__", "", phylum_label)
phylum_label_short = gsub(" ", "", phylum_label_short)

mapping_ASVandPhyla = data.frame("ASV" = file_asv_updatedName$OTUID, "PhylumLabels" = phylum_label_short)
table(mapping_ASVandPhyla$PhylumLabels)

mapping_ASVandPhyla$PhylumLabels[is.na(mapping_ASVandPhyla$PhylumLabels)] = "Unclassified"

#exclude the single ASV phylum
phlyum_groups = unique(mapping_ASVandPhyla$PhylumLabels)
singles = c("Arthropoda","Cyanobacteria","Desulfobacterota", "Vertebrata")
phlyum_groups_subset = phlyum_groups[! phlyum_groups %in% singles]

#rename the NA group in phylum list
phlyum_groups_subset_2 = phlyum_groups_subset
phlyum_groups_subset_2[7] = "Unclassified"

phlyum_groups_subset_withoutSparsePhyla = phlyum_groups_subset_2[phlyum_groups_subset_2 != "Patescibacteria" & ph
lyum_groups_subset_2 != "Campilobacterota"]

phlyum_groups_subset_withoutSparsePhyla[7] = "Unclassified"

myGroup_serialPassage_miceOnly = c("IMM-g1->KO", "IMM-g2->KO", "NIMM-g1->WT", "NIMM-g2->WT")
myGroup_serialPassage_miceandHuman = c("IMM-g1->KO", "IMM-g2->KO", "NIMM-g1->WT", "NIMM-g2->WT", "HM1->KO", "HM1-
>WT")
sampleID_for_myGroup_serialPassage_miceOnly = metamapping_updatedName$SampleID[metamapping_updatedName$FMTGroupFM
Tsource..Recipientbackground %in% myGroup_serialPassage_miceOnly[1]]

mygroup_outfile_labels = gsub("->", "_to_", myGroup_serialPassage_miceandHuman)

## for selected FMT groups, generate correlations (samples within the same FMT group, stratified at the phylum ta
xa level) and boxplots of those correlations
for(i in 1:length(myGroup_serialPassage_miceandHuman)){

  myCorr_withinGroup_pearson_phlyum_perIRGroup_OUTPUT=getCorrelationCoefficient_withinGroup_phylumLevel_byGroup(g
roupList=phlyum_groups_subset_withoutSparsePhyla, tableWithMeta=myASVandMetadata_newNames, tableCountsOnly=file_a
sv_updatedName_t_samplesOnly_lognorm, corrTestMethod="pearson", sampleGroup = myGroup_serialPassage_miceandHuman[
i], mappingMetaFile = mapping_ASVandPhyla,mappingMetaFile_taxaColName = "ASV", mappingMetaFile_phylumColName = "P
hylumLabels", tableWithMeta_sampleColName = "SampleID" ,tableWithMeta_groupColName = "FMTGroupFMTsourcegtRecipien
tbackground")

  pdf(paste0(here("boxplots","withinGroups_pearson", "boxplot_phylum_"),mygroup_outfile_labels[i] , ".pdf"), widt
h = 10)
  boxplot(myCorr_withinGroup_pearson_phlyum_perIRGroup_OUTPUT[[2]], names = phlyum_groups_subset_withoutSparsePhy
la, main=paste0("Pearson Correlation Coefficient (R), Phylum Level ", myGroup_serialPassage_miceandHuman[i]), ce
x.axis=0.65, yaxt="n")
  axis(side=2, at=seq(0, 1, by=0.1))
  dev.off()
}
```

```
## Warning in cor(x, y): the standard deviation is zero

## Warning in cor(x, y): the standard deviation is zero

## Warning in cor(x, y): the standard deviation is zero

## Warning in cor(x, y): the standard deviation is zero

## Warning in cor(x, y): the standard deviation is zero

## Warning in cor(x, y): the standard deviation is zero

## Warning in cor(x, y): the standard deviation is zero

## Warning in cor(x, y): the standard deviation is zero

## Warning in cor(x, y): the standard deviation is zero

## Warning in cor(x, y): the standard deviation is zero

## Warning in cor(x, y): the standard deviation is zero

## Warning in cor(x, y): the standard deviation is zero

## Warning in cor(x, y): the standard deviation is zero

## Warning in cor(x, y): the standard deviation is zero

## Warning in cor(x, y): the standard deviation is zero

## Warning in cor(x, y): the standard deviation is zero

## Warning in cor(x, y): the standard deviation is zero

## Warning in cor(x, y): the standard deviation is zero
```

```
## Warning in cor(x, y): the standard deviation is zero
## Warning in cor(x, y): the standard deviation is zero
## Warning in cor(x, y): the standard deviation is zero
## Warning in cor(x, y): the standard deviation is zero
## Warning in cor(x, y): the standard deviation is zero
## Warning in cor(x, y): the standard deviation is zero
## Warning in cor(x, y): the standard deviation is zero
## Warning in cor(x, y): the standard deviation is zero
## Warning in cor(x, y): the standard deviation is zero
## Warning in cor(x, y): the standard deviation is zero
## Warning in cor(x, y): the standard deviation is zero
## Warning in cor(x, y): the standard deviation is zero
## Warning in cor(x, y): the standard deviation is zero
## Warning in cor(x, y): the standard deviation is zero
## Warning in cor(x, y): the standard deviation is zero
## Warning in cor(x, y): the standard deviation is zero
## Warning in cor(x, y): the standard deviation is zero
## Warning in cor(x, y): the standard deviation is zero
## Warning in cor(x, y): the standard deviation is zero
## Warning in cor(x, y): the standard deviation is zero
## Warning in cor(x, y): the standard deviation is zero
## Warning in cor(x, y): the standard deviation is zero
## Warning in cor(x, y): the standard deviation is zero
## Warning in cor(x, y): the standard deviation is zero
## Warning in cor(x, y): the standard deviation is zero
## Warning in cor(x, y): the standard deviation is zero
## Warning in cor(x, y): the standard deviation is zero
## Warning in cor(x, y): the standard deviation is zero
## Warning in cor(x, y): the standard deviation is zero
## Warning in cor(x, y): the standard deviation is zero
## Warning in cor(x, y): the standard deviation is zero
## Warning in cor(x, y): the standard deviation is zero
## Warning in cor(x, y): the standard deviation is zero
## Warning in cor(x, y): the standard deviation is zero
## Warning in cor(x, y): the standard deviation is zero
## Warning in cor(x, y): the standard deviation is zero
## Warning in cor(x, y): the standard deviation is zero
## Warning in cor(x, y): the standard deviation is zero
## Warning in cor(x, y): the standard deviation is zero
## Warning in cor(x, y): the standard deviation is zero
## Warning in cor(x, y): the standard deviation is zero
## Warning in cor(x, y): the standard deviation is zero
## Warning in cor(x, y): the standard deviation is zero
## Warning in cor(x, y): the standard deviation is zero
## Warning in cor(x, y): the standard deviation is zero
## Warning in cor(x, y): the standard deviation is zero
## Warning in cor(x, y): the standard deviation is zero
## Warning in cor(x, y): the standard deviation is zero
## Warning in cor(x, y): the standard deviation is zero
## Warning in cor(x, y): the standard deviation is zero
## Warning in cor(x, y): the standard deviation is zero
## Warning in cor(x, y): the standard deviation is zero
## Warning in cor(x, y): the standard deviation is zero
## Warning in cor(x, y): the standard deviation is zero
## Warning in cor(x, y): the standard deviation is zero
## Warning in cor(x, y): the standard deviation is zero
## Warning in cor(x, y): the standard deviation is zero
## Warning in cor(x, y): the standard deviation is zero
## Warning in cor(x, y): the standard deviation is zero
## Warning in cor(x, y): the standard deviation is zero
## Warning in cor(x, y): the standard deviation is zero
## Warning in cor(x, y): the standard deviation is zero
## Warning in cor(x, y): the standard deviation is zero
## Warning in cor(x, y): the standard deviation is zero
## Warning in cor(x, y): the standard deviation is zero
## Warning in cor(x, y): the standard deviation is zero
## Warning in cor(x, y): the standard deviation is zero
## Warning in cor(x, y): the standard deviation is zero
## Warning in cor(x, y): the standard deviation is zero
## Warning in cor(x, y): the standard deviation is zero
## Warning in cor(x, y): the standard deviation is zero
## Warning in cor(x, y): the standard deviation is zero
## Warning in cor(x, y): the standard deviation is zero
## Warning in cor(x, y): the standard deviation is zero
## Warning in cor(x, y): the standard deviation is zero
## Warning in cor(x, y): the standard deviation is zero
## Warning in cor(x, y): the standard deviation is zero
## Warning in cor(x, y): the standard deviation is zero
## Warning in cor(x, y): the standard deviation is zero
## Warning in cor(x, y): the standard deviation is zero
## Warning in cor(x, y): the standard deviation is zero
## Warning in cor(x, y): the standard deviation is zero
## Warning in cor(x, y): the standard deviation is zero
## Warning in cor(x, y): the standard deviation is zero
## Warning in cor(x, y): the standard deviation is zero
## Warning in cor(x, y): the standard deviation is zero
## Warning in cor(x, y): the standard deviation is zero
## Warning in cor(x, y): the standard deviation is zero
## Warning in cor(x, y): the standard deviation is zero
## Warning in cor(x, y): the standard deviation is zero
## Warning in cor(x, y): the standard deviation is zero
## Warning in cor(x, y): the standard deviation is zero
## Warning in cor(x, y): the standard deviation is zero
## Warning in cor(x, y): the standard deviation is zero
## Warning in cor(x, y): the standard deviation is zero
## Warning in cor(x, y): the standard deviation is zero
## Warning in cor(x, y): the standard deviation is zero
## Warning in cor(x, y): the standard deviation is zero
## Warning in cor(x, y): the standard deviation is zero
## Warning in cor(x, y): the standard deviation is zero
## Warning in cor(x, y): the standard deviation is zero
## Warning in cor(x, y): the standard deviation is zero
## Warning in cor(x, y): the standard deviation is zero
## Warning in cor(x, y): the standard deviation is zero
## Warning in cor(x, y): the standard deviation is zero
## Warning in cor(x, y): the standard deviation is zero
## Warning in cor(x, y): the standard deviation is zero
## Warning in cor(x, y): the standard deviation is zero
## Warning in cor(x, y): the standard deviation is zero
## Warning in cor(x, y): the standard deviation is zero
## Warning in cor(x, y): the standard deviation is zero
## Warning in cor(x, y): the standard deviation is zero
## Warning in cor(x, y): the standard deviation is zero
## Warning in cor(x, y): the standard deviation is zero
## Warning in cor(x, y): the standard deviation is zero
## Warning in cor(x, y): the standard deviation is zero
## Warning in cor(x, y): the standard deviation is zero
## Warning in cor(x, y): the standard deviation is zero
## Warning in cor(x, y): the standard deviation is zero
## Warning in cor(x, y): the standard deviation is zero
## Warning in cor(x, y): the standard deviation is zero
## Warning in cor(x, y): the standard deviation is zero
## Warning in cor(x, y): the standard deviation is zero
## Warning in cor(x, y): the standard deviation is zero
## Warning in cor(x, y): the standard deviation is zero
## Warning in cor(x, y): the standard deviation is zero
## Warning in cor(x, y): the standard deviation is zero
## Warning in cor(x, y): the standard deviation is zero
## Warning in cor(x, y): the standard deviation is zero
## Warning in cor(x, y): the standard deviation is zero
## Warning in cor(x, y): the standard deviation is zero
## Warning in cor(x, y): the standard deviation is zero
## Warning in cor(x, y): the standard deviation is zero
## Warning in cor(x, y): the standard deviation is zero
## Warning in cor(x, y): the standard deviation is zero
## Warning in cor(x, y): the standard deviation is zero
## Warning in cor(x, y): the standard deviation is zero
## Warning in cor(x, y): the standard deviation is zero
## Warning in cor(x, y): the standard deviation is zero
## Warning in cor(x, y): the standard deviation is zero
## Warning in cor(x, y): the standard deviation is zero
## Warning in cor(x, y): the standard deviation is zero
## Warning in cor(x, y): the standard deviation is zero
## Warning in cor(x, y): the standard deviation is zero
## Warning in cor(x, y): the standard deviation is zero
## Warning in cor(x, y): the standard deviation is zero
## Warning in cor(x, y): the standard deviation is zero
## Warning in cor(x, y): the standard deviation is zero
## Warning in cor(x, y): the standard deviation is zero
## Warning in cor(x, y): the standard deviation is zero
## Warning in cor(x, y): the standard deviation is zero
## Warning in cor(x, y): the standard deviation is zero
## Warning in cor(x, y): the standard deviation is zero
## Warning in cor(x, y): the standard deviation is zero
## Warning in cor(x, y): the standard deviation is zero
## Warning in cor(x, y): the standard deviation is zero
## Warning in cor(x, y): the standard deviation is zero
## Warning in cor(x, y): the standard deviation is zero
## Warning in cor(x, y): the standard deviation is zero
## Warning in cor(x, y): the standard deviation is zero
## Warning in cor(x, y): the standard deviation is zero
## Warning in cor(x, y): the standard deviation is zero
## Warning in cor(x, y): the standard deviation is zero
## Warning in cor(x, y): the standard deviation is zero
## Warning in cor(x, y): the standard deviation is zero
## Warning in cor(x, y): the standard deviation is zero
## Warning in cor(x, y): the standard deviation is zero
## Warning in cor(x, y): the standard deviation is zero
## Warning in cor(x, y): the standard deviation is zero
## Warning in cor(x, y): the standard deviation is zero
## Warning in cor(x, y): the standard deviation is zero
## Warning in cor(x, y): the standard deviation is zero
## Warning in cor(x, y): the standard deviation is zero
## Warning in cor(x, y): the standard deviation is zero
## Warning in cor(x, y): the standard deviation is zero
## Warning in cor(x, y): the standard deviation is zero
## Warning in cor(x, y): the standard deviation is zero
## Warning in cor(x, y): the standard deviation is zero
## Warning in cor(x, y): the standard deviation is zero
## Warning in cor(x, y): the standard deviation is zero
## Warning in cor(x, y): the standard deviation is zero
## Warning in cor(x, y): the standard deviation is zero
## Warning in cor(x, y): the standard deviation is zero
## Warning in cor(x, y): the standard deviation is zero
## Warning in cor(x, y): the standard deviation is zero
## Warning in cor(x, y): the standard deviation is zero
## Warning in cor(x, y): the standard deviation is zero
## Warning in cor(x, y): the standard deviation is zero
## Warning in cor(x, y): the standard deviation is zero
## Warning in cor(x, y): the standard deviation is zero
## Warning in cor(x, y): the standard deviation is zero
## Warning in cor(x, y): the standard deviation is zero
## Warning in cor(x, y): the standard deviation is zero
## Warning in cor(x, y): the standard deviation is zero
## Warning in cor(x, y): the standard deviation is zero
## Warning in cor(x, y): the standard deviation is zero
## Warning in cor(x, y): the standard deviation is zero
## Warning in cor(x, y): the standard deviation is zero
## Warning in cor(x, y): the standard deviation is zero
## Warning in cor(x, y): the standard deviation is zero
## Warning in cor(x, y): the standard deviation is zero
## Warning in cor(x, y): the standard deviation is zero
## Warning in cor(x, y): the standard deviation is zero
## Warning in cor(x, y): the standard deviation is zero
## Warning in cor(x, y): the standard deviation is zero
## Warning in cor(x, y): the standard deviation is zero
## Warning in cor(x, y): the standard deviation is zero
## Warning in cor(x, y): the standard deviation is zero
## Warning in cor(x, y): the standard deviation is zero
## Warning in cor(x, y): the standard deviation is zero
## Warning in cor(x, y): the standard deviation is zero
## Warning in cor(x, y): the standard deviation is zero
## Warning in cor(x, y): the standard deviation is zero
## Warning in cor(x, y): the standard deviation is zero
## Warning in cor(x, y): the standard deviation is zero
## Warning in cor(x, y): the standard deviation is zero
## Warning in cor(x, y): the standard deviation is zero
## Warning in cor(x, y): the standard deviation is zero
## Warning in cor(x, y): the standard deviation is zero
## Warning in cor(x, y): the standard deviation is zero
## Warning in cor(x, y): the standard deviation is zero
## Warning in cor(x, y): the standard deviation is zero
## Warning in cor(x, y): the standard deviation is zero
## Warning in cor(x, y): the standard deviation is zero
## Warning in cor(x, y): the standard deviation is zero
## Warning in cor(x, y): the standard deviation is zero
## Warning in cor(x, y): the standard deviation is zero
## Warning in cor(x, y): the standard deviation is zero
## Warning in cor(x, y): the standard deviation is zero
## Warning in cor(x, y): the standard deviation is zero
## Warning in cor(x, y): the standard deviation is zero
## Warning in cor(x, y): the standard deviation is zero
## Warning in cor(x, y): the standard deviation is zero
## Warning in cor(x, y): the standard deviation is zero
## Warning in cor(x, y): the standard deviation is zero
## Warning in cor(x, y): the standard deviation is zero
## Warning in cor(x, y): the standard deviation is zero
## Warning in cor(x, y): the standard deviation is zero
## Warning in cor(x, y): the standard deviation is zero
## Warning in cor(x, y): the standard deviation is zero
## Warning in cor(x, y): the standard deviation is zero
## Warning in cor(x, y): the standard deviation is zero
## Warning in cor(x, y): the standard deviation is zero
## Warning in cor(x, y): the standard deviation is zero
## Warning in cor(x, y): the standard deviation is zero
## Warning in cor(x, y): the standard deviation is zero
## Warning in cor(x, y): the standard deviation is zero
## Warning in cor(x, y): the standard deviation is zero
## Warning in cor(x, y): the standard deviation is zero
## Warning in cor(x, y): the standard deviation is zero
## Warning in cor(x, y): the standard deviation is zero
## Warning in cor(x, y): the standard deviation is zero
## Warning in cor(x, y): the standard deviation is zero
## Warning in cor(x, y): the standard deviation is zero
## Warning in cor(x, y): the standard deviation is zero
## Warning in cor(x, y): the standard deviation is zero
## Warning in cor(x, y): the standard deviation is zero
## Warning in cor(x, y): the standard deviation is zero
## Warning in cor(x, y): the standard deviation is zero
## Warning in cor(x, y): the standard deviation is zero
## Warning in cor(x, y): the standard deviation is zero
## Warning in cor(x, y): the standard deviation is zero
## Warning in cor(x, y): the standard deviation is zero
## Warning in cor(x, y): the standard deviation is zero
## Warning in cor(x, y): the standard deviation is zero
## Warning in cor(x, y): the standard deviation is zero
## Warning in cor(x, y): the standard deviation is zero
## Warning in cor(x, y): the standard deviation is zero
## Warning in cor(x, y): the standard deviation is zero
## Warning in cor(x, y): the standard deviation is zero
## Warning in cor(x, y): the standard deviation is zero
## Warning in cor(x, y): the standard deviation is zero
## Warning in cor(x, y): the standard deviation is zero
## Warning in cor(x, y): the standard deviation is zero
## Warning in cor(x, y): the standard deviation is zero
## Warning in cor(x, y): the standard deviation is zero
## Warning in cor(x, y): the standard deviation is zero
## Warning in cor(x, y): the standard deviation is zero
## Warning in cor(x, y): the standard deviation is zero
## Warning in cor(x, y): the standard deviation is zero
## Warning in cor(x, y): the standard deviation is zero
## Warning in cor(x, y): the standard deviation is zero
## Warning in cor(x, y): the standard deviation is zero
## Warning in cor(x, y): the standard deviation is zero
## Warning in cor(x, y): the standard deviation is zero
## Warning in cor(x, y): the standard deviation is zero
## Warning in cor(x, y): the standard deviation is zero
## Warning in cor(x, y): the standard deviation is zero
## Warning in cor(x, y): the standard deviation is zero
## Warning in cor(x, y): the standard deviation is zero
## Warning in cor(x, y): the standard deviation is zero
## Warning in cor(x, y): the standard deviation is zero
## Warning in cor(x, y): the standard deviation is zero
## Warning in cor(x, y): the standard deviation is zero
## Warning in cor(x, y): the standard deviation is zero
## Warning in cor(x, y): the standard deviation is zero
## Warning in cor(x, y): the standard deviation is zero
## Warning in cor(x, y): the standard deviation is zero
## Warning in cor(x, y): the standard deviation is zero
## Warning in cor(x, y): the standard deviation is zero
## Warning in cor(x, y): the standard deviation is zero
## Warning in cor(x, y): the standard deviation is zero
## Warning in cor(x, y): the standard deviation is zero
## Warning in cor(x, y): the standard deviation is zero
## Warning in cor(x, y): the standard deviation is zero
## Warning in cor(x, y): the standard deviation is zero
## Warning in cor(x, y): the standard deviation is zero
## Warning in cor(x, y): the standard deviation is zero
## Warning in cor(x, y): the standard deviation is zero
## Warning in cor(x, y): the standard deviation is zero
## Warning in cor(x, y): the standard deviation is zero
## Warning in cor(x, y): the standard deviation is zero
## Warning in cor(x, y): the standard deviation is zero
## Warning in cor(x, y): the standard deviation is zero
## Warning in cor(x, y): the standard deviation is zero
## Warning in cor(x, y): the standard deviation is zero
## Warning in cor(x, y): the standard deviation is zero
## Warning in cor(x, y): the standard deviation is zero
## Warning in cor(x, y): the standard deviation is zero
## Warning in cor(x, y): the standard deviation is zero
## Warning in cor(x, y): the standard deviation is zero
## Warning in cor(x, y): the standard deviation is zero
## Warning in cor(x, y): the standard deviation is zero
## Warning in cor(x, y): the standard deviation is zero
## Warning in cor(x, y): the standard deviation is zero
## Warning in cor(x, y): the standard deviation is zero
## Warning in cor(x, y): the standard deviation is zero
## Warning in cor(x, y): the standard deviation is zero
## Warning in cor(x, y): the standard deviation is zero
## Warning in cor(x, y): the standard deviation is zero
## Warning in cor(x, y): the standard deviation is zero
## Warning in cor(x, y): the standard deviation is zero
## Warning in cor(x, y): the standard deviation is zero
## Warning in cor(x, y): the standard deviation is zero
## Warning in cor(x, y): the standard deviation is zero
## Warning in cor(x, y): the standard deviation is zero
## Warning in cor(x, y): the standard deviation is zero
## Warning in cor(x, y): the standard deviation is zero
## Warning in cor(x, y): the standard deviation is zero
## Warning in cor(x, y): the standard deviation is zero
## Warning in cor(x, y): the standard deviation is zero
## Warning in cor(x, y): the standard deviation is zero
## Warning in cor(x, y): the standard deviation is zero
## Warning in cor(x, y): the standard deviation is zero
## Warning in cor(x, y): the standard deviation is zero
## Warning in cor(x, y): the standard deviation is zero
## Warning in cor(x, y): the standard deviation is zero
## Warning in cor(x, y): the standard deviation is zero
## Warning in cor(x, y): the standard deviation is zero
## Warning in cor(x, y): the standard deviation is zero
## Warning in cor(x, y): the standard deviation is zero
## Warning in cor(x, y): the standard deviation is zero
## Warning in cor(x, y): the standard deviation is zero
## Warning in cor(x, y): the standard deviation is zero
## Warning in cor(x, y): the standard deviation is zero
## Warning in cor(x, y): the standard deviation is zero
## Warning in cor(x, y): the standard deviation is zero
## Warning in cor(x, y): the standard deviation is zero
## Warning in cor(x, y): the standard deviation is zero
## Warning in cor(x, y): the standard deviation is zero
## Warning in cor(x, y): the standard deviation is zero
## Warning in cor(x, y): the standard deviation is zero
## Warning in cor(x, y): the standard deviation is zero
## Warning in cor(x, y): the standard deviation is zero
## Warning in cor(x, y): the standard deviation is zero
## Warning in cor(x, y): the standard deviation is zero
## Warning in cor(x, y): the standard deviation is zero
## Warning in cor(x, y): the standard deviation is zero
## Warning in cor(x, y): the standard deviation is zero
## Warning in cor(x, y): the standard deviation is zero
## Warning in cor(x, y): the standard deviation is zero
## Warning in cor(x, y): the standard deviation is zero
## Warning in cor(x, y): the standard deviation is zero
## Warning in cor(x, y): the standard deviation is zero
## Warning in cor(x, y): the standard deviation is zero
## Warning in cor(x, y): the standard deviation is zero
## Warning in cor(x, y): the standard deviation is zero
## Warning in cor(x, y): the standard deviation is zero
## Warning in cor(x, y): the standard deviation is zero
## Warning in cor(x, y): the standard deviation is zero
## Warning in cor(x, y): the standard deviation is zero
## Warning in cor(x, y): the standard deviation is zero
## Warning in cor(x, y): the standard deviation is zero
## Warning in cor(x, y): the standard deviation is zero
## Warning in cor(x, y): the standard deviation is zero
```

```
## Warning in cor(x, y): the standard deviation is zero
## Warning in cor(x, y): the standard deviation is zero
## Warning in cor(x, y): the standard deviation is zero
## Warning in cor(x, y): the standard deviation is zero
## Warning in cor(x, y): the standard deviation is zero
## Warning in cor(x, y): the standard deviation is zero
## Warning in cor(x, y): the standard deviation is zero
## Warning in cor(x, y): the standard deviation is zero
## Warning in cor(x, y): the standard deviation is zero
## Warning in cor(x, y): the standard deviation is zero
## Warning in cor(x, y): the standard deviation is zero
## Warning in cor(x, y): the standard deviation is zero
## Warning in cor(x, y): the standard deviation is zero
## Warning in cor(x, y): the standard deviation is zero
## Warning in cor(x, y): the standard deviation is zero
## Warning in cor(x, y): the standard deviation is zero
## Warning in cor(x, y): the standard deviation is zero
## Warning in cor(x, y): the standard deviation is zero
```

Next, we calculate t-test p-values to compare the correlations for each phylum between WT and KO groups to determine if there are significant differences between phylum for each phenotype.

At this time, we performed the t-test on the HM1toWT vs HM1toKO group. Results are generated as a dataframe with unadjusted p-values for each phylum. This dataframe is written out as a tab-delimited text file.

```
## ttest to compare wt and ko phylum groups

myCorr_withinGroup_pearson_phlyum_immg1toKO=getCorrelationCoefficient_withinGroup_phylumLevel_byGroup(groupList=phlyum_groups_subset_withoutSparsePhyla, tableWithMeta=myASVandMetadata_newNames, tableCountsOnly=file_asv_updatedName_t_samplesOnly_lognorm, corrTestMethod="pearson", sampleGroup = myGroup_serialPassage_miceandHuman[1], mappingMetaFile = mapping_ASVandPhyla,mappingMetaFile_taxaColName = "ASV", mappingMetaFile_phylumColName = "PhylumLabels", tableWithMeta_sampleColName = "SampleID" ,tableWithMeta_groupColName = "FMTGroupFMTsourcegtRecipientbackground")
```

```
## Warning in cor(x, y): the standard deviation is zero
## Warning in cor(x, y): the standard deviation is zero
## Warning in cor(x, y): the standard deviation is zero
## Warning in cor(x, y): the standard deviation is zero
## Warning in cor(x, y): the standard deviation is zero
## Warning in cor(x, y): the standard deviation is zero
## Warning in cor(x, y): the standard deviation is zero
## Warning in cor(x, y): the standard deviation is zero
## Warning in cor(x, y): the standard deviation is zero
## Warning in cor(x, y): the standard deviation is zero
## Warning in cor(x, y): the standard deviation is zero
## Warning in cor(x, y): the standard deviation is zero
## Warning in cor(x, y): the standard deviation is zero
## Warning in cor(x, y): the standard deviation is zero
## Warning in cor(x, y): the standard deviation is zero
## Warning in cor(x, y): the standard deviation is zero
## Warning in cor(x, y): the standard deviation is zero
## Warning in cor(x, y): the standard deviation is zero
## Warning in cor(x, y): the standard deviation is zero
## Warning in cor(x, y): the standard deviation is zero
## Warning in cor(x, y): the standard deviation is zero
## Warning in cor(x, y): the standard deviation is zero
## Warning in cor(x, y): the standard deviation is zero
## Warning in cor(x, y): the standard deviation is zero
## Warning in cor(x, y): the standard deviation is zero
## Warning in cor(x, y): the standard deviation is zero
## Warning in cor(x, y): the standard deviation is zero
## Warning in cor(x, y): the standard deviation is zero
## Warning in cor(x, y): the standard deviation is zero
## Warning in cor(x, y): the standard deviation is zero
## Warning in cor(x, y): the standard deviation is zero
## Warning in cor(x, y): the standard deviation is zero
## Warning in cor(x, y): the standard deviation is zero
## Warning in cor(x, y): the standard deviation is zero
## Warning in cor(x, y): the standard deviation is zero
## Warning in cor(x, y): the standard deviation is zero
## Warning in cor(x, y): the standard deviation is zero
## Warning in cor(x, y): the standard deviation is zero
## Warning in cor(x, y): the standard deviation is zero
## Warning in cor(x, y): the standard deviation is zero
## Warning in cor(x, y): the standard deviation is zero
## Warning in cor(x, y): the standard deviation is zero
## Warning in cor(x, y): the standard deviation is zero
## Warning in cor(x, y): the standard deviation is zero
## Warning in cor(x, y): the standard deviation is zero
## Warning in cor(x, y): the standard deviation is zero
## Warning in cor(x, y): the standard deviation is zero
## Warning in cor(x, y): the standard deviation is zero
## Warning in cor(x, y): the standard deviation is zero
## Warning in cor(x, y): the standard deviation is zero
## Warning in cor(x, y): the standard deviation is zero
## Warning in cor(x, y): the standard deviation is zero
## Warning in cor(x, y): the standard deviation is zero
## Warning in cor(x, y): the standard deviation is zero
## Warning in cor(x, y): the standard deviation is zero
## Warning in cor(x, y): the standard deviation is zero
## Warning in cor(x, y): the standard deviation is zero
## Warning in cor(x, y): the standard deviation is zero
## Warning in cor(x, y): the standard deviation is zero
## Warning in cor(x, y): the standard deviation is zero
## Warning in cor(x, y): the standard deviation is zero
## Warning in cor(x, y): the standard deviation is zero
## Warning in cor(x, y): the standard deviation is zero
## Warning in cor(x, y): the standard deviation is zero
## Warning in cor(x, y): the standard deviation is zero
## Warning in cor(x, y): the standard deviation is zero
## Warning in cor(x, y): the standard deviation is zero
## Warning in cor(x, y): the standard deviation is zero
## Warning in cor(x, y): the standard deviation is zero
## Warning in cor(x, y): the standard deviation is zero
## Warning in cor(x, y): the standard deviation is zero
## Warning in cor(x, y): the standard deviation is zero
## Warning in cor(x, y): the standard deviation is zero
## Warning in cor(x, y): the standard deviation is zero
## Warning in cor(x, y): the standard deviation is zero
## Warning in cor(x, y): the standard deviation is zero
## Warning in cor(x, y): the standard deviation is zero
## Warning in cor(x, y): the standard deviation is zero
## Warning in cor(x, y): the standard deviation is zero
## Warning in cor(x, y): the standard deviation is zero
## Warning in cor(x, y): the standard deviation is zero
## Warning in cor(x, y): the standard deviation is zero
## Warning in cor(x, y): the standard deviation is zero
## Warning in cor(x, y): the standard deviation is zero
## Warning in cor(x, y): the standard deviation is zero
## Warning in cor(x, y): the standard deviation is zero
## Warning in cor(x, y): the standard deviation is zero
## Warning in cor(x, y): the standard deviation is zero
## Warning in cor(x, y): the standard deviation is zero
## Warning in cor(x, y): the standard deviation is zero
## Warning in cor(x, y): the standard deviation is zero
## Warning in cor(x, y): the standard deviation is zero
## Warning in cor(x, y): the standard deviation is zero
## Warning in cor(x, y): the standard deviation is zero
## Warning in cor(x, y): the standard deviation is zero
## Warning in cor(x, y): the standard deviation is zero
## Warning in cor(x, y): the standard deviation is zero
## Warning in cor(x, y): the standard deviation is zero
## Warning in cor(x, y): the standard deviation is zero
## Warning in cor(x, y): the standard deviation is zero
## Warning in cor(x, y): the standard deviation is zero
## Warning in cor(x, y): the standard deviation is zero
## Warning in cor(x, y): the standard deviation is zero
## Warning in cor(x, y): the standard deviation is zero
## Warning in cor(x, y): the standard deviation is zero
## Warning in cor(x, y): the standard deviation is zero
## Warning in cor(x, y): the standard deviation is zero
## Warning in cor(x, y): the standard deviation is zero
## Warning in cor(x, y): the standard deviation is zero
## Warning in cor(x, y): the standard deviation is zero
## Warning in cor(x, y): the standard deviation is zero
## Warning in cor(x, y): the standard deviation is zero
## Warning in cor(x, y): the standard deviation is zero
## Warning in cor(x, y): the standard deviation is zero
## Warning in cor(x, y): the standard deviation is zero
## Warning in cor(x, y): the standard deviation is zero
## Warning in cor(x, y): the standard deviation is zero
## Warning in cor(x, y): the standard deviation is zero
## Warning in cor(x, y): the standard deviation is zero
## Warning in cor(x, y): the standard deviation is zero
## Warning in cor(x, y): the standard deviation is zero
## Warning in cor(x, y): the standard deviation is zero
## Warning in cor(x, y): the standard deviation is zero
## Warning in cor(x, y): the standard deviation is zero
## Warning in cor(x, y): the standard deviation is zero
## Warning in cor(x, y): the standard deviation is zero
## Warning in cor(x, y): the standard deviation is zero
## Warning in cor(x, y): the standard deviation is zero
## Warning in cor(x, y): the standard deviation is zero
## Warning in cor(x, y): the standard deviation is zero
## Warning in cor(x, y): the standard deviation is zero
## Warning in cor(x, y): the standard deviation is zero
## Warning in cor(x, y): the standard deviation is zero
## Warning in cor(x, y): the standard deviation is zero
## Warning in cor(x, y): the standard deviation is zero
## Warning in cor(x, y): the standard deviation is zero
## Warning in cor(x, y): the standard deviation is zero
## Warning in cor(x, y): the standard deviation is zero
```

```
myCorr_withinGroup_pearson_phlyum_nimmg1toWT=getCorrelationCoefficient_withinGroup_phylumLevel_byGroup(groupList=phlyum_groups_subset_withoutSparsePhyla, tableWithMeta=myASVandMetadata_newNames, tableCountsOnly=file_asv_updatedName_t_samplesOnly_lognorm, corrTestMethod="pearson", sampleGroup = myGroup_serialPassage_miceandHuman[3], mappingMetaFile = mapping_ASVandPhyla, mappingMetaFile_taxaColName = "ASV", mappingMetaFile_phylumColName = "PhylumLabels", tableWithMeta_sampleColName = "SampleID" ,tableWithMeta_groupColName = "FMTGroupFMTsourcegtRecipientbackground")
```

```
## Warning in cor(x, y): the standard deviation is zero
## Warning in cor(x, y): the standard deviation is zero
## Warning in cor(x, y): the standard deviation is zero
## Warning in cor(x, y): the standard deviation is zero
## Warning in cor(x, y): the standard deviation is zero
## Warning in cor(x, y): the standard deviation is zero
## Warning in cor(x, y): the standard deviation is zero
## Warning in cor(x, y): the standard deviation is zero
## Warning in cor(x, y): the standard deviation is zero
## Warning in cor(x, y): the standard deviation is zero
## Warning in cor(x, y): the standard deviation is zero
## Warning in cor(x, y): the standard deviation is zero
## Warning in cor(x, y): the standard deviation is zero
## Warning in cor(x, y): the standard deviation is zero
## Warning in cor(x, y): the standard deviation is zero
## Warning in cor(x, y): the standard deviation is zero
## Warning in cor(x, y): the standard deviation is zero
## Warning in cor(x, y): the standard deviation is zero
## Warning in cor(x, y): the standard deviation is zero
## Warning in cor(x, y): the standard deviation is zero
## Warning in cor(x, y): the standard deviation is zero
## Warning in cor(x, y): the standard deviation is zero
## Warning in cor(x, y): the standard deviation is zero
## Warning in cor(x, y): the standard deviation is zero
## Warning in cor(x, y): the standard deviation is zero
## Warning in cor(x, y): the standard deviation is zero
## Warning in cor(x, y): the standard deviation is zero
## Warning in cor(x, y): the standard deviation is zero
## Warning in cor(x, y): the standard deviation is zero
## Warning in cor(x, y): the standard deviation is zero
## Warning in cor(x, y): the standard deviation is zero
## Warning in cor(x, y): the standard deviation is zero
## Warning in cor(x, y): the standard deviation is zero
## Warning in cor(x, y): the standard deviation is zero
## Warning in cor(x, y): the standard deviation is zero
## Warning in cor(x, y): the standard deviation is zero
## Warning in cor(x, y): the standard deviation is zero
## Warning in cor(x, y): the standard deviation is zero
## Warning in cor(x, y): the standard deviation is zero
## Warning in cor(x, y): the standard deviation is zero
## Warning in cor(x, y): the standard deviation is zero
## Warning in cor(x, y): the standard deviation is zero
## Warning in cor(x, y): the standard deviation is zero
## Warning in cor(x, y): the standard deviation is zero
## Warning in cor(x, y): the standard deviation is zero
## Warning in cor(x, y): the standard deviation is zero
## Warning in cor(x, y): the standard deviation is zero
## Warning in cor(x, y): the standard deviation is zero
## Warning in cor(x, y): the standard deviation is zero
## Warning in cor(x, y): the standard deviation is zero
## Warning in cor(x, y): the standard deviation is zero
## Warning in cor(x, y): the standard deviation is zero
## Warning in cor(x, y): the standard deviation is zero
## Warning in cor(x, y): the standard deviation is zero
## Warning in cor(x, y): the standard deviation is zero
## Warning in cor(x, y): the standard deviation is zero
## Warning in cor(x, y): the standard deviation is zero
## Warning in cor(x, y): the standard deviation is zero
## Warning in cor(x, y): the standard deviation is zero
## Warning in cor(x, y): the standard deviation is zero
## Warning in cor(x, y): the standard deviation is zero
## Warning in cor(x, y): the standard deviation is zero
## Warning in cor(x, y): the standard deviation is zero
## Warning in cor(x, y): the standard deviation is zero
## Warning in cor(x, y): the standard deviation is zero
## Warning in cor(x, y): the standard deviation is zero
## Warning in cor(x, y): the standard deviation is zero
## Warning in cor(x, y): the standard deviation is zero
## Warning in cor(x, y): the standard deviation is zero
## Warning in cor(x, y): the standard deviation is zero
## Warning in cor(x, y): the standard deviation is zero
## Warning in cor(x, y): the standard deviation is zero
## Warning in cor(x, y): the standard deviation is zero
## Warning in cor(x, y): the standard deviation is zero
## Warning in cor(x, y): the standard deviation is zero
## Warning in cor(x, y): the standard deviation is zero
## Warning in cor(x, y): the standard deviation is zero
## Warning in cor(x, y): the standard deviation is zero
## Warning in cor(x, y): the standard deviation is zero
## Warning in cor(x, y): the standard deviation is zero
## Warning in cor(x, y): the standard deviation is zero
## Warning in cor(x, y): the standard deviation is zero
## Warning in cor(x, y): the standard deviation is zero
## Warning in cor(x, y): the standard deviation is zero
## Warning in cor(x, y): the standard deviation is zero
## Warning in cor(x, y): the standard deviation is zero
## Warning in cor(x, y): the standard deviation is zero
## Warning in cor(x, y): the standard deviation is zero
## Warning in cor(x, y): the standard deviation is zero
## Warning in cor(x, y): the standard deviation is zero
## Warning in cor(x, y): the standard deviation is zero
## Warning in cor(x, y): the standard deviation is zero
## Warning in cor(x, y): the standard deviation is zero
## Warning in cor(x, y): the standard deviation is zero
## Warning in cor(x, y): the standard deviation is zero
## Warning in cor(x, y): the standard deviation is zero
## Warning in cor(x, y): the standard deviation is zero
## Warning in cor(x, y): the standard deviation is zero
## Warning in cor(x, y): the standard deviation is zero
## Warning in cor(x, y): the standard deviation is zero
## Warning in cor(x, y): the standard deviation is zero
## Warning in cor(x, y): the standard deviation is zero
## Warning in cor(x, y): the standard deviation is zero
## Warning in cor(x, y): the standard deviation is zero
## Warning in cor(x, y): the standard deviation is zero
## Warning in cor(x, y): the standard deviation is zero
## Warning in cor(x, y): the standard deviation is zero
## Warning in cor(x, y): the standard deviation is zero
## Warning in cor(x, y): the standard deviation is zero
## Warning in cor(x, y): the standard deviation is zero
## Warning in cor(x, y): the standard deviation is zero
## Warning in cor(x, y): the standard deviation is zero
## Warning in cor(x, y): the standard deviation is zero
## Warning in cor(x, y): the standard deviation is zero
## Warning in cor(x, y): the standard deviation is zero
## Warning in cor(x, y): the standard deviation is zero
## Warning in cor(x, y): the standard deviation is zero
## Warning in cor(x, y): the standard deviation is zero
## Warning in cor(x, y): the standard deviation is zero
## Warning in cor(x, y): the standard deviation is zero
## Warning in cor(x, y): the standard deviation is zero
## Warning in cor(x, y): the standard deviation is zero
## Warning in cor(x, y): the standard deviation is zero
## Warning in cor(x, y): the standard deviation is zero
## Warning in cor(x, y): the standard deviation is zero
## Warning in cor(x, y): the standard deviation is zero
## Warning in cor(x, y): the standard deviation is zero
## Warning in cor(x, y): the standard deviation is zero
## Warning in cor(x, y): the standard deviation is zero
## Warning in cor(x, y): the standard deviation is zero
## Warning in cor(x, y): the standard deviation is zero
## Warning in cor(x, y): the standard deviation is zero
## Warning in cor(x, y): the standard deviation is zero
## Warning in cor(x, y): the standard deviation is zero
## Warning in cor(x, y): the standard deviation is zero
## Warning in cor(x, y): the standard deviation is zero
## Warning in cor(x, y): the standard deviation is zero
## Warning in cor(x, y): the standard deviation is zero
## Warning in cor(x, y): the standard deviation is zero
## Warning in cor(x, y): the standard deviation is zero
## Warning in cor(x, y): the standard deviation is zero
## Warning in cor(x, y): the standard deviation is zero
## Warning in cor(x, y): the standard deviation is zero
## Warning in cor(x, y): the standard deviation is zero
## Warning in cor(x, y): the standard deviation is zero
## Warning in cor(x, y): the standard deviation is zero
## Warning in cor(x, y): the standard deviation is zero
## Warning in cor(x, y): the standard deviation is zero
## Warning in cor(x, y): the standard deviation is zero
## Warning in cor(x, y): the standard deviation is zero
## Warning in cor(x, y): the standard deviation is zero
## Warning in cor(x, y): the standard deviation is zero
## Warning in cor(x, y): the standard deviation is zero
## Warning in cor(x, y): the standard deviation is zero
## Warning in cor(x, y): the standard deviation is zero
## Warning in cor(x, y): the standard deviation is zero
## Warning in cor(x, y): the standard deviation is zero
## Warning in cor(x, y): the standard deviation is zero
## Warning in cor(x, y): the standard deviation is zero
## Warning in cor(x, y): the standard deviation is zero
## Warning in cor(x, y): the standard deviation is zero
## Warning in cor(x, y): the standard deviation is zero
## Warning in cor(x, y): the standard deviation is zero
## Warning in cor(x, y): the standard deviation is zero
## Warning in cor(x, y): the standard deviation is zero
## Warning in cor(x, y): the standard deviation is zero
## Warning in cor(x, y): the standard deviation is zero
## Warning in cor(x, y): the standard deviation is zero
## Warning in cor(x, y): the standard deviation is zero
## Warning in cor(x, y): the standard deviation is zero
## Warning in cor(x, y): the standard deviation is zero
## Warning in cor(x, y): the standard deviation is zero
## Warning in cor(x, y): the standard deviation is zero
## Warning in cor(x, y): the standard deviation is zero
## Warning in cor(x, y): the standard deviation is zero
## Warning in cor(x, y): the standard deviation is zero
## Warning in cor(x, y): the standard deviation is zero
## Warning in cor(x, y): the standard deviation is zero
## Warning in cor(x, y): the standard deviation is zero
## Warning in cor(x, y): the standard deviation is zero
```

```
myCorr_withinGroup_pearson_phlyum_hm1toWT=getCorrelationCoefficient_withinGroup_phylumLevel_byGroup(groupList=phlyum_groups_subset_withoutSparsePhyla, tableWithMeta=myASVandMetadata_newNames, tableCountsOnly=file_asv_updatedName_t_samplesOnly_lognorm, corrTestMethod="pearson", sampleGroup = myGroup_serialPassage_miceandHuman[6], mappingMetaFile = mapping_ASVandPhyla,mappingMetaFile_taxaColName = "ASV", mappingMetaFile_phylumColName = "PhylumLabels", tableWithMeta_sampleColName = "SampleID" ,tableWithMeta_groupColName = "FMTGroupFMTsourcegtRecipientbackground")
```

```
## Warning in cor(x, y): the standard deviation is zero
## Warning in cor(x, y): the standard deviation is zero
## Warning in cor(x, y): the standard deviation is zero
## Warning in cor(x, y): the standard deviation is zero
## Warning in cor(x, y): the standard deviation is zero
## Warning in cor(x, y): the standard deviation is zero
## Warning in cor(x, y): the standard deviation is zero
## Warning in cor(x, y): the standard deviation is zero
## Warning in cor(x, y): the standard deviation is zero
## Warning in cor(x, y): the standard deviation is zero
## Warning in cor(x, y): the standard deviation is zero
## Warning in cor(x, y): the standard deviation is zero
## Warning in cor(x, y): the standard deviation is zero
## Warning in cor(x, y): the standard deviation is zero
## Warning in cor(x, y): the standard deviation is zero
## Warning in cor(x, y): the standard deviation is zero
## Warning in cor(x, y): the standard deviation is zero
## Warning in cor(x, y): the standard deviation is zero
## Warning in cor(x, y): the standard deviation is zero
## Warning in cor(x, y): the standard deviation is zero
```

```
myCorr_withinGroup_pearson_phlyum_hm1toKO=getCorrelationCoefficient_withinGroup_phylumLevel_byGroup(groupList=phlyum_groups_subset_withoutSparsePhyla, tableWithMeta=myASVandMetadata_newNames, tableCountsOnly=file_asv_updatedName_t_samplesOnly_lognorm, corrTestMethod="pearson", sampleGroup = myGroup_serialPassage_miceandHuman[5], mappingMetaFile = mapping_ASVandPhyla,mappingMetaFile_taxaColName = "ASV", mappingMetaFile_phylumColName = "PhylumLabels", tableWithMeta_sampleColName = "SampleID" ,tableWithMeta_groupColName = "FMTGroupFMTsourcegtRecipientbackground")
```

```
## Warning in cor(x, y): the standard deviation is zero
## Warning in cor(x, y): the standard deviation is zero
## Warning in cor(x, y): the standard deviation is zero
## Warning in cor(x, y): the standard deviation is zero
## Warning in cor(x, y): the standard deviation is zero
## Warning in cor(x, y): the standard deviation is zero
## Warning in cor(x, y): the standard deviation is zero
## Warning in cor(x, y): the standard deviation is zero
## Warning in cor(x, y): the standard deviation is zero
## Warning in cor(x, y): the standard deviation is zero
## Warning in cor(x, y): the standard deviation is zero
## Warning in cor(x, y): the standard deviation is zero
## Warning in cor(x, y): the standard deviation is zero
## Warning in cor(x, y): the standard deviation is zero
## Warning in cor(x, y): the standard deviation is zero
## Warning in cor(x, y): the standard deviation is zero
## Warning in cor(x, y): the standard deviation is zero
## Warning in cor(x, y): the standard deviation is zero
## Warning in cor(x, y): the standard deviation is zero
## Warning in cor(x, y): the standard deviation is zero
## Warning in cor(x, y): the standard deviation is zero
## Warning in cor(x, y): the standard deviation is zero
## Warning in cor(x, y): the standard deviation is zero
## Warning in cor(x, y): the standard deviation is zero
## Warning in cor(x, y): the standard deviation is zero
## Warning in cor(x, y): the standard deviation is zero
## Warning in cor(x, y): the standard deviation is zero
## Warning in cor(x, y): the standard deviation is zero
## Warning in cor(x, y): the standard deviation is zero
## Warning in cor(x, y): the standard deviation is zero
## Warning in cor(x, y): the standard deviation is zero
## Warning in cor(x, y): the standard deviation is zero
## Warning in cor(x, y): the standard deviation is zero
## Warning in cor(x, y): the standard deviation is zero
## Warning in cor(x, y): the standard deviation is zero
## Warning in cor(x, y): the standard deviation is zero
## Warning in cor(x, y): the standard deviation is zero
## Warning in cor(x, y): the standard deviation is zero
## Warning in cor(x, y): the standard deviation is zero
## Warning in cor(x, y): the standard deviation is zero
## Warning in cor(x, y): the standard deviation is zero
## Warning in cor(x, y): the standard deviation is zero
## Warning in cor(x, y): the standard deviation is zero
## Warning in cor(x, y): the standard deviation is zero
## Warning in cor(x, y): the standard deviation is zero
## Warning in cor(x, y): the standard deviation is zero
## Warning in cor(x, y): the standard deviation is zero
## Warning in cor(x, y): the standard deviation is zero
## Warning in cor(x, y): the standard deviation is zero
## Warning in cor(x, y): the standard deviation is zero
## Warning in cor(x, y): the standard deviation is zero
## Warning in cor(x, y): the standard deviation is zero
## Warning in cor(x, y): the standard deviation is zero
## Warning in cor(x, y): the standard deviation is zero
## Warning in cor(x, y): the standard deviation is zero
## Warning in cor(x, y): the standard deviation is zero
## Warning in cor(x, y): the standard deviation is zero
## Warning in cor(x, y): the standard deviation is zero
## Warning in cor(x, y): the standard deviation is zero
## Warning in cor(x, y): the standard deviation is zero
## Warning in cor(x, y): the standard deviation is zero
## Warning in cor(x, y): the standard deviation is zero
## Warning in cor(x, y): the standard deviation is zero
## Warning in cor(x, y): the standard deviation is zero
## Warning in cor(x, y): the standard deviation is zero
## Warning in cor(x, y): the standard deviation is zero
## Warning in cor(x, y): the standard deviation is zero
## Warning in cor(x, y): the standard deviation is zero
## Warning in cor(x, y): the standard deviation is zero
## Warning in cor(x, y): the standard deviation is zero
## Warning in cor(x, y): the standard deviation is zero
## Warning in cor(x, y): the standard deviation is zero
## Warning in cor(x, y): the standard deviation is zero
## Warning in cor(x, y): the standard deviation is zero
## Warning in cor(x, y): the standard deviation is zero
## Warning in cor(x, y): the standard deviation is zero
## Warning in cor(x, y): the standard deviation is zero
## Warning in cor(x, y): the standard deviation is zero
## Warning in cor(x, y): the standard deviation is zero
## Warning in cor(x, y): the standard deviation is zero
## Warning in cor(x, y): the standard deviation is zero
## Warning in cor(x, y): the standard deviation is zero
```

```
## Warning in cor(x, y): the standard deviation is zero

## Warning in cor(x, y): the standard deviation is zero

## Warning in cor(x, y): the standard deviation is zero

## Warning in cor(x, y): the standard deviation is zero

## Warning in cor(x, y): the standard deviation is zero

## Warning in cor(x, y): the standard deviation is zero

## Warning in cor(x, y): the standard deviation is zero

## Warning in cor(x, y): the standard deviation is zero

## Warning in cor(x, y): the standard deviation is zero

## Warning in cor(x, y): the standard deviation is zero

## Warning in cor(x, y): the standard deviation is zero

## Warning in cor(x, y): the standard deviation is zero

## Warning in cor(x, y): the standard deviation is zero

## Warning in cor(x, y): the standard deviation is zero

## Warning in cor(x, y): the standard deviation is zero

## Warning in cor(x, y): the standard deviation is zero

## Warning in cor(x, y): the standard deviation is zero
```

```
ttestpval_list = NULL
for(i in 1:length(phlyum_groups_subset_withoutSparsePhyla)){

  a = unlist(myCorr_withinGroup_pearson_phlyum_hm1toWT[[2]][i])
  b = unlist(myCorr_withinGroup_pearson_phlyum_hm1toKO[[2]][i])
  if(sum(is.na(a)) >=1 | sum(is.na(b)) >=1){print(paste0(phlyum_groups_subset_withoutSparsePhyla[i], " : NA"))} e
lse {

  ttest_pval = t.test(a,b)$p.value
  resultstoprint = paste0(phlyum_groups_subset_withoutSparsePhyla[i],
                          ": ", ttest_pval)
  ttestpval_list[i] = ttest_pval

  }
}

mydf = data.frame("Phylum"=phlyum_groups_subset_withoutSparsePhyla,
                   "Pvalue" = ttestpval_list)

write.table(mydf, here("boxplots","withinGroups_pearson","phylum_boxplot_ttest_hm1toWT_vs_hm1toKO.txt"), sep = "\
t", quote = F, row.names = F)
```

The code below generates scatterplots of samples across all groups with phylum-level counts data. This differs from the above figures, which were based on ASV counts, but was sorted according to phyla taxonomic classification.

```
dir.create(here("asvAbundance_scatterplots","acrossGroups_pearson_phylumCounts"), recursive = T)
```

```
## Warning in dir.create(here("asvAbundance_scatterplots",
## "acrossGroups_pearson_phylumCounts"), :
## 'C:\Users\aduong\Desktop\Balfour2022\asvAbundance_scatterplots\acrossGroups_pearson_phylumCounts'
## already exists
```

```
corr_acrossGroups_pearson_PhylumCounts_withHuman = getCorrelationCoefficient_acrossGroup(groupList=labels_serialp
assage_newNames, tableWithMeta=phylumCountsandMetadata, tableCountsOnly=phylumCounts_samplesOnly_df_lognorm,
corrTestMethod="pearson", variableName = "FMTGroupFMTsourcegtRecipientbackground", sampleColumnName = "SampleID")

generateScatterplots_acrossGroup(groupList=labels_serialpassage_newNames, tableWithMeta=phylumCountsandMetadata,c
orrTestMethod = "pearson"  ,
                                variableName = "FMTGroupFMTsourcegtRecipientbackground", sampleColumnName ="Samp
leID" ,tableCountsOnly=phylumCounts_samplesOnly_df_lognorm, resultsDirectory = here("asvAbundance_scatterplots","
acrossGroups_pearson_phylumCounts","scatterplot_"))
```

##### Testing homogeneity of variances

Bartlett's test checks for equality of variances between two or more groups.

Levene's test is the non-parametric version of variance equality testing.

After testing the data for normality (Shapiro-Wilkes test), the data presented with a normal distribution.

However, there were some outlier scores within some groups, so we also performed the non-parametric Levene's test for a more robust assessment of the histology scores.

```
histologyScores_hm1immg1immg2toKO = read.table(here("files", "histologyScores_hm1immg1immg2toKO.txt"), sep = "\
t", header = T)

phenotype_groups = unique(histologyScores_hm1immg1immg2toKO$Group)
colnames(histologyScores_hm1immg1immg2toKO)
regions_list = c("Ileum", "Cecum","Proximal.Colon","Distal.Colon","Rectum")

bartlettResults_list = list()
for(i in 1:length(regions_list)){
  bartlettPvalues= calculateBartlettPvalue_histologyScoreData(histologyScores_hm1immg1immg2toKO,
                                                              regionColName = regions_list[i],      groupColName = "
Group", phenotypeNames=phenotype_groups)

  bartlettResults_list[[i]]= bartlettPvalues
}

names(bartlettResults_list)=regions_list


barlettResult_totalColonScore = calculateBartlettPvalue_histologyScoreData(histologyScores_hm1immg1immg2toKO,
                                             regionColName = "Total.1",
                                             groupColName = "Group", phenotypeNames=phenotype_groups)

levenesTest_totalColonScore=calculateLevenesTestPvalue_histologyScoreData(histologyScores_hm1immg1immg2toKO,regio
nColName = "Total.1", groupColName = "Group",phenotypeNames=phenotype_groups)


colnames(histologyScores_hm1immg1immg2toKO)


leveneResults_list = list()
for(i in 1:length(regions_list)){
  levenePvalues= calculateLevenesTestPvalue_histologyScoreData(histologyScores_hm1immg1immg2toKO,
                                                              regionColName = regions_list[i], groupColName = "Grou
p",phenotypeNames=phenotype_groups)

  leveneResults_list[[i]]= levenePvalues
}

names(leveneResults_list)=regions_list
```

##### Cage Effect

We assess transfer inefficiencies for samples within the same groups per cage. Since the transfer efficiency trends were consistent with the same figure that was generated without cage data, we suggest that our initially observed transfer efficiency values were not driven by cage effect.

```
metadataNames= c("SampleID","UniversalCageNumber","Background", "FMTGroupFMTsourcegtRecipientbackground",
               "Passge", "Sex", "Agewk", "SampleType")
fmtList_withinGroup = c("HM1->WT","NIMM-g1->WT", "NIMM-g2->WT",
                        "HM1->KO", "IMM-g1->KO", "IMM-g2->KO")
withinGroups_perCage = getCorrelationCoefficient_withinGroup_perCage(tableWithMeta=myASVandMetadata_newNames,
                                                                     corrTestMethod="pearson",
                                                                     phenotypeGroup=fmtList_withinGroup,
                                                                     tableWithMeta_sampleColName="SampleID",
                                                                     tableWithMeta_groupColName="FMTGroupFMTsourcegtR
ecipientbackground",

                                                                     tableWithMeta_cageColName="UniversalCageNumber",
                                                                    metadataNames = metadataNames)



pdf(here("boxplots","Boxplots_WithinGroups_PerCage.pdf"))

par(mar=c(6, 4, 2, 2))
boxplot(withinGroups_perCage[[2]], names = withinGroups_perCage[[1]], las=2)
stripchart(withinGroups_perCage[[2]], method="jitter",add=T, pch=16, vertical = T)

dev.off()
```

##### Protein Assay: fLCN2

We compared log10 transformed and non-transformed protein assay data with both t-test and Wilcoxon test.

```
# protein assay data: fLCN2

#histology scores, not in log space

fLCN2_immg1toko = c(228.75,178.26,489.264,396.486,260.886,4679.712,381.708,
                    507.51,3328.215,1100.715,300.915,43.543,11174.805,199.605,477.075
                    )
fLCN2_nimmg1toko = c(76.281,92.0298,6.3465,13.908,195.102,370.494,142.946,292.641,
                     177.647,9.036,221.677,237.329,47.194,910.155,77.11
                      )


#log10 + 1
fLCN2_immg1toko_log10 = log10(fLCN2_immg1toko + 1)
fLCN2_nimmg1toko_log10 = log10(fLCN2_nimmg1toko + 1)

#histogram
par(mfrow=c(2,2))
hist(fLCN2_immg1toko, main="",xlim=c(0,10000))
hist(fLCN2_nimmg1toko, main="",xlim=c(0,10000))
hist(fLCN2_immg1toko_log10, main="",xlim=c(0,5))
hist(fLCN2_nimmg1toko_log10, main="",xlim=c(0,5))
```

```
# stat tests
ttest_pvals = t.test(fLCN2_immg1toko, fLCN2_nimmg1toko)$p.value
wilcoxon_pvals = wilcox.test(fLCN2_immg1toko, fLCN2_nimmg1toko)$p.value

ttest_pvals_log10input = t.test(fLCN2_immg1toko_log10, fLCN2_nimmg1toko_log10)$p.value
wilcoxon_pvals_log10input = wilcox.test(fLCN2_immg1toko_log10, fLCN2_nimmg1toko_log10)$p.value

#boxplots
pdf(here("boxplots","fLCN2_histologyScores.pdf"))

par(mfrow=c(1,2))
boxplot(list(fLCN2_immg1toko, fLCN2_nimmg1toko), names=c("IMMg1toKO", "NIMMg1toKO"),
        main=paste0("fLCN2 (Non-transformed Data)", "\nT-test Pval=", format(ttest_pvals, digits = 4),
                    ", Wilcoxon test Pval=", format(wilcoxon_pvals, digits = 4)),
        cex.main=0.85, col=c("red","royalblue"))
stripchart(list(fLCN2_immg1toko, fLCN2_nimmg1toko),
           add=TRUE, vertical = T, pch=16)

boxplot(list(fLCN2_immg1toko_log10, fLCN2_nimmg1toko_log10), names=c("IMMg1toKO", "NIMMg1toKO"),
        main=paste0("fLCN2 (Log10 transformed Data)", "\nT-test Pval=", format(ttest_pvals_log10input, digits =
4),
                    ", Wilcoxon test Pval=", format(wilcoxon_pvals_log10input, digits = 4)),
        cex.main=0.85, col=c("red","royalblue"))
stripchart(list(fLCN2_immg1toko_log10, fLCN2_nimmg1toko_log10),
           add=TRUE, vertical = T, pch=16)

dev.off()
```

### Contents


- **Aim**
- **Main Python Methods**
- **Data Processing Steps**
  - Step 1: Retrieve original counts table
  - Step 2: Retrieve original metadata table
  - Step 3: Create dictionary from metadata table
  - Step 4: Calculate total read counts per sample
  - Step 5: Extract counts for all six sample groups
  - Step 6: Calculate Spearman Correlation Coefficients for each sample group
  - Step 7: Generate final figure
  - Step 8: Compare original and regenerated figures


### Aim: Replicate Figure 1D

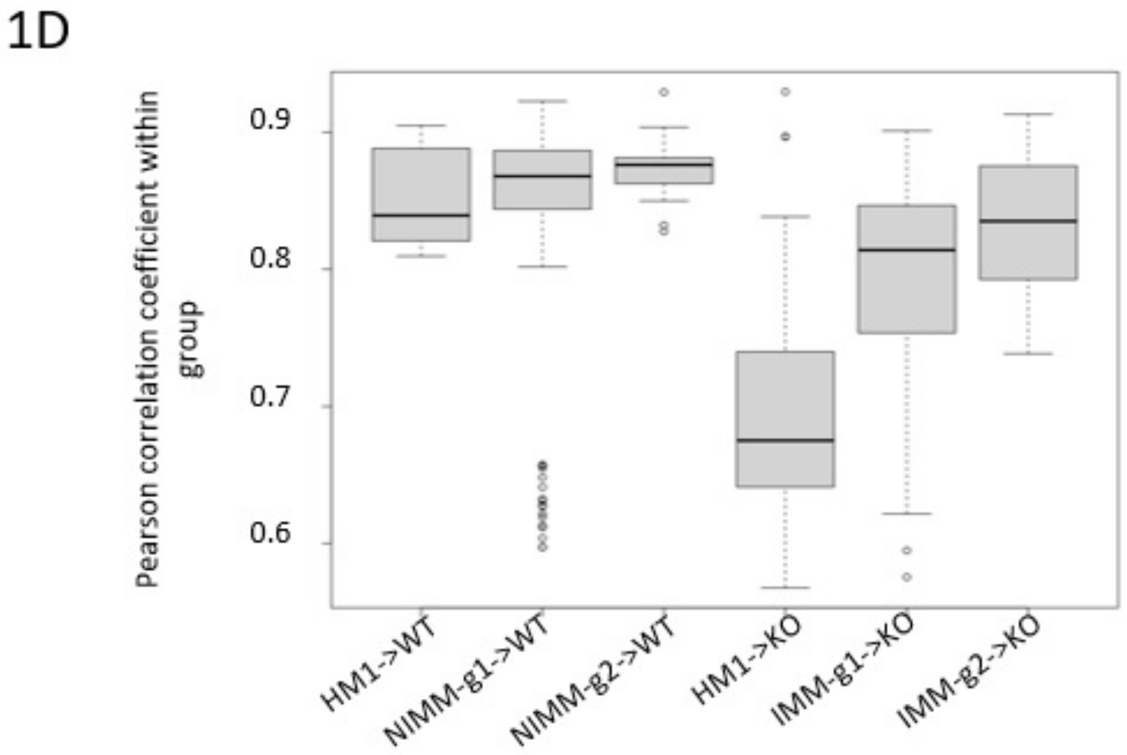


### Main Python Methods

In [1]:
```
import numpy as np
import pandas as pd
import matplotlib.pyplot as plt
import os
import re
from scipy import stats
from scipy.special import comb

# ******************************** MAIN METHODS ********************************
```

```
# Retrieve tab delimited file
def read_csv_file(file_path, skiprows=None, header = 0, sep = '\t', index_col=
False):
    df = pd.read_csv(file_path, skiprows=skiprows, sep=sep, header=header, ind
ex_col=index_col)

    return df

# Create dictionary from metadata table
def get_dict_from_metadata(input_df):
    mydict = {}
    for row in input_df.iterrows():
        obj = row[1]
        sample_id = obj['SampleID']
        key = obj['FMTGroupFMTsourcegtRecipientbackground']
        if key not in mydict:
            mydict[key] = [sample_id]
        else:
            mydict[key].append(sample_id)
    return mydict

# Calculate Pearson correlation coefficients from input set of samples
def get_pearson_corr_values(input_df, ser_sample_count_sums):
    overall_mean_count = ser_sample_count_sums.mean()
    col_names = input_df.columns
    df_input_norm_logged = _get_norm_counts(input_df, col_names, ser_sample_co
unt_sums, overall_mean_count)
    final_corr_result = _get_corr_values(col_names, df_input_norm_logged)
    return final_corr_result

# ****************************** HELPER METHODS ******************************
**
# Normalize counts table
def _get_norm_counts(input_df, col_names, ser_sample_count_sums, overall_mean_
count):
    df_norm_logged =    pd.DataFrame()
    for col_name in col_names:
        df_norm_logged.loc[:, col_name] = \
        np.log10(((input_df.loc[:, col_name] / ser_sample_count_sums[col_name
]) * overall_mean_count) + 1)

    return df_norm_logged

# Calculate Pearson Correlation Coefficients
def _get_corr_values(col_names, input_norm_logged_df):
    corr_result = []
    tup_list = _get_tup_list(col_names)

    for tup in tup_list:
        x_col_name = col_names[tup[0]]
        y_col_name = col_names[tup[1]]
        rho_value = \
        stats.pearsonr(input_norm_logged_df.loc[:, x_col_name], input_norm_log
ged_df.loc[:, y_col_name]).statistic
        corr_result.append(rho_value)

    # Remove nan values
    corr_result = [val for val in corr_result if not np.isnan(val)]

    return corr_result

# Generate tuple list of sample column names
```

```
def _get_tup_list(col_names):
    tup_list = []
    corr_result = []
    col_name_length = len(col_names)
    for i in range(col_name_length):
        for j in range(i+1, col_name_length):
            tup_list.append((i, j))

    return tup_list

# ******************************* INPUT FILE PATHS ***************************
****
# Get current working directory
current_working_dir = os.getcwd()
# original counts table
asv_tbl_file_path = os.path.join(current_working_dir, 'asv_biom-with-taxonomy.
txt')
# original metadata table
metadata_file_path = os.path.join(current_working_dir, 'mappingMetadata.txt')
```

### Data Processing Steps

### Step 1: Retrieve original counts table

In [2]:

```
df_asv = read_csv_file(asv_tbl_file_path, 1)
df_asv = df_asv.astype({col:'int32' for col in df_asv.columns[1:-1] }, copy=Fa
lse)
df_asv
```

Out[2]:

| | #OTU ID | 1gKO.1 | 1gKO.2 | 1gKO.3 | 1gWT.1 | 1gWT.2 | 1gWT.3 | 2gKO.1 |
|---|---|---|---|---|---|---|---|---|
| **0** | 1ba8c796d07406783c96d016a6a5cace | 13615 | 16637 | 17148 | 20227 | 23630 | 25656 | 14832 |
| **1** | a6c38249aff7768283faf6cfbdeb05a8 | 26439 | 30129 | 19743 | 8955 | 10759 | 7074 | 18489 |
| **2** | 062f38ff92cfaee0654200b6f5be5ddf | 7451 | 8774 | 8754 | 174 | 214 | 148 | 21958 |
| **3** | 1183cc23f552d81e63c93ca9fcba2f2c | 225 | 223 | 184 | 13762 | 16856 | 18692 | 269 |
| **4** | 5e15ecfb579e72bf87c0bea3920bbf42 | 10108 | 12117 | 8633 | 10027 | 11910 | 7424 | 5979 |
| **...** | ... | ... | ... | ... | ... | ... | ... | ... |
| **4070** | 92bb8f4683ef5c8651e7d34dbb37ab2e | 0 | 0 | 0 | 0 | 0 | 0 | 0 |
| **4071** | 92f09070a4fd5786bb34e756217e6ee1 | 0 | 0 | 0 | 0 | 0 | 0 | 0 |
| **4072** | 919b82324c41ed0046323c63aa1550da | 0 | 0 | 0 | 0 | 0 | 0 | 0 |
| **4073** | dbc0dad15ec1c8ad9d826cab94e18696 | 0 | 0 | 0 | 0 | 0 | 0 | 0 |

| 4074 | 1ff2d07d10264c23dc43e08d3097cd7c | 0 | 0 | 0 | 0 | 0 | 0 | 0 |
|---|---|---|---|---|---|---|---|---|

4075 rows × 112 columns

### Step 2: Retrieve original metadata table

In [3]:
```
df_metadata = read_csv_file(metadata_file_path)
df_metadata
```

Out[3]:

| | SampleID | UniversalCageNumber | Background | FMTGroupFMTsourcegtRecipientbackground | Passage |
|---|---|---|---|---|---|
| 0 | F8-1 | F8-cage-1 | 129.IL10KO | 1gKOgtKO | 8 |
| 1 | F8-2 | F8-cage-1 | 129.IL10KO | 1gKOgtKO | 8 |
| 2 | F8-3 | F8-cage-2 | 129.IL10KO | 1gKOgtKO | 8 |
| 3 | F8-4 | F8-cage-2 | 129.IL10KO | 1gKOgtKO | 8 |
| 4 | F8-5 | F8-cage-3 | 129.IL10KO | 1gKOgtKO | 8 |
| ... | ... | ... | ... | ... | ... |
| 105 | 1gWT.2 | NaN | NaN | 1gWTinput | 1gWT |
| 106 | 1gWT.3 | NaN | NaN | 1gWTinput | 1gWT |
| 107 | 2gWT.1 | NaN | NaN | 2gWTinput | 2gWT |
| 108 | 2gWT.2 | NaN | NaN | 2gWTinput | 2gWT |
| 109 | 2gWT.3 | NaN | NaN | 2gWTinput | 2gWT |

110 rows × 8 columns

### Step 3: Create dictionary from metadata table

In [4]:
```
dict_metadata = get_dict_from_metadata(df_metadata)
dict_metadata.keys()
```

Out[4]:
```
dict_keys(['1gKOgtKO', '2gKOgtKO', '1gWTgtKO', '1gWTgtWT', '2gWTgtWT', 'hFM
T.1.2.3.gtKO', 'hFMT.3.4.5.gtKO', 'hFMT.1.2.3.gtWT', 'hFMT.1.2.3.input', 'hF
MT.3.4.5.input', '1gKOinput', '2gKOinput', '1gWTinput', '2gWTinput'])
```

### Step 4: Calculate total read counts per sample

In [5]:
```
sample_count_sums = df_asv.iloc[:, 1:-1].sum(axis=0)
sample_count_sums
```

Out[5]:
```
1gKO.1      118256
1gKO.2      141891
1gKO.3      123292
1gWT.1      119717
1gWT.2      146158
             ...
h1-2-3.2    135326
h1-2-3.3    133745
```

```
h3-4-5.1     129613
h3-4-5.2     140316
h3-4-5.3     132984
Length: 110, dtype: int64
```

### Step 5: Extract counts for all six sample groups

In [6]:

```python
# 'hFMT.1.2.3.gtWT' -->  'HM1->WT'
key_name = 'hFMT.1.2.3.gtWT'
print('Grouped columns: {}'.format(dict_metadata[key_name]))
print('Number of expected correlation values: {}'.format(int(comb(len(dict_met
adata[key_name]), 2))))
HM1_WT = df_asv[[col_name for col_name in dict_metadata[key_name] ]]
HM1_WT
```

```
Grouped columns: ['F1-16', 'F1-17', 'F1-18', 'F1-19', 'F1-20', 'F1-21', 'F1-
22']
Number of expected correlation values: 21
```

Out[6]:

| | F1-16 | F1-17 | F1-18 | F1-19 | F1-20 | F1-21 | F1-22 |
|---|---|---|---|---|---|---|---|
| **0** | 19186 | 10727 | 29599 | 32588 | 32950 | 18853 | 5199 |
| **1** | 2938 | 6816 | 3233 | 1118 | 1710 | 1047 | 2202 |
| **2** | 9072 | 10348 | 5289 | 176 | 196 | 207 | 269 |
| **3** | 247 | 258 | 275 | 18297 | 22544 | 19687 | 21014 |
| **4** | 2389 | 3858 | 1904 | 3904 | 2928 | 5286 | 4884 |
| **...** | ... | ... | ... | ... | ... | ... | ... |
| **4070** | 0 | 0 | 0 | 0 | 0 | 0 | 0 |
| **4071** | 0 | 0 | 0 | 0 | 0 | 0 | 0 |
| **4072** | 0 | 0 | 0 | 0 | 0 | 0 | 0 |
| **4073** | 0 | 0 | 0 | 0 | 0 | 0 | 0 |
| **4074** | 0 | 0 | 0 | 0 | 0 | 0 | 0 |

4075 rows × 7 columns

In [7]:

```python
# '1gWTgtWT' --> 'NIMM-g1->WT'
key_name = '1gWTgtWT'
print('Grouped columns: {}'.format(dict_metadata[key_name]))
print('Number of expected correlation values: {}'.format(int(comb(len(dict_met
adata[key_name]), 2))))

NIMM_g1_WT = df_asv[[col_name for col_name in dict_metadata[key_name] ]]
NIMM_g1_WT
```

```
Grouped columns: ['F8-27', 'F8-28', 'F8-29', 'F8-30', 'F8-31', 'F8-32', 'F8-
33', 'F11-1', 'F11-2', 'F11-3', 'F11-4', 'F11-5', 'F11-6', 'F11-7', 'F11-8',
'F11-9', 'F11-10', 'F11-11']
Number of expected correlation values: 153
```

Out[7]:

| | F8-27 | F8-28 | F8-29 | F8-30 | F8-31 | F8-32 | F8-33 | F11-1 | F11-2 | F11-3 | F11-4 | F11-5 | F11-6 | F[illegible] |
|---|---|---|---|---|---|---|---|---|---|---|---|---|---|---|
| **0** | 21313 | 6113 | 24573 | 19347 | 18028 | 10482 | 11761 | 34742 | 24360 | 31173 | 45582 | 18929 | 22349 | 1[illegible] |
| **1** | [illegible] | [illegible] | [illegible] | [illegible] | [illegible] | [illegible] | [illegible] | [illegible] | [illegible] | [illegible] | [illegible] | [illegible] | [illegible] | [illegible] |

| | | | | | | | | | | | | | | |
|---|---|---|---|---|---|---|---|---|---|---|---|---|---|---|
| **1** | 300 | 972 | 397 | 332 | 573 | 825 | 1995 | 410 | 434 | 376 | 376 | 370 | 337 | |
| **2** | 135 | 243 | 197 | 193 | 99 | 207 | 237 | 129 | 291 | 167 | 282 | 412 | 190 | |
| **3** | 21071 | 16272 | 27247 | 21302 | 26135 | 23762 | 22465 | 25848 | 26231 | 19490 | 22181 | 18524 | 12915 | 2 |
| **4** | 2245 | 5227 | 5760 | 1189 | 2327 | 14279 | 7911 | 1416 | 2031 | 1899 | 2725 | 2679 | 1920 | [illegible] |
| **...** | ... | ... | ... | ... | ... | ... | ... | ... | ... | ... | ... | ... | ... | |
| **4070** | 0 | 0 | 0 | 0 | 0 | 0 | 0 | 0 | 0 | 0 | 0 | 0 | 0 | |
| **4071** | 0 | 0 | 0 | 0 | 0 | 0 | 0 | 0 | 0 | 0 | 0 | 0 | 0 | |
| **4072** | 0 | 0 | 0 | 0 | 0 | 0 | 0 | 0 | 0 | 0 | 0 | 0 | 0 | |
| **4073** | 0 | 0 | 0 | 0 | 0 | 0 | 0 | 0 | 0 | 0 | 0 | 0 | 0 | |
| **4074** | 0 | 0 | 0 | 0 | 0 | 0 | 0 | 0 | 0 | 0 | 0 | 0 | 0 | |

4075 rows × 18 columns

In [8]:

```
# '2gWTgtWT' --> 'NIMM-g2->WT'
key_name = '2gWTgtWT'
print('Grouped columns: {}'.format(dict_metadata[key_name]))
print('Number of expected correlation values: {}'.format(int(comb(len(dict_met
adata[key_name]), 2))))
NIMM_g2_WT = df_asv[[col_name for col_name in dict_metadata[key_name] ]]
NIMM_g2_WT
```

```
Grouped columns: ['F8-34', 'F8-35', 'F8-36', 'F8-37', 'F8-38', 'F8-39', 'F8-
40', 'F8-41']
Number of expected correlation values: 28
```

Out[8]:

| | F8-34 | F8-35 | F8-36 | F8-37 | F8-38 | F8-39 | F8-40 | F8-41 |
|---|---|---|---|---|---|---|---|---|
| **0** | 23089 | 23716 | 26435 | 13911 | 32356 | 18934 | 23958 | 18944 |
| **1** | 329 | 537 | 320 | 362 | 473 | 1848 | 258 | 282 |
| **2** | 172 | 254 | 238 | 146 | 225 | 135 | 189 | 66 |
| **3** | 11455 | 20177 | 11798 | 13910 | 13422 | 24886 | 20107 | 18889 |
| **4** | 4360 | 10659 | 3839 | 3613 | 5501 | 13735 | 2324 | 12215 |
| **...** | ... | ... | ... | ... | ... | ... | ... | ... |
| **4070** | 0 | 0 | 0 | 0 | 0 | 0 | 0 | 0 |
| **4071** | 0 | 0 | 0 | 0 | 0 | 0 | 0 | 0 |
| **4072** | 0 | 0 | 0 | 0 | 0 | 0 | 0 | 0 |
| **4073** | 0 | 0 | 0 | 0 | 0 | 0 | 0 | 0 |
| **4074** | 0 | 0 | 0 | 0 | 0 | 0 | 0 | 0 |

4075 rows × 8 columns

In [9]:

```
# 'hFMT.1.2.3.gtKO' --> 'HM1->KO'
key_name = 'hFMT.1.2.3.gtKO'
print('Grouped columns: {}'.format(dict_metadata[key_name]))
print('Number of expected correlation values: {}'.format(int(comb(len(dict_met
adata[key_name]), 2))))
HM1_KO = df_asv[[col_name for col_name in dict_metadata[key_name] ]]
HM1_KO
```

```
Grouped columns: ['F3-7', 'F3-8', 'F3-9', 'F3-10', 'F3-11', 'F3-12', 'F1-7',
'F1-8', 'F1-9', 'F1-10', 'F1-11', 'F1-12', 'F1-13', 'F1-14', 'F1-15']
```

```
Number of expected correlation values: 105
```

Out[9]:

| | F3-7 | F3-8 | F3-9 | F3-10 | F3-11 | F3-12 | F1-7 | F1-8 | F1-9 | F1-10 | F1-11 | F1-12 | F1-13 | F |
|---|---|---|---|---|---|---|---|---|---|---|---|---|---|---|
| **0** | 10097 | 574 | 12959 | 18737 | 17857 | 12650 | 10738 | 9576 | 7304 | 14409 | 14382 | 8787 | 7694 | |
| **1** | 35102 | 51633 | 31840 | 22887 | 20446 | 33188 | 23565 | 21707 | 12802 | 26719 | 5701 | 15715 | 25125 | |
| **2** | 1328 | 91 | 2203 | 189 | 182 | 244 | 33299 | 13739 | 53077 | 17570 | 34094 | 18922 | 19204 | 4 |
| **3** | 137 | 315 | 305 | 134 | 221 | 290 | 203 | 348 | 363 | 238 | 219 | 205 | 255 | |
| **4** | 34960 | 13573 | 4109 | 3811 | 1155 | 28938 | 1144 | 6973 | 666 | 8230 | 1287 | 11346 | 13195 | [illegible] |
| **...** | ... | ... | ... | ... | ... | ... | ... | ... | ... | ... | ... | ... | ... | |
| **4070** | 0 | 0 | 0 | 0 | 0 | 0 | 0 | 0 | 0 | 0 | 0 | 0 | 0 | |
| **4071** | 0 | 0 | 0 | 0 | 0 | 0 | 0 | 0 | 0 | 0 | 0 | 0 | 0 | |
| **4072** | 0 | 0 | 0 | 0 | 0 | 0 | 0 | 0 | 0 | 0 | 0 | 0 | 0 | |
| **4073** | 0 | 0 | 0 | 0 | 0 | 0 | 0 | 0 | 0 | 0 | 0 | 0 | 0 | |
| **4074** | 0 | 0 | 0 | 0 | 0 | 0 | 0 | 0 | 0 | 0 | 0 | 0 | 0 | |

4075 rows × 15 columns

In [10]:

```
# '1gKOgtKO' --> 'IMM-g1->KO'
key_name = '1gKOgtKO'
print('Grouped columns: {}'.format(dict_metadata[key_name]))
print('Number of expected correlation values: {}'.format(int(comb(len(dict_met
adata[key_name]), 2))))
IMM_g1_KO = df_asv[[col_name for col_name in dict_metadata[key_name] ]]
IMM_g1_KO
```

```
Grouped columns: ['F8-1', 'F8-2', 'F8-3', 'F8-4', 'F8-5', 'F8-6', 'F8-7', 'F
8-8', 'F4-1', 'F4-2', 'F4-3', 'F4-4', 'F4-5', 'F3-1', 'F3-2', 'F3-4', 'F3-
5', 'F3-6']
Number of expected correlation values: 153
```

Out[10]:

| | F8-1 | F8-2 | F8-3 | F8-4 | F8-5 | F8-6 | F8-7 | F8-8 | F4-1 | F4-2 | F4-3 | F4-4 | F4-5 | |
|---|---|---|---|---|---|---|---|---|---|---|---|---|---|---|
| **0** | 3339 | 15223 | 11423 | 18205 | 12486 | 10761 | 9339 | 466 | 29629 | 20134 | 27588 | 26741 | 25644 | 2 |
| **1** | 10283 | 6602 | 9314 | 13651 | 8789 | 16118 | 12045 | 13841 | 3687 | 6998 | 6904 | 7796 | 4011 | |
| **2** | 41532 | 40602 | 52337 | 11038 | 16781 | 8926 | 25758 | 31166 | 1627 | 37042 | 23825 | 26508 | 30376 | |
| **3** | 240 | 276 | 331 | 302 | 341 | 231 | 226 | 306 | 275 | 198 | 213 | 271 | 172 | 2 |
| **4** | 1806 | 6778 | 6725 | 12734 | 3901 | 10948 | 10347 | 1234 | 8891 | 3718 | 9188 | 4419 | 8315 | |
| **...** | ... | ... | ... | ... | ... | ... | ... | ... | ... | ... | ... | ... | ... | |
| **4070** | 0 | 0 | 0 | 0 | 0 | 0 | 0 | 0 | 0 | 0 | 0 | 0 | 0 | |
| **4071** | 0 | 0 | 0 | 0 | 0 | 0 | 0 | 0 | 0 | 0 | 0 | 0 | 0 | |
| **4072** | 0 | 0 | 0 | 0 | 0 | 0 | 0 | 0 | 0 | 0 | 0 | 0 | 0 | |
| **4073** | 0 | 0 | 0 | 0 | 0 | 0 | 0 | 0 | 0 | 0 | 0 | 0 | 0 | |
| **4074** | 0 | 0 | 0 | 0 | 0 | 0 | 0 | 0 | 0 | 0 | 0 | 0 | 0 | |

4075 rows × 18 columns

In [11]:

```
# '2gKOgtKO' --> 'IMM-g2->KO'
key_name = '2gKOgtKO'
```

```
print('Grouped columns: {}'.format(dict_metadata[key_name]))
print('Number of expected correlation values: {}'.format(int(comb(len(dict_met
adata[key_name]), 2))))
IMM_g2_KO = df_asv[[col_name for col_name in dict_metadata[key_name] ]]
IMM_g2_KO
```

```
Grouped columns: ['F8-9', 'F8-10', 'F8-11', 'F8-12', 'F8-13', 'F8-14', 'F8-1
5', 'F8-16']
Number of expected correlation values: 28
```

Out[11]:

| | F8-9 | F8-10 | F8-11 | F8-12 | F8-13 | F8-14 | F8-15 | F8-16 |
|---|---|---|---|---|---|---|---|---|
| **0** | 23451 | 24742 | 31312 | 13542 | 4593 | 10228 | 14477 | 23819 |
| **1** | 8030 | 14203 | 27872 | 26539 | 12663 | 13720 | 23316 | 14430 |
| **2** | 19653 | 22616 | 4603 | 28535 | 20096 | 20133 | 14650 | 24062 |
| **3** | 266 | 344 | 300 | 269 | 257 | 330 | 249 | 226 |
| **4** | 6490 | 6143 | 12353 | 2229 | 6367 | 11645 | 9607 | 10622 |
| **...** | ... | ... | ... | ... | ... | ... | ... | ... |
| **4070** | 0 | 0 | 0 | 0 | 0 | 0 | 0 | 0 |
| **4071** | 0 | 0 | 0 | 0 | 0 | 0 | 0 | 0 |
| **4072** | 0 | 0 | 0 | 0 | 0 | 0 | 0 | 0 |
| **4073** | 0 | 0 | 0 | 0 | 0 | 0 | 0 | 0 |
| **4074** | 0 | 0 | 0 | 0 | 0 | 0 | 0 | 0 |

4075 rows × 8 columns

### Step 6: Calculate Spearman Correlation Coefficients for each sample group

In [12]:

```
HM1_WT_data = get_pearson_corr_values(HM1_WT, sample_count_sums)
NIMM_g1_WT_data = get_pearson_corr_values(NIMM_g1_WT, sample_count_sums)
NIMM_g2_WT_data = get_pearson_corr_values(NIMM_g2_WT, sample_count_sums)
HM1_KO_data = get_pearson_corr_values(HM1_KO, sample_count_sums)
IMM_g1_KO_data = get_pearson_corr_values(IMM_g1_KO, sample_count_sums)
IMM_g2_KO_data = get_pearson_corr_values(IMM_g2_KO, sample_count_sums)
```

### Step 7: Generate final figure

In [13]:

```
data = [HM1_WT_data,
        NIMM_g1_WT_data,
        NIMM_g2_WT_data,
        HM1_KO_data,
        IMM_g1_KO_data,
       IMM_g2_KO_data]

sample_groups = ['HM->WT',
                'NIMM-g1->WT',
                 'NIMM-g2->WT',
                 'HM1->KO',
                 'IMM-g1->KO',
                 'IMM-g2->KO']
```

```
# Multiple box plots on one Axes
fig, ax = plt.subplots()
ax.boxplot(data, widths=0.7)
ax.set_title('Regenerated Figure 1D')
ax.set_yticks([0.6, 0.7, 0.8, 0.9])
ax.set_xticklabels(sample_groups,
                   rotation=45, fontsize=12)

plt.show()
```

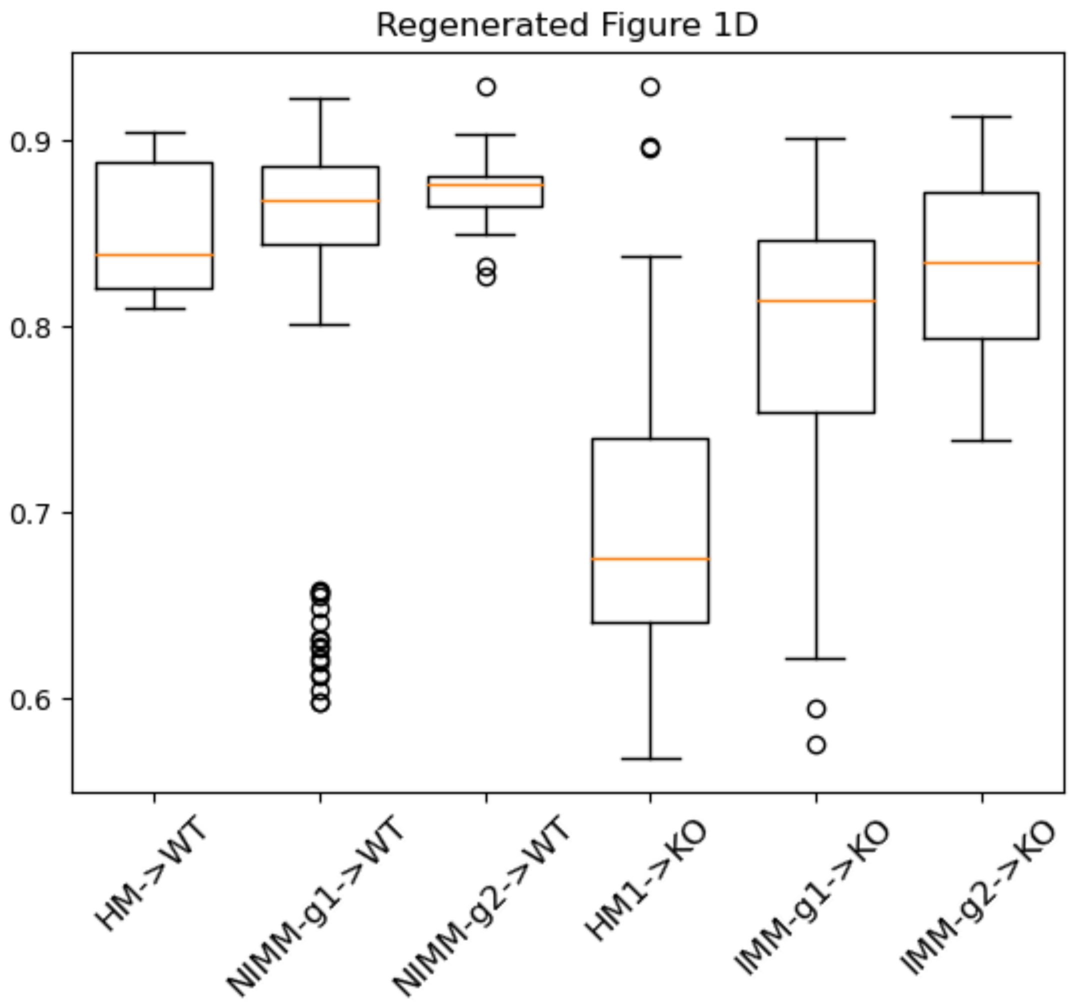


#### Step 8: Compare original and regenerated figures

1D

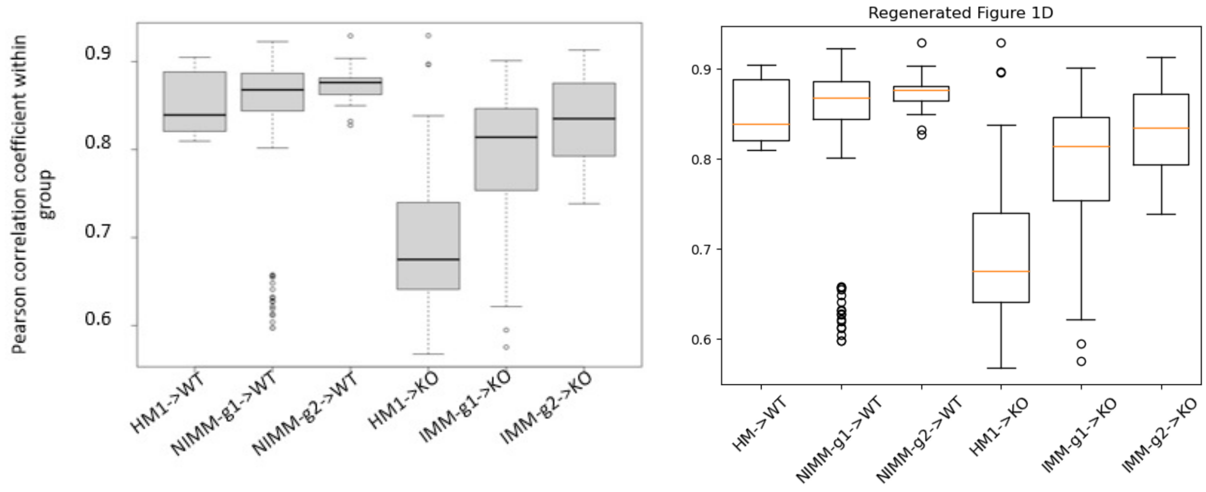

```
# Multiple box plots on one Axes
fig, ax = plt.subplots()
ax.boxplot(data, widths=0.7)
ax.set_title('Regenerated Figure 1D')
ax.set_yticks([0.6, 0.7, 0.8, 0.9])
ax.set_xticklabels(sample_groups,
                   rotation=45, fontsize=12)

plt.show()
```

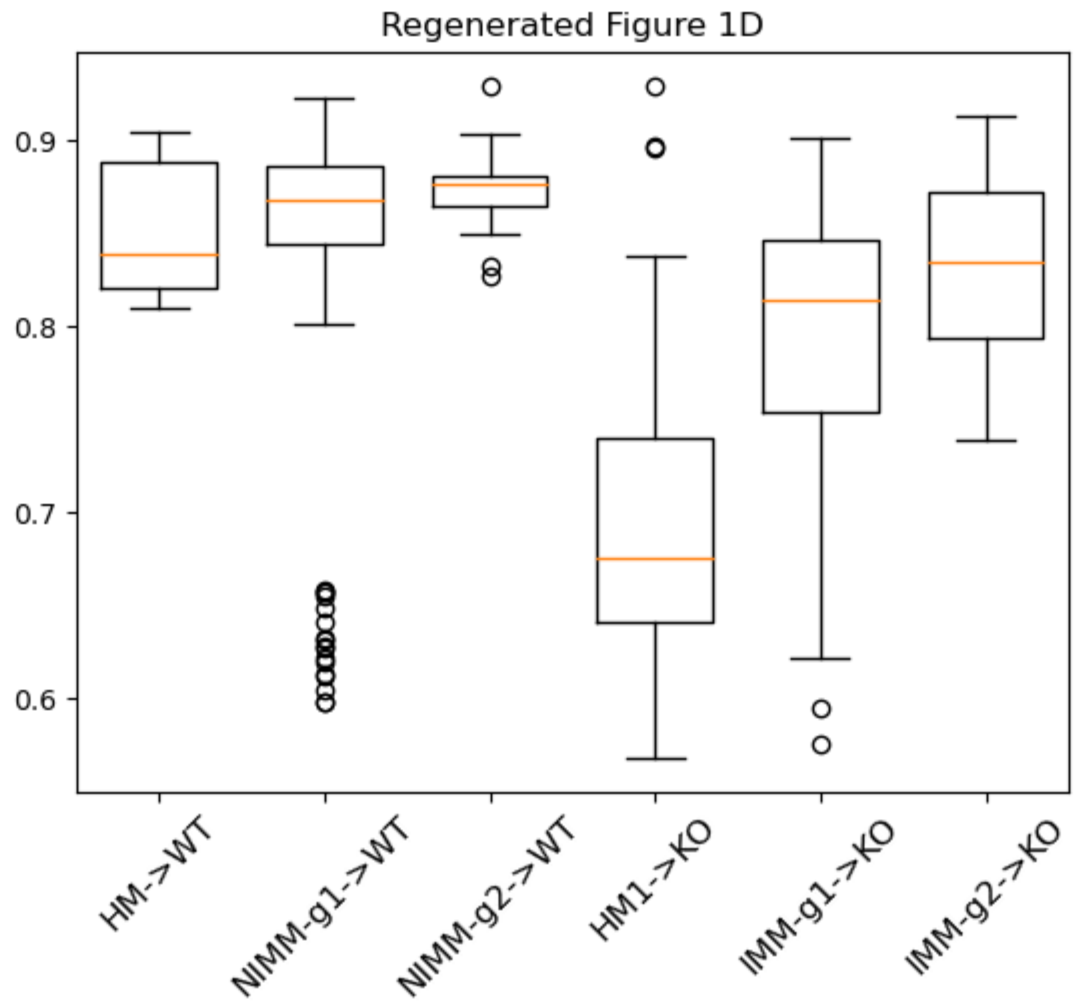


#### Step 8: Compare original and regenerated figures

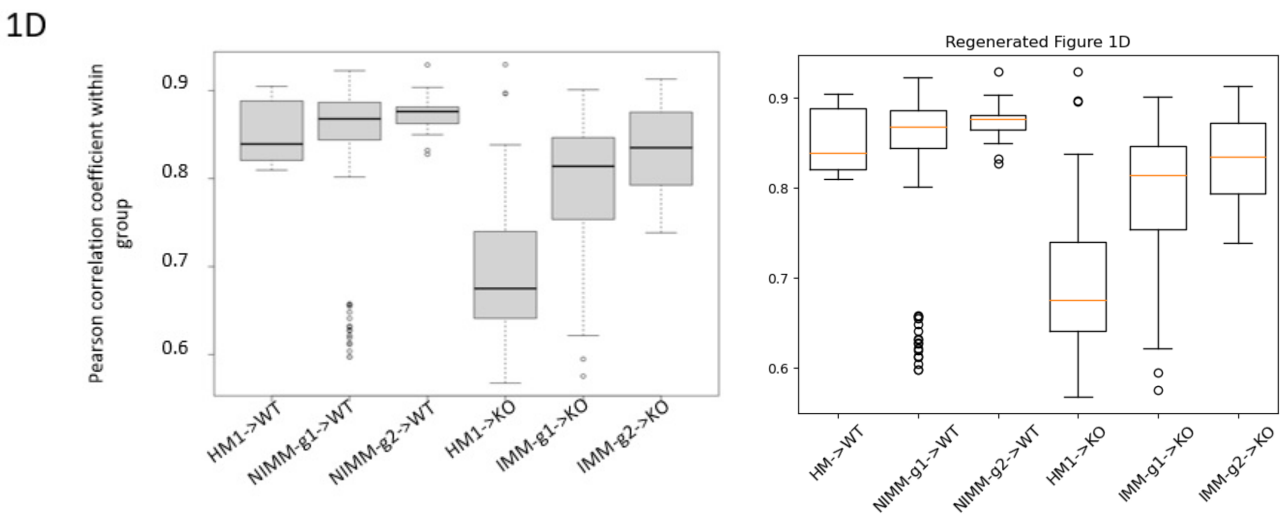

### Contents


- **Aim**
- **Main Python Methods**
- **Data Processing Steps**
  - Step 1: Retrieve original metadata table
  - Step 2: Create dictionary from metadata table
  - Step 3: Retrieve feature counts table with filtered sample columns
  - Steo 4: Calculate total read counts per sample
  - Step 5: Generate normalized counts feature table
  - Step 6: Generate Bray-Curtis dissimilarity distance matrix
  - Step 7: Perform Principal Coordinate Analysis (PCoA)
  - Step 8: Plot PCoA Results
  - Step 9: Compare Original and Regenerated Figures


### Aim: Replicate Figure 3A

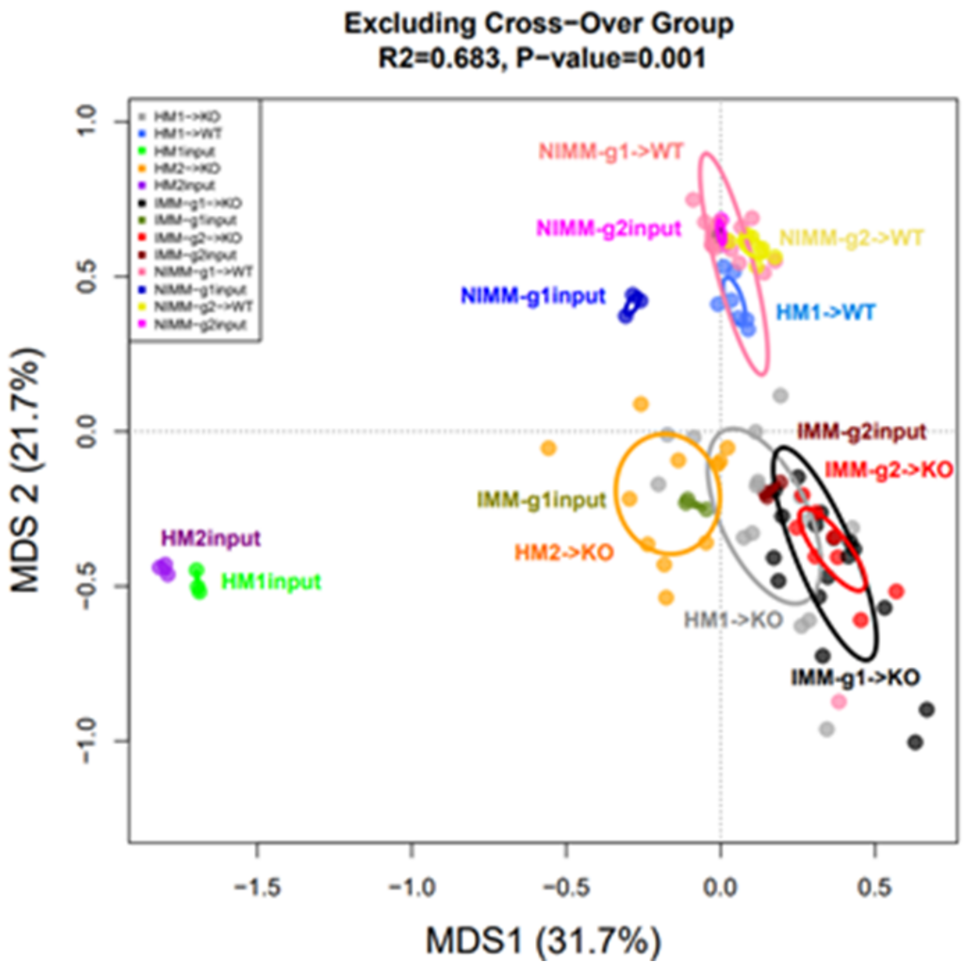


### Main Python Methods

In [1]:
```
import numpy as np
import pandas as pd
```

```
import matplotlib.pyplot as plt
import os
import re
import skbio
from scipy import stats
from scipy.spatial import distance
from matplotlib.patches import Ellipse
import matplotlib.transforms as transforms

# ******************************** MAIN METHODS ********************************
# Retrieve tab delimited file
def read_csv_file(file_path, skiprows=None, header = 0, sep = '\t', index_col=
False):
    df = pd.read_csv(file_path, skiprows=skiprows, sep=sep, header=header, ind
ex_col=index_col)

    return df

# Create dictionary from metadata table
def get_dict_from_metadata(input_df):
    mydict = {}
    for row in input_df.iterrows():
        obj = row[1]
        sample_id = obj['SampleID']
        key = obj['FMTGroupFMTsourcegtRecipientbackground']
        if key not in mydict:
            mydict[key] = [sample_id]
        else:
            mydict[key].append(sample_id)
    return mydict

def write_dataframe(input_df, file_path):
    input_df.to_csv(file_path, sep='\t', header=True, index=False)

# Reference:  Function "get_cov_ellipse" extracted from the following URL:
# https://scipython.com/book/chapter-7-matplotlib/examples/bmi-data-with-confi
dence-ellipses/
def get_cov_ellipse(cov, centre, nstd, **kwargs):
    """
    Return a matplotlib Ellipse patch representing the covariance matrix
    cov centred at centre and scaled by the factor nstd.
    """
    # Find and sort eigenvalues and eigenvectors into descending order
    eigvals, eigvecs = np.linalg.eigh(cov)
    order = eigvals.argsort()[::-1]
    eigvals, eigvecs = eigvals[order], eigvecs[:, order]

    # The anti-clockwise angle to rotate our ellipse by
    vx, vy = eigvecs[:,0][0], eigvecs[:,0][1]
    theta = np.arctan2(vy, vx)

    # Width and height of ellipse to draw
    width, height = 2 * nstd * np.sqrt(eigvals)
    return Ellipse(xy=centre, width=width, height=height,
                   angle=np.degrees(theta), fill=False, **kwargs)

# ******************************** INPUT FILE PATHS ***************************
****
# Get current working directory
current_working_dir = os.getcwd()
# original counts table
feature_tbl_file_path = os.path.join(current_working_dir, 'feature-table_balfo
```

```
ur.txt')
# original metadata table
metadata_file_path = os.path.join(current_working_dir, 'mappingMetadata.txt')

# cross reference dictionary that maps original sample group nomenclature to r
evised nomenclature
cross_ref_dict = { '1gKOgtKO':'IMM-g1->KO',
                    '2gKOgtKO':'IMM-g2->KO',
                    '1gWTgtKO':'NIMM-g1->KO',
                    '1gWTgtWT':'NIMM-g1->WT',
                    '2gWTgtWT':'NIMM-g2->WT',
                    'hFMT.1.2.3.gtKO':'HM1->KO',
                    'hFMT.3.4.5.gtKO':'HM2->KO',
                    'hFMT.1.2.3.gtWT':'HM1->WT',
                    'hFMT.1.2.3.input':'HM1input',
                    'hFMT.3.4.5.input': 'HM2input',
                    '1gKOinput':'IMM-g1input',
                    '2gKOinput':'IMM-g2input',
                    '1gWTinput':'NIMM-g1input',
                    '2gWTinput':'NIMM-g2input'
                 }

# target sample group names for Figure 3A
dict_keys_ordered = ['hFMT.1.2.3.gtKO',
                     'hFMT.1.2.3.gtWT',
                     'hFMT.1.2.3.input',
                     'hFMT.3.4.5.gtKO',
                     'hFMT.3.4.5.input',
                     '1gKOgtKO',
                     '1gKOinput',
                     '2gKOgtKO',
                     '2gKOinput',
                     '1gWTgtWT',
                     '1gWTinput',
                     '2gWTgtWT',
                     '2gWTinput']
```

### Data Processing Steps

### Step 1: Retrieve original metadata table

In [2]:

```
df_metadata = read_csv_file(metadata_file_path)
df_metadata
```

Out[2]:

| | SampleID | UniversalCageNumber | Background | FMTGroupFMTsourcegtRecipientbackground | Passage |
|---|---|---|---|---|---|
| **0** | F8-1 | F8-cage-1 | 129.IL10KO | 1gKOgtKO | 8 |
| **1** | F8-2 | F8-cage-1 | 129.IL10KO | 1gKOgtKO | 8 |
| **2** | F8-3 | F8-cage-2 | 129.IL10KO | 1gKOgtKO | 8 |
| **3** | F8-4 | F8-cage-2 | 129.IL10KO | 1gKOgtKO | 8 |
| **4** | F8-5 | F8-cage-3 | 129.IL10KO | 1gKOgtKO | 8 |
| **...** | ... | ... | ... | ... | ... |
| **105** | 1gWT.2 | NaN | NaN | 1gWTinput | 1gWT |
| **106** | 1gWT.3 | NaN | NaN | 1gWTinput | 1gWT |

| | | | | | |
|---|---|---|---|---|---|
| **107** | 2gWT.1 | NaN | NaN | 2gWTinput | 2gWT |
| **108** | 2gWT.2 | NaN | NaN | 2gWTinput | 2gWT |
| **109** | 2gWT.3 | NaN | NaN | 2gWTinput | 2gWT |

110 rows × 8 columns

### Step 2: Create dictionary from metadata table

In [3]:

```
dict_metadata = get_dict_from_metadata(df_metadata)
dict_keys_filtered = dict_keys_ordered
filtered_columns = [col for sublist in [dict_metadata[key] for key in dict_key
s_filtered] for col in sublist]
dict_keys_filtered
```

Out[3]:

```
['hFMT.1.2.3.gtKO',
 'hFMT.1.2.3.gtWT',
 'hFMT.1.2.3.input',
 'hFMT.3.4.5.gtKO',
 'hFMT.3.4.5.input',
 '1gKOgtKO',
 '1gKOinput',
 '2gKOgtKO',
 '2gKOinput',
 '1gWTgtWT',
 '1gWTinput',
 '2gWTgtWT',
 '2gWTinput']
```

### Step 3: Retrieve feature counts table with filtered sample columns

In [4]:

```
df_feature_counts = read_csv_file(feature_tbl_file_path, 1)
df_feature_counts = df_feature_counts.astype({col:'int32' for col in df_featur
e_counts.columns[1:]})
df_feature_counts
```

Out[4]:

| | #OTU ID | 1gKO.1 | 1gKO.2 | 1gKO.3 | 1gWT.1 | 1gWT.2 | 1gWT.[illegible] |
|---|---|---|---|---|---|---|---|
| **0** | d__Bacteria;p__Verrucomicrobiota;c__Verrucomic... | 11370 | 14120 | 14648 | 16632 | 19817 | 2170[illegible] |
| **1** | d__Bacteria;p__Firmicutes;c__Clostridia;o__Lac... | 22545 | 25836 | 17048 | 7547 | 9272 | 599[illegible] |
| **2** | d__Bacteria;p__Proteobacteria;c__Gammaproteoba... | 5919 | 6481 | 6714 | 31 | 31 | 3[illegible] |
| **3** | d__Bacteria;p__Firmicutes;c__Clostridia;o__Lac... | 8134 | 9827 | 7090 | 8264 | 10100 | 632[illegible] |
| **4** | d__Bacteria;p__Firmicutes;c__Clostridia;o__Lac... | 6280 | 7649 | 5719 | 9715 | 11851 | 804[illegible] |
| **...** | ... | ... | ... | ... | ... | ... | .. |
| **191** | d__Bacteria;p__Firmicutes;c__Clostridia;o__Pep... | 0 | 0 | 0 | 0 | 0 | [illegible] |
| **192** | d__Bacteria;p__Firmicutes;c__Clostridia;o__Clo... | 0 | 0 | 0 | 0 | 0 | [illegible] |
| **193** | d__Bacteria;p__Firmicutes;c__Clostridia;o__Chr... | 0 | 0 | 0 | 0 | 0 | [illegible] |
| **194** | d__Bacteria;p__Firmicutes;c__Incertae_Sedis;o_... | 0 | 0 | 0 | 0 | 0 | [illegible] |

| 195 | d__Bacteria;p__Firmicutes;c__Bacilli;o__Lactob... | 0 | 0 | 0 | 0 | 0 | ( |
|---|---|---|---|---|---|---|---|

196 rows × 111 columns

In [5]:

```
df_feature_counts_filtered = df_feature_counts[filtered_columns]
df_feature_counts_filtered
```

Out[5]:

| | F3-7 | F3-8 | F3-9 | F3-10 | F3-11 | F3-12 | F1-7 | F1-8 | F1-9 | F1-10 | ... | F8-35 | F8-36 | F8-37 |
|---|---|---|---|---|---|---|---|---|---|---|---|---|---|---|
| **0** | 8445 | 319 | 10551 | 15735 | 14058 | 10445 | 8869 | 8035 | 5953 | 11460 | ... | 20081 | 21965 | 11552 |
| **1** | 34435 | 43961 | 31682 | 20382 | 17233 | 27241 | 19991 | 23011 | 12533 | 27427 | ... | 374 | 113 | 200 |
| **2** | 978 | 13 | 1555 | 34 | 27 | 51 | 24450 | 10453 | 39194 | 12486 | ... | 64 | 55 | 14 |
| **3** | 28326 | 10652 | 3300 | 3168 | 900 | 23043 | 886 | 5723 | 480 | 6491 | ... | 8678 | 3211 | 2961 |
| **4** | 3647 | 10380 | 3639 | 3886 | 3693 | 3755 | 655 | 2478 | 205 | 5444 | ... | 6252 | 5577 | 7567 |
| **...** | ... | ... | ... | ... | ... | ... | ... | ... | ... | ... | ... | ... | ... | ... |
| **191** | 0 | 0 | 0 | 0 | 0 | 0 | 0 | 0 | 0 | 0 | ... | 0 | 0 | 0 |
| **192** | 0 | 0 | 0 | 0 | 0 | 0 | 0 | 0 | 0 | 0 | ... | 0 | 0 | 0 |
| **193** | 0 | 0 | 0 | 0 | 0 | 0 | 0 | 0 | 0 | 0 | ... | 0 | 0 | 0 |
| **194** | 0 | 0 | 0 | 0 | 0 | 0 | 0 | 0 | 0 | 0 | ... | 0 | 0 | 0 |
| **195** | 0 | 0 | 0 | 0 | 0 | 0 | 0 | 0 | 0 | 0 | ... | 0 | 0 | 0 |

196 rows × 103 columns

### Step 4: Calculate total read counts per sample

In [6]:

```
sample_count_sums = df_feature_counts_filtered.sum(axis=0)
sample_count_sums
```

Out[6]:

```
F3-7       99046
F3-8      112057
F3-9      121801
F3-10      98907
F3-11      93879
           ...
F8-40      63669
F8-41      72906
2gWT.1     86473
2gWT.2     76015
2gWT.3     78810
Length: 103, dtype: int64
```

### Step 5: Generate normalized counts feature table

In [7]:

```
df_feature_counts_filtered_norm = np.log10((df_feature_counts_filtered/sample_count_sums)*sample_count_sums.mean() + 1)
df_feature_counts_filtered_norm
```

Out[7]:

| | F3-7 | F3-8 | F3-9 | F3-10 | F3-11 | F3-12 | F1-7 | F1-8 | F1-9 | F1-1 |
|---|---|---|---|---|---|---|---|---|---|---|
| **0** | 3.904009 | 2.429163 | 3.910888 | 4.174860 | 4.148578 | 3.988447 | 3.906037 | 3.851763 | 3.766695 | 4.00565 |

| | | | | | | | | | | |
|---|---|---|---|---|---|---|---|---|---|---|
| **1** | 4.514368 | 4.566832 | 4.388371 | 4.287234 | 4.237009 | 4.404734 | 4.258967 | 4.308672 | 4.089975 | 4.38462 |
| **2** | 2.968161 | 1.075794 | 3.079633 | 1.522678 | 1.447759 | 1.686105 | 4.346407 | 3.966004 | 4.585116 | 4.04289 |
| **3** | 4.429556 | 3.951232 | 3.406225 | 3.478894 | 2.955347 | 4.332053 | 2.906082 | 3.704425 | 2.674047 | 3.75881 |
| **4** | 3.539416 | 3.940000 | 3.448677 | 3.567585 | 3.568120 | 3.544228 | 2.775080 | 3.341015 | 2.305792 | 3.68243 |
| **...** | ... | ... | ... | ... | ... | ... | ... | ... | ... | . |
| **191** | 0.000000 | 0.000000 | 0.000000 | 0.000000 | 0.000000 | 0.000000 | 0.000000 | 0.000000 | 0.000000 | 0.00000 |
| **192** | 0.000000 | 0.000000 | 0.000000 | 0.000000 | 0.000000 | 0.000000 | 0.000000 | 0.000000 | 0.000000 | 0.00000 |
| **193** | 0.000000 | 0.000000 | 0.000000 | 0.000000 | 0.000000 | 0.000000 | 0.000000 | 0.000000 | 0.000000 | 0.00000 |
| **194** | 0.000000 | 0.000000 | 0.000000 | 0.000000 | 0.000000 | 0.000000 | 0.000000 | 0.000000 | 0.000000 | 0.00000 |
| **195** | 0.000000 | 0.000000 | 0.000000 | 0.000000 | 0.000000 | 0.000000 | 0.000000 | 0.000000 | 0.000000 | 0.00000 |

196 rows × 103 columns

### Step 6: Generate Bray-Curtis dissimilarity distance matrix

In [8]:

```
col_name_list = df_feature_counts_filtered_norm.columns
num_of_cols = len(col_name_list)
dist_mat = np.zeros((num_of_cols, num_of_cols))
tup_list = [(col_name_list.get_loc(col2), col_name_list.get_loc(col1))  for co
l1 in col_name_list for col2 in col_name_list if col_name_list.get_loc(col2) >
 col_name_list.get_loc(col1)]

for tup in tup_list:
    col_1 = tup[0]
    col_2 = tup[1]
    dist = distance.braycurtis(df_feature_counts_filtered_norm.iloc[:, col_1],
 df_feature_counts_filtered_norm.iloc[:, col_2])
    dist_mat[col_1, col_2] = dist

dist_mat_sym = dist_mat + dist_mat.T
print('Dimensions of matrix: {}'.format(dist_mat_sym.shape))
dist_mat_sym
```

```
Dimensions of matrix: (103, 103)
```

Out[8]:

```
array([[0.        , 0.17459747, 0.34079072, ..., 0.32584445, 0.34141039,
        0.35352788],
       [0.17459747, 0.        , 0.38958066, ..., 0.34970683, 0.36941559,
        0.37073948],
       [0.34079072, 0.38958066, 0.        , ..., 0.30164799, 0.30201668,
        0.31426109],
       ...,
       [0.32584445, 0.34970683, 0.30164799, ..., 0.        , 0.03575365,
        0.04063024],
       [0.34141039, 0.36941559, 0.30201668, ..., 0.03575365, 0.        ,
        0.03552113],
       [0.35352788, 0.37073948, 0.31426109, ..., 0.04063024, 0.03552113,
        0.        ]])
```

### Step 7: Perform Principal Coordinate Analysis (PCoA)

In [9]:
```
my_pcoa = skbio.stats.ordination.pcoa(dist_mat_sym)
df_pcoa = my_pcoa.samples[['PC1', 'PC2']]
my_pcoa.proportion_explained
```

```
C:\ProgramData\Anaconda3\lib\site-packages\skbio\stats\ordination\_principal
_coordinate_analysis.py:143: RuntimeWarning: The result contains negative ei
genvalues. Please compare their magnitude with the magnitude of some of the
largest positive eigenvalues. If the negative ones are smaller, it's probabl
y safe to ignore them, but if they are large in magnitude, the results won't
be useful. See the Notes section for more details. The smallest eigenvalue i
s -0.03986325960166524 and the largest is 1.9638240904410669.
  warn(
```

Out[9]:
```
PC1      0.317344
PC2      0.216448
PC3      0.100538
PC4      0.059908
PC5      0.035616
           ...
PC99     0.000000
PC100    0.000000
PC101    0.000000
PC102    0.000000
PC103    0.000000
Length: 103, dtype: float64
```

### Step 8: Plot PCoA Results

In [10]:
```
group_counts = [len(dict_metadata[key]) for key in dict_keys_filtered]
group_names = [cross_ref_dict[key] for key in dict_keys_filtered]
colors = ["#A6A6A6", "#4169E1", "#00FF00", "#FFA500", "#A020F0",
                    "#000000", "#6B8E23", "#FF0000", "#8B0000",
                     "#FF82AB", "#0000CD", "#EEEE00", "#FF00FF"]
text_tup = [(-0.05, -0.09),
            (0.04, -0.02),
            (0.02, -0.01),
            (-0.13,-0.06),
            (0.0, 0.02),
            (-0.02, -0.1),
            (-0.175, -0.01),
            (-0.01, 0.06),
            (0.03, 0.04),
            (-0.15, 0.08),
            (-0.13, 0.0),
            (0.03, 0.0),
            (-0.14, 0.0)]

fig, ax = plt.subplots()
ax.set_ylim((-0.3, 0.3))
ax.set_xlabel('MDS1 ({}%)'.format(np.round(my_pcoa.proportion_explained.PC1 *
100, 1)))
ax.set_ylabel('MDS2 ({}%)'.format(np.round(my_pcoa.proportion_explained.PC2 *
100, 1)))
ax.set_title('Regenerated Figure 3A')

end = 0
for idx, group_count in enumerate(group_counts):
    start = end
```

```
    end = start + group_count

    # multiply by -1 to rotate vector 180 degrees in order to match figure 3A
    pc1 = df_pcoa.iloc[start:end, 0] * -1.0
    # multiply by -1 to rotate vector 180 degrees in order to match figure 3A
    pc2 = df_pcoa.iloc[start:end, 1] * -1.0

    group_name = group_names[idx]
    ax.scatter(pc1, pc2, c=colors[idx],  label=group_name,
               alpha=1.0, edgecolors='none')
    cov = np.cov(pc1,pc2)
    x_mean = pc1.mean()
    y_mean = pc2.mean()
    e = get_cov_ellipse(cov, (x_mean, y_mean), 1,
                        ec=colors[idx], linewidth=2.0, alpha=0.4)
    ax.text(x_mean + text_tup[idx][0], y_mean + text_tup[idx][1], group_name,
fontsize='x-small', c=colors[idx])
    ax.add_artist(e)

ax.legend(fontsize=6, loc='upper left')
plt.show()
```

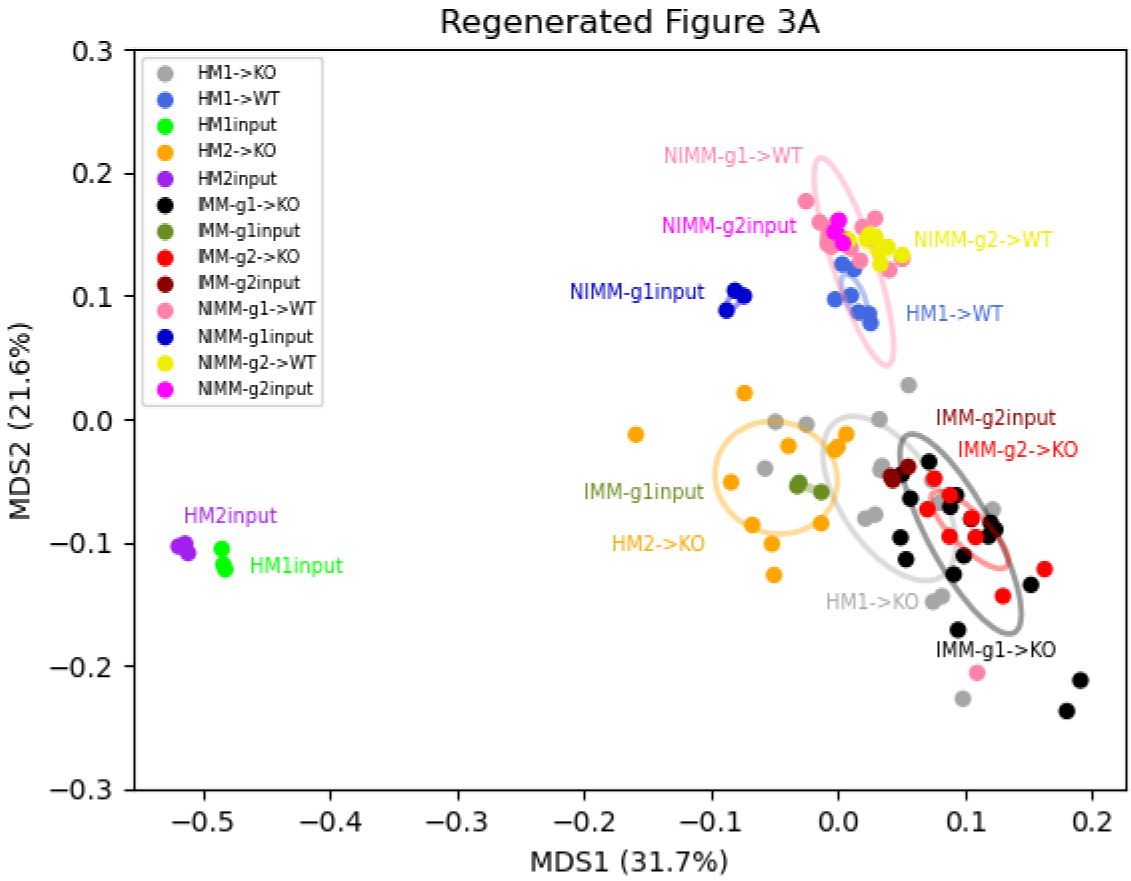


#### Step 9: Compare Original and Regenerated Figures

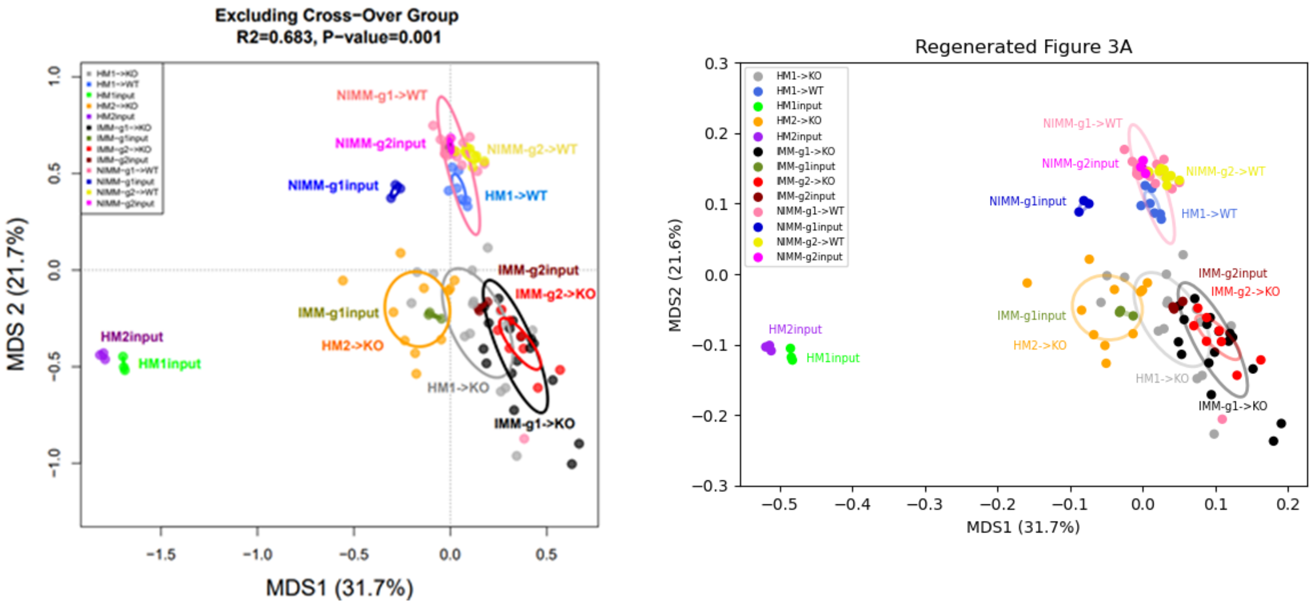
Excluding Cross-Over Group
R2=0.683, P-value=0.001
MDS1 (31.7%)
MDS 2 (21.7%)
NIMM-g1->WT
NIMM-g2input
NIMM-g2->WT
NIMM-g1input
HM1->WT
IMM-g2input
IMM-g2->KO
IMM-g1input
HM2->KO
HM1->KO
IMM-g1->KO
HM2input
HM1input
Regenerated Figure 3A
MDS2 (21.6%)
HM1->KO
HM1->WT
HM1input
HM2->KO
HM2input
IMM-g1->KO
IMM-g1input
IMM-g2->KO
IMM-g2input
NIMM-g1->WT
NIMM-g1input
NIMM-g2->WT
NIMM-g2input

### Contents


- **Aim**
- **Main Python Methods**
- **Data Processing Steps**
  - Step 1: Retrieve original counts table
  - Step 2: Retrieve original metadata table
  - Step 3: Create dictionary from metadata table
  - Step 4: Calculate total read counts per sample
  - Step 5: Extract counts for all six sample groups
  - Step 6: Normalize count values for all six sample groups
  - Step 7: Calculate Spearman Correlation Coefficients for each sample group
  - Step 8: Generate final figure
  - Step 9: Compare original and regenerated figures


### Aim: Replicate Figure 4B

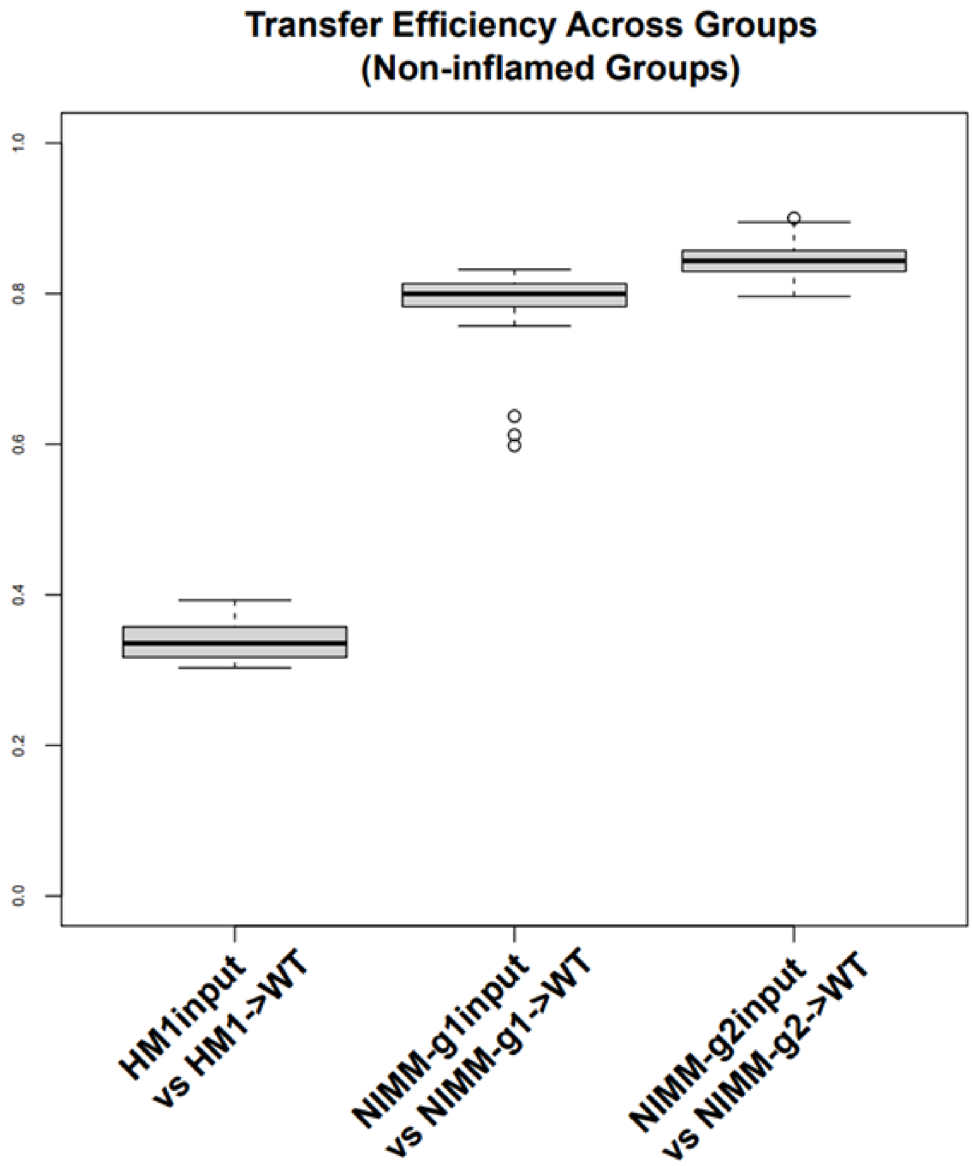

#### Main Python Methods

In [20]:
```
import numpy as np
import pandas as pd
import matplotlib.pyplot as plt
import os
import re
from scipy import stats
from scipy.special import comb

# ****************************** MAIN METHODS ******************************
# calculate Pearson correlation coefficients from input set of samples
def get_inter_corr_values(df1, df2):
    corr_values = [stats.pearsonr(df1.loc[:, col_A], df2.loc[:, col_B]).statis
tic \
            for col_A in df1.columns for col_B in df2.columns]
    # Remove nan values
    corr_values = [val for val in corr_values if not np.isnan(val)]

    return corr_values

# Retrieve tab delimited file
def read_csv_file(file_path, skiprows=None, header = 0, sep = '\t', index_col=
False):
    df = pd.read_csv(file_path, skiprows=skiprows, sep=sep, header=header, ind
ex_col=index_col)

    return df

# Create dictionary from metadata table
def get_dict_from_metadata(input_df):
    mydict = {}
    for row in input_df.iterrows():
        obj = row[1]
        sample_id = obj['SampleID']
        key = obj['FMTGroupFMTsourcegtRecipientbackground']
        if key not in mydict:
            mydict[key] = [sample_id]
        else:
            mydict[key].append(sample_id)
    return mydict

# ****************************** HELPER METHODS *****************************
**
# Normalize sample counts
def _get_norm_counts(input_df, ser_sample_count_sums):
    overall_mean_count = ser_sample_count_sums.mean()
    df_norm_logged =    pd.DataFrame()
    for col_name in input_df.columns:
        df_norm_logged.loc[:, col_name] = \
        np.log10(((input_df.loc[:, col_name] / ser_sample_count_sums[col_name
]) * overall_mean_count) + 1)

    return df_norm_logged

# Get tuples of sample column names
def _get_tup_list(col_names):
    tup_list = []
    corr_result = []
    col_name_length = len(col_names)
    for i in range(col_name_length):
```

```
        for j in range(i+1, col_name_length):
            tup_list.append((i, j))

    return tup_list

# ******************************** INPUT FILE PATHS ****************************
****
# Get current working directory
current_working_dir = os.getcwd()
# original counts table
asv_tbl_file_path = os.path.join(current_working_dir, 'asv_biom-with-taxonomy.
txt')
# original metadata table
metadata_file_path = os.path.join(current_working_dir, 'mappingMetadata.txt')
```

### Data Processing Steps

### Step 1: Retrieve original counts table

In [21]:

```
df_asv = read_csv_file(asv_tbl_file_path, 1)
df_asv = df_asv.astype({col:'int32' for col in df_asv.columns[1:-1] }, copy=Fa
lse)
df_asv
```

Out[21]:

| | #OTU ID | 1gKO.1 | 1gKO.2 | 1gKO.3 | 1gWT.1 | 1gWT.2 | 1gWT.3 | 2gKO.1 |
|---|---|---|---|---|---|---|---|---|
| **0** | 1ba8c796d07406783c96d016a6a5cace | 13615 | 16637 | 17148 | 20227 | 23630 | 25656 | 14832 |
| **1** | a6c38249aff7768283faf6cfbdeb05a8 | 26439 | 30129 | 19743 | 8955 | 10759 | 7074 | 18489 |
| **2** | 062f38ff92cfaee0654200b6f5be5ddf | 7451 | 8774 | 8754 | 174 | 214 | 148 | 21958 |
| **3** | 1183cc23f552d81e63c93ca9fcba2f2c | 225 | 223 | 184 | 13762 | 16856 | 18692 | 269 |
| **4** | 5e15ecfb579e72bf87c0bea3920bbf42 | 10108 | 12117 | 8633 | 10027 | 11910 | 7424 | 5979 |
| **...** | ... | ... | ... | ... | ... | ... | ... | ... |
| **4070** | 92bb8f4683ef5c8651e7d34dbb37ab2e | 0 | 0 | 0 | 0 | 0 | 0 | 0 |
| **4071** | 92f09070a4fd5786bb34e756217e6ee1 | 0 | 0 | 0 | 0 | 0 | 0 | 0 |
| **4072** | 919b82324c41ed0046323c63aa1550da | 0 | 0 | 0 | 0 | 0 | 0 | 0 |
| **4073** | dbc0dad15ec1c8ad9d826cab94e18696 | 0 | 0 | 0 | 0 | 0 | 0 | 0 |
| **4074** | 1ff2d07d10264c23dc43e08d3097cd7c | 0 | 0 | 0 | 0 | 0 | 0 | 0 |

4075 rows × 112 columns

### Step 2: Retrieve original metadata table

In [22]:
```
df_metadata = read_csv_file(metadata_file_path)
df_metadata
```

Out[22]:

| | SampleID | UniversalCageNumber | Background | FMTGroupFMTsourcegtRecipientbackground | Passage |
|---|---|---|---|---|---|
| **0** | F8-1 | F8-cage-1 | 129.IL10KO | 1gKOgtKO | 8 |
| **1** | F8-2 | F8-cage-1 | 129.IL10KO | 1gKOgtKO | 8 |
| **2** | F8-3 | F8-cage-2 | 129.IL10KO | 1gKOgtKO | 8 |
| **3** | F8-4 | F8-cage-2 | 129.IL10KO | 1gKOgtKO | 8 |
| **4** | F8-5 | F8-cage-3 | 129.IL10KO | 1gKOgtKO | 8 |
| **...** | ... | ... | ... | ... | ... |
| **105** | 1gWT.2 | NaN | NaN | 1gWTinput | 1gWT |
| **106** | 1gWT.3 | NaN | NaN | 1gWTinput | 1gWT |
| **107** | 2gWT.1 | NaN | NaN | 2gWTinput | 2gWT |
| **108** | 2gWT.2 | NaN | NaN | 2gWTinput | 2gWT |
| **109** | 2gWT.3 | NaN | NaN | 2gWTinput | 2gWT |

110 rows × 8 columns

### Step 3: Create dictionary from metadata table

In [4]:
```
dict_metadata = get_dict_from_metadata(df_metadata)
dict_metadata.keys()
```

Out[4]:
```
dict_keys(['1gKOgtKO', '2gKOgtKO', '1gWTgtKO', '1gWTgtWT', '2gWTgtWT', 'hFM
T.1.2.3.gtKO', 'hFMT.3.4.5.gtKO', 'hFMT.1.2.3.gtWT', 'hFMT.1.2.3.input', 'hF
MT.3.4.5.input', '1gKOinput', '2gKOinput', '1gWTinput', '2gWTinput'])
```

### Step 4: Calculate total read counts per sample

In [5]:
```
sample_count_sums = df_asv.iloc[:, 1:-1].sum(axis=0)
sample_count_sums
```

Out[5]:
```
1gKO.1      118256
1gKO.2      141891
1gKO.3      123292
1gWT.1      119717
1gWT.2      146158
             ...
h1-2-3.2    135326
h1-2-3.3    133745
h3-4-5.1    129613
h3-4-5.2    140316
h3-4-5.3    132984
Length: 110, dtype: int64
```

### Step 5: Extract counts for target sample groups

In [6]:
```
# 'hFMT.1.2.3.input' -->  'HM1Input'
key_name ='hFMT.1.2.3.input'
print('Grouped columns: {}'.format(dict_metadata[key_name]))
HM1_Input = df_asv[[col_name for col_name in dict_metadata[key_name] ]]
HM1_Input
```

```
Grouped columns: ['h1-2-3.1', 'h1-2-3.2', 'h1-2-3.3']
```

Out[6]:

| | h1-2-3.1 | h1-2-3.2 | h1-2-3.3 |
|---|---|---|---|
| **0** | 1483 | 1515 | 1402 |
| **1** | 1129 | 691 | 725 |
| **2** | 2927 | 3523 | 3254 |
| **3** | 298 | 231 | 238 |
| **4** | 218 | 160 | 111 |
| **...** | ... | ... | ... |
| **4070** | 0 | 0 | 0 |
| **4071** | 0 | 0 | 0 |
| **4072** | 0 | 0 | 0 |
| **4073** | 0 | 0 | 0 |
| **4074** | 0 | 0 | 0 |

4075 rows × 3 columns

In [7]:
```
# 'hFMT.1.2.3.gtWT' -->  'HM1->WT'
key_name = 'hFMT.1.2.3.gtWT'
print('Grouped columns: {}'.format(dict_metadata[key_name]))
print('Number of expected correlation values: {}'.format(int(comb(len(dict_met
adata[key_name]), 2))))
HM1_WT = df_asv[[col_name for col_name in dict_metadata[key_name] ]]
HM1_WT
```

```
Grouped columns: ['F1-16', 'F1-17', 'F1-18', 'F1-19', 'F1-20', 'F1-21', 'F1-
22']
Number of expected correlation values: 21
```

Out[7]:

| | F1-16 | F1-17 | F1-18 | F1-19 | F1-20 | F1-21 | F1-22 |
|---|---|---|---|---|---|---|---|
| **0** | 19186 | 10727 | 29599 | 32588 | 32950 | 18853 | 5199 |
| **1** | 2938 | 6816 | 3233 | 1118 | 1710 | 1047 | 2202 |
| **2** | 9072 | 10348 | 5289 | 176 | 196 | 207 | 269 |
| **3** | 247 | 258 | 275 | 18297 | 22544 | 19687 | 21014 |
| **4** | 2389 | 3858 | 1904 | 3904 | 2928 | 5286 | 4884 |
| **...** | ... | ... | ... | ... | ... | ... | ... |
| **4070** | 0 | 0 | 0 | 0 | 0 | 0 | 0 |
| **4071** | 0 | 0 | 0 | 0 | 0 | 0 | 0 |
| **4072** | 0 | 0 | 0 | 0 | 0 | 0 | 0 |

| | | | | | | | |
|---|---|---|---|---|---|---|---|
| **4072** | 0 | 0 | 0 | 0 | 0 | 0 | 0 |
| **4073** | 0 | 0 | 0 | 0 | 0 | 0 | 0 |
| **4074** | 0 | 0 | 0 | 0 | 0 | 0 | 0 |

4075 rows × 7 columns

In [8]:
```
# '1gWTinput' -->  'NIMM-g1input'
key_name ='1gWTinput'
print('Grouped columns: {}'.format(dict_metadata[key_name]))
NIMM_g1input = df_asv[[col_name for col_name in dict_metadata[key_name] ]]
NIMM_g1input
```

```
Grouped columns: ['1gWT.1', '1gWT.2', '1gWT.3']
```

Out[8]:

| | 1gWT.1 | 1gWT.2 | 1gWT.3 |
|---|---|---|---|
| **0** | 20227 | 23630 | 25656 |
| **1** | 8955 | 10759 | 7074 |
| **2** | 174 | 214 | 148 |
| **3** | 13762 | 16856 | 18692 |
| **4** | 10027 | 11910 | 7424 |
| **...** | ... | ... | ... |
| **4070** | 0 | 0 | 0 |
| **4071** | 0 | 0 | 0 |
| **4072** | 0 | 0 | 0 |
| **4073** | 0 | 0 | 0 |
| **4074** | 0 | 0 | 0 |

4075 rows × 3 columns

In [9]:
```
# '1gWTgtWT' -->  'NIMM-g1->WT'
key_name ='1gWTgtWT'
print('Grouped columns: {}'.format(dict_metadata[key_name]))
NIMM_g1_WT = df_asv[[col_name for col_name in dict_metadata[key_name] ]]
NIMM_g1_WT
```

```
Grouped columns: ['F8-27', 'F8-28', 'F8-29', 'F8-30', 'F8-31', 'F8-32', 'F8-
33', 'F11-1', 'F11-2', 'F11-3', 'F11-4', 'F11-5', 'F11-6', 'F11-7', 'F11-8',
'F11-9', 'F11-10', 'F11-11']
```

Out[9]:

| | F8-27 | F8-28 | F8-29 | F8-30 | F8-31 | F8-32 | F8-33 | F11-1 | F11-2 | F11-3 | F11-4 | F11-5 | F11-6 | F |
|---|---|---|---|---|---|---|---|---|---|---|---|---|---|---|
| **0** | 21313 | 6113 | 24573 | 19347 | 18028 | 10482 | 11761 | 34742 | 24360 | 31173 | 45582 | 18929 | 22349 | 1 |
| **1** | 300 | 972 | 397 | 332 | 573 | 825 | 1995 | 410 | 434 | 376 | 376 | 370 | 337 | |
| **2** | 135 | 243 | 197 | 193 | 99 | 207 | 237 | 129 | 291 | 167 | 282 | 412 | 190 | |
| **3** | 21071 | 16272 | 27247 | 21302 | 26135 | 23762 | 22465 | 25848 | 26231 | 19490 | 22181 | 18524 | 12915 | 2 |
| **4** | 2245 | 5227 | 5760 | 1189 | 2327 | 14279 | 7911 | 1416 | 2031 | 1899 | 2725 | 2679 | 1920 | [illegible] |
| **...** | ... | ... | ... | ... | ... | ... | ... | ... | ... | ... | ... | ... | ... | |
| **4070** | 0 | 0 | 0 | 0 | 0 | 0 | 0 | 0 | 0 | 0 | 0 | 0 | 0 | |
| **4071** | 0 | 0 | 0 | 0 | 0 | 0 | 0 | 0 | 0 | 0 | 0 | 0 | 0 | |

| | | | | | | | | | | | | | |
|---|---|---|---|---|---|---|---|---|---|---|---|---|---|
| **4071** | 0 | 0 | 0 | 0 | 0 | 0 | 0 | 0 | 0 | 0 | 0 | 0 | 0 |
| **4072** | 0 | 0 | 0 | 0 | 0 | 0 | 0 | 0 | 0 | 0 | 0 | 0 | 0 |
| **4073** | 0 | 0 | 0 | 0 | 0 | 0 | 0 | 0 | 0 | 0 | 0 | 0 | 0 |
| **4074** | 0 | 0 | 0 | 0 | 0 | 0 | 0 | 0 | 0 | 0 | 0 | 0 | 0 |

4075 rows × 18 columns

In [10]:

```
# '2gWTinput' -->  'NIMM-g2input'
key_name ='2gWTinput'
print('Grouped columns: {}'.format(dict_metadata[key_name]))
NIMM_g2input = df_asv[[col_name for col_name in dict_metadata[key_name] ]]
NIMM_g2input
```

```
Grouped columns: ['2gWT.1', '2gWT.2', '2gWT.3']
```

Out[10]:

| | 2gWT.1 | 2gWT.2 | 2gWT.3 |
|---|---|---|---|
| **0** | 26154 | 27846 | 29545 |
| **1** | 487 | 298 | 403 |
| **2** | 221 | 152 | 77 |
| **3** | 18419 | 20399 | 21089 |
| **4** | 2446 | 1550 | 1249 |
| **...** | ... | ... | ... |
| **4070** | 0 | 0 | 0 |
| **4071** | 0 | 0 | 0 |
| **4072** | 0 | 0 | 0 |
| **4073** | 0 | 0 | 0 |
| **4074** | 0 | 0 | 0 |

4075 rows × 3 columns

In [11]:

```
# '2gWTgtWT' -->  'NIMM-g2->WT'
key_name ='2gWTgtWT'
print('Grouped columns: {}'.format(dict_metadata[key_name]))
NIMM_g2_WT = df_asv[[col_name for col_name in dict_metadata[key_name] ]]
NIMM_g2_WT
```

```
Grouped columns: ['F8-34', 'F8-35', 'F8-36', 'F8-37', 'F8-38', 'F8-39', 'F8-
40', 'F8-41']
```

Out[11]:

| | F8-34 | F8-35 | F8-36 | F8-37 | F8-38 | F8-39 | F8-40 | F8-41 |
|---|---|---|---|---|---|---|---|---|
| **0** | 23089 | 23716 | 26435 | 13911 | 32356 | 18934 | 23958 | 18944 |
| **1** | 329 | 537 | 320 | 362 | 473 | 1848 | 258 | 282 |
| **2** | 172 | 254 | 238 | 146 | 225 | 135 | 189 | 66 |
| **3** | 11455 | 20177 | 11798 | 13910 | 13422 | 24886 | 20107 | 18889 |
| **4** | 4360 | 10659 | 3839 | 3613 | 5501 | 13735 | 2324 | 12215 |
| **...** | ... | ... | ... | ... | ... | ... | ... | ... |
| **4070** | 0 | 0 | 0 | 0 | 0 | 0 | 0 | 0 |
| **4071** | 0 | 0 | 0 | 0 | 0 | 0 | 0 | 0 |
| **4072** | 0 | 0 | 0 | 0 | 0 | 0 | 0 | 0 |

| | | | | | | | | |
|---|---|---|---|---|---|---|---|---|
| **4072** | 0 | 0 | 0 | 0 | 0 | 0 | 0 | 0 |
| **4073** | 0 | 0 | 0 | 0 | 0 | 0 | 0 | 0 |
| **4074** | 0 | 0 | 0 | 0 | 0 | 0 | 0 | 0 |

4075 rows × 8 columns

### Step 6: Normalize count values for target sample groups

In [12]:

```
HM1_Input_norm = _get_norm_counts(HM1_Input, sample_count_sums)
HM1_Input_norm
```

Out[12]:

| | h1-2-3.1 | h1-2-3.2 | h1-2-3.3 |
|---|---|---|---|
| **0** | 3.170992 | 3.168281 | 3.139741 |
| **1** | 3.052636 | 2.827698 | 2.853624 |
| **2** | 3.466129 | 3.534613 | 3.505231 |
| **3** | 2.475230 | 2.353116 | 2.371106 |
| **4** | 2.340004 | 2.194478 | 2.041962 |
| **...** | ... | ... | ... |
| **4070** | 0.000000 | 0.000000 | 0.000000 |
| **4071** | 0.000000 | 0.000000 | 0.000000 |
| **4072** | 0.000000 | 0.000000 | 0.000000 |
| **4073** | 0.000000 | 0.000000 | 0.000000 |
| **4074** | 0.000000 | 0.000000 | 0.000000 |

4075 rows × 3 columns

In [13]:

```
HM1_WT_norm = _get_norm_counts(HM1_WT, sample_count_sums)
HM1_WT_norm
```

Out[13]:

| | F1-16 | F1-17 | F1-18 | F1-19 | F1-20 | F1-21 | F1-22 |
|---|---|---|---|---|---|---|---|
| **0** | 4.245267 | 3.999519 | 4.464795 | 4.552695 | 4.457225 | 4.238876 | 3.673620 |
| **1** | 3.430471 | 3.802595 | 3.503245 | 3.088422 | 3.172643 | 2.983868 | 3.300643 |
| **2** | 3.920013 | 3.983899 | 3.716960 | 2.287385 | 2.234151 | 2.281717 | 2.389137 |
| **3** | 2.356868 | 2.382421 | 2.434436 | 4.302027 | 4.292408 | 4.257674 | 4.280140 |
| **4** | 3.340672 | 3.555481 | 3.273402 | 3.631237 | 3.406096 | 3.686687 | 3.646482 |
| **...** | ... | ... | ... | ... | ... | ... | ... |
| **4070** | 0.000000 | 0.000000 | 0.000000 | 0.000000 | 0.000000 | 0.000000 | 0.000000 |
| **4071** | 0.000000 | 0.000000 | 0.000000 | 0.000000 | 0.000000 | 0.000000 | 0.000000 |
| **4072** | 0.000000 | 0.000000 | 0.000000 | 0.000000 | 0.000000 | 0.000000 | 0.000000 |
| **4073** | 0.000000 | 0.000000 | 0.000000 | 0.000000 | 0.000000 | 0.000000 | 0.000000 |
| **4074** | 0.000000 | 0.000000 | 0.000000 | 0.000000 | 0.000000 | 0.000000 | 0.000000 |

4075 rows × 7 columns

In [14]:
```
NIMM_g1input_norm = _get_norm_counts(NIMM_g1input, sample_count_sums)
NIMM_g1input_norm
```

Out[14]:

| | 1gWT.1 | 1gWT.2 | 1gWT.3 |
|---|---|---|---|
| **0** | 4.346750 | 4.327617 | 4.390865 |
| **1** | 3.992909 | 3.985949 | 3.831387 |
| **2** | 2.283615 | 2.286796 | 2.154970 |
| **3** | 4.179510 | 4.180916 | 4.253338 |
| **4** | 4.042010 | 4.030085 | 3.852357 |
| **...** | ... | ... | ... |
| **4070** | 0.000000 | 0.000000 | 0.000000 |
| **4071** | 0.000000 | 0.000000 | 0.000000 |
| **4072** | 0.000000 | 0.000000 | 0.000000 |
| **4073** | 0.000000 | 0.000000 | 0.000000 |
| **4074** | 0.000000 | 0.000000 | 0.000000 |

4075 rows × 3 columns

In [15]:
```
NIMM_g1_WT_norm = _get_norm_counts(NIMM_g1_WT, sample_count_sums)
NIMM_g1_WT_norm
```

Out[15]:

| | F8-27 | F8-28 | F8-29 | F8-30 | F8-31 | F8-32 | F8-33 | F11-1 | F11-2 | F11 |
|---|---|---|---|---|---|---|---|---|---|---|
| **0** | 4.269712 | 3.868181 | 4.305071 | 4.313638 | 4.241945 | 4.043398 | 4.160521 | 4.485555 | 4.333532 | 4.4697 |
| **1** | 2.419820 | 3.069904 | 2.514711 | 2.549369 | 2.744909 | 2.939868 | 3.390166 | 2.558671 | 2.585453 | 2.5523 |
| **2** | 2.075047 | 2.468951 | 2.211733 | 2.314670 | 1.986115 | 2.340871 | 2.466282 | 2.059083 | 2.412411 | 2.2014 |
| **3** | 4.264752 | 4.293331 | 4.349929 | 4.355443 | 4.403212 | 4.398815 | 4.441569 | 4.357132 | 4.365668 | 4.2658 |
| **4** | 3.292482 | 3.800189 | 3.675105 | 3.102528 | 3.352961 | 4.177641 | 3.988323 | 3.096097 | 3.254786 | 3.2547 |
| **...** | ... | ... | ... | ... | ... | ... | ... | ... | ... | |
| **4070** | 0.000000 | 0.000000 | 0.000000 | 0.000000 | 0.000000 | 0.000000 | 0.000000 | 0.000000 | 0.000000 | 0.0000 |
| **4071** | 0.000000 | 0.000000 | 0.000000 | 0.000000 | 0.000000 | 0.000000 | 0.000000 | 0.000000 | 0.000000 | 0.0000 |
| **4072** | 0.000000 | 0.000000 | 0.000000 | 0.000000 | 0.000000 | 0.000000 | 0.000000 | 0.000000 | 0.000000 | 0.0000 |
| **4073** | 0.000000 | 0.000000 | 0.000000 | 0.000000 | 0.000000 | 0.000000 | 0.000000 | 0.000000 | 0.000000 | 0.0000 |
| **4074** | 0.000000 | 0.000000 | 0.000000 | 0.000000 | 0.000000 | 0.000000 | 0.000000 | 0.000000 | 0.000000 | 0.0000 |

4075 rows × 18 columns

In [16]:
```
NIMM_g2input_norm = _get_norm_counts(NIMM_g2input, sample_count_sums)
NIMM_g2input_norm
```

Out[16]:

| | 2gWT.1 | 2gWT.2 | 2gWT.3 |
|---|---|---|---|
| **0** | 4.410161 | 4.472699 | 4.474499 |
| **1** | 2.681041 | 2.503502 | 2.610372 |
| **2** | 2.338993 | 2.212436 | 1.896044 |

| | | | |
|---|---|---|---|
| **3** | 4.257896 | 4.337551 | 4.328077 |
| **4** | 3.381243 | 3.218516 | 3.100908 |
| **...** | ... | ... | ... |
| **4070** | 0.000000 | 0.000000 | 0.000000 |
| **4071** | 0.000000 | 0.000000 | 0.000000 |
| **4072** | 0.000000 | 0.000000 | 0.000000 |
| **4073** | 0.000000 | 0.000000 | 0.000000 |
| **4074** | 0.000000 | 0.000000 | 0.000000 |

4075 rows × 3 columns

In [17]:
```
NIMM_g2_WT_norm = _get_norm_counts(NIMM_g2_WT, sample_count_sums)
NIMM_g2_WT_norm
```

Out[17]:

| | F8-34 | F8-35 | F8-36 | F8-37 | F8-38 | F8-39 | F8-40 | F8-41 |
|---|---|---|---|---|---|---|---|---|
| **0** | 4.326180 | 4.345537 | 4.473561 | 4.148775 | 4.388825 | 4.288827 | 4.446174 | 4.297041 |
| **1** | 2.481386 | 2.701315 | 2.557721 | 2.565278 | 2.554926 | 3.278493 | 2.479769 | 2.471266 |
| **2** | 2.201025 | 2.377136 | 2.429562 | 2.172667 | 2.233579 | 2.145017 | 2.345136 | 1.845337 |
| **3** | 4.021791 | 4.275356 | 4.123208 | 4.148744 | 4.006713 | 4.407534 | 4.370073 | 4.295778 |
| **4** | 3.602349 | 3.998236 | 3.635686 | 3.563373 | 3.619399 | 4.149421 | 3.433104 | 4.106475 |
| **...** | ... | ... | ... | ... | ... | ... | ... | ... |
| **4070** | 0.000000 | 0.000000 | 0.000000 | 0.000000 | 0.000000 | 0.000000 | 0.000000 | 0.000000 |
| **4071** | 0.000000 | 0.000000 | 0.000000 | 0.000000 | 0.000000 | 0.000000 | 0.000000 | 0.000000 |
| **4072** | 0.000000 | 0.000000 | 0.000000 | 0.000000 | 0.000000 | 0.000000 | 0.000000 | 0.000000 |
| **4073** | 0.000000 | 0.000000 | 0.000000 | 0.000000 | 0.000000 | 0.000000 | 0.000000 | 0.000000 |
| **4074** | 0.000000 | 0.000000 | 0.000000 | 0.000000 | 0.000000 | 0.000000 | 0.000000 | 0.000000 |

4075 rows × 8 columns

### Step 7: Calculate Spearman Correlation Coefficients for each sample group

In [18]:
```
corr_value1 = get_inter_corr_values(HM1_Input_norm, HM1_WT_norm)
corr_value2 = get_inter_corr_values(NIMM_g1input_norm, NIMM_g1_WT_norm)
corr_value3 = get_inter_corr_values(NIMM_g2input_norm, NIMM_g2_WT_norm)
```

### Step 8: Generate final figure

In [25]:
```
data = []
x_tick_labels = ['HM1input\nvs HM1->WT',
                 'NIMM-g1input\nvs NIMM-g1->WT',
                 'NIMM-g2input\nvs NIMM-g2->WT']

data.extend([corr_value1, corr_value2, corr_value3])
fig, ax = plt.subplots()
```

```
ax.boxplot(data, widths=0.7)
ax.set_title('Regenerated Figure 4B')
ax.set_yticks([0.0, 0.2, 0.4, 0.6, 0.8, 1.0])
ax.set_xticklabels(x_tick_labels,
                   rotation=45, fontsize=10, fontweight='bold')

plt.show()
```

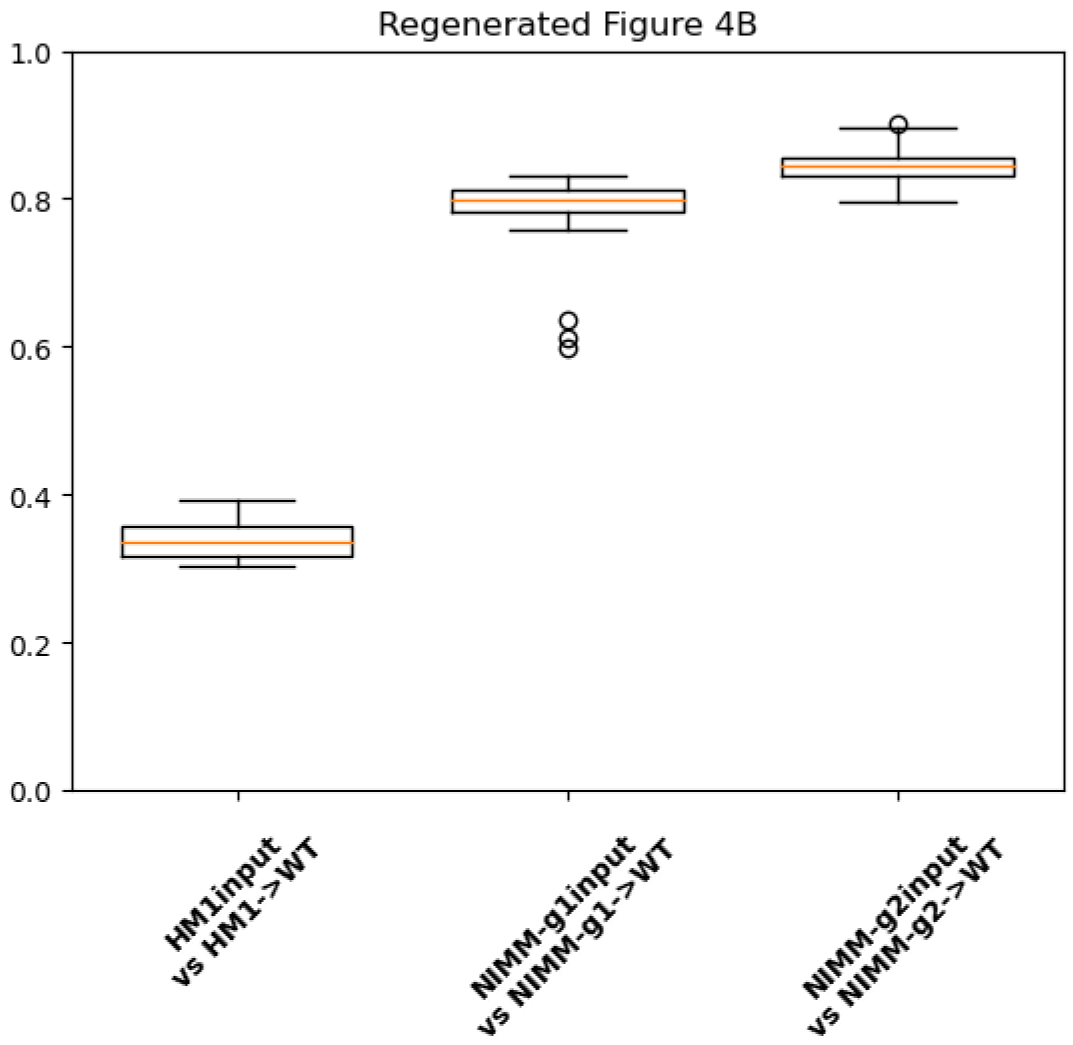


#### Step 9: Compare original and regenerated figures

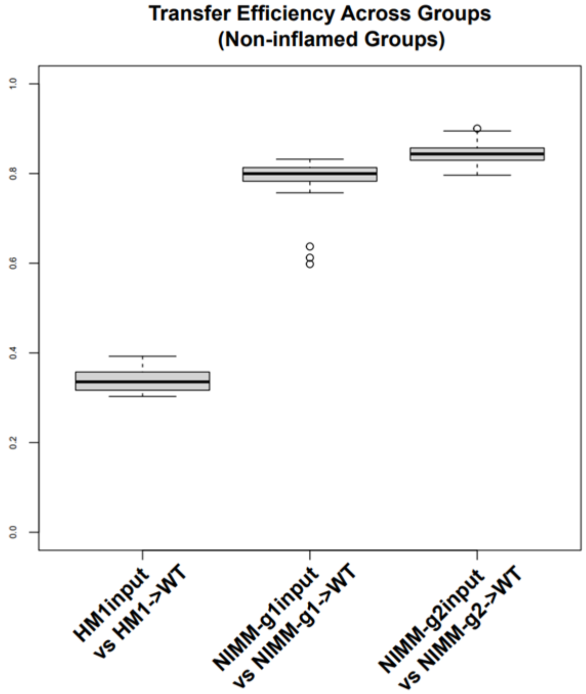
Transfer Efficiency Across Groups
(Non-inflamed Groups)
1.0
0.8
0.6
0.4
0.2
0.0
HM1input
vs HM1->WT
NIMM-g1input
vs NIMM-g1->WT
NIMM-g2input
vs NIMM-g2->WT

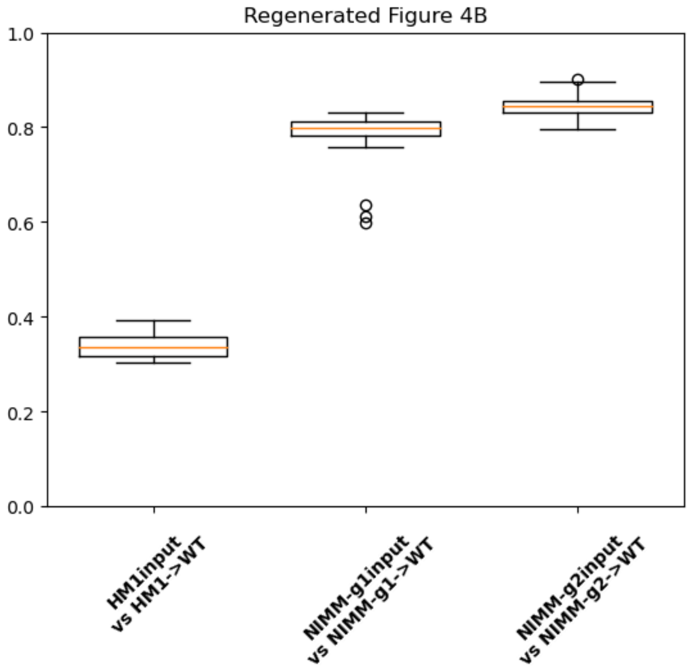
Regenerated Figure 4B
1.0
0.8
0.6
0.4
0.2
0.0
HM1input
vs HM1->WT
NIMM-g1input
vs NIMM-g1->WT
NIMM-g2input
vs NIMM-g2->WT

In [ ]:

### Contents


- **Aim**
- **Main Python Methods**
- **Data Processing Steps**
    - Step 1: Retrieve original counts table
    - Step 2: Retrieve original metadata table
    - Step 3: Create dictionary from metadata table
    - Step 4: Calculate total read counts per sample
    - Step 5: Extract counts for all six sample groups
    - Step 6: Normalize count values for all six sample groups
    - Step 7: Calculate Spearman Correlation Coefficients for each sample group
    - Step 8: Generate final figure
    - Step 9: Compare original and regenerated figures


### Aim: Replicate Figure 4D

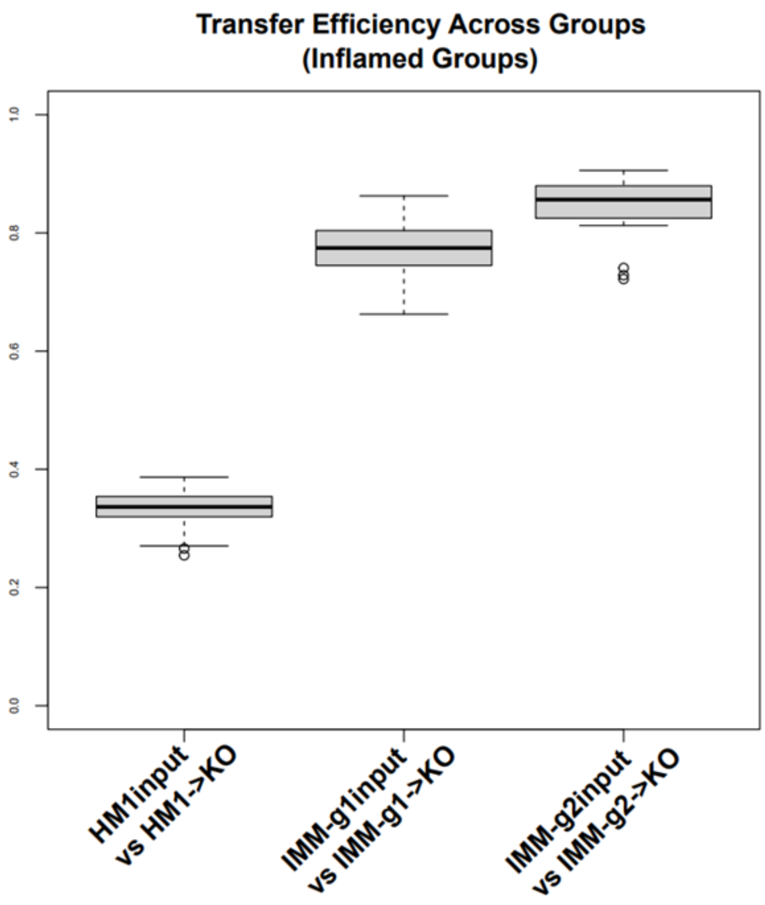


### Main Python Methods

In [1]:
```
import numpy as np
import pandas as pd
import matplotlib.pyplot as plt
import os
```

```
import re
from scipy import stats
from scipy.special import comb

# ******************************* MAIN METHODS ******************************
# Calculate Pearson correlation coefficients from input set of samples
def get_inter_corr_values(df1, df2):
    corr_values = [stats.pearsonr(df1.loc[:, col_A], df2.loc[:, col_B]).statis
tic \
            for col_A in df1.columns for col_B in df2.columns]
    # Remove nan values
    corr_values = [val for val in corr_values if not np.isnan(val)]

    return corr_values

# Retrieve tab delimited file
def read_csv_file(file_path, skiprows=None, header = 0, sep = '\t', index_col=
False):
    df = pd.read_csv(file_path, skiprows=skiprows, sep=sep, header=header, ind
ex_col=index_col)

    return df

# Create dictionary from metadata table
def get_dict_from_metadata(input_df):
    mydict = {}
    for row in input_df.iterrows():
        obj = row[1]
        sample_id = obj['SampleID']
        key = obj['FMTGroupFMTsourcegtRecipientbackground']
        if key not in mydict:
            mydict[key] = [sample_id]
        else:
            mydict[key].append(sample_id)
    return mydict

# ******************************* HELPER METHODS *****************************
**
# Normalize values in counts table
def _get_norm_counts(input_df, ser_sample_count_sums):
    overall_mean_count = ser_sample_count_sums.mean()
    df_norm_logged =    pd.DataFrame()
    for col_name in input_df.columns:
        df_norm_logged.loc[:, col_name] = \
        np.log10(((input_df.loc[:, col_name] / ser_sample_count_sums[col_name
]) * overall_mean_count) + 1)

    return df_norm_logged

# ******************************* INPUT FILE PATHS ***************************
****
current_working_dir = os.getcwd()
base_dir = r'C:\Users\young\Documents\anh_validation'
# original counts table
asv_tbl_file_path = os.path.join(base_dir, 'asv_biom-with-taxonomy.txt')
# original metadata table
metadata_file_path = os.path.join(base_dir, 'mappingMetadata.txt')
```

### Data Processing Steps

### Step 1: Retrieve original counts table

In [2]:
```
df_asv = read_csv_file(asv_tbl_file_path, 1)
df_asv = df_asv.astype({col:'int32' for col in df_asv.columns[1:-1] }, copy=Fa
lse)
df_asv
```

Out[2]:

| | #OTU ID | 1gKO.1 | 1gKO.2 | 1gKO.3 | 1gWT.1 | 1gWT.2 | 1gWT.3 | 2gKO.1 |
|---|---|---|---|---|---|---|---|---|
| **0** | 1ba8c796d07406783c96d016a6a5cace | 13615 | 16637 | 17148 | 20227 | 23630 | 25656 | 14832 |
| **1** | a6c38249aff7768283faf6cfbdeb05a8 | 26439 | 30129 | 19743 | 8955 | 10759 | 7074 | 18489 |
| **2** | 062f38ff92cfaee0654200b6f5be5ddf | 7451 | 8774 | 8754 | 174 | 214 | 148 | 21958 |
| **3** | 1183cc23f552d81e63c93ca9fcba2f2c | 225 | 223 | 184 | 13762 | 16856 | 18692 | 269 |
| **4** | 5e15ecfb579e72bf87c0bea3920bbf42 | 10108 | 12117 | 8633 | 10027 | 11910 | 7424 | 5979 |
| **...** | ... | ... | ... | ... | ... | ... | ... | ... |
| **4070** | 92bb8f4683ef5c8651e7d34dbb37ab2e | 0 | 0 | 0 | 0 | 0 | 0 | 0 |
| **4071** | 92f09070a4fd5786bb34e756217e6ee1 | 0 | 0 | 0 | 0 | 0 | 0 | 0 |
| **4072** | 919b82324c41ed0046323c63aa1550da | 0 | 0 | 0 | 0 | 0 | 0 | 0 |
| **4073** | dbc0dad15ec1c8ad9d826cab94e18696 | 0 | 0 | 0 | 0 | 0 | 0 | 0 |
| **4074** | 1ff2d07d10264c23dc43e08d3097cd7c | 0 | 0 | 0 | 0 | 0 | 0 | 0 |

4075 rows × 112 columns

### Step 2: Retrieve original metadata table

In [3]:
```
df_metadata = read_csv_file(metadata_file_path)
df_metadata
```

Out[3]:

| | SampleID | UniversalCageNumber | Background | FMTGroupFMTsourcegtRecipientbackground | Passage |
|---|---|---|---|---|---|
| **0** | F8-1 | F8-cage-1 | 129.IL10KO | 1gKOgtKO | 8 |
| **1** | F8-2 | F8-cage-1 | 129.IL10KO | 1gKOgtKO | 8 |
| **2** | F8-3 | F8-cage-2 | 129.IL10KO | 1gKOgtKO | 8 |
| **3** | F8-4 | F8-cage-2 | 129.IL10KO | 1gKOgtKO | 8 |
| **4** | F8-5 | F8-cage-3 | 129.IL10KO | 1gKOgtKO | 8 |
| **...** | ... | ... | ... | ... | ... |

| | | | | | |
|---|---|---|---|---|---|
| **105** | 1gWT.2 | NaN | NaN | 1gWTinput | 1gWT |
| **106** | 1gWT.3 | NaN | NaN | 1gWTinput | 1gWT |
| **107** | 2gWT.1 | NaN | NaN | 2gWTinput | 2gWT |
| **108** | 2gWT.2 | NaN | NaN | 2gWTinput | 2gWT |
| **109** | 2gWT.3 | NaN | NaN | 2gWTinput | 2gWT |

110 rows × 8 columns

### Step 3: Create dictionary from metadata table

In [4]:

```
dict_metadata = get_dict_from_metadata(df_metadata)
dict_metadata.keys()
```

Out[4]:

```
dict_keys(['1gKOgtKO', '2gKOgtKO', '1gWTgtKO', '1gWTgtWT', '2gWTgtWT', 'hFM
T.1.2.3.gtKO', 'hFMT.3.4.5.gtKO', 'hFMT.1.2.3.gtWT', 'hFMT.1.2.3.input', 'hF
MT.3.4.5.input', '1gKOinput', '2gKOinput', '1gWTinput', '2gWTinput'])
```

### Step 4: Calculate total read counts per sample

In [5]:

```
sample_count_sums = df_asv.iloc[:, 1:-1].sum(axis=0)
sample_count_sums
```

Out[5]:

```
1gKO.1       118256
1gKO.2       141891
1gKO.3       123292
1gWT.1       119717
1gWT.2       146158
              ...
h1-2-3.2     135326
h1-2-3.3     133745
h3-4-5.1     129613
h3-4-5.2     140316
h3-4-5.3     132984
Length: 110, dtype: int64
```

### Step 5: Extract counts for target sample groups

In [6]:

```
# 'hFMT.1.2.3.input' -->  'HM1Input'
key_name ='hFMT.1.2.3.input'
print('Grouped columns: {}'.format(dict_metadata[key_name]))
HM1_Input = df_asv[[col_name for col_name in dict_metadata[key_name] ]]
HM1_Input
```

```
Grouped columns: ['h1-2-3.1', 'h1-2-3.2', 'h1-2-3.3']
```

Out[6]:

| | h1-2-3.1 | h1-2-3.2 | h1-2-3.3 |
|---|---|---|---|
| **0** | 1483 | 1515 | 1402 |
| **1** | 1129 | 691 | 725 |
| **2** | 2927 | 3523 | 3254 |
| **3** | 298 | 231 | 238 |

| | | | |
|---|---|---|---|
| [illegible] | [illegible] | [illegible] | [illegible] |
| **4** | 218 | 160 | 111 |
| **...** | ... | ... | ... |
| **4070** | 0 | 0 | 0 |
| **4071** | 0 | 0 | 0 |
| **4072** | 0 | 0 | 0 |
| **4073** | 0 | 0 | 0 |
| **4074** | 0 | 0 | 0 |

4075 rows × 3 columns

In [7]:
```python
# 'hFMT.1.2.3.gtKO' -->  'HM1->KO'
key_name = 'hFMT.1.2.3.gtKO'
print('Grouped columns: {}'.format(dict_metadata[key_name]))
HM1_KO = df_asv[[col_name for col_name in dict_metadata[key_name] ]]
HM1_KO
```

```
Grouped columns: ['F3-7', 'F3-8', 'F3-9', 'F3-10', 'F3-11', 'F3-12', 'F1-7',
'F1-8', 'F1-9', 'F1-10', 'F1-11', 'F1-12', 'F1-13', 'F1-14', 'F1-15']
```

Out[7]:

| | F3-7 | F3-8 | F3-9 | F3-10 | F3-11 | F3-12 | F1-7 | F1-8 | F1-9 | F1-10 | F1-11 | F1-12 | F1-13 | F |
|---|---|---|---|---|---|---|---|---|---|---|---|---|---|---|
| **0** | 10097 | 574 | 12959 | 18737 | 17857 | 12650 | 10738 | 9576 | 7304 | 14409 | 14382 | 8787 | 7694 | |
| **1** | 35102 | 51633 | 31840 | 22887 | 20446 | 33188 | 23565 | 21707 | 12802 | 26719 | 5701 | 15715 | 25125 | |
| **2** | 1328 | 91 | 2203 | 189 | 182 | 244 | 33299 | 13739 | 53077 | 17570 | 34094 | 18922 | 19204 | 4 |
| **3** | 137 | 315 | 305 | 134 | 221 | 290 | 203 | 348 | 363 | 238 | 219 | 205 | 255 | |
| **4** | 34960 | 13573 | 4109 | 3811 | 1155 | 28938 | 1144 | 6973 | 666 | 8230 | 1287 | 11346 | 13195 | [illegible] |
| **...** | ... | ... | ... | ... | ... | ... | ... | ... | ... | ... | ... | ... | ... | |
| **4070** | 0 | 0 | 0 | 0 | 0 | 0 | 0 | 0 | 0 | 0 | 0 | 0 | 0 | |
| **4071** | 0 | 0 | 0 | 0 | 0 | 0 | 0 | 0 | 0 | 0 | 0 | 0 | 0 | |
| **4072** | 0 | 0 | 0 | 0 | 0 | 0 | 0 | 0 | 0 | 0 | 0 | 0 | 0 | |
| **4073** | 0 | 0 | 0 | 0 | 0 | 0 | 0 | 0 | 0 | 0 | 0 | 0 | 0 | |
| **4074** | 0 | 0 | 0 | 0 | 0 | 0 | 0 | 0 | 0 | 0 | 0 | 0 | 0 | |

4075 rows × 15 columns

In [8]:
```python
# '1gKOinput' -->  'IMM-g1input'
key_name = '1gKOinput'
print('Grouped columns: {}'.format(dict_metadata[key_name]))
IMM_g1input = df_asv[[col_name for col_name in dict_metadata[key_name] ]]
IMM_g1input
```

```
Grouped columns: ['1gKO.1', '1gKO.2', '1gKO.3']
```

Out[8]:

| | 1gKO.1 | 1gKO.2 | 1gKO.3 |
|---|---|---|---|
| **0** | 13615 | 16637 | 17148 |
| **1** | 26439 | 30129 | 19743 |
| **2** | 7451 | 8774 | 8754 |
| **3** | 225 | 223 | 184 |
| **4** | 10108 | 12117 | 8633 |

| | | | |
|---|---|---|---|
| **4** | [illegible] | [illegible] | [illegible] |
| **...** | ... | ... | ... |
| **4070** | 0 | 0 | 0 |
| **4071** | 0 | 0 | 0 |
| **4072** | 0 | 0 | 0 |
| **4073** | 0 | 0 | 0 |
| **4074** | 0 | 0 | 0 |

4075 rows × 3 columns

In [9]:

```
# '1gKOgtKO' -->  'IMM-g1->KO'
key_name = '1gKOgtKO'
print('Grouped columns: {}'.format(dict_metadata[key_name]))
IMM_g1_KO = df_asv[[col_name for col_name in dict_metadata[key_name] ]]
IMM_g1_KO
```

```
Grouped columns: ['F8-1', 'F8-2', 'F8-3', 'F8-4', 'F8-5', 'F8-6', 'F8-7', 'F
8-8', 'F4-1', 'F4-2', 'F4-3', 'F4-4', 'F4-5', 'F3-1', 'F3-2', 'F3-4', 'F3-
5', 'F3-6']
```

Out[9]:

| | F8-1 | F8-2 | F8-3 | F8-4 | F8-5 | F8-6 | F8-7 | F8-8 | F4-1 | F4-2 | F4-3 | F4-4 | F4-5 | |
|---|---|---|---|---|---|---|---|---|---|---|---|---|---|---|
| **0** | 3339 | 15223 | 11423 | 18205 | 12486 | 10761 | 9339 | 466 | 29629 | 20134 | 27588 | 26741 | 25644 | 2 |
| **1** | 10283 | 6602 | 9314 | 13651 | 8789 | 16118 | 12045 | 13841 | 3687 | 6998 | 6904 | 7796 | 4011 | |
| **2** | 41532 | 40602 | 52337 | 11038 | 16781 | 8926 | 25758 | 31166 | 1627 | 37042 | 23825 | 26508 | 30376 | |
| **3** | 240 | 276 | 331 | 302 | 341 | 231 | 226 | 306 | 275 | 198 | 213 | 271 | 172 | 2 |
| **4** | 1806 | 6778 | 6725 | 12734 | 3901 | 10948 | 10347 | 1234 | 8891 | 3718 | 9188 | 4419 | 8315 | |
| **...** | ... | ... | ... | ... | ... | ... | ... | ... | ... | ... | ... | ... | ... | |
| **4070** | 0 | 0 | 0 | 0 | 0 | 0 | 0 | 0 | 0 | 0 | 0 | 0 | 0 | |
| **4071** | 0 | 0 | 0 | 0 | 0 | 0 | 0 | 0 | 0 | 0 | 0 | 0 | 0 | |
| **4072** | 0 | 0 | 0 | 0 | 0 | 0 | 0 | 0 | 0 | 0 | 0 | 0 | 0 | |
| **4073** | 0 | 0 | 0 | 0 | 0 | 0 | 0 | 0 | 0 | 0 | 0 | 0 | 0 | |
| **4074** | 0 | 0 | 0 | 0 | 0 | 0 | 0 | 0 | 0 | 0 | 0 | 0 | 0 | |

4075 rows × 18 columns

In [10]:

```
# '2gKOinput' -->  'IMM-g2input'
key_name = '2gKOinput'
print('Grouped columns: {}'.format(dict_metadata[key_name]))
IMM_g2input = df_asv[[col_name for col_name in dict_metadata[key_name] ]]
IMM_g2input
```

```
Grouped columns: ['2gKO.1', '2gKO.2', '2gKO.3']
```

Out[10]:

| | 2gKO.1 | 2gKO.2 | 2gKO.3 |
|---|---|---|---|
| **0** | 14832 | 14973 | 13992 |
| **1** | 18489 | 18178 | 9122 |
| **2** | 21958 | 22435 | 17629 |
| **3** | 269 | 153 | 230 |
| **4** | 5979 | 4360 | 2762 |

| | | | |
|---|---|---|---|
| 4 | [illegible] | [illegible] | [illegible] |
| ... | ... | ... | ... |
| 4070 | 0 | 0 | 0 |
| 4071 | 0 | 0 | 0 |
| 4072 | 0 | 0 | 0 |
| 4073 | 0 | 0 | 0 |
| 4074 | 0 | 0 | 0 |

4075 rows × 3 columns

In [11]:
```
# '2gKOgtKO' -->  'IMM-g2->KO'
key_name = '2gKOgtKO'
print('Grouped columns: {}'.format(dict_metadata[key_name]))
IMM_g2_KO = df_asv[[col_name for col_name in dict_metadata[key_name] ]]
IMM_g2_KO
```

```
Grouped columns: ['F8-9', 'F8-10', 'F8-11', 'F8-12', 'F8-13', 'F8-14', 'F8-1
5', 'F8-16']
```

Out[11]:

| | F8-9 | F8-10 | F8-11 | F8-12 | F8-13 | F8-14 | F8-15 | F8-16 |
|---|---|---|---|---|---|---|---|---|
| 0 | 23451 | 24742 | 31312 | 13542 | 4593 | 10228 | 14477 | 23819 |
| 1 | 8030 | 14203 | 27872 | 26539 | 12663 | 13720 | 23316 | 14430 |
| 2 | 19653 | 22616 | 4603 | 28535 | 20096 | 20133 | 14650 | 24062 |
| 3 | 266 | 344 | 300 | 269 | 257 | 330 | 249 | 226 |
| 4 | 6490 | 6143 | 12353 | 2229 | 6367 | 11645 | 9607 | 10622 |
| ... | ... | ... | ... | ... | ... | ... | ... | ... |
| 4070 | 0 | 0 | 0 | 0 | 0 | 0 | 0 | 0 |
| 4071 | 0 | 0 | 0 | 0 | 0 | 0 | 0 | 0 |
| 4072 | 0 | 0 | 0 | 0 | 0 | 0 | 0 | 0 |
| 4073 | 0 | 0 | 0 | 0 | 0 | 0 | 0 | 0 |
| 4074 | 0 | 0 | 0 | 0 | 0 | 0 | 0 | 0 |

4075 rows × 8 columns

### Step 6: Normalize count values for target sample groups

In [12]:
```
HM1_Input_norm = _get_norm_counts(HM1_Input, sample_count_sums)
HM1_Input_norm
```

Out[12]:

| | h1-2-3.1 | h1-2-3.2 | h1-2-3.3 |
|---|---|---|---|
| 0 | 3.170992 | 3.168281 | 3.139741 |
| 1 | 3.052636 | 2.827698 | 2.853624 |
| 2 | 3.466129 | 3.534613 | 3.505231 |
| 3 | 2.475230 | 2.353116 | 2.371106 |
| 4 | 2.340004 | 2.194478 | 2.041962 |

| | | | |
|---|---|---|---|
| ... | ... | ... | ... |
| **4070** | 0.000000 | 0.000000 | 0.000000 |
| **4071** | 0.000000 | 0.000000 | 0.000000 |
| **4072** | 0.000000 | 0.000000 | 0.000000 |
| **4073** | 0.000000 | 0.000000 | 0.000000 |
| **4074** | 0.000000 | 0.000000 | 0.000000 |

4075 rows × 3 columns

In [13]:
```
HM1_KO_norm = _get_norm_counts(HM1_KO, sample_count_sums)
HM1_KO_norm
```

Out[13]:

| | F3-7 | F3-8 | F3-9 | F3-10 | F3-11 | F3-12 | F1-7 | F1-8 | F1-9 | F1- |
|---|---|---|---|---|---|---|---|---|---|---|
| **0** | 4.023995 | 2.715632 | 4.028643 | 4.300323 | 4.281648 | 4.108629 | 4.026117 | 3.977907 | 3.888433 | 4.1297 |
| **1** | 4.565105 | 4.668820 | 4.419021 | 4.387208 | 4.340445 | 4.527498 | 4.367439 | 4.333297 | 4.132126 | 4.3978 |
| **2** | 3.143272 | 1.920176 | 3.259285 | 2.306214 | 2.292111 | 2.395644 | 4.517597 | 4.134664 | 4.749730 | 4.2158 |
| **3** | 2.159501 | 2.455717 | 2.402055 | 2.157737 | 2.376040 | 2.470375 | 2.304807 | 2.539512 | 2.585851 | 2.3495 |
| **4** | 4.563345 | 4.088595 | 3.529896 | 3.608747 | 3.092750 | 4.467988 | 3.053962 | 3.840159 | 2.848906 | 3.8865 |
| ... | ... | ... | ... | ... | ... | ... | ... | ... | ... | ... |
| **4070** | 0.000000 | 0.000000 | 0.000000 | 0.000000 | 0.000000 | 0.000000 | 0.000000 | 0.000000 | 0.000000 | 0.0000 |
| **4071** | 0.000000 | 0.000000 | 0.000000 | 0.000000 | 0.000000 | 0.000000 | 0.000000 | 0.000000 | 0.000000 | 0.0000 |
| **4072** | 0.000000 | 0.000000 | 0.000000 | 0.000000 | 0.000000 | 0.000000 | 0.000000 | 0.000000 | 0.000000 | 0.0000 |
| **4073** | 0.000000 | 0.000000 | 0.000000 | 0.000000 | 0.000000 | 0.000000 | 0.000000 | 0.000000 | 0.000000 | 0.0000 |
| **4074** | 0.000000 | 0.000000 | 0.000000 | 0.000000 | 0.000000 | 0.000000 | 0.000000 | 0.000000 | 0.000000 | 0.0000 |

4075 rows × 15 columns

In [14]:
```
IMM_g1input_norm = _get_norm_counts(IMM_g1input, sample_count_sums)
IMM_g1input_norm
```

Out[14]:

| | 1gKO.1 | 1gKO.2 | 1gKO.3 |
|---|---|---|---|
| **0** | 4.180178 | 4.188104 | 4.262257 |
| **1** | 4.468392 | 4.446001 | 4.323454 |
| **2** | 3.918399 | 3.910251 | 3.970273 |
| **3** | 2.400047 | 2.317401 | 2.295045 |
| **4** | 4.050836 | 4.050434 | 3.964229 |
| ... | ... | ... | ... |
| **4070** | 0.000000 | 0.000000 | 0.000000 |
| **4071** | 0.000000 | 0.000000 | 0.000000 |
| **4072** | 0.000000 | 0.000000 | 0.000000 |
| **4073** | 0.000000 | 0.000000 | 0.000000 |
| **4074** | 0.000000 | 0.000000 | 0.000000 |

4075 rows × 3 columns

In [15]:
```
IMM_g1_KO_norm = _get_norm_counts(IMM_g1_KO, sample_count_sums)
IMM_g1_KO_norm
```

Out[15]:

| | F8-1 | F8-2 | F8-3 | F8-4 | F8-5 | F8-6 | F8-7 | F8-8 | F4-1 | F4 |
|---|---|---|---|---|---|---|---|---|---|---|
| **0** | 3.530045 | 4.171099 | 4.067413 | 4.239122 | 4.320109 | 4.065412 | 3.978399 | 2.753708 | 4.509011 | 4.2994 |
| **1** | 4.018461 | 3.808313 | 3.978778 | 4.114104 | 4.167634 | 4.240858 | 4.088895 | 4.225749 | 3.604062 | 3.8405 |
| **2** | 4.624693 | 4.597128 | 4.728413 | 4.021838 | 4.448499 | 3.984224 | 4.418981 | 4.578248 | 3.248913 | 4.5642 |
| **3** | 2.388291 | 2.431092 | 2.530706 | 2.460420 | 2.757180 | 2.398871 | 2.364043 | 2.571443 | 2.478060 | 2.2944 |
| **4** | 3.263255 | 3.819737 | 3.837351 | 4.083907 | 3.814908 | 4.072893 | 4.022908 | 3.176160 | 3.986276 | 3.5659 |
| **...** | ... | ... | ... | ... | ... | ... | ... | ... | ... | |
| **4070** | 0.000000 | 0.000000 | 0.000000 | 0.000000 | 0.000000 | 0.000000 | 0.000000 | 0.000000 | 0.000000 | 0.0000 |
| **4071** | 0.000000 | 0.000000 | 0.000000 | 0.000000 | 0.000000 | 0.000000 | 0.000000 | 0.000000 | 0.000000 | 0.0000 |
| **4072** | 0.000000 | 0.000000 | 0.000000 | 0.000000 | 0.000000 | 0.000000 | 0.000000 | 0.000000 | 0.000000 | 0.0000 |
| **4073** | 0.000000 | 0.000000 | 0.000000 | 0.000000 | 0.000000 | 0.000000 | 0.000000 | 0.000000 | 0.000000 | 0.0000 |
| **4074** | 0.000000 | 0.000000 | 0.000000 | 0.000000 | 0.000000 | 0.000000 | 0.000000 | 0.000000 | 0.000000 | 0.0000 |

4075 rows × 18 columns

In [16]:
```
IMM_g2input_norm = _get_norm_counts(IMM_g2input, sample_count_sums)
IMM_g2input_norm
```

Out[16]:

| | 2gKO.1 | 2gKO.2 | 2gKO.3 |
|---|---|---|---|
| **0** | 4.143317 | 4.154522 | 4.242762 |
| **1** | 4.239025 | 4.238754 | 4.056985 |
| **2** | 4.313700 | 4.330129 | 4.343104 |
| **3** | 2.403557 | 2.166842 | 2.460093 |
| **4** | 3.748792 | 3.618774 | 3.538206 |
| **...** | ... | ... | ... |
| **4070** | 0.000000 | 0.000000 | 0.000000 |
| **4071** | 0.000000 | 0.000000 | 0.000000 |
| **4072** | 0.000000 | 0.000000 | 0.000000 |
| **4073** | 0.000000 | 0.000000 | 0.000000 |
| **4074** | 0.000000 | 0.000000 | 0.000000 |

4075 rows × 3 columns

In [17]:
```
IMM_g2_KO_norm = _get_norm_counts(IMM_g2_KO, sample_count_sums)
IMM_g2_KO_norm
```

Out[17]:

| | F8-9 | F8-10 | F8-11 | F8-12 | F8-13 | F8-14 | F8-15 | F8-16 |
|---|---|---|---|---|---|---|---|---|
| **0** | 4.480880 | 4.378895 | 4.379459 | 4.143025 | 3.698748 | 4.105404 | 4.195335 | 4.353703 |
| **1** | 4.015462 | 4.137854 | 4.328918 | 4.435212 | 4.139133 | 4.232959 | 4.402300 | 4.136058 |
| **2** | 4.404150 | 4.339877 | 3.546894 | 4.466704 | 4.339694 | 4.399505 | 4.200494 | 4.358111 |

| | | | | | | | | |
|---|---|---|---|---|---|---|---|---|
| **3** | 2.536849 | 2.523304 | 2.362739 | 2.442634 | 2.448049 | 2.615148 | 2.432436 | 2.332891 |
| **4** | 3.923001 | 3.773895 | 3.975548 | 3.359611 | 3.840562 | 4.161749 | 4.017259 | 4.003009 |
| **...** | ... | ... | ... | ... | ... | ... | ... | ... |
| **4070** | 0.000000 | 0.000000 | 0.000000 | 0.000000 | 0.000000 | 0.000000 | 0.000000 | 0.000000 |
| **4071** | 0.000000 | 0.000000 | 0.000000 | 0.000000 | 0.000000 | 0.000000 | 0.000000 | 0.000000 |
| **4072** | 0.000000 | 0.000000 | 0.000000 | 0.000000 | 0.000000 | 0.000000 | 0.000000 | 0.000000 |
| **4073** | 0.000000 | 0.000000 | 0.000000 | 0.000000 | 0.000000 | 0.000000 | 0.000000 | 0.000000 |
| **4074** | 0.000000 | 0.000000 | 0.000000 | 0.000000 | 0.000000 | 0.000000 | 0.000000 | 0.000000 |

4075 rows × 8 columns

#### Step 7: Calculate Spearman Correlation Coefficients for each sample group

In [18]:

```
corr_value1 = get_inter_corr_values(HM1_Input_norm, HM1_KO_norm)
corr_value2 = get_inter_corr_values(IMM_g1input_norm, IMM_g1_KO_norm)
corr_value3 = get_inter_corr_values(IMM_g2input_norm, IMM_g2_KO_norm)
```

#### Step 8: Generate final figure

In [19]:

```
data = [corr_value1, corr_value2, corr_value3]
x_tick_labels = ['HM1Input\nvs HM1->KO',
                 'IMM-g1input\nvs IMM-g1->KO',
                 'IMM-g2input\nvs IMM-g2->KO']

fig, ax = plt.subplots()
ax.boxplot(data, widths=0.8)
ax.set_title('Regenerated Figure 4D')
ax.set_yticks([0.0, 0.2, 0.4, 0.6, 0.8, 1.0])
ax.set_xticklabels(x_tick_labels,
                   rotation=45, fontsize=10, weight='bold')

plt.show()
```

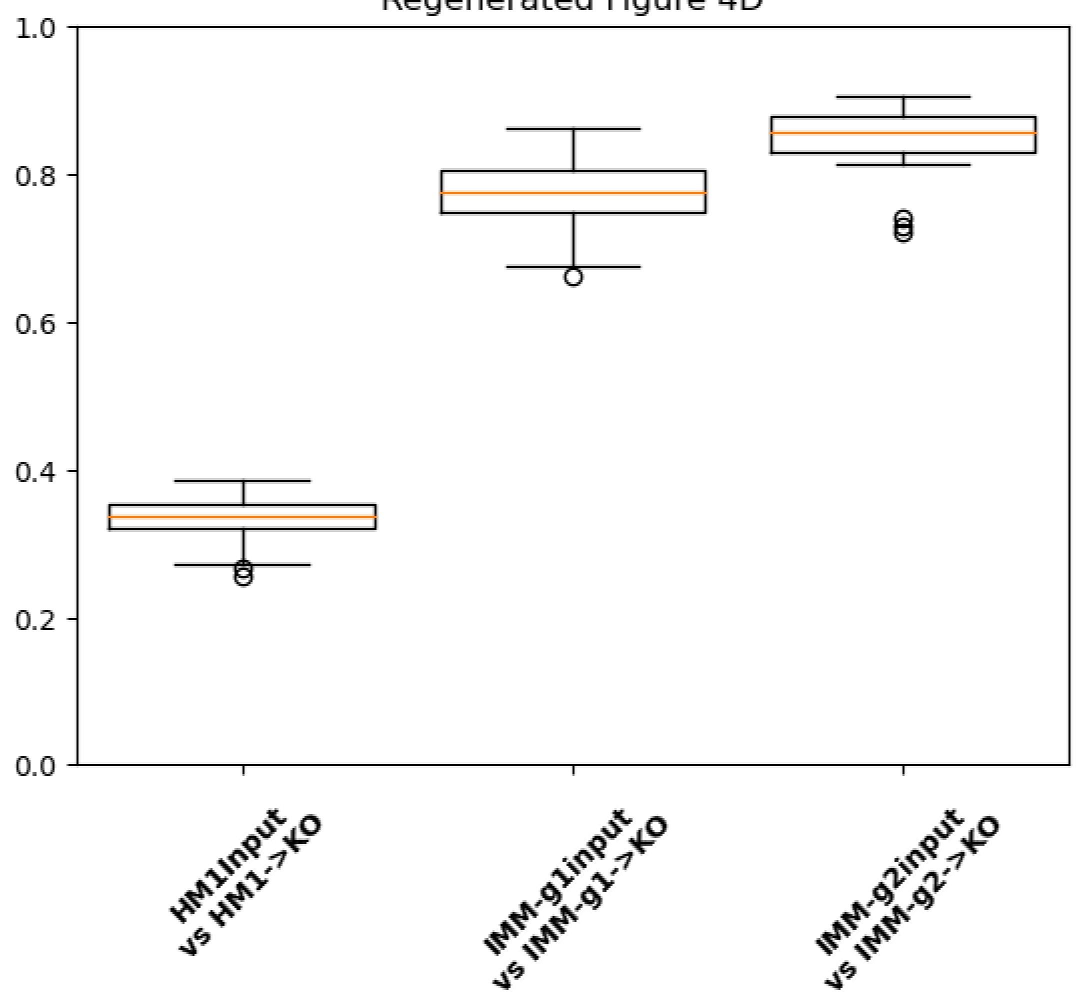


#### Step 9: Compare original and regenerated figures

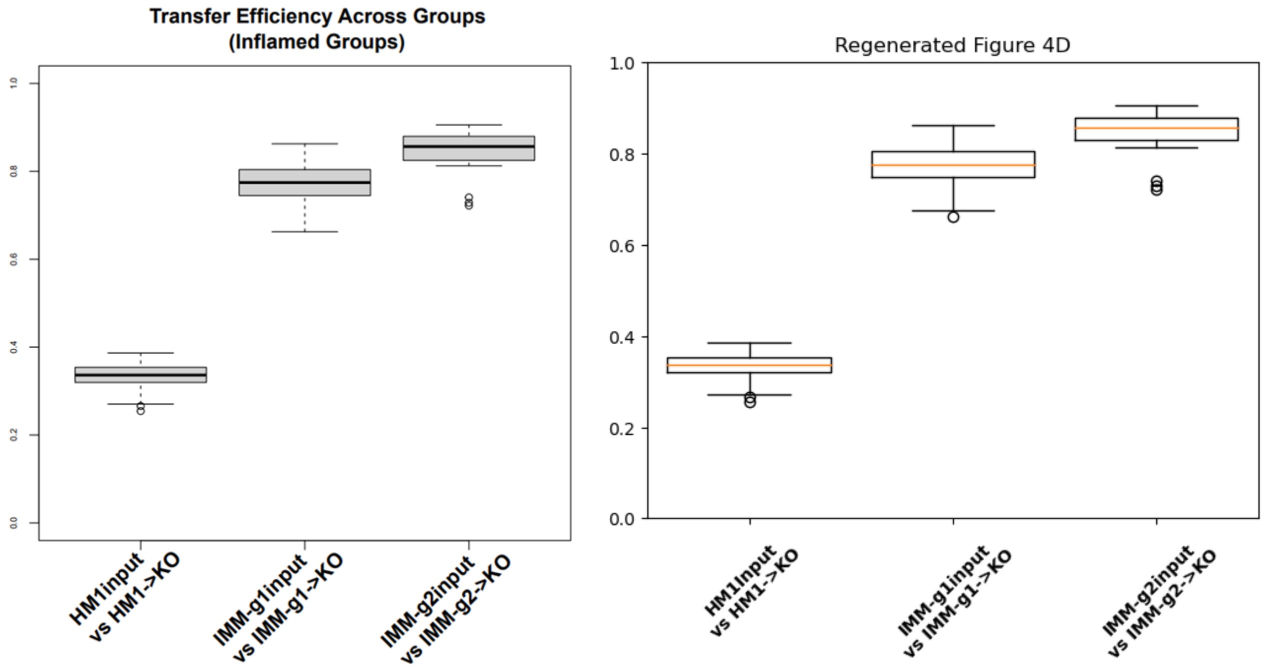

### Contents


- **Aim**
- **Main Python Methods**
- **Data Processing Steps**
  - Step 1: Retrieve original counts table
  - Step 2: Retrieve original metadata table
  - Step 3: Create dictionary from metadata table
  - Step 4: Calculate total read counts per sample
  - Step 5: Extract counts for sample group HM1->KO
  - Step 6: Normalize count values for sample group HM1->KOO
  - Step 7: Calculate Spearman Correlation Coefficients for each sample pair
  - Step 8: Generate final figure
  - Step 9: Compare original and regenerated figures


### Aim: Replicate Figure 5D

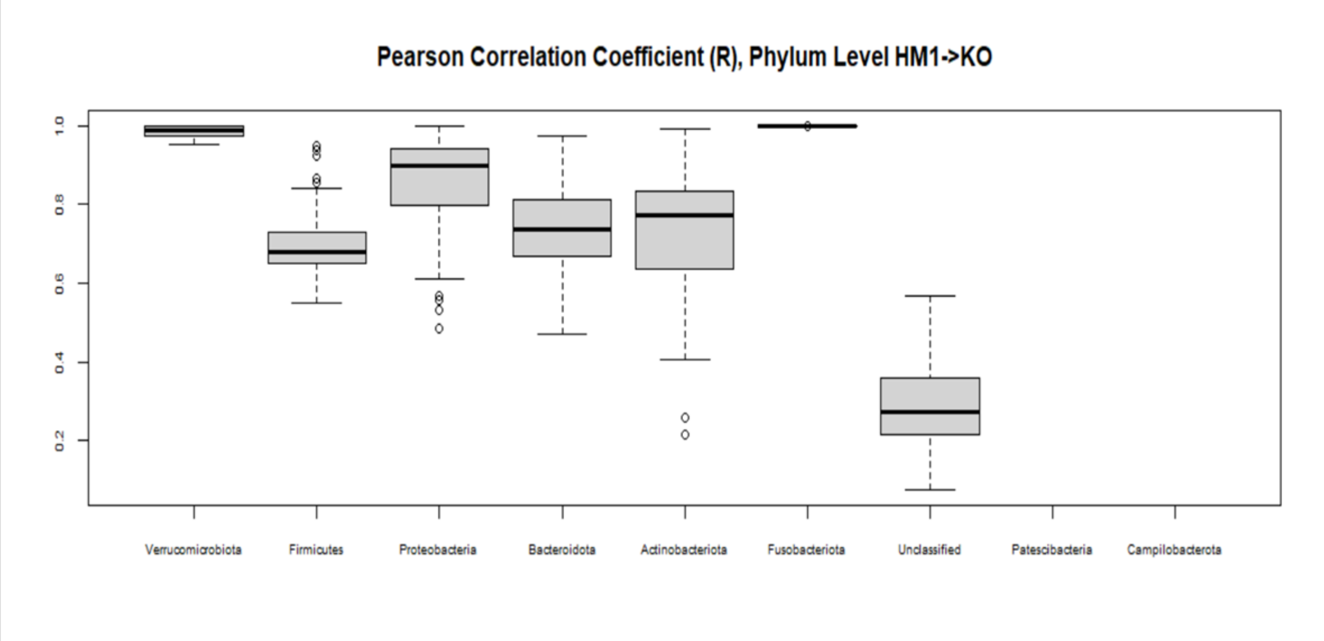


### Main Python Methods

In [1]:

```
import numpy as np
import pandas as pd
import matplotlib.pyplot as plt
import os
import re
from scipy import stats
from scipy.special import comb

# ******************************* MAIN METHODS *******************************
# Filter rows for target taxa
def filter_rows(taxa_query, taxa_target):
    final_result = False
```

```python
    search_result = re.search(taxa_target, taxa_query)
    if search_result:
        final_result = True

    return final_result
# Calculate Pearson correlation coefficients from input set of samples
def get_inter_corr_values(input_df):
    tup_list_column_names = [(col1, col2) for col1 in input_df.columns for col
2 in input_df.columns if input_df.columns.get_loc(col2) > input_df.columns.get
_loc(col1)]
    corr_values = [stats.pearsonr(input_df.loc[:, tup[0]], input_df.loc[:, tup
[1]]).statistic \
            for tup in tup_list_column_names]
    # Remove nan values
    corr_values = [val for val in corr_values if not np.isnan(val)]
    return corr_values

# Retrieve tab delimited file
def read_csv_file(file_path, skiprows=None, header = 0, sep = '\t', index_col=
False):
    df = pd.read_csv(file_path, skiprows=skiprows, sep=sep, header=header, ind
ex_col=index_col)

    return df

# Create dictionary from metadata table
def get_dict_from_metadata(input_df):
    mydict = {}
    for row in input_df.iterrows():
        obj = row[1]
        sample_id = obj['SampleID']
        key = obj['FMTGroupFMTsourcegtRecipientbackground']
        if key not in mydict:
            mydict[key] = [sample_id]
        else:
            mydict[key].append(sample_id)
    return mydict

# ******************************* HELPER METHODS *****************************
**
# Normalize sample count values
def _get_norm_counts(input_df, ser_sample_count_sums):
    overall_mean_count = ser_sample_count_sums.mean()
    df_norm_logged =    pd.DataFrame()
    for col_name in input_df.columns:
        df_norm_logged.loc[:, col_name] = \
        np.log10(((input_df.loc[:, col_name] / ser_sample_count_sums[col_name
]) * overall_mean_count) + 1)

    return df_norm_logged

# ******************************* INPUT FILE PATHS ***************************
****
# Get current working directory
current_working_dir = os.getcwd()
# original counts table
asv_tbl_file_path = os.path.join(current_working_dir, 'asv_biom-with-taxonomy.
txt')
# original metadata table
metadata_file_path = os.path.join(current_working_dir, 'mappingMetadata.txt')
```

### Data Processing Steps

### Step 1: Retrieve original counts table

In [2]:
```
df_asv = read_csv_file(asv_tbl_file_path, 1)
df_asv
```

Out[2]:

| | #OTU ID | 1gKO.1 | 1gKO.2 | 1gKO.3 | 1gWT.1 | 1gWT.2 | 1gWT.3 | 2gKO.1 |
|---|---|---|---|---|---|---|---|---|
| **0** | 1ba8c796d07406783c96d016a6a5cace | 13615.0 | 16637.0 | 17148.0 | 20227.0 | 23630.0 | 25656.0 | 14832.0 |
| **1** | a6c38249aff7768283faf6cfbdeb05a8 | 26439.0 | 30129.0 | 19743.0 | 8955.0 | 10759.0 | 7074.0 | 18489.0 |
| **2** | 062f38ff92cfaee0654200b6f5be5ddf | 7451.0 | 8774.0 | 8754.0 | 174.0 | 214.0 | 148.0 | 21958.0 |
| **3** | 1183cc23f552d81e63c93ca9fcba2f2c | 225.0 | 223.0 | 184.0 | 13762.0 | 16856.0 | 18692.0 | 269.0 |
| **4** | 5e15ecfb579e72bf87c0bea3920bbf42 | 10108.0 | 12117.0 | 8633.0 | 10027.0 | 11910.0 | 7424.0 | 5979.0 |
| **...** | ... | ... | ... | ... | ... | ... | ... | ... |
| **4070** | 92bb8f4683ef5c8651e7d34dbb37ab2e | 0.0 | 0.0 | 0.0 | 0.0 | 0.0 | 0.0 | 0.0 |
| **4071** | 92f09070a4fd5786bb34e756217e6ee1 | 0.0 | 0.0 | 0.0 | 0.0 | 0.0 | 0.0 | 0.0 |
| **4072** | 919b82324c41ed0046323c63aa1550da | 0.0 | 0.0 | 0.0 | 0.0 | 0.0 | 0.0 | 0.0 |
| **4073** | dbc0dad15ec1c8ad9d826cab94e18696 | 0.0 | 0.0 | 0.0 | 0.0 | 0.0 | 0.0 | 0.0 |
| **4074** | 1ff2d07d10264c23dc43e08d3097cd7c | 0.0 | 0.0 | 0.0 | 0.0 | 0.0 | 0.0 | 0.0 |

4075 rows × 112 columns

In [3]:
```
df_asv = read_csv_file(asv_tbl_file_path, 1)
df_asv = df_asv.astype({col:'int32' for col in df_asv.columns[1:-1] }, copy=Fa
lse)
df_asv
```

Out[3]:

| | #OTU ID | 1gKO.1 | 1gKO.2 | 1gKO.3 | 1gWT.1 | 1gWT.2 | 1gWT.3 | 2gKO.1 |
|---|---|---|---|---|---|---|---|---|
| **0** | 1ba8c796d07406783c96d016a6a5cace | 13615 | 16637 | 17148 | 20227 | 23630 | 25656 | 14832 |
| **1** | a6c38249aff7768283faf6cfbdeb05a8 | 26439 | 30129 | 19743 | 8955 | 10759 | 7074 | 18489 |
| **2** | 062f38ff92cfaee0654200b6f5be5ddf | 7451 | 8774 | 8754 | 174 | 214 | 148 | 21958 |

| | | | | | | | | |
|---|---|---|---|---|---|---|---|---|
| **3** | 1183cc23f552d81e63c93ca9fcba2f2c | 225 | 223 | 184 | 13762 | 16856 | 18692 | 269 |
| **4** | 5e15ecfb579e72bf87c0bea3920bbf42 | 10108 | 12117 | 8633 | 10027 | 11910 | 7424 | 5979 |
| **...** | ... | ... | ... | ... | ... | ... | ... | ... |
| **4070** | 92bb8f4683ef5c8651e7d34dbb37ab2e | 0 | 0 | 0 | 0 | 0 | 0 | 0 |
| **4071** | 92f09070a4fd5786bb34e756217e6ee1 | 0 | 0 | 0 | 0 | 0 | 0 | 0 |
| **4072** | 919b82324c41ed0046323c63aa1550da | 0 | 0 | 0 | 0 | 0 | 0 | 0 |
| **4073** | dbc0dad15ec1c8ad9d826cab94e18696 | 0 | 0 | 0 | 0 | 0 | 0 | 0 |
| **4074** | 1ff2d07d10264c23dc43e08d3097cd7c | 0 | 0 | 0 | 0 | 0 | 0 | 0 |

4075 rows × 112 columns

### Step 2: Retrieve original metadata table

In [4]:

```
df_metadata = read_csv_file(metadata_file_path)
df_metadata
```

Out[4]:

| | SampleID | UniversalCageNumber | Background | FMTGroupFMTsourcegtRecipientbackground | Passage |
|---|---|---|---|---|---|
| **0** | F8-1 | F8-cage-1 | 129.IL10KO | 1gKOgtKO | 8 |
| **1** | F8-2 | F8-cage-1 | 129.IL10KO | 1gKOgtKO | 8 |
| **2** | F8-3 | F8-cage-2 | 129.IL10KO | 1gKOgtKO | 8 |
| **3** | F8-4 | F8-cage-2 | 129.IL10KO | 1gKOgtKO | 8 |
| **4** | F8-5 | F8-cage-3 | 129.IL10KO | 1gKOgtKO | 8 |
| **...** | ... | ... | ... | ... | ... |
| **105** | 1gWT.2 | NaN | NaN | 1gWTinput | 1gWT |
| **106** | 1gWT.3 | NaN | NaN | 1gWTinput | 1gWT |
| **107** | 2gWT.1 | NaN | NaN | 2gWTinput | 2gWT |
| **108** | 2gWT.2 | NaN | NaN | 2gWTinput | 2gWT |
| **109** | 2gWT.3 | NaN | NaN | 2gWTinput | 2gWT |

110 rows × 8 columns

### Step 3: Create dictionary from metadata table

In [5]:

```
dict_metadata = get_dict_from_metadata(df_metadata)
dict_metadata.keys()
```

Out[5]:

```
dict_keys(['1gKOgtKO', '2gKOgtKO', '1gWTgtKO', '1gWTgtWT', '2gWTgtWT', 'hFM
T.1.2.3.gtKO', 'hFMT.3.4.5.gtKO', 'hFMT.1.2.3.gtWT', 'hFMT.1.2.3.input', 'hF
MT.3.4.5.input', '1gKOinput', '2gKOinput', '1gWTinput', '2gWTinput'])
```

### Step 4: Calculate total read counts per sample

In [6]:
```
sample_count_sums = df_asv.iloc[:, 1:-1].sum(axis=0)
sample_count_sums
```

Out[6]:
```
1gKO.1      118256
1gKO.2      141891
1gKO.3      123292
1gWT.1      119717
1gWT.2      146158
             ...
h1-2-3.2    135326
h1-2-3.3    133745
h3-4-5.1    129613
h3-4-5.2    140316
h3-4-5.3    132984
Length: 110, dtype: int64
```

### Step 5: Extract counts for sample group HM1->KO

In [7]:
```
# 'hFMT.1.2.3.gtKO' -->  'HM1->KO'
key_name = 'hFMT.1.2.3.gtKO'
group_columns = dict_metadata[key_name]
group_columns.extend(['#OTU ID', 'taxonomy'])
key_name = 'hFMT.1.2.3.gtKO'
HM1_KO = df_asv[[col_name for col_name in group_columns ]]
HM1_KO
```

Out[7]:

| | F3-7 | F3-8 | F3-9 | F3-10 | F3-11 | F3-12 | F1-7 | F1-8 | F1-9 | F1-10 | F1-11 | F1-12 | F1-13 | F |
|---|---|---|---|---|---|---|---|---|---|---|---|---|---|---|
| **0** | 10097 | 574 | 12959 | 18737 | 17857 | 12650 | 10738 | 9576 | 7304 | 14409 | 14382 | 8787 | 7694 | |
| **1** | 35102 | 51633 | 31840 | 22887 | 20446 | 33188 | 23565 | 21707 | 12802 | 26719 | 5701 | 15715 | 25125 | |
| **2** | 1328 | 91 | 2203 | 189 | 182 | 244 | 33299 | 13739 | 53077 | 17570 | 34094 | 18922 | 19204 | 4 |
| **3** | 137 | 315 | 305 | 134 | 221 | 290 | 203 | 348 | 363 | 238 | 219 | 205 | 255 | |
| **4** | 34960 | 13573 | 4109 | 3811 | 1155 | 28938 | 1144 | 6973 | 666 | 8230 | 1287 | 11346 | 13195 | [illegible] |
| **...** | ... | ... | ... | ... | ... | ... | ... | ... | ... | ... | ... | ... | ... | |
| **4070** | 0 | 0 | 0 | 0 | 0 | 0 | 0 | 0 | 0 | 0 | 0 | 0 | 0 | |
| **4071** | 0 | 0 | 0 | 0 | 0 | 0 | 0 | 0 | 0 | 0 | 0 | 0 | 0 | |
| **4072** | 0 | 0 | 0 | 0 | 0 | 0 | 0 | 0 | 0 | 0 | 0 | 0 | 0 | |
| **4073** | 0 | 0 | 0 | 0 | 0 | 0 | 0 | 0 | 0 | 0 | 0 | 0 | 0 | |
| **4074** | 0 | 0 | 0 | 0 | 0 | 0 | 0 | 0 | 0 | 0 | 0 | 0 | 0 | |

4075 rows × 17 columns

In [8]:
```
target_phylum = 'p__Verrucomicrobiota'
df_Verrucomicrobiota = HM1_KO[HM1_KO.taxonomy.apply(lambda x: filter_rows(x, t
arget_phylum))]
print('Number of rows: {}'.format(len(df_Verrucomicrobiota.index)))
print('Columns: {}'.format(', '.join(df_Verrucomicrobiota.columns)))
```

```
Number of rows: 24
Columns: F3-7, F3-8, F3-9, F3-10, F3-11, F3-12, F1-7, F1-8, F1-9, F1-10, F1-
11, F1-12, F1-13, F1-14, F1-15, #OTU ID, taxonomy
```

In [9]:
```
target_phylum = 'p__Firmicutes'
df_Firmicutes = HM1_KO[HM1_KO.taxonomy.apply(lambda x: filter_rows(x, target_p
hylum))]
print('Number of rows: {}'.format(len(df_Firmicutes.index)))
print('Columns: {}'.format(', '.join(df_Firmicutes.columns)))
```

```
Number of rows: 830
Columns: F3-7, F3-8, F3-9, F3-10, F3-11, F3-12, F1-7, F1-8, F1-9, F1-10, F1-
11, F1-12, F1-13, F1-14, F1-15, #OTU ID, taxonomy
```

In [10]:
```
target_phylum = 'p__Proteobacteria'
df_Proteobacteria = HM1_KO[HM1_KO.taxonomy.apply(lambda x: filter_rows(x, targ
et_phylum))]
print('Number of rows: {}'.format(len(df_Proteobacteria.index)))
print('Columns: {}'.format(', '.join(df_Proteobacteria.columns)))
```

```
Number of rows: 51
Columns: F3-7, F3-8, F3-9, F3-10, F3-11, F3-12, F1-7, F1-8, F1-9, F1-10, F1-
11, F1-12, F1-13, F1-14, F1-15, #OTU ID, taxonomy
```

In [11]:
```
target_phylum = 'p__Bacteroidota'
df_Bacteroidota = HM1_KO[HM1_KO.taxonomy.apply(lambda x: filter_rows(x, target
_phylum))]
print('Number of rows: {}'.format(len(df_Bacteroidota.index)))
print('Columns: {}'.format(', '.join(df_Bacteroidota.columns)))
```

```
Number of rows: 91
Columns: F3-7, F3-8, F3-9, F3-10, F3-11, F3-12, F1-7, F1-8, F1-9, F1-10, F1-
11, F1-12, F1-13, F1-14, F1-15, #OTU ID, taxonomy
```

In [12]:
```
target_phylum = 'p__Actinobacteriota'
df_Actinobacteriota = HM1_KO[HM1_KO.taxonomy.apply(lambda x: filter_rows(x, ta
rget_phylum))]
print('Number of rows: {}'.format(len(df_Actinobacteriota.index)))
print('Columns: {}'.format(', '.join(df_Actinobacteriota.columns)))
```

```
Number of rows: 71
Columns: F3-7, F3-8, F3-9, F3-10, F3-11, F3-12, F1-7, F1-8, F1-9, F1-10, F1-
11, F1-12, F1-13, F1-14, F1-15, #OTU ID, taxonomy
```

In [13]:
```
target_phylum = 'p__Fusobacteriota'
df_Fusobacteriota = HM1_KO[HM1_KO.taxonomy.apply(lambda x: filter_rows(x, targ
et_phylum))]
print('Number of rows: {}'.format(len(df_Fusobacteriota.index)))
print('Columns: {}'.format(', '.join(df_Fusobacteriota.columns)))
```

```
Number of rows: 11
Columns: F3-7, F3-8, F3-9, F3-10, F3-11, F3-12, F1-7, F1-8, F1-9, F1-10, F1-
11, F1-12, F1-13, F1-14, F1-15, #OTU ID, taxonomy
```

In [14]:
```
target_phylum = 'p__Patescibacteria'
df_Patescibacteria = HM1_KO[HM1_KO.taxonomy.apply(lambda x: filter_rows(x, tar
get_phylum))]
print('Number of rows: {}'.format(len(df_Patescibacteria.index)))
print('Columns: {}'.format(', '.join(df_Patescibacteria.columns)))
```

```
Number of rows: 3
Columns: F3-7, F3-8, F3-9, F3-10, F3-11, F3-12, F1-7, F1-8, F1-9, F1-10, F1-
11, F1-12, F1-13, F1-14, F1-15, #OTU ID, taxonomy
```

In [15]:
```
target_phylum = 'p__Campilobacterota'
df_Campilobacterota = HM1_KO[HM1_KO.taxonomy.apply(lambda x: filter_rows(x, ta
rget_phylum))]
print('Number of rows: {}'.format(len(df_Campilobacterota.index)))
print('Columns: {}'.format(', '.join(df_Campilobacterota.columns)))
```

```
Number of rows: 3
Columns: F3-7, F3-8, F3-9, F3-10, F3-11, F3-12, F1-7, F1-8, F1-9, F1-10, F1-
11, F1-12, F1-13, F1-14, F1-15, #OTU ID, taxonomy
```

In [16]:
```
target_phylum = 'Unassigned'
df_Unassigned = HM1_KO[HM1_KO.taxonomy.apply(lambda x: filter_rows(x, target_p
hylum))]
print('Number of rows: {:,}'.format(len(df_Unassigned.index)))
print('Columns: {}'.format(', '.join(df_Unassigned.columns)))
```

```
Number of rows: 2,196
Columns: F3-7, F3-8, F3-9, F3-10, F3-11, F3-12, F1-7, F1-8, F1-9, F1-10, F1-
11, F1-12, F1-13, F1-14, F1-15, #OTU ID, taxonomy
```

### Step 6: Normalize count values for sample group HM1->KO

In [17]:
```
df_Verrucomicrobiota_norm = _get_norm_counts(df_Verrucomicrobiota.iloc[:, :-2
], sample_count_sums)
print('Number of rows: {}'.format(len(df_Verrucomicrobiota_norm.index)))
print('Columns: {}'.format(', '.join(df_Verrucomicrobiota_norm.columns)))
```

```
Number of rows: 24
Columns: F3-7, F3-8, F3-9, F3-10, F3-11, F3-12, F1-7, F1-8, F1-9, F1-10, F1-
11, F1-12, F1-13, F1-14, F1-15
```

In [18]:
```
df_Firmicutes_norm = _get_norm_counts(df_Firmicutes.iloc[:, :-2], sample_count
_sums)
print('Number of rows: {}'.format(len(df_Firmicutes_norm.index)))
print('Columns: {}'.format(', '.join(df_Firmicutes_norm.columns)))
```

```
Number of rows: 830
Columns: F3-7, F3-8, F3-9, F3-10, F3-11, F3-12, F1-7, F1-8, F1-9, F1-10, F1-
11, F1-12, F1-13, F1-14, F1-15
```

In [19]:
```
df_Proteobacteria_norm = _get_norm_counts(df_Proteobacteria.iloc[:, :-2], samp
le_count_sums)
```

```
print('Number of rows: {}'.format(len(df_Proteobacteria_norm.index)))
print('Columns: {}'.format(', '.join(df_Proteobacteria_norm.columns)))
```

```
Number of rows: 51
Columns: F3-7, F3-8, F3-9, F3-10, F3-11, F3-12, F1-7, F1-8, F1-9, F1-10, F1-
11, F1-12, F1-13, F1-14, F1-15
```

In [20]:
```
df_Bacteroidota_norm = _get_norm_counts(df_Bacteroidota.iloc[:, :-2], sample_c
ount_sums)
print('Number of rows: {}'.format(len(df_Bacteroidota_norm.index)))
print('Columns: {}'.format(', '.join(df_Bacteroidota_norm.columns)))
```

```
Number of rows: 91
Columns: F3-7, F3-8, F3-9, F3-10, F3-11, F3-12, F1-7, F1-8, F1-9, F1-10, F1-
11, F1-12, F1-13, F1-14, F1-15
```

In [21]:
```
df_Actinobacteriota_norm = _get_norm_counts(df_Actinobacteriota.iloc[:, :-2],
sample_count_sums)
print('Number of rows: {}'.format(len(df_Actinobacteriota_norm.index)))
print('Columns: {}'.format(', '.join(df_Actinobacteriota_norm.columns)))
```

```
Number of rows: 71
Columns: F3-7, F3-8, F3-9, F3-10, F3-11, F3-12, F1-7, F1-8, F1-9, F1-10, F1-
11, F1-12, F1-13, F1-14, F1-15
```

In [22]:
```
df_Fusobacteriota_norm = _get_norm_counts(df_Fusobacteriota.iloc[:, :-2], samp
le_count_sums)
print('Number of rows: {}'.format(len(df_Fusobacteriota_norm.index)))
print('Columns: {}'.format(', '.join(df_Fusobacteriota_norm.columns)))
```

```
Number of rows: 11
Columns: F3-7, F3-8, F3-9, F3-10, F3-11, F3-12, F1-7, F1-8, F1-9, F1-10, F1-
11, F1-12, F1-13, F1-14, F1-15
```

In [23]:
```
df_Unassigned_norm = _get_norm_counts(df_Unassigned.iloc[:, :-2], sample_count
_sums)
print('Number of rows: {:,}'.format(len(df_Unassigned_norm.index)))
print('Columns: {}'.format(', '.join(df_Unassigned_norm.columns)))
```

```
Number of rows: 2,196
Columns: F3-7, F3-8, F3-9, F3-10, F3-11, F3-12, F1-7, F1-8, F1-9, F1-10, F1-
11, F1-12, F1-13, F1-14, F1-15
```

In [24]:
```
df_Patescibacteria_norm = _get_norm_counts(df_Patescibacteria.iloc[:, :-2], sa
mple_count_sums)
print('Number of rows: {:,}'.format(len(df_Patescibacteria_norm.index)))
print('Columns: {}'.format(', '.join(df_Patescibacteria_norm.columns)))
```

```
Number of rows: 3
Columns: F3-7, F3-8, F3-9, F3-10, F3-11, F3-12, F1-7, F1-8, F1-9, F1-10, F1-
11, F1-12, F1-13, F1-14, F1-15
```

In [25]:
```
df_Campilobacterota_norm = _get_norm_counts(df_Campilobacterota.iloc[:, :-2],
sample_count_sums)
print('Number of rows: {:,}'.format(len(df_Campilobacterota_norm.index)))
print('Columns: {}'.format(', '.join(df_Campilobacterota_norm.columns)))
```

```
Number of rows: 3

Columns: F3-7, F3-8, F3-9, F3-10, F3-11, F3-12, F1-7, F1-8, F1-9, F1-10, F1-
11, F1-12, F1-13, F1-14, F1-15
```

### Step 7: Calculate Spearman Correlation Coefficients for each sample pair

In [26]:

```
corr_Verrucomicrobiota_norm = get_inter_corr_values(df_Verrucomicrobiota_norm)
norm_Firmicutes_norm = get_inter_corr_values(df_Firmicutes_norm)
corr_Proteobacteria_norm = get_inter_corr_values(df_Proteobacteria_norm)
corr_Bacteroidota_norm = get_inter_corr_values(df_Bacteroidota_norm)
corr_Actinobacteriota_norm = get_inter_corr_values(df_Actinobacteriota_norm)
corr_Fusobacteriota_norm = get_inter_corr_values(df_Fusobacteriota_norm)
corr_Unassigned_norm = get_inter_corr_values(df_Unassigned_norm)
corr_Patescibacteria_norm = get_inter_corr_values(df_Patescibacteria_norm)
corr_Campilobacterota_norm = get_inter_corr_values(df_Campilobacterota_norm)
```

```
C:\ProgramData\Anaconda3\lib\site-packages\scipy\stats\_stats_py.py:4424: Co
nstantInputWarning: An input array is constant; the correlation coefficient
is not defined.
  warnings.warn(stats.ConstantInputWarning(msg))
```

### Step 8: Generate final figure

In [27]:

```
data = [corr_Verrucomicrobiota_norm,
            norm_Firmicutes_norm,
            corr_Proteobacteria_norm,
            corr_Bacteroidota_norm,
            corr_Actinobacteriota_norm,
            corr_Fusobacteriota_norm,
            corr_Unassigned_norm,
            corr_Patescibacteria_norm,
            corr_Campilobacterota_norm]

x_tick_labels = ['Verrucomicrobiota',
                 'Firmicutes',
                 'Proteobacteria',
                 'Bacteroidota',
                 'Actinobacteriota',
                 'Fusobacteriota',
                 'Unassigned',
                 'Patescibacteria',
                 'Campilobacterota']
fig, ax = plt.subplots(figsize=(9, 3))
ax.boxplot(data, widths=0.8)
ax.set_title('Regenerated Figure 5D')
ax.set_yticks([0.2, 0.4, 0.6, 0.8, 1.0])
ax.set_xticklabels(x_tick_labels,
                    rotation=0, fontsize=6)
plt.show()
```

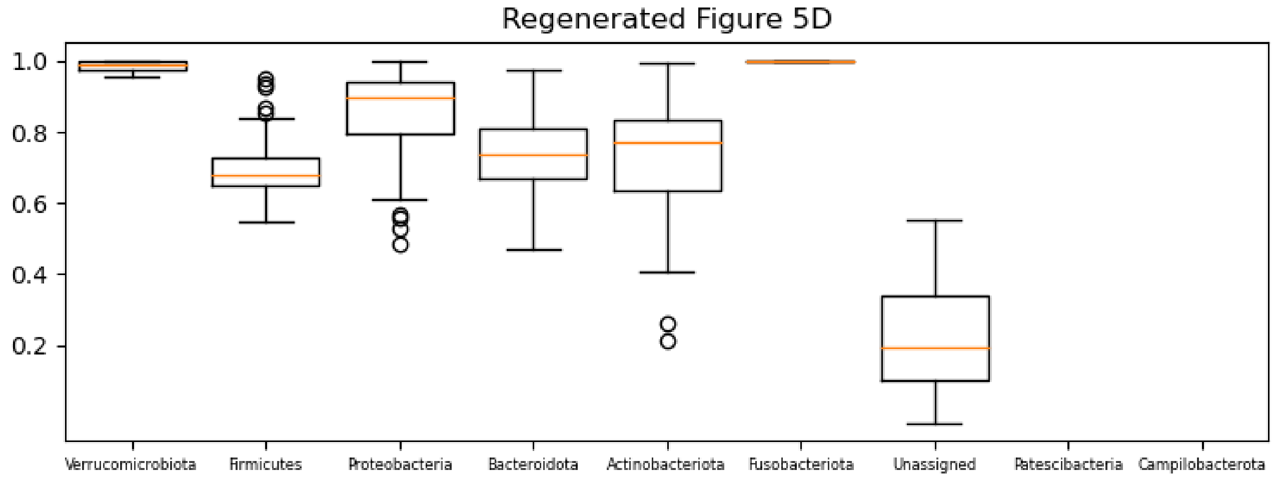


#### Step 9: Compare original and regenerated figures

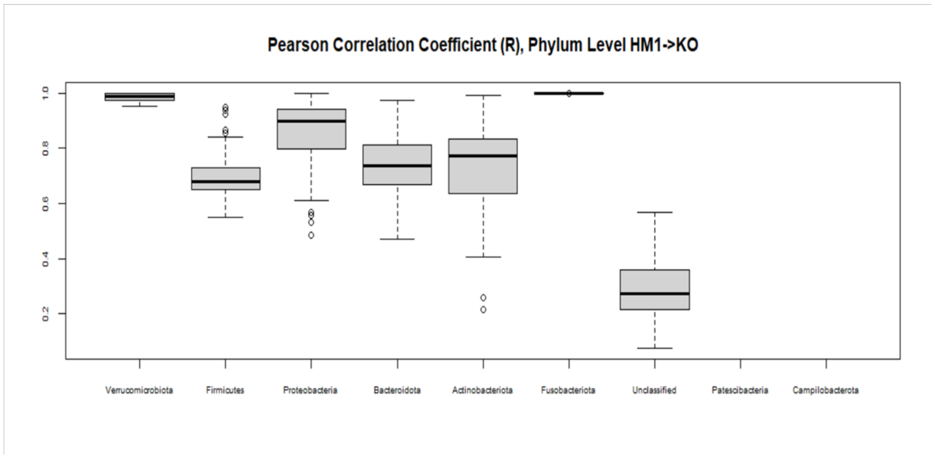


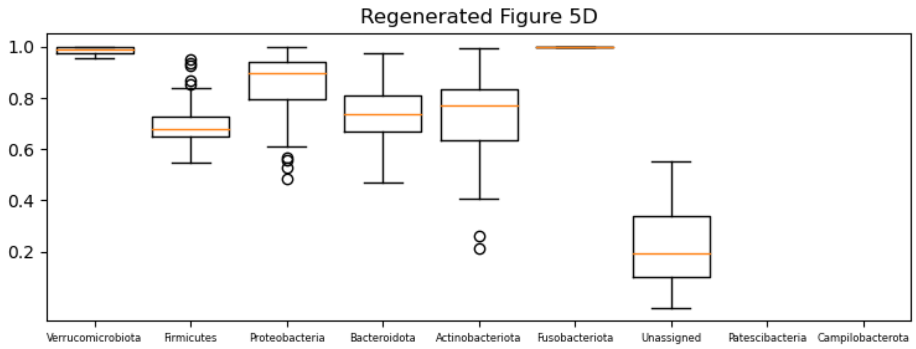

### Figure Validation Summary

#### Figure 1D Comparision (source code at figure_validation_1D.pdf)

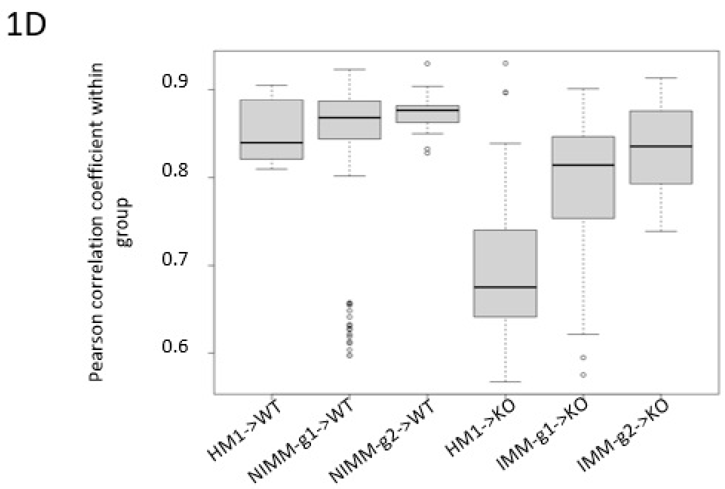


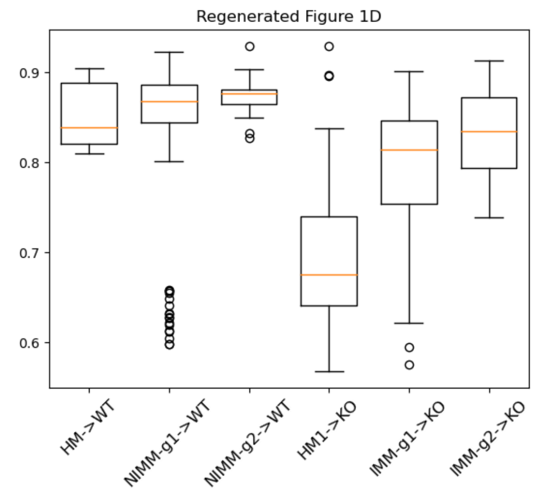


#### Figure 3A Comparision (source code at figure_validation_3A.pdf)

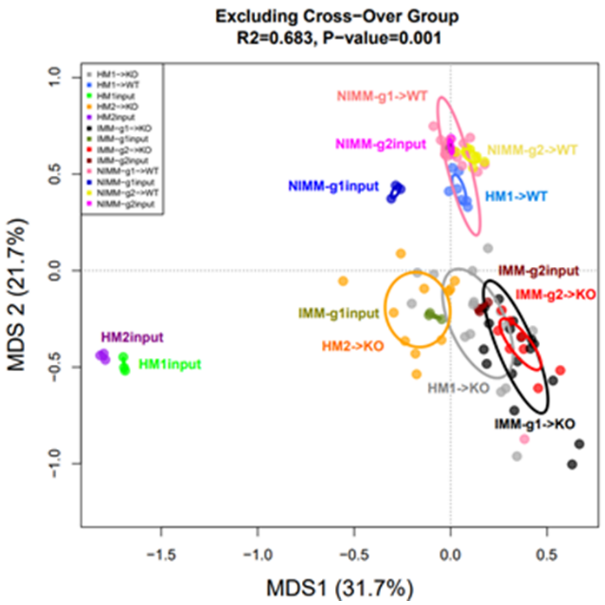


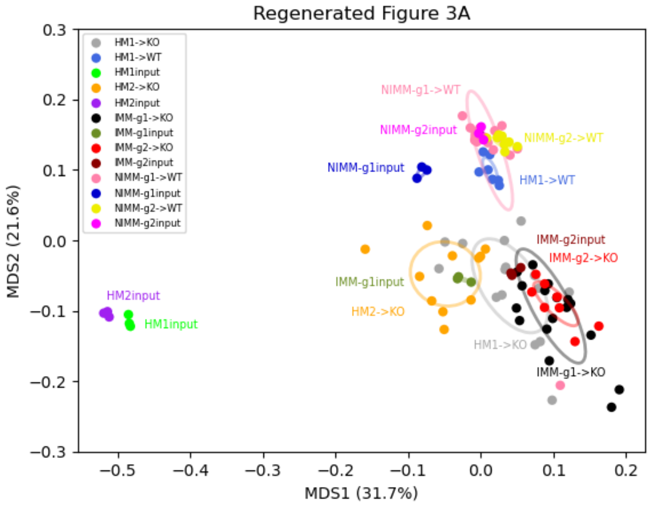


#### Figure 4B Comparision (source code at figure_validation_4B.pdf)

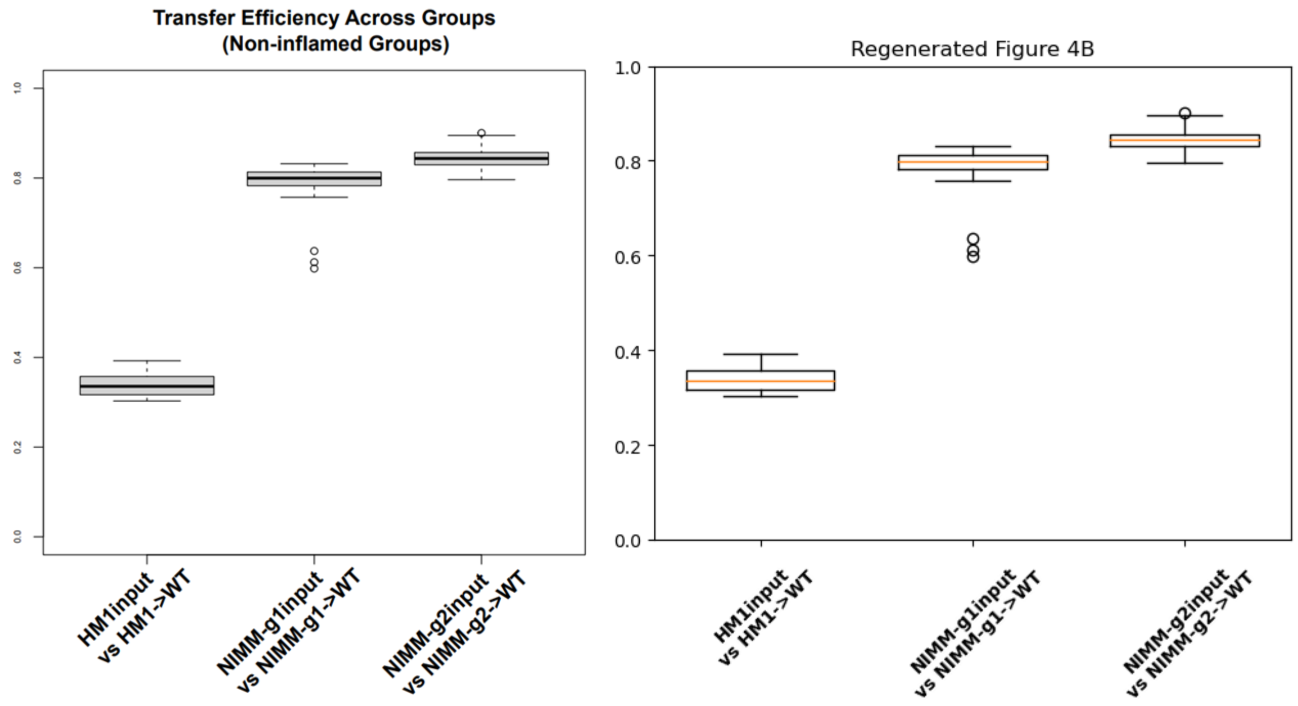


### Figure 4D Comparision (source code at figure_validation_4D.pdf)

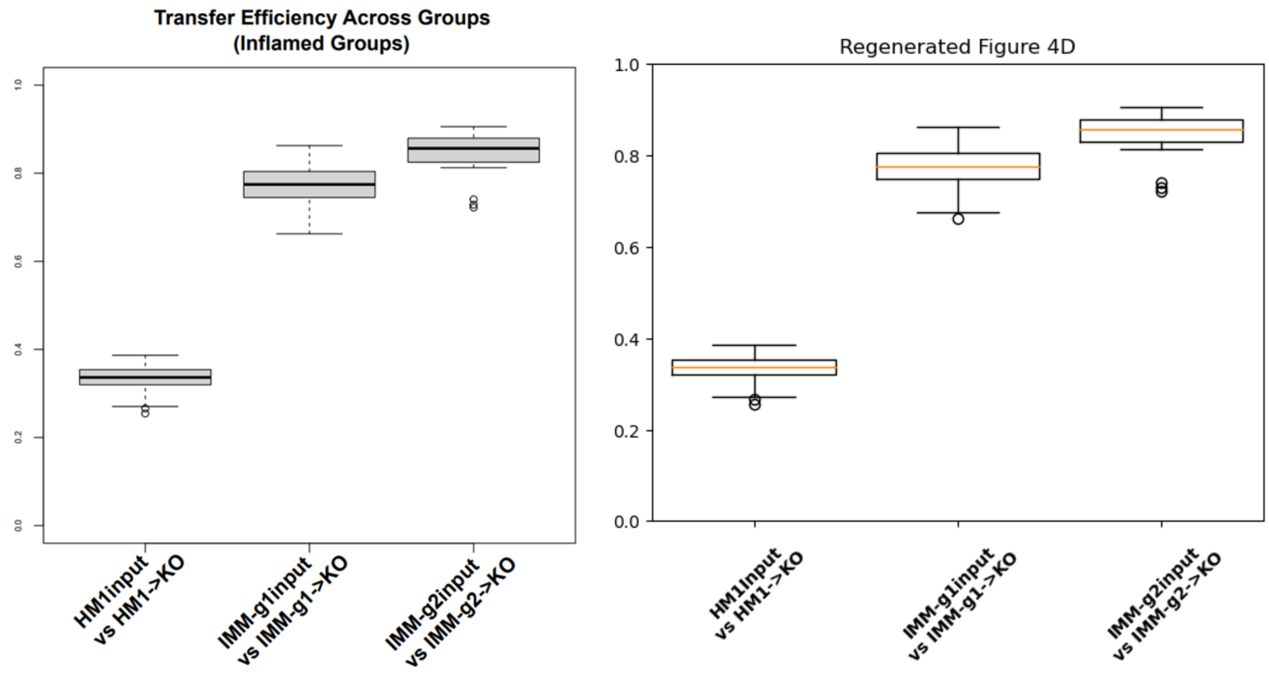


### Figure 5D Comparision (source code at figure_validation_5D.pdf)

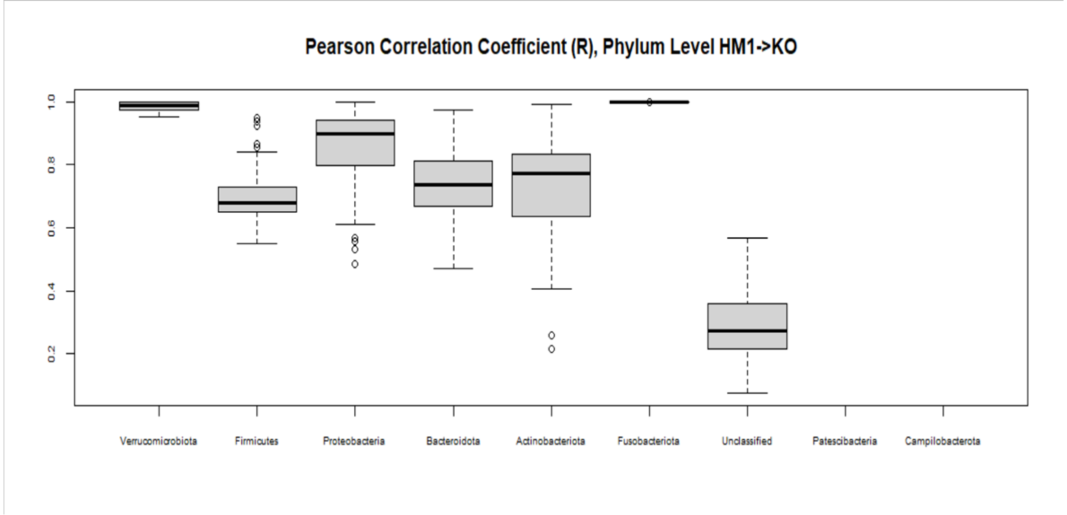

Pearson Correlation Coefficient (R), Phylum Level HM1->KO
1.0
0.8
0.6
0.4
0.2
Verrucomicrobiota
Firmicutes
Proteobacteria
Bacteroidota
Actinobacteriota
Fusobacteriota
Unclassified
Patescibacteria
Campilobacterota


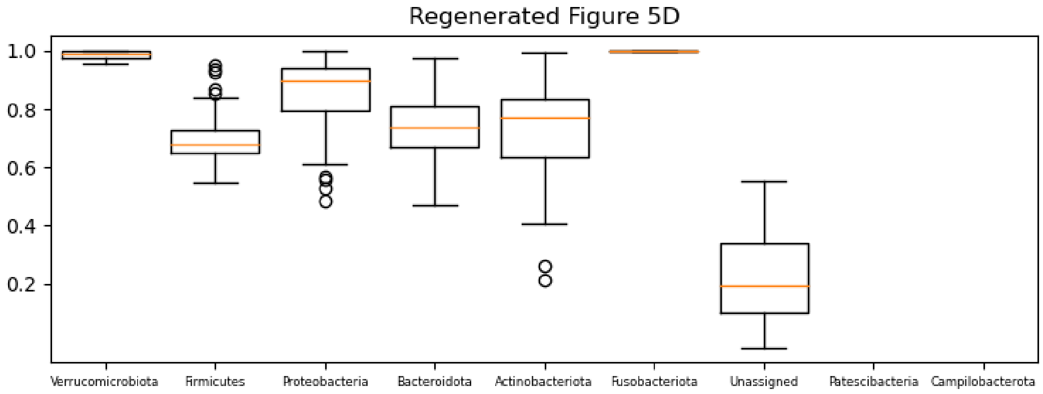

Regenerated Figure 5D
1.0
0.8
0.6
0.4
0.2
Verrucomicrobiota
Firmicutes
Proteobacteria
Bacteroidota
Actinobacteriota
Fusobacteriota
Unassigned
Patescibacteria
Campilobacterota
